# SMURF: soft-segmentation for single-cell reconstruction and topological analysis of spatial transcriptomic data

**DOI:** 10.1101/2025.05.28.656357

**Authors:** Juanru Guo, Simona Sarafinovska, Ryan A. Hagenson, Mark C. Valentine, David Y. Chen, William H. McCoy, Joseph D. Dougherty, Robi D. Mitra, Brian D. Muegge

## Abstract

High-resolution spatial transcriptomics requires new computational methods to accurately assign transcripts to individual cells. We developed SMURF (Segmentation and Manifold UnRolling Framework), a novel, cross-platform, soft-segmentation algorithm that maps mRNAs from barcoded capture spots to nearby nuclei. SMURF also “unrolls” complex tissue architectures by projecting cells onto Cartesian coordinates, enabling analysis of cell-type organization and gene-expression gradients in intact tissues. We benchmarked SMURF and found it assigns mRNAs to single cells more accurately than existing approaches. We evaluated SMURF’s ability to unroll complex tissues at cell-type resolution across multiple tissues and platforms. This analysis revealed previously unrecognized zonation of gene-expression programs, identified the transcription factors that regulate these patterns, and provided evidence that regional gene expression at the intestinal tip is reprogrammed by luminal environmental signals. Together, these results establish SMURF as a powerful framework for analyzing gene expression of cells within their native tissue contexts.

## Introduction

Biology is poised to address a fundamental challenge: describing the molecular features of all cell types in the body at single-cell resolution. However, a full understanding of how cells function cannot be achieved without considering how they exist within their microenvironments, as cells of the same subtype may have substantially different properties depending on their niche. Spatial transcriptomics (ST) addresses this challenge by allowing researchers to measure cellular gene expression while preserving spatial context.

First-generation capture-based ST platforms used relatively coarse, multi-cell capture spots. Emerging high-resolution methods are now approaching single-cell or even subcellular granularity. High-resolution capture-based ST platforms (e.g., Visium HD, Stereo-seq, Slide-seq/Slide-seqV2, Seq-Scope, PIXEL-seq, and DBiT-seq) dramatically increase spatial granularity but introduce new analytical challenges [1, 2, 3, 4, 5, 6, 7]. First, the capture elements (spots/pixels/beads/tiles) are not intrinsically co-registered to cell boundaries; many lie in extracellular space and therefore capture few or no transcripts. Second, transcripts from a single cell can be dispersed across adjacent capture elements, and conversely, individual capture elements near cell edges often contain transcripts from multiple neighboring cells. Most existing high-resolution segmentation/deconvolution methods implicitly assume one cell per capture element [8, 9], an assumption frequently violated at these resolutions. Thus, there is a need for new computational approaches that accurately segment and deconvolve mRNA in capture spots and assign them to single cells.

Another obstacle to fully harnessing the power of spatial transcriptomics is that most tissues exhibit complex three-dimensional architectures. Many organs—including the intestine, retina, and cerebral cortex—are composed of repeating or curved substructures that can be conceptualized as biological manifolds [10, 11]. Within these manifolds, cells are arranged along defined spatial axes that reflect essential biological gradients such as differentiation, signaling, and metabolic specialization [12, 13, 14]. However, current analytical tools are poorly suited to interrogate transcriptional organization along these non-Cartesian geometries. This challenge is compounded by tissue deformation and distortion during sample preparation, which obscures native spatial relationships. A computational framework capable of “unrolling” such complex tissue structures onto standard Cartesian coordinates would therefore enable quantitative analyses of regional gene-expression programs across diverse tissue substructures.

To address these challenges, we developed the Segmentation and Manifold UnRolling Framework (SMURF). SMURF introduces the concept of soft segmentation, in which mRNA counts within a capture spot can be assigned wholly to a single cell or fractionally distributed among multiple nearby cells in a manner that optimizes the cosine similarity of each cell to its corresponding gene-expression cluster. This approach increases the number of mRNAs assigned to individual cells and improves both clustering precision and transcript recovery, yielding state-of-the-art performance across diverse datasets. Beyond segmentation, SMURF provides a platform-independent framework—compatible with both sequencing- and imaging-based spatial transcriptomics—to computationally “unroll” complex tissue architectures using manifold theory, thereby revealing transcriptional organization along continuous biological axes that are otherwise hidden in conventional two-dimensional tissue sections.

We demonstrate the biological power of SMURF to uncover the spatial organization of gene-expression programs across diverse tissues and spatial transcriptomic platforms. Applying SMURF to sequencing-based datasets (e.g., Visium HD and Stereo-seq) and imaging-based datasets (e.g., CosMx, MERFISH, and Xenium Prime), we recovered fine-grained spatial structure and cell-type organization in brain, retina, skin, and intestinal tissues. Using Visium HD mouse intestine, SMURF resolved transcriptional programs along both the crypt-to-villus and proximal-to-distal axes at single-cell resolution, and identified the transcription factors (TFs) that regulate these programs. We revealed that environmental cues from the intestinal lumen are key determinants of regional gene-expression identity. Together, these analyses show that SMURF robustly segments high-resolution spatial data, sensitively detects rare cell types, and computationally unrolls cells onto Cartesian coordinates to enable manifold-level analysis of tissue architecture. By integrating manifold learning with soft segmentation in a platform-independent framework, SMURF establishes a general computational foundation for analyzing transcriptional organization in complex tissue geometries.

## Results

### Soft-segmentation for spatial transcriptomics

The capture elements, or “spots”, of high-resolution ST methods are not aligned with cellular boundaries. Some spots collect mRNA from multiple cells, and conversely, most cells have transcripts that are distributed across multiple spots. This makes it difficult to reliably assign transcripts to individual cells. SMURF overcomes this limitation using a novel three-step “soft segmentation” approach.

In the first step, SMURF identifies nuclei from H&E or DAPI/IF images using established segmentation methods [15, 16, 17, 18]. It then constructs a gene expression matrix by assigning mRNA counts from the capture spots that directly overlie each nucleus. Nuclei are then clustered by expression similarity using Scanpy [19, 20, 21]. Simultaneously, SMURF builds a nearest-neighbor network based on nuclear positions that will be used in the second step to assign surrounding spots to nearby nuclei (Figure 1a).

**Fig. 1:**
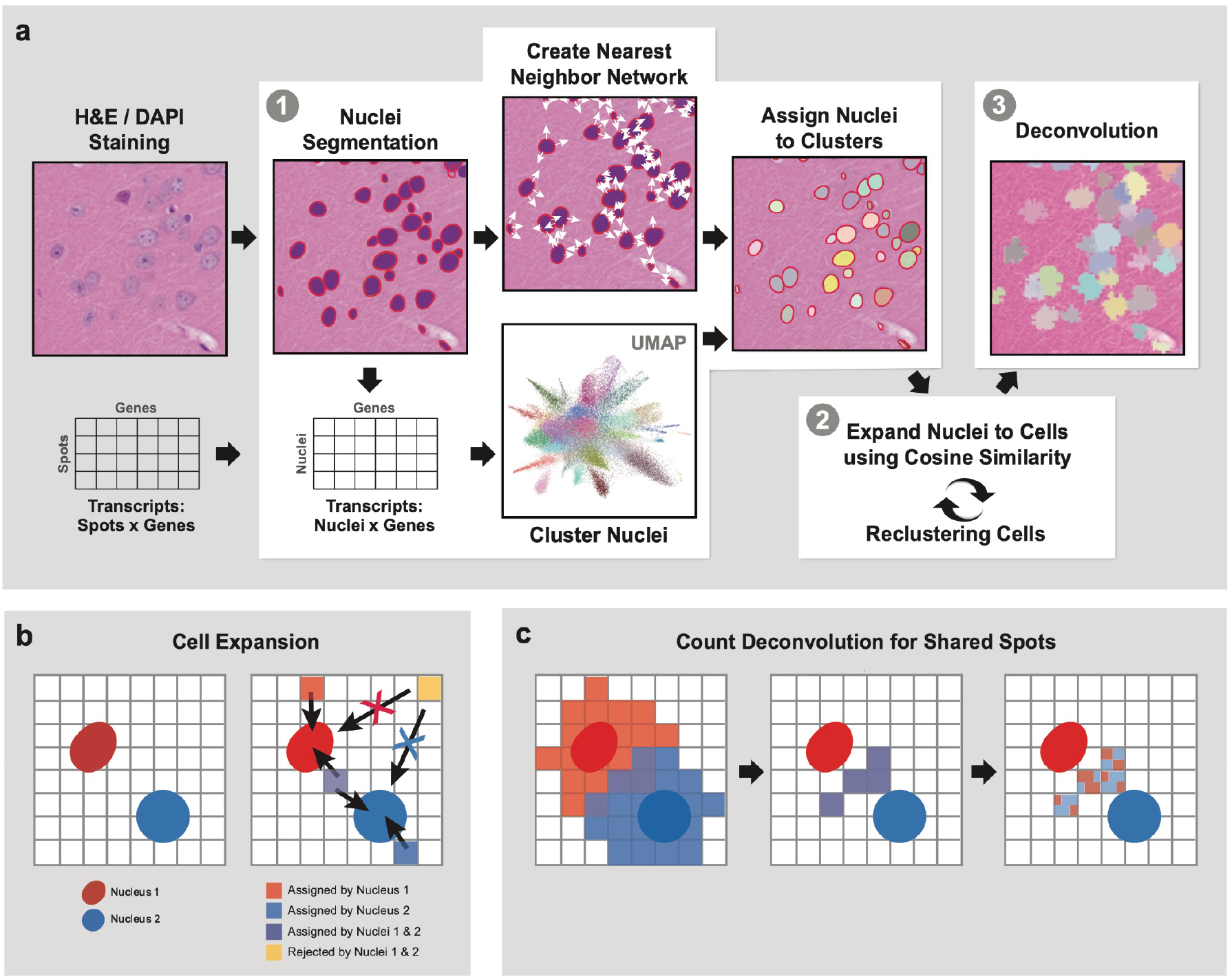
SMURF uses soft-segmentation to reconstruct single-cell transcriptomes. (a) Schematic of SMURF’s soft-segmentation algorithm. SMURF segments nuclei from the image data, assigns transcripts to nuclei from overlapping capture spots and clusters the resulting nuclei-by-genes matrix (Step 1). The algorithm then iteratively expands cell boundaries to assign transcripts from capture spots that do not overlap nuclei to cells (Step 2). Finally, SMURF leverages deep learning to deconvolve spots assigned to multiple cells during the expansion step (Step 3). (b) Example of capture spots assignment to cells. Capture spots can be assigned to zero, one, or multiple nuclei during cell expansion (Step 2), depending on their proximity and expression similarity. (c) Count deconvolution for shared spots. After the cell expansion step, capture spots may be shared between multiple cells (purple squares in the first two panels). Transcripts from shared capture spots are divided among cells using a quadratic programming or deep learning algorithm (see Methods).

Because nuclei occupy only a small fraction of total cell volume, the initial single-nucleus gene expression matrix is sparse, with low UMI counts [22]. Thus, the second step of soft-segmentation expands the nucleus-anchored gene expression by incrementally assigning nearby spots to nuclei through concentric expansion. Nearby spots are added only if they increase the cosine similarity between the expression profiles of the nucleus and its cluster. This process is then repeated iteratively – nucleus assignments and cluster identities are alternately updated until gene expression clustering stabilizes using normalized mutual information (NMI) [23]. At this step, SMURF allows flexible assignment: a spot can be linked to one nucleus, several nuclei, or none (Figure 1b).

The third and final step of SMURF’s soft-segmentation is count deconvolution for spots shared by multiple cells. SMURF deconvolves individual mRNA counts from capture spots assigned to multiple cells, using a combination of quadratic programming and deep learning [24, 25] (Methods, Figure 1c).

### SMURF improves single-cell reconstruction of high-resolution ST data

We benchmarked SMURF’s soft segmentation algorithm using Visium HD data from the mouse brain and distal small intestine (the ileum) [26, 27]. We compared SMURF to three commonly used methods (Figure 2a-b). The 8 *µ*m method, default from 10x Genomics, aggregates 16 2-*µ*m-spots into 8 *µ*m ×8 *µ*m “cells” to increase UMI counts for clustering and differential expression. It does not consider nuclear boundaries, often assigning transcripts to “cells” that lack nuclei, only partially cover a nucleus, or span multiple nuclei (Figure 2a-b, 2nd column). Bin2cell, a recently described method for single-cell segmentation, performs better than the 8 *µ*m in the mouse brain [9] (Figure 2a, 3rd column). However, Bin2cell often merges multiple cells or misses nuclei entirely in the densely packed small intestine (Figure 2b, 3rd column). The Nuclei segmentation method only uses spots directly overlapping nuclei; this results in a sparse expression matrix, as it excludes many transcripts from the surrounding cytoplasm (Figure 2a-b, 4th column). In contrast, SMURF achieves reliable single-cell resolution while accurately assigning cytoplasmic transcripts (Figure 2a-b, last column).

**Fig. 2:**
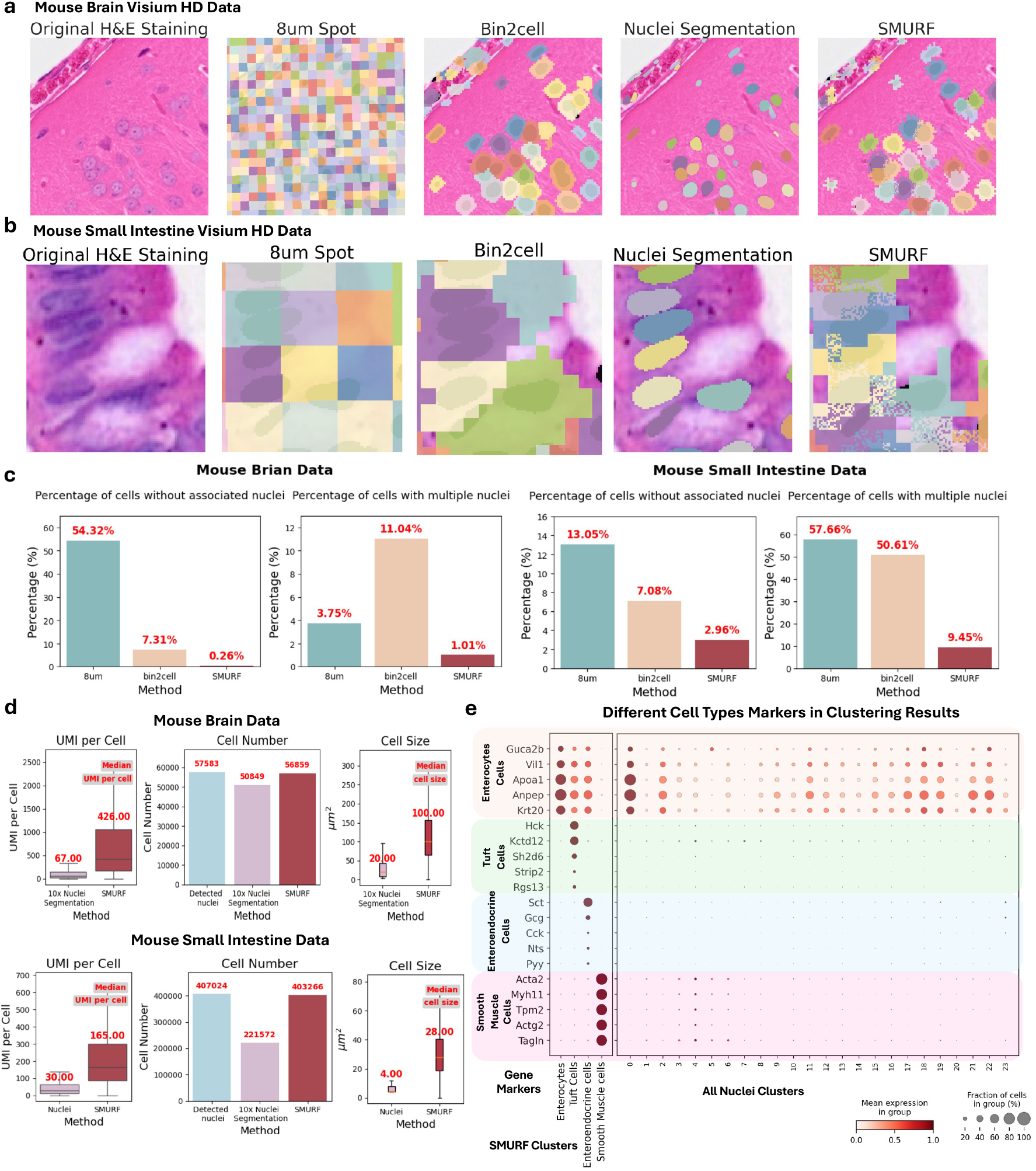
SMURF accurately assigns transcripts to cells in mouse brain and mouse small intestine data. (a,b) Representative images illustrating how SMURF and other ST segmentation methods aggregate Visium HD capture spots to define cells in the mouse brain (top row) and mouse small intestine (bottom row). Capture spots are shaded according to their assigned cell. SMURF uniquely manages to both assign only one nucleus per cell and analyze captures spots under cell bodies. (c) Percentage of cells without an associated nucleus or containing multiple nuclei in the mouse brain (left panel) and mouse small intestine (right panel) using different ST segmentation strategies. (d) Median number of UMIs assigned to cells, total cells detected, and median cell size using SMURF or nuclei-based segmentation. (e) Cell type detection in mouse small intestine using clusters generated by SMURF or nuclei-segmentation. Expression of marker genes for four intestinal cell types is plotted across clusters. SMURF but not nuclei segmentation has clusters that clearly correspond to each cell type. Abbreviations: UMI, Unique Molecular Identifier.

We next quantified segmentation accuracy by measuring the percentage of cells lacking a nucleus and the percentage containing multiple nuclei for each method (Figure 2c). SMURF was the most accurate in both brain and small intestine data. Only 0.26% of SMURF cells in the mouse brain lacked a nucleus, and only 1.01% contained multiple nuclei. This was significantly better than Bin2cell or the 8 *µ*m method. SMURF also outperformed these methods in the densely populated small intestine, with the lowest rates of both nucleus-less (2.96%) and multi-nucleus cells (9.45%, Figure 2c, right panels), demonstrating that SMURF more accurately assigns transcripts to cells at single-cell resolution than existing methods. We also found that SMURF’s cell-level normalization corrects a severe count imbalance artifact observed in transcript-sparse regions when using the 8 *µ*m method (Supplementary Notes, Extended Data Figure 1a). Compared to the nuclei segmentation approach, SMURF assigned 5-7 times more UMIs to each cell, and the average cell size was bigger (Figure 2d). Together, these results demonstrate that SMURF accurately assigns Visium HD counts to single cells, with substantially more counts per cell than nuclei segmentation alone.

We next evaluated whether SMURF’s soft segmentation generalizes across high-resolution, capture-based spatial transcriptomic platforms by applying it to Stereo-seq mouse brain and Seq-Scope mouse liver datasets. In both, SMURF assigned substantially more UMIs per cell than nuclei-based segmentation (Extended Data Figure 1c). To benchmark against another nuclei expansion approach, we compared SMURF soft segmentation to results obtained with the Spateo morphological expansion algorithm [2]. SMURF’s probabilistic “soft-expansion” produced higher correlations between nuclei masks and their corresponding expanded regions than morphological expansion (Extended Data Figure 1d). Together, these results demonstrate that SMURF effectively works across capture-based spatial transcriptomic datasets and provides more accurate transcript assignment than conventional morphological methods.

### SMURF improves cell type identification and capture of rare cell types

We hypothesized that SMURF’s higher transcript assignment per cell would lead to more accurate cell-type identification and improved estimation of their relative abundances. To test this, we used cell-type proportions from a probe-based ST study of mouse cortex as a reference standard [28]. Using Kullback–Leibler (KL) divergence [29], we quantified how closely SMURF and nuclei segmentation (without expansion) reproduced the probe-based ST-derived proportions. SMURF showed substantially lower KL divergence (0.123 vs. 0.738), indicating a more accurate representation of cell-type abundances (Extended Data Figure 1b).

Comparable ground-truth data for the small intestine are limited, so we clustered and annotated cell types independently for nuclei segmentation without expansion and SMURF. Both approaches detected enterocytes, the dominant epithelial population (Figure 2e). However, only SMURF reliably identified rarer epithelial cell types, such as tuft cells and enteroendocrine cells (EECs), as well as smooth muscle cells from the muscularis layer. These cell types were not captured by nuclei segmentation, potentially due to irregular nuclear morphology, low nuclear transcript abundance, or mRNA polarization—factors known to affect these cell populations.

Taken together, our results show that SMURF accurately assigns transcripts to individual cells with single-cell resolution, retaining transcripts from both the nucleus and cytoplasm. Compared to nuclei segmentation, SMURF recovers more transcripts per cell and improves the identification of known cell types in Visium HD data. SMURF’s increased count depth enhances statistical power for downstream analyses and enables more reliable detection of rare or morphologically irregular cell types.

### SMURF applies manifold detection to project ST data from complex tissue substructures onto Cartesian coordinates

Having assigned transcripts to individual cells, the next challenge is to reconstruct how these cells are spatially arranged within the intricate geometries of real tissues. Spatial relationships are critical for interpreting gene-expression patterns, but most tissues contain curved or layered substructures that do not align with standard Cartesian grids, hampering the analysis of cell organization or gene expression gradients in these substructures. To address this, we developed a manifold detection approach that projects cells from complex tissue shapes onto a novel coordinate system aligned with the structure of interest (Methods). Manifold detection is a mathematical concept that assigns local Euclidean coordinates to small areas of a complex surface. SMURF uses cell-resolved transcriptomes to identify tissue structures (e.g., cortical layers of the brain or the muscularis mucosa of the intestine) and defines a manifold through that structure. SMURF then “unrolls” this manifold, projecting the cells onto standard Cartesian axes. This functionality enables quantitative analyses of cellular organization and regional specification within a coordinate system most interpretable for each tissue.

We evaluated SMURF’s manifold detection by analyzing the mouse brain cortex. We first identified a one-dimensional manifold path representing the spatial distribution of cells in the cortical basement membrane using a locally linear embedding algorithm [30] (Figure 3a). We then extended the manifold to two dimensions to include all cortical cells, assigning each an (*x, y*) coordinates based on its position relative to the basement membrane (Figure 3b-c). This effectively projected the cortex onto a rectilinear coordinate system—”unrolling” the tissue for spatial analysis.

**Fig. 3:**
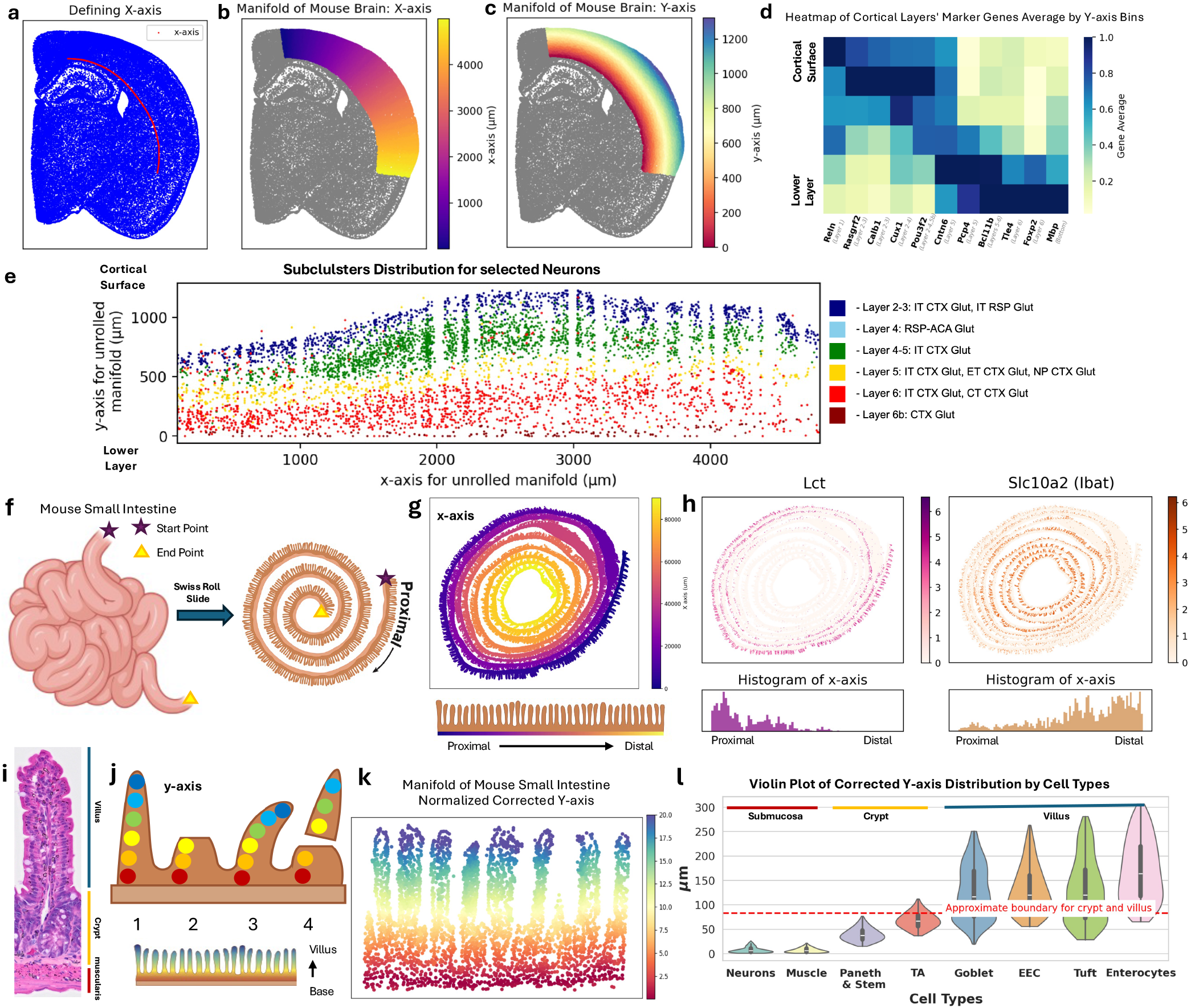
Manifold unrolling of Visium HD data in the cortex and small intestine. (a) Selection of the x-axis contour in the mouse cortex based on the positions of cells in the basement membrane cluster. (b) Overlay of cortical manifold onto mouse brain section, colored by x-axis (*µ*m). (c) Overlay of cortical manifold onto mouse brain section, colored by y-axis (*µ*m). (d) Heatmap of marker genes for cortical layers across different y-axis bins of the mouse brain manifold. (e) Distribution of cortical layer-specific neurons in the unrolled rectilinear coordinate system. (f) Schematic illustration of the mouse small intestine and its “Swiss roll” histological orientation. (g) Overlay of mouse small intestine data, colored by unrolled x-axis based on cells in the muscle layer (*µ*m). (h) Expression of selected proximal and distal enriched genes in the original ST image and histogram plot of those genes on the unrolled x-axis of the mouse small intestine. (i) Illustration of the crypt-villus axis in the mouse small intestine. (j) Schematic of four common villus structures encountered during construction of the y-axis for mouse small intestine data. (k) Overlay of intestinal manifold in selected villi with cells colored by normalized, corrected y-axis. (l) Cell type distribution along the corrected y-axis. Red dashed line indicates the approximate boundary between crypt and villus regions.

To illustrate how SMURF’s manifold algorithm can reveal regional patterning of cell types and gene expression, we divided the unrolled y-axis of the cortex into five bins spanning from the cortical base to the surface. We plotted the expression of known cortical layer markers across these bins as a heatmap [31] (Figure 3d). Layer-specific gene expression aligned precisely with SMURF-assigned locations: markers of layers 2–4 neurons, such as Cux1 and Pou3f2, were enriched nearer the cortical surface, while markers of layers 5 and 6, including Bcl11b and Foxp2, were expressed near the base. Next, we assigned every cortical cell in the ST data a cell-type identity based on its transcriptional profile, using the Allen Brain Cell Atlas as a reference [32, 33]. As expected, cell types stratified along the projected y-axis, with Layer 2 and 3 neurons at the cortical surface and Layer 6 neurons at the base (Figure 3e). Non-neuronal cell types also projected according to their known anatomic distributions [34, 35] (Extended Data Figure 2). Taken together, these results demonstrate that SMURF accurately “unrolls” a manifold of cells onto rectilinear coordinates while preserving patterns of gene expression and cellular organization.

### SMURF accurately transforms complex and discontinuous tissue onto rectilinear coordinates

We next applied SMURF to investigate regional transcriptional programs in the mouse small intestine, a tissue characterized by complex geometry and continuous spatial gradients. Gene expression, cell-type composition, and physiological function vary systematically along both the proximal–distal (longitudinal) and crypt–villus (radial) axes.

We analyzed Visium HD data from healthy mouse ileum (distal small intestine) prepared in a Swiss roll configuration [27] (Figure 3f). SMURF soft-segmented over 40,000 cells and applied its generalized unrolling algorithm to reconstruct the tissue’s native longitudinal layout (Figure 3g). Pseudobulked epithelial gene expression along the reconstructed x-axis correlated strongly with a recently published single-cell RNA-seq atlas of the mouse small intestine [36] (Extended Data Figure 3b). SMURF accurately localized canonical regional markers, including proximally enriched Lactase (Lct) and distally enriched Ileal bile acid transporter (Ibat) [36] (Figure 3h).

We next assigned cells to coordinates along the crypt-to-villus axis of the intestine. This dataset contained over 1,000 villi, presenting several analytical challenges (Figure 3i). Individual villi often bend from perpendicular alignment with the muscularis base, and some are partially severed or completely detached during sectioning. To correct these distortions, we developed an iterative method inspired by dynamic programming that identifies sequential layers of cells to reconstruct a corrected villus-specific y-axis representing a projected perpendicular alignment for each villus (Methods, Figure 3k, Extended Data Figure 3c, Extended Data Figure 4). SMURF accurately localized expected cell types along this reconstructed axis: neuronal and muscle cells in the muscularis propria underneath the epithelium, stem and Paneth cells at the crypt base, proliferative transit-amplifying (TA) cells above the stem cell zone, and differentiated epithelial populations along the villus (Figure 3i). Marker genes exhibited the anticipated spatial order, with Olfm4 marking stem cells in the crypt, Krt19 expressing TA cells above the crypt, Fabp6 enterocytes along the villus, and mature Ada expressing cells at the villus tip (Extended Data Figure 3d-e). Interestingly, some enteroendocrine (EEC) and tuft cells were detected near the base of the TA zone, suggesting occasional escape from the typical upward migration of differentiated cells. While retrograde migration has been reported for EECs, it has not previously been described for tuft cells [37, 38]. Collectively, these results show that SMURF precisely defines manifolds and “unrolls” complex and discontinuous tissue structures, projecting cells onto rectilinear coordinates that preserve in vivo spatial relationships, even in regions that are bent, torn, or partially missing.

### SMURF projects high-resolution ST data from multiple platforms and tissues

We next asked whether SMURF’s unrolling and projection algorithms generalize across different high-resolution spatial transcriptomics platforms. We first applied SMURF to another sequencing-based method, Stereo-seq, using data from the mouse olfactory bulb [2]. The olfactory bulb is organized concentrically into ring-like layers (Figure 4a). Using the authors’ provided cell segmentation, we anchored the SMURF projection on the outermost olfactory nerve layer and unwrapped the tissue into a two-dimensional coordinate space (Figure 4b). SMURF successfully resolved the expected radial progression of cell types from outermost to innermost, including the glomerular layer, mitral cell layer, and deep granule cell layer (Figure 4c). These results demonstrate that SMURF generalizes across other sequencing-based ST platforms and complex tissue geometries.

**Fig. 4:**
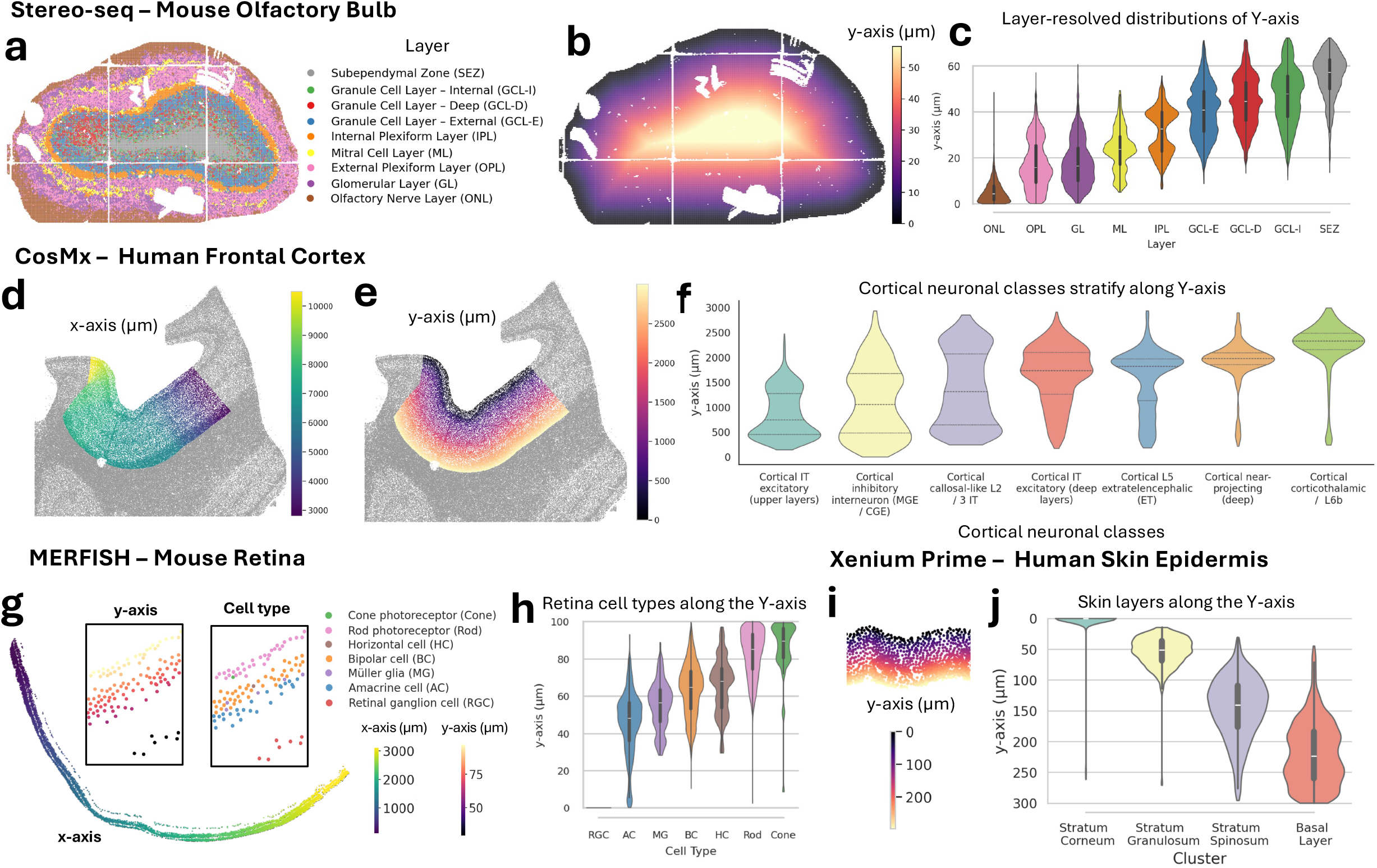
SMURF accurately projects high-resolution ST data from multiple platforms and tissues. (a–c) *Stereo-seq, mouse olfactory bulb*. (a) Tissue map with spots colored by annotated layers; (b) unrolled coordinate field showing y-axis manifold position across the olfactory bulb; (c) layer-resolved distributions of the unrolled y-axis. (d–f) *CosMx, human frontal cortex*. (d, e) Segmented cells with colors indicating the unrolled x-axis or y-axis coordinate (*µ*m); (f) distributions of cortical neuronal classes on the unrolled y-axis. (g–h) *MERFISH, mouse retina*. (g) Low-dimensional manifold of retinal cells colored by x-axis (left to right) and y-axis (inner to outer axes), with insets showing the same embedding labeled by major cell classes; (h) distributions of cell types on the unrolled y-axis. (i–j) *Xenium Prime, human skin epidermis*. (i) Segmented epidermal cells colored by the unrolled y-axis (*µ*m), using the outermost stratum corneum as the manifold base; (j) distributions of the unrolled y-axis across epidermal strata.

Sequencing-based technologies such as Visium HD and Stereo-seq capture and amplify mRNAs using spatially barcoded oligonucleotides that are mapped to histologic images to infer location [1, 6]. In contrast, imaging-based ST methods directly quantify gene expression in situ at subcellular resolution using labeled probes. To test SMURF’s performance on these datasets, we applied it to several imaging-based platforms, including CosMx, MERFISH, and 10x Xenium Prime [39, 40, 7]. We used the supplied cell segmentations and applied SMURF’s manifold unrolling algorithm to project each dataset into standardized Cartesian coordinates. Using CosMx data from the human frontal cortex, SMURF successfully recapitulated the laminar organization of cortical neurons [41]. The SMURF y-axis stratified excitatory and inhibitory neuronal subclasses in line with known cortical layers (Figure 4d-f). We next analyzed a MERFISH mouse retina dataset [42]. Anchoring SMURF’s manifold detection on the innermost retinal layer (ganglion cells) revealed the expected organization of cell types along the y-axis: ganglion and amacrine cells at the base, bipolar and horizontal cells in the inner nuclear layer, and photoreceptor rods and cones in the outermost layer (Figure 4g-h). Finally, we applied SMURF to a 10x Xenium Prime human skin melanoma dataset, focusing on the non-malignant epidermis [43]. We annotated epidermal keratinocyte subtypes using canonical marker genes [44]. SMURF unrolled and correctly ordered cells along the stratified epidermal axis from the stratum corneum at the surface, through the stratum granulosum and stratum spinosum, to the basal layer at the epidermal–dermal junction (Figure 4i-j). Extending this analysis, SMURF projected the tissue along an isodepth axis anchored at the basal layer, revealing distinct layer-specific organization near the surface, while deeper regions beyond the basal boundary lacked clear laminar structure (Extended Data Figure 5). These results demonstrate that SMURF can integrate data collected using a variety of ST platforms across a number of tissue types into a common coordinate space, enabling the quantitative analysis of cell organization and gene-expression gradients in complex tissue architectures.

### SMURF identifies the zonation of transcriptional programs in intestinal development

To demonstrate the potential of SMURF to discover new biological insights, we returned to the intestinal data. Stem cells at the crypt base continuously give rise to rapidly dividing TA cells, which differentiate into specialized digestive and barrier functions as they migrate up the villus [45] (Figure 5a). We hypothesized that SMURF’s projection of complex tissue structures onto rectilinear coordinates could be used to quantitatively examine the relationship between intestinal epithelial cells’ developmental state and their spatial position on the crypt-villus axis. We focused on the developmental trajectories of the most abundant epithelial cell type, enterocytes, from stem and TA cells. Pseudotime trajectories were measured using diffusion maps and visualized with partition-based graph abstraction (PAGA) and force-directed graph layouts [46, 47, 48] (Figure 5b). Pseudotime trajectory captured the expected transition from stem and Paneth cells to mature villus cells (left and middle panels). SMURF-assigned y-axis position followed a similar distribution (right panel), and was significantly correlated with pseudotime (*corr* = 0.80, p-value *<* 10^−10^; Extended Data Figure 6a). These findings demonstrate SMURF can recapitulate the spatial-temporal coupling of cell differentiation along the crypt-villus axis solely from spatial location.

**Fig. 5:**
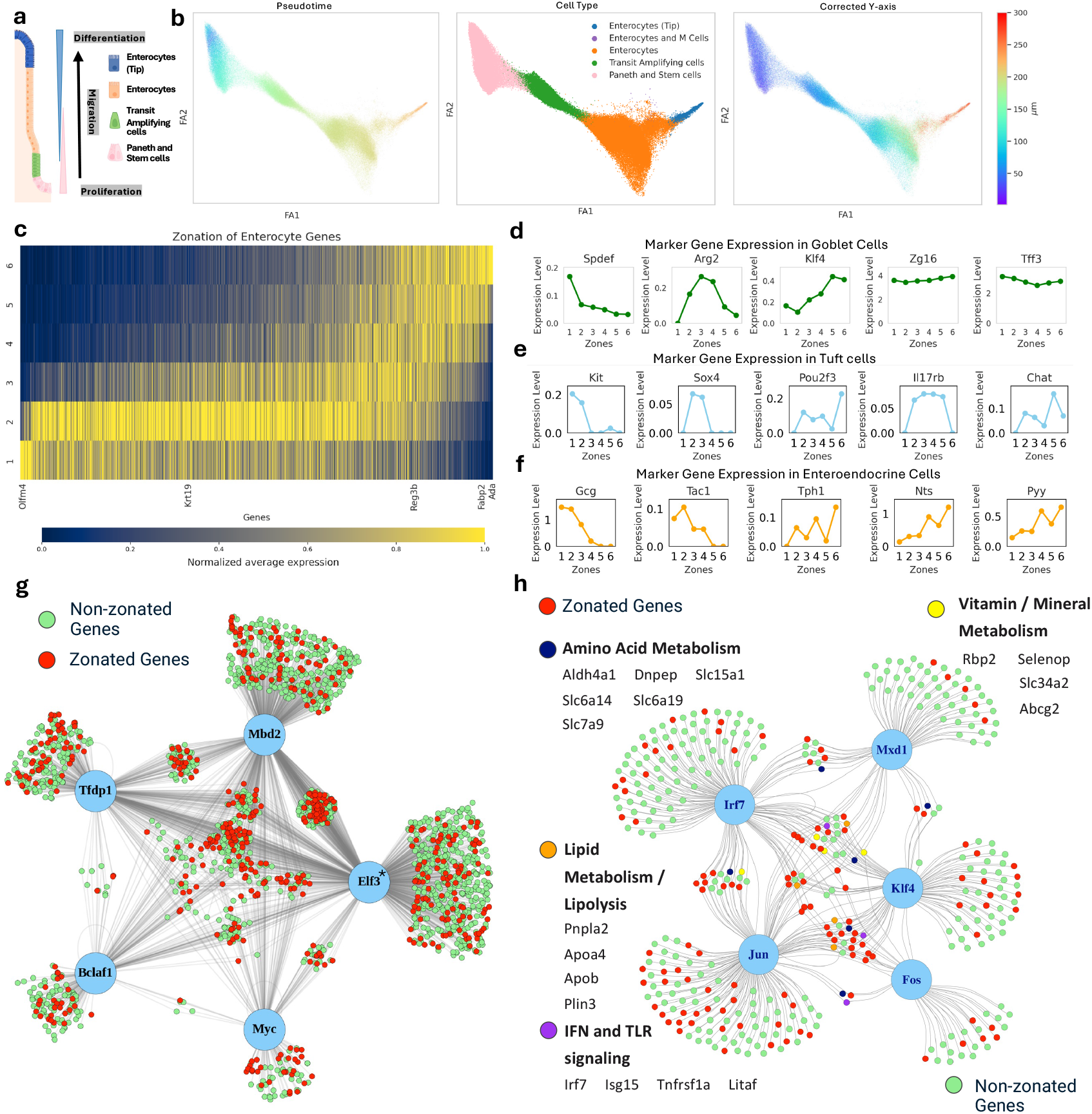
Spatial, temporal, and transcription-factor program organization along the ileal crypt–villus axis. (a) Schematic illustrating the differentiation trajectory of the epithelial lineage in intestinal villi from proliferative stem cells (bottom) to differentiated enterocytes (top). (b) Concordance of pseudotime trajectory and corrected y-axis position during enterocyte maturation. PAGA embedding of intestinal stem and Paneth cells, TA, and enterocytes, colored by pseudotime position (left panel), cell type (center panel), and corrected y-axis position (right panel). (c) Heatmap of normalized gene expression across zones. Gene expression in all stem and Paneth cells, TA, and enterocytes is included. The heatmap displays normalized expression of each gene (columns) in each zone (rows). (d-f) Zonation profiles of selected genes in goblet cells (d); tuft cells (e); and EECs (f). (g) Integrated TF target networks for Zone 2. TFs are depicted in blue circles. Lines connect TFs to their gene targets. Selected target genes are color-coded by their predominant functional category. *: 500 of the 1032 Elf3 targets are shown for legibility. (h) Integrated TF target networks for Zone 6. Coloring as in panel g.

We next sought to identify the gene programs that change along the crypt-villus axis during epithelial cell maturation in the ileum. We binned villi into 6 zones spanning from the crypt to the villus tip based on the corrected y-axis positions (Extended Data Figure 6b). We pseudobulked gene expression from the stem/Paneth, TA, and enterocyte clusters for each zone. Normalized gene expression was highly zonated across the villi (Figure 5c). We identified 2,997 genes that were significantly zonated by differential expression analysis (Methods, Supplementary Table 1). We further refined this list to a set of “landmark” genes highly specific for each zone by algorithmic and manual curation (Supplementary Notes, Extended Data Figure 7). KEGG pathway enrichment analysis revealed specialized transcriptional programs in each zone, including WNT signaling and proliferation in the crypts, with emerging capacities for carbohydrate, protein, and lipid absorption as enterocytes matured along the villus (Extended Data Figure 6c). Recent reports in the mouse jejunum and human duodenum showed functional specialization along the crypt-villus, but the genes and pathways in the ileum are distinct, highlighting the regionally specialized nature of intestinal metabolism [49, 50].

We next turned our attention to crypt-to-villus zonation in less abundant secretory epithelial cell types. Goblet cells, tuft cells, and the heterogeneous EECs are individually scattered throughout the villus and cannot be isolated by LCM or conventional spatial analysis with large bin sizes. Using the same six zones, SMURF analysis identified significant zonation along the crypt-to-villus axis for each secretory cell type (Figure 5d-f). In contrast to a recent study of the human small intestine that showed tuft cell markers zonated towards the bottom of the villus [50], the markers were variably expressed along the crypt-villus axis in mice. The broad tuft marker Kit was enriched at the crypt base. Sox4 is known to initiate the Tuft cell program, including the lineage-specifying TF Pou2f3. SMURF projections recovered this relationship with Sox4 at the base and Pou2f3 diffusely expressed (Figure 5e) [51]. Immune and sensory receptors, including Il17rb and Chat, had variable expression along the villus axis, in line with recent reports of spatial segregation of Tuft cell types in the intestine [52, 53, 54, 55] (Figure 5e). Similarly, EEC hormone expression was highly zonated. EECs in crypts were expressed Substance P (Tac1) and GLP-1 (Gcg), while EEC expressing Serotonin (Tph1), Neurotensin (Nts), and Peptide YY (Pyy) were only detected in villus zones, consistent with prior immunohistochemical studies [56, 57] (Figure 5f).

This is the first detailed description of ileal transcriptional zonation using ST data and provides an important resource of landmark genes in this organ. The functional progression of metabolic functions in enterocytes is concordant with prior studies in the jejunum [49], but our ST methods additionally resolve zonation in multiple rare cell types.

### Zonated transcription factor regulatory networks along the villus

The network of transcription factors regulating zonated gene expression in the mature villus remains poorly defined. Using SCENIC, we identified active TFs and their targets in each zone based on expression profiles from stem/Paneth, TA, and enterocyte cells [58]. We detected 118 TFs with average activity scores in at least one of the six zones (≥ 0.05, Supplementary Notes Table S3).

Cooperative TF binding is a well-established mechanism for regulating gene expression; we therefore mapped TF interaction networks within each zone. In Zone 2, a highly interconnected network of 5 TFs emerged, anchored by Elf3 (Figure 5g). Elf3 mediates Wnt-independent *β*-catenin activation in colon cancer [59, 60]. Other members of this network include Myc and Tfdp1, which are known to regulate proliferation and cell-cycle progression in the intestine, as well as Bclaf1, and Mbd2, which are not as well studied in intestinal zonation [61, 62, 63]. Genes with enriched expression in Zone 2 were significantly more likely to be targets of multiple TFs in this network (p-value = 1.56 × 10^−7^, Methods, Figure 5g). Genes co-targeted by two or more TFs were enriched for RNA biogenesis and mitochondrial metabolism, core TA cell functions. Thus, we identified a novel cooperative TF network controlling TA cell proliferation.

At the villus tip, we identified a distinct TF network, including AP-1 complex members Jun and Fos, previously linked to tip identity, and novel TFs Irf7, Mxd1, and Klf4 [50, 49, 64, 65]. As in Zone 2, tip-zonated genes in Zone 6 were also over-represented among those targeted by multiple TFs (p-value *<* 10^−10^, Methods, Figure 5h). These genes are involved in amino acid, lipid, and vitamin metabolism, as well as host defense, suggesting that AP-1 complex and environmental sensors such as Irf7 cooperate to drive tip-specific functions.

### Regionally specific gene programs accumulate in the epithelial villus tip

The intestine is functionally specialized along its proximal-distal axis, yet the molecular mechanisms that maintain regional identity remain unclear. To address this, we divided villi from the proximal and distal ileum by the x-axis and performed differential expression analysis on pseudobulked counts from each villus zone. Surprisingly, more genes were differentially expressed in upper villus zones than at the base [36, 11, 66] (Figure 6a). Our ST data are derived from the mouse ileum. To test the idea that villus bottom zonation is preserved across regions while villus top zonation is regionally specific, we plotted markers of jejunal crypt-villus zonation identified previously by LCM [49] (Extended Data Figure 8a-b). The genes identified as bottom markers in the jejunum were also expressed predominantly in Zones 2 and 3 of the ileum. In contrast, the jejunal villus top markers were not segregated at the top of the ileum, supporting the idea that villus tips have regionally specific transcriptional programs. Proximal ileal villus tips upregulated genes involved in typical proximal metabolic pathways, including lipid transporters Apoa4 and Apoc3, consistent with the proximal ileum’s role in absorbing excess dietary fats [67] (Figure 6b). In contrast, distal ileal tips are enriched for known distal ileum genes, such as the beta-alanine transporter Slc6a14, along with genes involved in bile acid absorption and steroid biosynthesis [68] (Figure 6c).

**Fig. 6:**
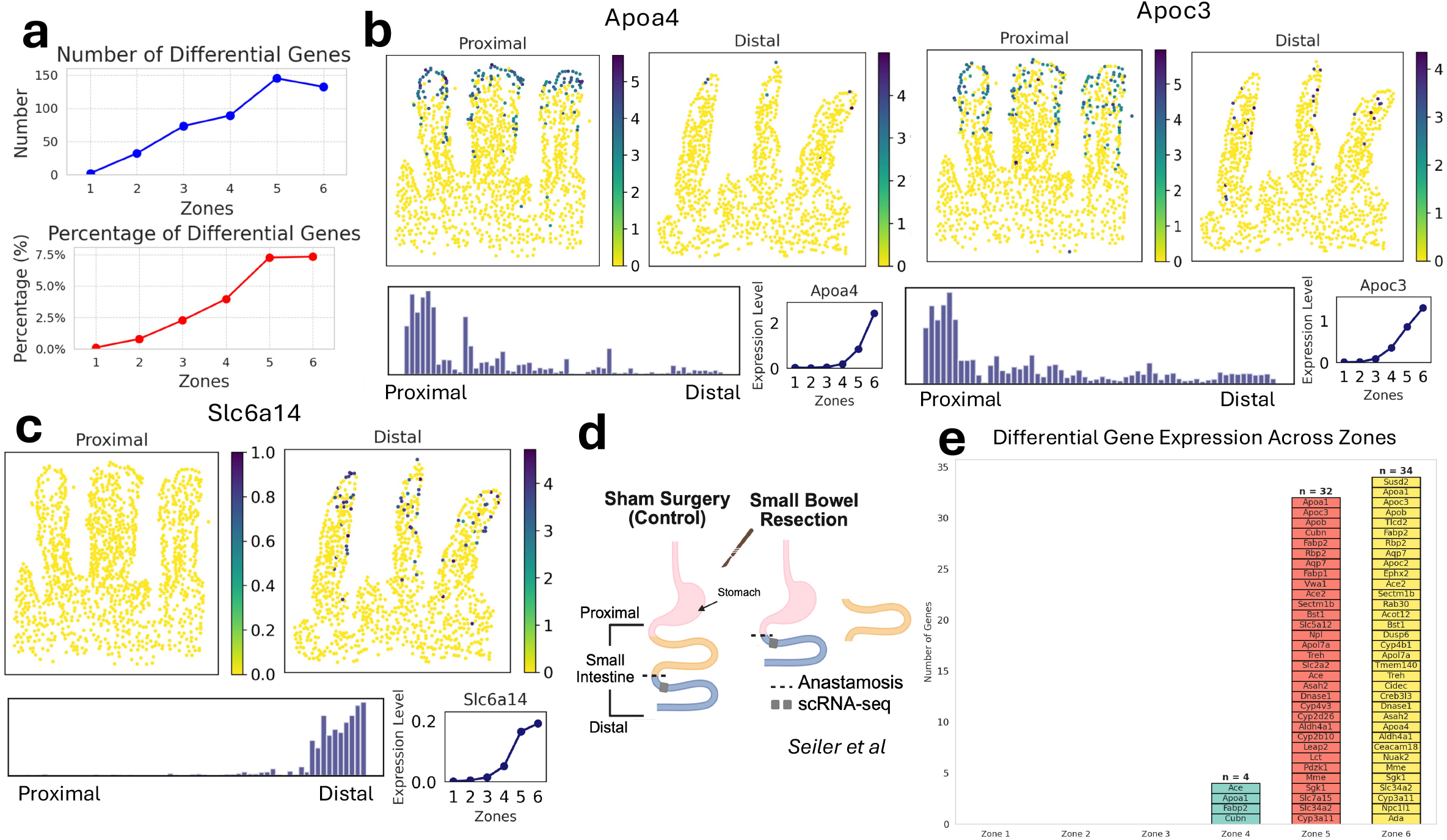
Regionally specific gene programs along the proximal-distal intestinal axis accumulate in the villus tip. (a) Number (top) and fraction (bottom) of genes that are differentially expressed in each zone, comparing proximal and distal regions of villi. Only genes expressed in more than 30 cells are considered. (b) Expression of proximal-tip enriched genes (*Apoa4* and *Apoc3* ). Data shows expression in the original ST data, as well as expression along the unrolled x and y axes. (c) Expression of distal-tip enriched gene (*Slc6a14*. Data shows expression in the original ST data, as well as expression along the unrolled x and y axes. (d) Schematic of small bowel resection and scRNA-seq dataset (Seiler et al [71]). (e) Tip cells, but not cells at the base of crypts, are reprogrammed one week after small bowel resection. For each zone, the number and identity of differentially expressed genes from the single-cell data are shown.

These data suggest that proximal–distal cellular identity is not dictated solely by stem cell-intrinsic transcriptional programs. Although intestinal stem cells encode much of their progeny’s regional identity, recent evidence challenges this view [36, 69, 70]. We therefore hypothesized that luminal gradients of environmental signals, including dietary components, xenobiotics, and microbes, interact with intestinal TF networks to establish regional identity.

To test this, we examined a small bowel resection (SBR) model, in which 50% of the small intestine is surgically removed and the remaining segments are joined, exposing the distal ileum to proximal luminal contents [71] (Figure 6d). Our hypothesis predicts that this exposure would induce proximal transcriptional programs at distal villi tips. We re-analyzed previously reported scRNA-seq from SBR ileal tissue and sham-operated controls, identified differentially expressed genes, and mapped these genes to spatial zones in healthy ileum using our ST dataset. As predicted, genes upregulated after resection were significantly enriched in the tip zonated genes (empirical chi-square test, p-value = 9.99 × 10^−6^), Figure 6e). None of the induced genes were expressed in stem and TA cells, indicating that most transcriptional changes occur downstream of any stem cell reprogramming. As reported, the SBR upregulated genes were enriched for proximal lipid metabolism pathways [71]. Together, these results support our hypothesis that environmental cues, rather than stem cell-intrinsic programs alone, shape regional identity. However, we cannot exclude additional effects from epithelial proliferation or hormonal signaling triggered by surgery.

## Discussion

The emergence of high-resolution, capture-based ST platforms has created an urgent need for algorithms that can accurately assign transcripts in densely tiled capture spots to individual cells. Here, we showed that SMURF’s novel “soft-segmentation” method more accurately and comprehensively assigns mRNAs to cells, segments cells with a higher accuracy, and more reliably detects some rare cell types than existing methods. SMURF also “unrolls” tissue substructures, even when parts of them are damaged or missing, to place cells on Cartesian coordinates, allowing for quantitative analysis of the *in vivo* spatial organization of cell types and transcriptional programs. SMURF’s unrolling functionality is platform-agnostic; we successfully unrolled and analyzed cellular gene expression from datasets collected using Visium HD, Stereo-seq, CosMx, MERFISH, and Xenium Prime platforms.

We applied SMURF to analyze ST data from the small intestine and gained new insights into the spatial organization of this organ. Prior work in this area has revealed important principles, but was constrained by coarse resolution or the use of labor-intensive methods [49, 72, 50, 73, 74]. Using SMURF, we identified cell-type-specific zonated gene expression programs and novel TF regulatory networks along the crypt-villus axis, many of which were not previously identified. We also used SMURF to investigate a fundamental question about small intestine development: are proximal-to-distal gradients in gene expression predetermined or established in response to environmental signals? We found that regional gene expression differences are largely restricted to villus tip cells and can be reprogrammed by relocating the distal intestine to a proximal environment, demonstrating that proximal–distal patterning in adult tissue is shaped more by environmental cues than by intrinsic stem cell programming.

We note that the current version of SMURF may require optimization for some applications. When assigning transcripts from capture spots near cell boundaries, SMURF cannot always resolve cells in Visium HD datasets that share a high degree of expression similarity and are closely adjacent, such as the stem and Paneth cells in the intestinal crypt. Although SMURF still outperforms other methods in this regard, imaging-based methods with inherent sub-cellular resolution followed by SMURF analysis may be preferred in these instances. Additionally, ST datasets, including associated histology images, can be extremely large, making the analysis computationally demanding. SMURF’s full implementation employs a deep-learning algorithm for deconvolution, which requires GPUs to run efficiently [75]. To make SMURF more widely accessible, we have developed a “lite” version of SMURF, which can run on most workstations (Supplementary Notes).

As ST continues to evolve, new platforms will employ even higher resolution capture spots (for capture methods), analyze more genes (for imaging-based methods), and profile larger numbers of cells. The full potential of these technologies will only be achieved with approaches like SMURF that consider all capture spots, have the ability to assign mRNAs in the same capture spot to different cells, and can unroll tissues in a platform-agnostic manner. SMURF’s segmentation method is inherently extensible to emerging technologies for three-dimensional tissue ST. SMURF’s segmentation and unrolling approach complements existing methods to study signaling gradients in ST [76, 77, 78]. Finally, SMURF’s unrolling algorithm could be extended to protein- or metabolite-based spatial technologies.

Our study highlights SMURF’s utility to uncover principles of tissue organization during normal development. An important future direction will be to deploy this approach to study disease models at single-cell resolution. Many diseases disrupt normal tissue structures, such as the loss of normal crypt-villus architecture in inflammatory bowel disease, or recruit novel cell types, such as vascularization in cancer. SMURF offers a framework to dissect how multiple cell types interact in these contexts, including “4D” studies of evolving responses over time. We have demonstrated SMURF’s utility in multiple tissues (Supplementary Notes) and anticipate that SMURF will provide insight into other spatially complex organs such as the kidney, retina, and lung.

## Methods

### Soft Segmentation Analysis

#### Introduction

In this section, we provide a detailed description of the soft-segmentation method implemented in SMURF. Cells are denoted by the index *i*, genes by *j*, and spots by *k*. Since cells are typically associated with exactly one nucleus, we denote both a cell and its corresponding nucleus by *i*, emphasizing their one-to-one correspondence. The inputs to the soft segmentation algorithm consist of: (1) the 2 *µ*m spot gene expression matrix *S*; (2) the H&E or DAPI/IF tissue image *Img*^*o*^; and (3) the “spot to pixel” data frame.

We define the 2 *µ*m spot gene expression matrix *S* to have dimensions *K*_spots_ × *J*_genes_, where *K*_spots_ represents the number of spots and *J*_genes_ the number of genes, with each element *S*_*kj*_ ≥ 0. Each pixel in the original image *Img* ^*o*^ represents a grayscale value (0–255) for DAPI/IF images or an RGB array for H&E stained images. The matrix *S* is linked to the processed image *Img* via the “spot to pixel” data frame. Note that some pixels in *Img*^*o*^ are not covered by spots, and each spot encompasses multiple pixels; hence, we focus on the area covered by spots, defined as *Img*. We denote by *Img*_*k*_ the set of pixels corresponding to spot *k*.

SMURF requires users to generate their own nuclei segmentation results. To facilitate this, we provide tutorials for DAPI/IF images (using Cellpose [15, 79]) and H&E-stained images (using StarDist [16, 17, 18]); users may also choose their preferred segmentation tools. The final nuclei segmentation output, denoted as *Nuclei_img*, is a two-dimensional matrix with the same dimensions as *Img*. In *Nuclei_img*, regions without nuclei are assigned a value of 0, whereas regions containing nuclei receive unique numerical identifiers.

Given the substantial volume of data and resulting outputs, SMURF defines a new Python object, the Spatial Object (*SO*), initially constructed from the three inputs. The *SO* records the nuclei segmentation results, detailed information for each cell, and visualization outputs. For clarity, we refer to the components of *SO* by their object names rather than describing each individual element of *SO* here.

The soft-segmentation in SMURF involves three main steps: nuclei analysis, cell expansion, and count deconvolution for shared spots. For the first step, a nucleus-by-gene matrix is constructed and clustered to provide initial cell type annotations, and a nearest neighbor network is constructed to estimate approximate cell size; second, cells are expanded by iteratively assigning adjacent spots based on their similarity to the cell’s cluster identity, with periodic reclustering to refine assignments; finally, overlapping spots that are shared by multiple cells are resolved through optimization methods (quadratic programming and deep learning), ensuring accurate gene expression profiles for each cell. Finally, SMURF outputs a spatial single-cell <monospace>AnnData </monospace>object containing the gene expression matrix *G* (*I*_cells_ × *J*_genes_). Each cell is annotated with its gene expression profile, spatial coordinates, preliminary cluster assignment, and the cosine similarity to its assigned cluster.

#### Step one: Nuclei Analysis

The nuclei analysis includes mainly three parts: building the nuclei gene expression array, clustering nuclei, and estimating the potential sizes of cells based on location. It takes in the 2*µ*m spot gene expression matrix *S* and nuclei segmentation results *Nuclei_img* from the image.

Before the analysis, we first filter the 2 *µ*m spot gene expression matrix *S* based on the criterion

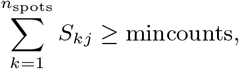

with the default threshold set at mincounts = 1000. We denote the resulting filtered matrix as *S*^*′*^.

The `Nuclei_img` contains pixel-level data, and each 2 *µ*m spot corresponds to approximately 50 pixels. Consequently, many nuclei overlap multiple capture spots and, conversely, some capture spots are shared by multiple nuclei. We define *pct*_*ik*_ = (number of pixels in nucleus *i* of spot *k*)*/Img*_*k*_, with 0 ≤ *pct*_*ik*_ ≤ 1, to denote the percentage of pixels in spot *k* that overlap the nucleus of cell *i*. For each nucleus *i*, we record all spot IDs for which the fraction of overlapping pixels is at least a specified minimum (the default fraction is 1*/*6). The transcripts captured in all spots that meet this criterion for nucleus *i* are assigned to that nucleus. Using this initial assignment of transcripts to cells, we create the gene count matrix *X*^(0)^, defined as:

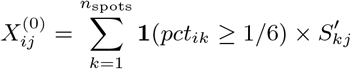

where *X*^(0)^ is an *I*_cells_ × *J*_genes_ filtered_ matrix and 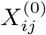 indicates the count expression of gene *j* in nucleus *i*.

In the next part, we filter out nuclei with few transcripts assigned to them, since extremely low counts could be due to errors like waterdrops on the capture surface and influence the accuracy of further analysis. We then analyze matrix *X*^(0)^ using the standard Scanpy [19, 20] workflow, including filtering out genes that are not expressed in some minimum number of cells, count normalization, dimensionality reduction via principal component analysis (PCA), and Leiden clustering. Through this approach, we obtain initial clustering results *C*^(0)^ based on the *X*^(0)^, where *C*^(0)^ is a 1 * *n*_*cells*_ array that assigns each nucleus into one of *n*^(0)^ clusters based on gene expression profile. These initial cluster assignments are based solely on transcripts assigned to nuclei from overlapping capture spots.

Finally, we estimate the potential number of spots belonging to a cell based on its cell density. We calculate the center of each nucleus by averaging the locations of the identified spots associated with that nucleus. We employ the k-nearest neighbors (KNN) algorithm [80], linking each nucleus to its three closest neighbors. For each nucleus *i*, we record *size*_*i*_ as the number of spots occupied by the nucleus and the distances *d*_*io*_ (for *o* = 1, 2, 3) from the center of nucleus *i* to the centers of its three nearest neighbors. Assuming that the nuclei are approximately circular, we calculate 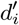 as:

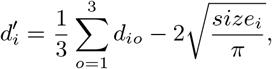

which represents the average distance between nearby cells minus twice the estimated radius (calculated from the cell size), thus approximating the maximum extent of the spots.

#### Step two: Cell Expansion

Having assigned transcripts to nuclei from all overlapping capture spots, we next sought to assign transcripts from nearby, non-overlapping capture spots, effectively expanding these nuclei into cells. During this nuclei expansion phase, we use an alternating optimization strategy to expand the nuclei, which generates robust cell gene profiles and reliable clustering results. In essence, we iteratively assign adjacent spots to cells based on whether their addition increases the cell’s similarity to the cluster to which it is assigned. We perform periodic reclustering to refine the assignments.

Here, we use the iteration variable *r* (starting from *r* = 1) to record each iteration and define the initial nuclei results as the starting point (*r* = 0). For example, as previously introduced, *X*^(0)^ represents the nuclei-by-gene expression matrix, *C*^(0)^ denotes the nuclei clustering results.

In the beginning of each iteration *r*, we calculate the average expression of each cluster, denoted by *w*^(*r*−1)^, which has dimensions *n*^(*r*−1)^ × *n*_genes_filtered_. For each cluster 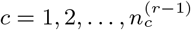 and for each gene *j*, the average expression is defined as

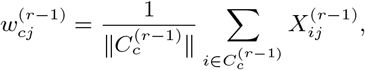

where 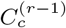 denotes the set of cells belonging to cluster *c* in *C*^(*r*−1)^ and 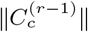 represents the number of cells in this set. Then, we normalize the gene expression by

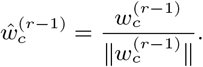

Additionally, we calculate

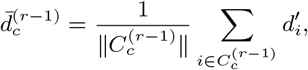

to decide the maximum expansion of spots for each nuclei. By default, we use

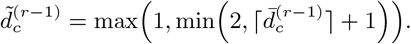

In the second stage, our goal is to derive the matrix *X*^(*r*)^ (*n*_*cells*_ × *n*_genes_ filtered_) so that cells within the same cluster exhibit high cosine similarity. We denote the gene expression vector for cell *i* in the current iteration as 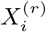 and let 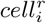 represent the spots assigned to cell *i* that were not previously associated with nucleus *i* in iteration *r*. The objective function is defined as follows:

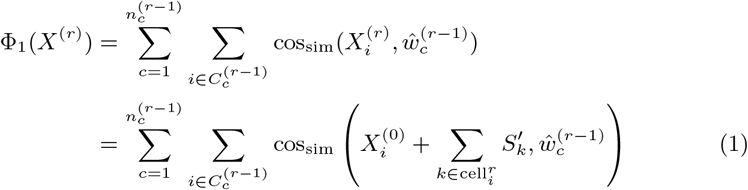

Here, our objective is to determine *cell*^*r*^ for *i* = 1, 2, …, *n*_*cells*_. For each spot *k* included in 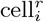, the distance from the spot to the nucleus must be less than 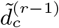, and the spots within a cell must form a connected region.

To achieve this within limited memory resources, we employ a greedy algorithm that expands the spots one-by-one in a specified order. A spot is added if its inclusion improves the objective function (1), and expansion in a particular direction ceases if the previously added spot is excluded.

In the next phase of iteration *r*, we analyze *X*^(*r*)^ following the Scanpy tutorial as described earlier, thereby obtaining the clustering information *C*^(*r*)^. To compare the clustering outcomes between iterations *r* and *r* − 1, we calculate the normalized mutual information score (NMI) using the formula:

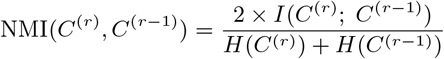

where *I*(*C*^(*r*)^; *C*^(*r*−1)^) denotes the mutual information between the clustering iterations, defined as:

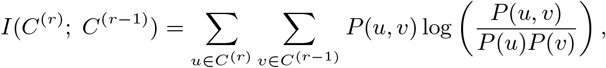

*H*(*C*^(*r*)^) and *H*(*C*^(*r*−1)^) represent the Shannon entropies of the respective clustering iterations, defined as:

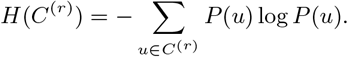

This calculation quantifies the consistency of clustering results between iterations *r* and *r* − 1, with higher NMI values indicating greater similarity. Typically, the NMI increases as the iterations progress, and by default, the process terminates when there is no improvement in the NMI between two consecutive iterations.

#### Step three: Count Deconvolution for Shared Spots

The previous step assigns transcripts from spots that do not overlap nuclei to cells; however, transcripts from a single spot may be assigned to multiple cells. In this step, we deconvolve counts from shared spots so that every transcript is assigned to at most one cell. We denote these assignments as *r* = *. At this stage, we isolate spots that are exclusively associated with a single cell and sum their gene counts to obtain 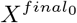. Next, we focus on the ambiguous spots, which can be categorized into two types: those shared by cells from different clusters, and those shared by cells within the same cluster. We first address the former by transforming situations where multiple clusters share spots into scenarios resembling the latter case. Finally, we apply deep learning optimization to the objective function (Equation 1) to determine the proportion of each cell cluster present within the shared spots.

For the first scenario, we simplify the problem to a case where cells from different clusters, denoted by the set *u*_*k*_, share a single spot *k*, with each cluster’s contribution represented by the weight matrix 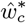. Due to the resolution of Spatial technologies, *u*_*k*_ typically contains 2 or 3 clusters. We then employ sequential least-squares programming and define the equation as:

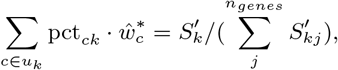

where 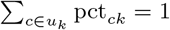 ensures that the proportions sum up to unity and 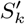 represents the gene expression array for spot *k*. After obtaining the values of pct_*ck*_, we employ a binomial model to determine the precise count via random number generation. We then define 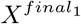 as 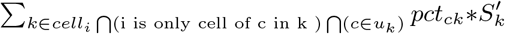.

Here, each specific cell *i* contains spots with predetermined area percentages. However, there remains a subset of spots *v*_*k*_ that are shared by at least two cells within the same cluster. We refine the objective function (2) to incorporate both the already assigned spots and those yet to be assigned, as follows:

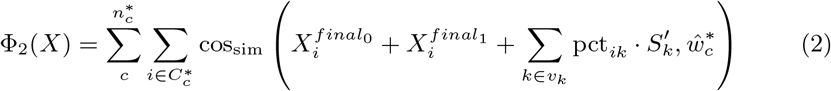

pct_*ik*_ satisfies the condition:

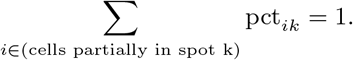

The maximization of Equation (2) is achieved through gradient-based optimization using the PyTorch library. Subsequently, we employ a binomial model to precisely determine counts via random number generation. Moreover, since each cell comprises a spot and its associated proportion, we revert to the unfiltered *S* to recover the gene expression count matrix for all genes.

#### Unrolling Analysis

Many tissues possess intricate biological structures that are challenging to study, especially since some tissues can become deformed on slides, obscuring their native organization. SMURF’s soft segmentation algorithm identifies distinct clusters of cells that include spatial information. Using these clusters to define a manifold, we propose a framework to unroll and elucidate the underlying biological architecture of tissue substructures.

We first select a specific cluster or the boundary of the cluster/tissue from the soft-segmentation output to guide the unrolling process. Because the relevant cluster may not perfectly correspond to the boundary of a tissue substructure, the set of cell coordinates can be augmented by user-defined points and by inserting the midpoints between existing cells. Cells that are isolated or located at the periphery are filtered out if their average distance to the nearest neighboring cells exceeds a specified threshold. Next, we apply Locally Linear Embedding (LLE) to preserve local neighborhood structures, yielding a new x-axis represented by the matrix *L* (of dimensions *n*_*l*_ × 2), where *n*_*l*_ denotes the number of selected cells. The LLE algorithm comprises several key steps: neighborhood selection, weight matrix construction, and low-dimensional embedding calculation. For each data point *l*_*i*_ in *L*, its local neighborhood is identified using the k-nearest neighbors method:

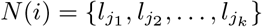

where *N* (*i*) denotes the neighborhood of the data point *l*_*i*_. For each data point *l*_*i*_, we compute weights *p*_*ij*_ that best approximate *l*_*i*_ using its neighbors *N* (*i*), with the objective to minimize the reconstruction error:

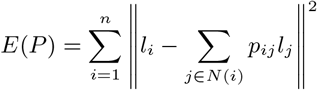

subject to the constraint ∑_*j*∈*N*(*i*)_ *p*_*ij*_ = 1, which guarantees that the weights sum to one and maintain local linearity. Once the weights *p*_*ij*_ are determined, the algorithm seeks a low-dimensional embedding *q*_*i*_ for each *l*_*i*_ that preserves the local geometry defined by these weights. The embedding is then optimized by minimizing the following cost function:

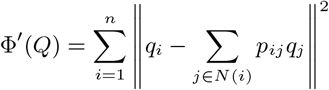

After obtaining *Q*, we reorder *L* according to the magnitude of *q*_*i*_ for each element in *L*. To present a smooth result with consistent distances between units on the x-axis, we first smooth the points *L*^*′*^ ((*n*_*l*_ − *n*_*avg*_ + 1) × 2) by averaging the closest *n*_*avg*_ points:

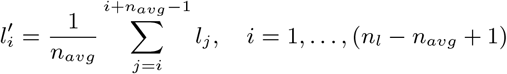

Next, we use interpolation functions to create an x-axis such that the actual distance between each unit point is consistent. We first calculate the Euclidean distances *d*_*i*_ between two nearby units in *L*^*′*^. The cumulative distance *D*_*i*_ is the total distance from the first point to the *i*-th point. Specifically, *D*_0_ = 0, and for *i* = 1, 2, …, (*n*_*l*_ − *n*_*avg*_), it is given by:

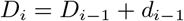

The total distance 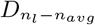 represents the distance from the starting point to the last point. A cubic interpolation function *f* is established for the coordinates of *L*^*′*^. We define a new sequence of distances containing *n*_*l*_ − *n*_*avg*_ points evenly distributed points 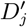, where:

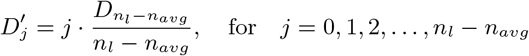

The new coordinates 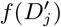 are computed using the interpolation functions for each 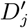, forming new points 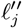. Thus, we create evenly and smoothly distributed points for the x-axis.

Depending on the tissue type, the y-axis is constructed either based on absolute distances or on cell connectivity. In the first case, for each cell, we identify its nearest unit on the x-axis and calculate the Euclidean distance from the cell to that unit, assigning this distance as the y-coordinate. In the second case, cells that are in close proximity are considered connected. For each cell, we determine the shortest connected path to the x-axis, define the y-layer as the number of cells along that path, and compute the y-coordinate by summing the distances along this path.

However, given the limited computational resources and the vast number of cells, calculating the shortest path for each cell individually is time-consuming. Moreover, the available paths are normally restricted to a limited set of directions. To overcome these challenges, we progressively generate y-layers by iteration. We begin by constructing the first layer using an iterative approach. For each unit on the x-axis, we identify the 18 closest cells within a specified distance and label them as Layer 1. Then, for each cell in Layer 1, we find the nearest neighboring cells among the remaining cells within the limited distance. To ensure that the cells are captured in the correct direction, we use a central reference point and require that the direction from the center to the original cell is consistent with the direction from the center to the Layer 1 cell. If a cell is selected multiple times by different Layer 1 cells, we retain the path with the shortest total length and record that path as the shortest one, assigning the corresponding x-axis coordinate to that cell. This process continues by iterations: in each iteration, the next layer is generated following the same procedure until no qualified cells are left.

### Mouse Brain data Analysis from 10x Genomics [26]

#### Nuclei Segmentation

SMURF requires nuclei segmentation results as input. In this study, we utilized Squidpy (v1.4.1) [81] with Stardist (v0.8.5) [18] to perform nuclei segmentation on mouse brain H&E data. We first used SMURF to get the exact spot-occupied area of the H&E plot. Since a small portion of the H&E staining in this data was damaged and exhibited abnormally dark red regions, we converted these areas to a pink hue similar to that of the blank regions. Subsequently, we divided the entire image into 20 blocks with a small overlapping area between adjacent sections. For each block, the “2D versatile he” model from Stardist was applied, and the segmentation results were then combined. To account for split cells at the block boundaries, we ensured that the overlapping regions were preserved and only retained cells that were completely contained within a block. These refined nuclei segmentation results were used as input for the SMURF analysis of the mouse brain data.

#### SMURF Analysis

We input the nuclei segmentation data along with the original output from 10x Genomics and followed the SMURF protocol to generate the final gene expression matrix along with primary clustering results. For the unrolling of the mouse cortex, we first selected the cluster corresponding to the basement membrane of the cortex, then filtered out isolated cells within this cluster and used the filtered cells to build the x-axis. We used the Euclidean distance to define the y-axis and selected a center point to determine the orientation. Next, we constructed both the x-axis and y-axis based on the spatial coordinates of the cells. The y-axis was evenly divided into five sections, and we subsequently plotted a heatmap of gene expression markers that delineated the distinct layers of the cortex [31, 82].

#### Cell type annotation

We uploaded the complete gene expression dataset directly from the SMURF output to MapMyCells (RRID: SCR 024672) to align cells with the 10X Whole Mouse Brain (CCN20230722) reference [33]. We then selected layer-specific neurons with subclass bootstrapping probabilities greater than 0.6 and plotted these cells along the new axes, with colors indicating layer information. Similarly, we filtered for specific non-neuronal cell types with subclass bootstrapping probabilities exceeding 0.6 and visualized them on the same coordinate system.

### Mouse Small Intestine Data Analysis from 10x Genomics [27]

#### Nuclei Segmentation

SMURF requires nuclei segmentation results as input. In this study, we utilized Squidpy (v1.4.1) [81] with Stardist (v0.8.5) [18] to perform nuclei segmentation on mouse small intestine H&E data. We first used SMURF to determine the exact spot-occupied areas of the H&E image. The entire image was then divided into 36 blocks with a small overlapping area between adjacent sections. For each block, the “2D versatile he” model from Stardist was applied, and the segmentation results were subsequently combined. To account for split cells at block boundaries, we preserved the overlapping regions and retained only those cells that were completely contained within a block. These refined nuclei segmentation results were used as input for the SMURF analysis of the small intestine data.

#### SMURF analysis

We followed the SMURF protocol to obtain the final gene expression matrix along with primary clustering results. For the unrolling analysis, we manually removed regions where the small intestine was inadvertently cut or damaged at both the beginning and the end. We then selected the clusters corresponding to the muscularis externa and myenteric plexus to define the x-axis, and filtered out isolated cells within these clusters. Finally, we adhered to the SMURF pipeline to define the y-axis layer by layer.

#### Single-Cell Downstream Analysis

We followed the Scanpy (v. 1.10.0) [19] pipeline to analyze the raw counts generated by SMURF. Initial filtering removed low-count cells based on SMURF’s cluster assignments: for cells in Peyer’s patches we required total counts to be at least smaller of 50 and 5 % of that cluster’s total counts, and for all other clusters we required total counts to be at least the larger of 50 and 10 % of that cluster’s total counts. We then excluded cells with 2000 or more detected genes, 2000 or more total counts, a mitochondrial gene fraction of 25 % or higher, or a cell size of 45 or greater. These quality-control thresholds ensured the removal of low-quality or abnormal cells, enabling robust downstream analysis.

We then normalized the cells by scaling each cell’s total counts to 1000 and performed log-normalization to stabilize variance across cells. Next, we retained the top 4000 variable genes and two key genes (Pyy and Sct). The data were scaled such that the maximum expression value for each gene was set to 10. We performed PCA to reduce the dimensionality to 40 components, followed by the construction of a nearest-neighbor graph using 25 neighbors. For visualization, we applied UMAP, and cell clustering was conducted using the Leiden algorithm with a resolution of 3 [83, 21]. Differential expression analysis was carried out using the Scanpy (scanpy.tl.rank genes groups, wilcoxon) [19]. We annotate the cell types by gene markers according to reference data and knowledge with the help of location information [36, 84, 85].

To detect the crypt–villus boundary, we first performed a rough division based on a normalized distance threshold of 9, temporarily assigning cells with lower values to the crypt group and those with higher values to the villus group. We then conducted differential gene expression analysis between these two groups, selecting genes with an adjusted p-value below 1 × 10^−5^. Using these genes, we built a random forest classifier by XGBoost (v 2.1.1) [86] to distinguish between crypt and villus cells based on their expression profiles. Recognizing that the crypt–villus continuum is a continuous structure, we further examined the local neighborhood of each cell. We assumed that the distribution of normalized y-axis positions for the crypt and villus groups follows a mixed Gaussian distribution and used the resulting boundary to precisely define the crypt–villus interface at each location.

#### Pseudotime Analysis

We used Scanpy (v1.10.0) [19] to perform pseudotime analysis on a subset of cells, including Paneth and stem cells, TA cells, and enterocytes. First, we computed diffusion maps to capture the underlying cellular trajectories. One cell was randomly selected from the Paneth and stem cells group as the putative starting point, and pseudotime was computed using the diffusion pseudotime function (sc.tl.dpt) [46]. To better visualize the structure of this cell subset, we employed PAGA (sc.tl.paga) [47] and force-directed graph drawing (sc.tl.draw_graph) [48] to generate a layout on which the pseudotime values were mapped.

We then selected cells that were not part of the highly concentrated Peyer’s patches and normalized both the pseudotime values and the normalized corrected y-axis to a scale from 0 to 1. Using pseudotime (0-1) as the independent variable and the normalized corrected y-axis (0-1) as the dependent variable, we performed linear regression analysis and computed the Pearson correlation coefficient to assess the strength and significance of their relationship.

#### Zonation Analysis

We selected only intact complete villi, including tip enterocytes, for zonation analysis. The intestine was divided into small sections along the x-axis, and we applied a Kolmogorov–Smirnov test to evaluate the distribution of tip enterocytes in each section, following a uniform distribution. To ensure comprehensive coverage, multiple candidate sections from each area that passed the test were retained, and the final selection was refined through manual curation. We retained Paneth and stem cells, TA cells, and enterocytes. We then subdivided the villi into six distinct zones according to their normalized, corrected y-axis, ensuring that each zone contained an equivalent number of cells. Gene count were pseudobulked in each zone. and only genes with counts greater than 10 were retained (12,005 genes in total). We normalized the average gene expression of each gene across the zones to a maximum expression of 1. To identify the genes that were significantly zonated, we used pair-wise differential expression analysis of each zone against all other zones using Scanpy (scanpy.tl.rank_genes_groups, wilcoxon) [19]. We considered a gene as significantly zonated if its absolute log2-fold-change (LFC) was greater than 1.5 and the adjusted p-value was below 1 × 10^−8^ compared to all other zones.

#### KEGG Analysis

We used gprofiler-official (v1.0.0) [87] to analyze KEGG pathway enrichment for each zone. KEGG pathways were retained if any zone exhibited an adjusted p-value below 1 × 10^−8^ and an LFC of 1.5. We generated bubble plots of significant KEGG pathways across all zones, where the bubble size indicates the fraction of genes from that pathway that were significantly enriched in the zone, and the bubble color represents the corresponding p-value.

#### Gene Expression by Zone

We used the y-axis boundaries identified in zonation analysis for all plots of cell-type-specific gene expression by zones. Cells in the designated cluster were split by zone and pseudobulked. Plots show the average log expression of the gene in each zone.

We then sought to capture proximal–distal differential genes across different zones. Cells were first divided along the x-axis, with the first half designated as the proximal group and the second half as the distal group. Only the subset of Paneth and stem cells, TA cells, and enterocytes was used, with each zone containing an approximately equal number of cells. For each zone, mitochondrial genes were filtered out, and only genes expressed in at least 100 cells were retained. Differential expression analysis was performed using an LFC threshold of 1.5 and an adjusted p-value cutoff of 0.001. The number of differential genes was then normalized by dividing by the total number of genes remaining after filtration. Finally, we generated plots of the differential gene count and the fraction of differential genes versus zones. KEGG pathways of the proximal and distal zone 6 genes were calculated if they exhibited an adjusted p-value below 0.001 and an LFC of 1.5. Line-plot trajectories were drawn for important marker genes across the zonation axis. Because stem, Paneth, and TA cells are predominantly confined to Zones 1–2, we displayed these three populations together with enterocytes for direct comparison. For every other lineage, the plot was restricted to cells of that specific type.

#### Transcription Factor Analysis

We selected Paneth cells, stem cells, TA cells, and enterocytes — each annotated with villus-zone information — from the previous section, and then followed the pySCENIC protocol (https://github.com/aertslab/SCENICprotocol) by applying `pyscenic` to infer TF–target gene regulatory networks and to identify and score regulons significantly enriched in our dataset by comparison with established reference databases [58].

A total of 186 regulons have been called by the results. We selected zone-specific regulons (LFC *>* 0.3 and adjusted *p*-value *<* 1 × 10^−10^) and plotted the TF–target network for all six zones. For Zone 2, we built a TF–target network for the seven regulons whose activities differed significantly across zones by differential t-test from Scanpy. In the graph, large blue nodes denote the TF genes, small green nodes mark non-zonated targets, red nodes mark zonated targets, and nodes in additional colours highlight zonated targets enriched for specific functional categories. To probe zonation enrichment, we applied Fisher’s exact tests to compare genes targeted by at least two TFs with all remaining targets and to compare each TF, comparing its unique targets with the pool of all other target genes separately. We completed the same analysis for Zone 6 with five differential regulons.

#### Single-cell Analysis of Seiler et al [71]

We downloaded the dataset from GEO (GSE130113), which contains raw counts from three sham and two SBR replicates. We first filtered cells by requiring a minimum of 200 genes per cell, and filtered genes by retaining only those expressed in at least 3 cells. The data were then normalized to a total of 10,000 counts per cell and log-normalized. We selected highly variable genes, scaled the data, performed PCA, and applied Harmony for batch correction. UMAP was used for dimensionality reduction, and clustering was performed using the Leiden algorithm [83, 21]. Differential expression analysis was carried out using a t-test. Although the differential results were not exactly identical to those reported in the original paper, they were extremely similar—the top five genes were exactly the same, and all the top 10 genes reported in the paper appeared as the top 11 in our analysis. Finally, we selected differential genes using a threshold of adjusted p-value less than 1 × 10^−50^ and an absolute LFC larger than 1, resulting in a total of 77 genes significantly differentially expressed in the SBR group.

We then mapped the differential genes to the spatial dataset, and 72 of them were expressed in the spatial data. Next, we sought to determine the specific zone in which each gene was predominantly expressed by defining a gene as expressed in a particular zone if its LFC in that zone was greater than 1.5 and its adjusted p-value was less than 1 × 10^−8^. Finally, we plotted the zones in which all SBR differential genes were expressed in the spatial dataset.

### Casting cells across different tissues and platforms

#### Mouse Olfactory Bulb—Stereo-seq Dataset [2]

We cropped one hemibulb and selected the boundary part of the hemibulb as the unfolding reference for the x-axis. The y-axis was computed as Euclidean distances to the ONL in two segments and then merged as above. We selected a contiguous region bounded by isocontours and visualized spatial ordering with violin plots of y-axis distributions across major cell types.

#### Human Frontal Cortex—NanoString CosMx Dataset [41]

We directly used the cell-level segmentation, quality control, and annotation results as input. We treated the `astro.1` boundary cells as the reference trajectory for the x-axis. For each cell, y-axis was defined as the Euclidean distance to the nearest point on this axis. To accommodate local curvature, the tissue was partitioned into two segments; distances were computed within each segment and then concatenated to preserve a consistent origin and units. We used MapMyCells (RRID: SCR 024672) to annotate the cells. We then selected a contiguous region bounded by density isocontours and summarized layer-specific organization using violin plots of the y-axis distributions for cortical cell types.

#### Mouse Retina—MERFISH Dataset [42]

We analyzed one of the retinas from sample **VZG105a WT2**. Retinal ganglion cells were used to define the x-axis for tissue unrolling. For all other cells, y-axis positions were computed as the Euclidean distance to the RGC axis, and cells at the lateral margins were removed. Because retinal cell types are arranged in layers, we summarized y-axis distributions by cell type and marker genes using violin plots.

#### Human Skin Melanoma—10x Xenium Prime Dataset [43]

We used the clustering results from *10x Genomics* website. Cluster 28 defined the x-axis reference, and y-axis was computed as Euclidean distances to this axis. We generated violin plots for the four principal epidermal clusters. In a complementary unrolling, the basal layer (cluster 20) was set as the origin (*y* = 0), with epidermal layers mapped to *y* ≤ 0 and extra-epidermal areas to *y >* 0.

#### Comparison Analysis for Results from Different Methods

We compared the results of 8 *µ*m spot data, Bin2cell results, nuclei segmentation results, and SMURF results. For each dataset, we used the direct output of 4×4 spots (corresponding to an 8 *µ*m × 8 *µ*m spot size) as the 8 *µ*m spot data. For Bin2cell, we followed the instructions provided in the Bin2cell GitHub repository [88]. Nuclei segmentation results were obtained using Squidpy (v1.4.1) [81] with Stardist (v0.8.5) [18].

In the analysis of the percentage of cells covered by nuclei, we encountered the issue that a single spot might encompass several nuclei, leading to multiple nuclei being assigned to a cell, an outcome that is not accurate. To address this, we defined a cell as covered by a nucleus only if at least 30% of the pixels in the cell region are occupied by that nucleus.

For the definition of nuclei data in the Mouse Brain dataset, we considered a spot to belong to a nucleus if at least 90% of the spot’s counts were within that nucleus. We then uploaded the raw counts matrix of nuclei, SMURF, and Xenium results from 10x [28] to MapMyCells [32, 33] by selecting the “10x Whole Mouse Brain (CCN20230722)” as the reference dataset. Using the Hierarchical Mapping strategy available in MapMyCells, we obtained the cell type results for the three datasets. We calculated the KL divergence according to the “class name” from the MapMyCells results. In the mouse small intestine dataset, we applied the same segmentation and clustering pipeline as provided by 10x Genomics [89].

## Data Availability

The public mouse brain Visium HD and mouse small intestine Visium HD datasets are available on the 10x Genomics website ([mouse brain Visium HD, mouse intestine Visium HD](https://www.10xgenomics.com/datasets/visium-hd-cytassist-gene-expression-libraries-of-mouse-brain-he)). Public mouse brain 10x Xenium data can be accessed on the 10x Genomics website (fresh-frozen mouse brain replicates). Public Mouse retina MER-FISH data (sample ID `VZG105a_WT2`) is from Choi et al. (2023) [42] and is available via Zenodo ([Zenodo record](https://doi.org/10.5281/zenodo.8144355)). The public adult mouse olfactory bulb Stereo-seq dataset is from Chen et al. (2022) [2] and can be downloaded from the STOmics/MOSTA portal ([MOSTA download (Mouse_olfa_S2.h5ad)](https://db.cngb.org/stomics/mosta/download/)). The public human frontal cortex NanoString CosMx SMI dataset can be accessed on the NanoString website ([Human frontal cortex CosMx](https://nanostring.com/products/cosmx-spatial-molecular-imager/ffpe-dataset/human-frontal-cortex-ffpe-dataset/)) [41]. Public Human Skin Melanoma 10x Xenium prime dataset is available on the 10x Genomics website[Human Skin Melanoma 10x Xenium prime](https://www.10xgenomics.com/datasets/xenium-prime-ffpe-human-skin)) [43]. The public SBR experiment dataset from Seiler et al (2019) [71]. is available from GEO under accession number GSE130113. The ISH staining for gene Slc1a2 was obtained from the Allen Brain Atlas (https://mouse.brain-map.org/gene/show/20273).

## Code Availability

The SMURF package is available at https://github.com/The-Mitra-Lab/SMURF. The code to reproduce the results presented in this study can be accessed at https://github.com/The-Mitra-Lab/SMURF_figures. The documentation for SMURF is available at https://the-mitra-lab.github.io/SMURF/.

## Computational resources

All analyses were executed on a single Ubuntu 20.04.6 LTS workstation equipped with an Intel^®^ Core™ i9–10920X CPU (12 cores/24 threads), 251 GiB RAM, and an NVIDIA GeForce RTX 3090 GPU (24 GiB VRAM).

## Acknowledgements

We thank Hao Fan for discussions that helped shape this project, and Jennifer Hayden for scientific illustrations. The contents of this manuscript do not represent the views of the Department of Veterans Affairs or the United States Government.

## Extended Data Figures

**Extended Data Figure 1:**
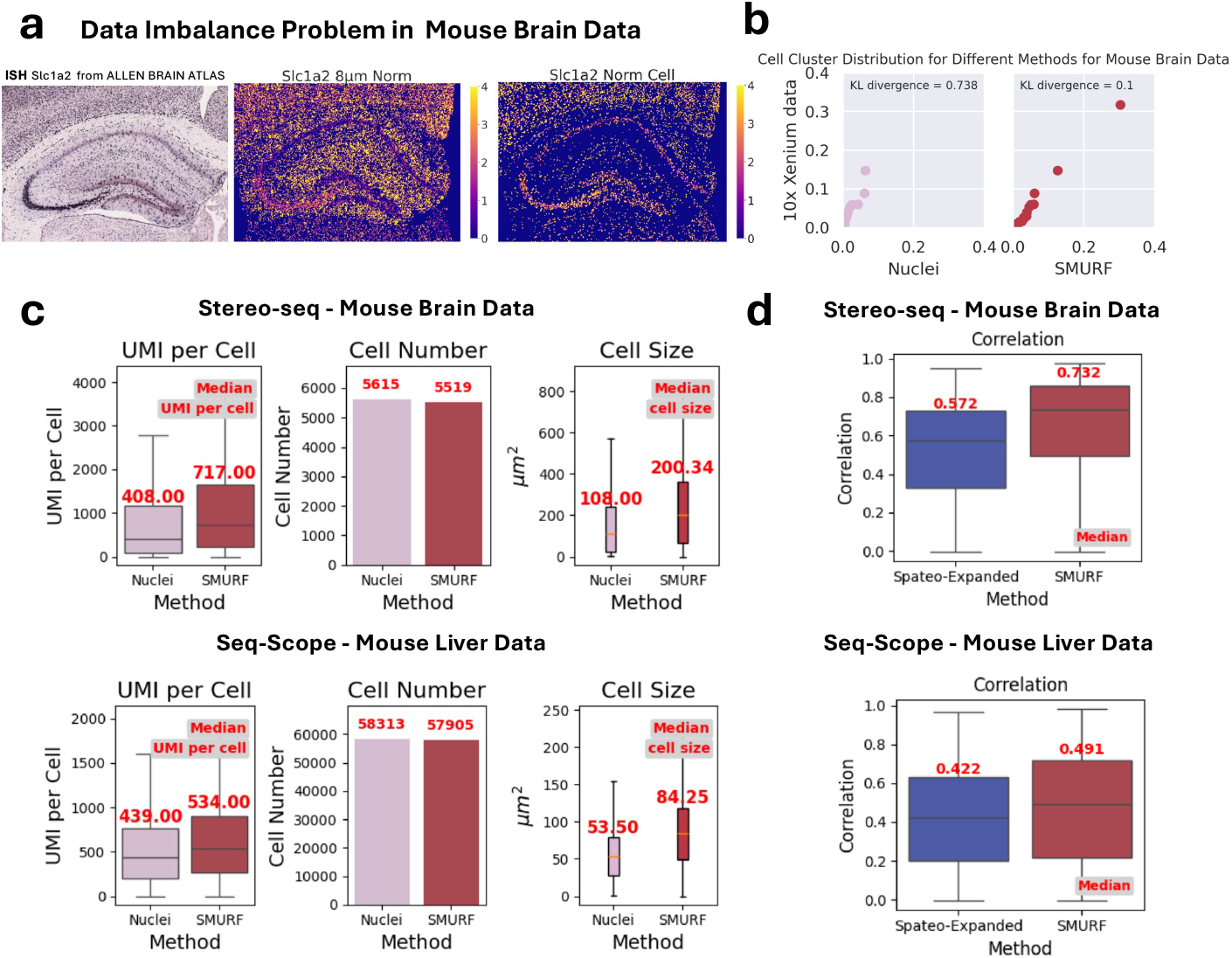
Soft segmentation with SMURF. (a) Illustration of the data imbalance issue when the standard normalized 8 *µ*m Spot method is used. The spatial expression pattern of Slc1a2 is discordant between Visium HD mouse brain data analyzed by the 8 *µ*m Spot method and ISH data from the Allen Brain Atlas. In contrast, when analyzed by SMURF, there is concordance. (b) Cell type proportions annotated by MapMyCells in probe-based mouse brain data, comparing nuclei results and SMURF results. KL divergence values quantify differences in cell type distributions. Lower KL divergence indicates better concordance. (c) Median number of UMIs assigned to cells, total cells detected, and median cell size using SMURF or nuclei-based segmentation (d) Per-cell UMI correlation between nuclei segmentation and Spatio/SMURF expansion.

**Extended Data Figure 2:**
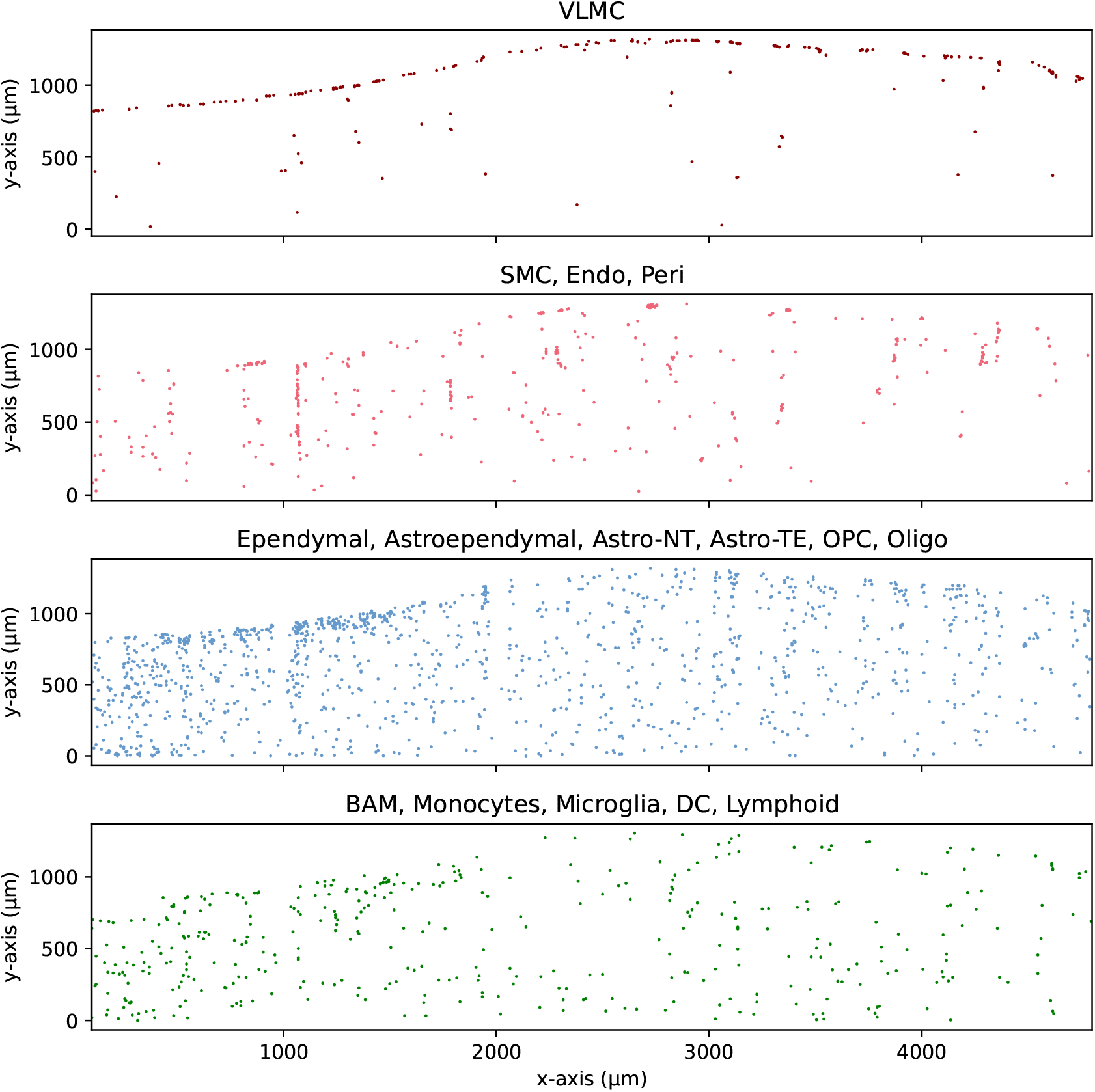
Distribution of cortical layer-specific non-neuronal cells in the unrolled rectilinear coordinate system. Cell segmentation and annotation of mouse cortical cells as in main text Figure 2.

**Extended Data Figure 3:**
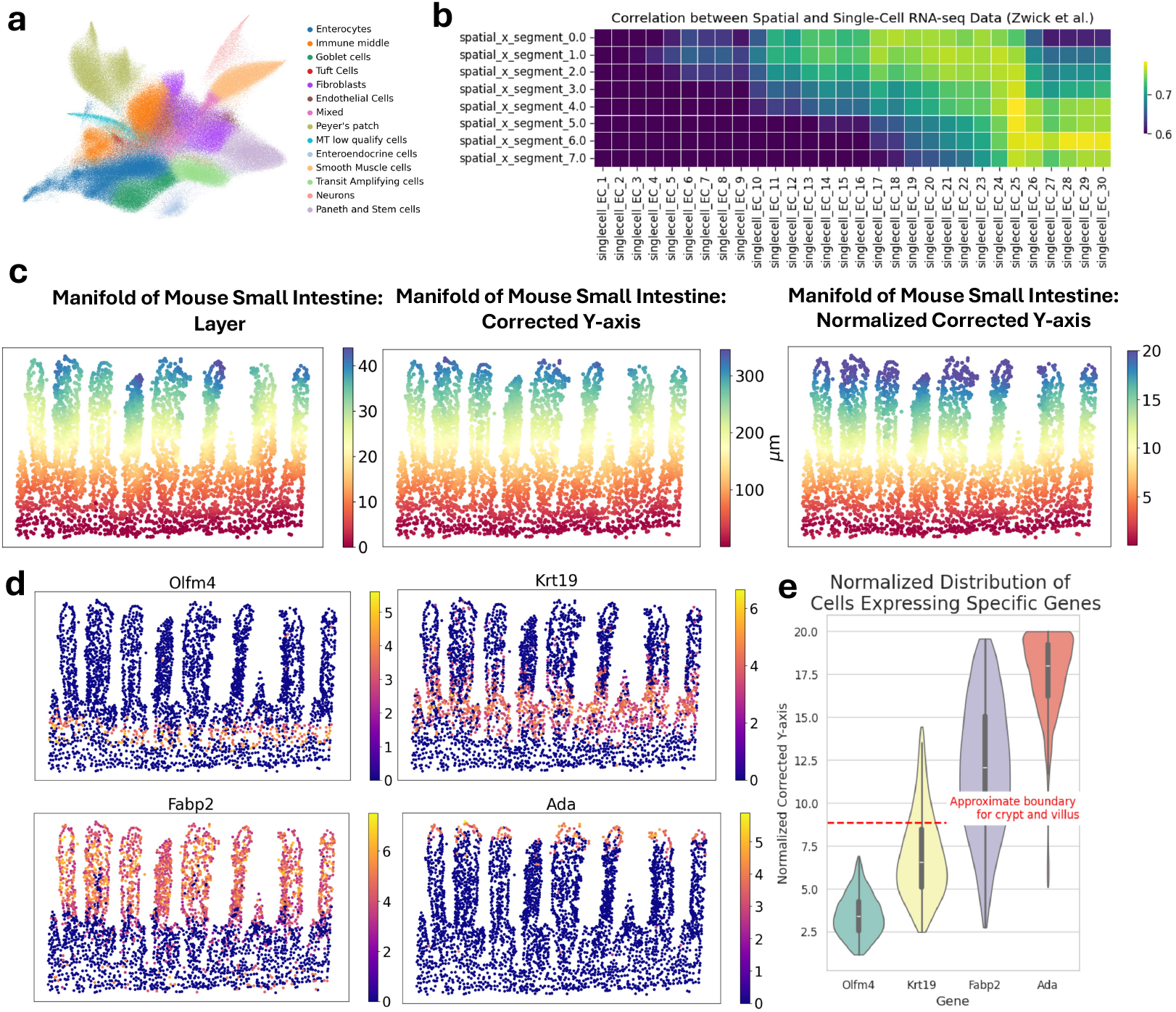
Mouse small intestine analysis. (a) UMAP embedding of intestinal cells, colored by annotated cell types. (b) Correlation between single-cell transcriptomes from Zwick et al. [36] and spatial transcriptomic profiles across intestinal sections. (c) Overlay of selected mouse small intestine data with cells colored by coordinate value along the y-axis at different stages of “unrolling”: raw layer positioning, corrected Y (*µ*m), and normalized corrected Y. (d) Marker gene expression visualized in the selected unrolled intestinal tissue (*Olfm4, Krt19, Fabp2*, and *Ada*). (e) Normalized corrected y-axis distribution of cells expressing selected genes with red dashed lines indicating the approximate boundary between crypt and villus regions.

**Extended Data Figure 4:**
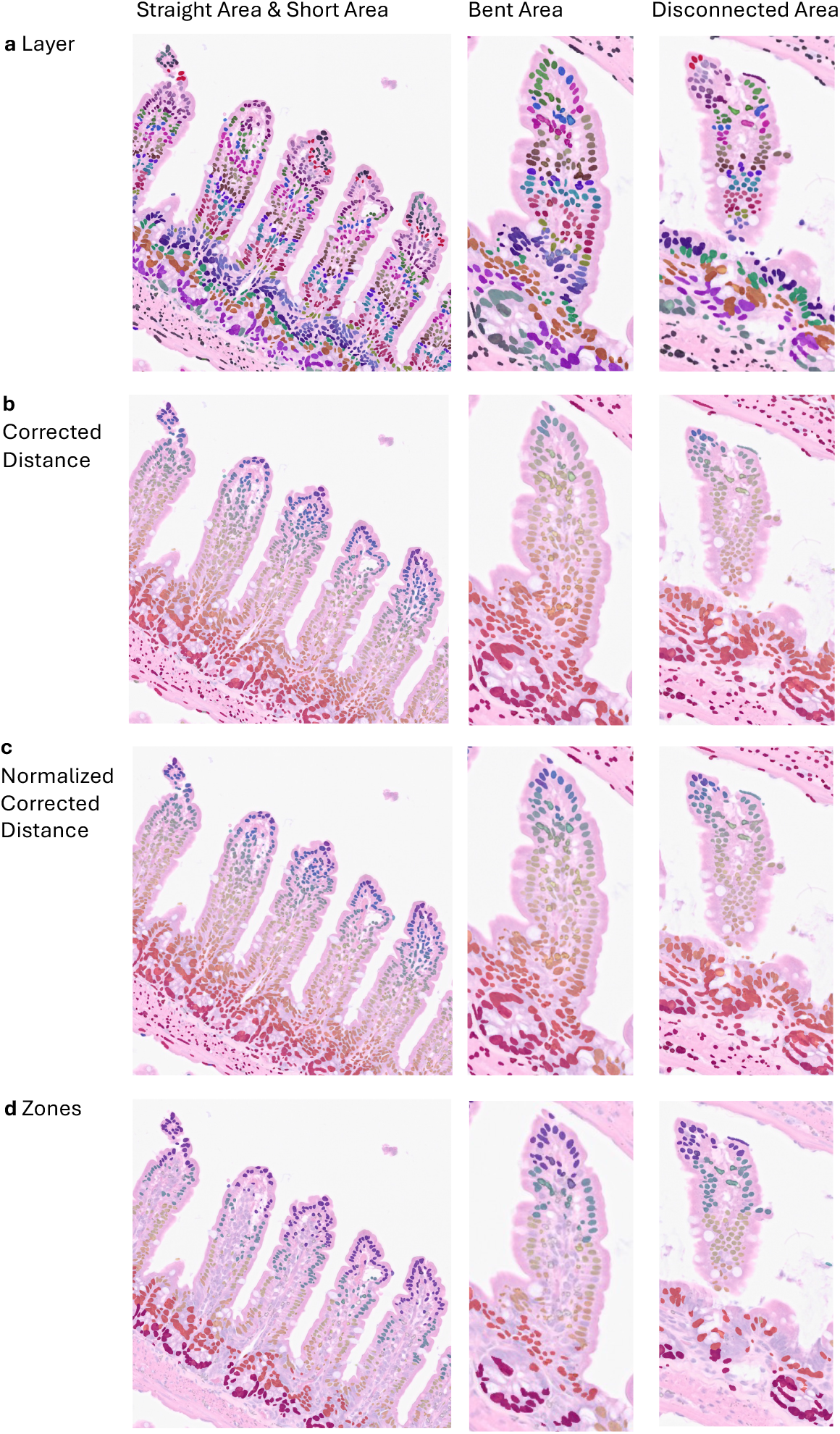
Different Distances Examples Mapped on Real H&E Plot. (a) Layer. (b) Corrected Distance. (c) Normalized Corrected Distance. (d) Zone.

**Extended Data Figure 5:**
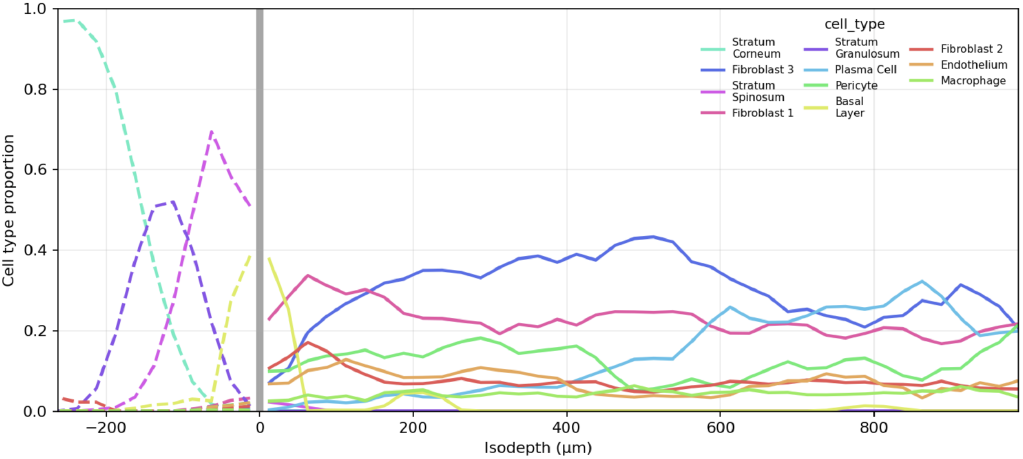
Epidermal zonation across isodepth (basal layer anchored). (a) Proportions of different cell clusters as a function of the isodepth with the basal layer as the zero axis.

**Extended Data Figure 6:**
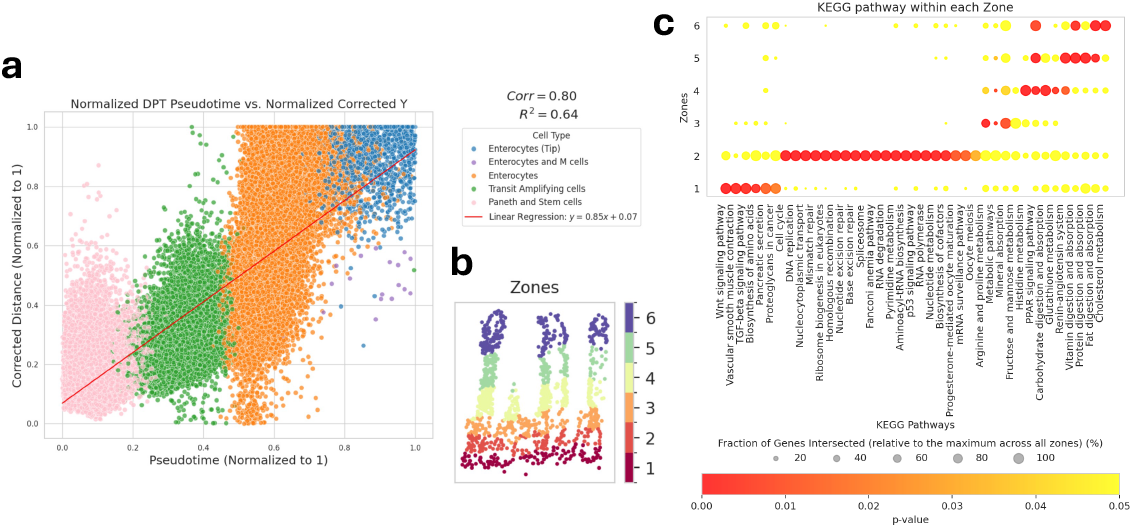
Spatial zoning of enterocytes: pseudotime–depth concordance and KEGG enrichment. (a) Scatter plot comparing the normalized pseudotime progression score and the normalized corrected y-axis. Each point represents a single cell. The dashed red line indicates a linear fit (Pearson correlation = 0.80). (b) Example of the discrete binning of cells of the intestinal villus into 6 zones along the y-axis. (c) Bubble plot displaying KEGG pathway enrichment of significantly zonated genes.

**Extended Data Figure 7:**
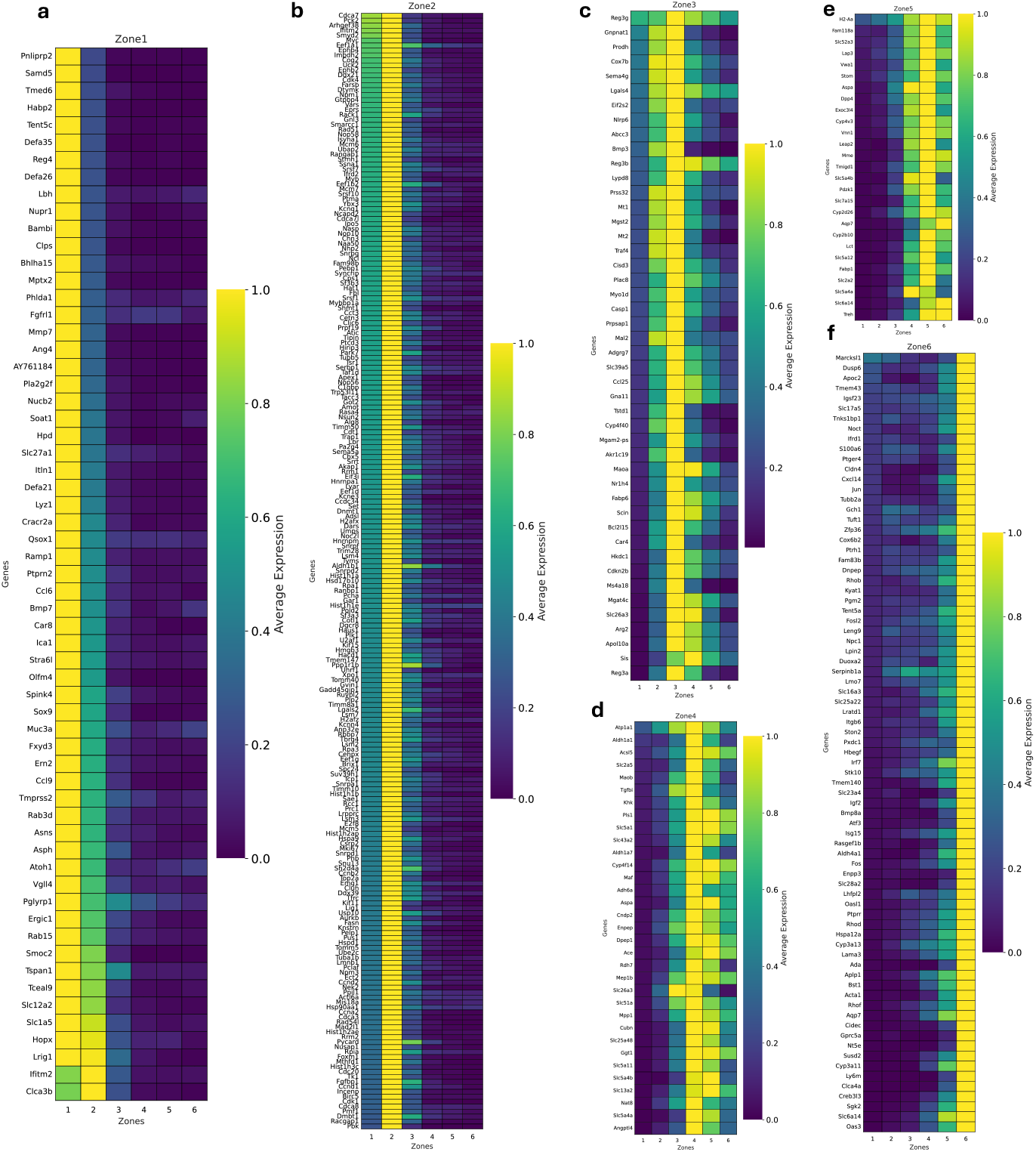
Highly zonated enterocyte genes. Statistically enriched genes were identified in each zone. These lists were manually inspected to select the most illustrative examples of zonated genes. Heatmaps depict normalized gene expression for Zone 1 (a), Zone 2 (b), Zone 3 (c), Zone 4 (d), Zone 5 (e), and Zone 6 (f).

**Extended Data Figure 8:**
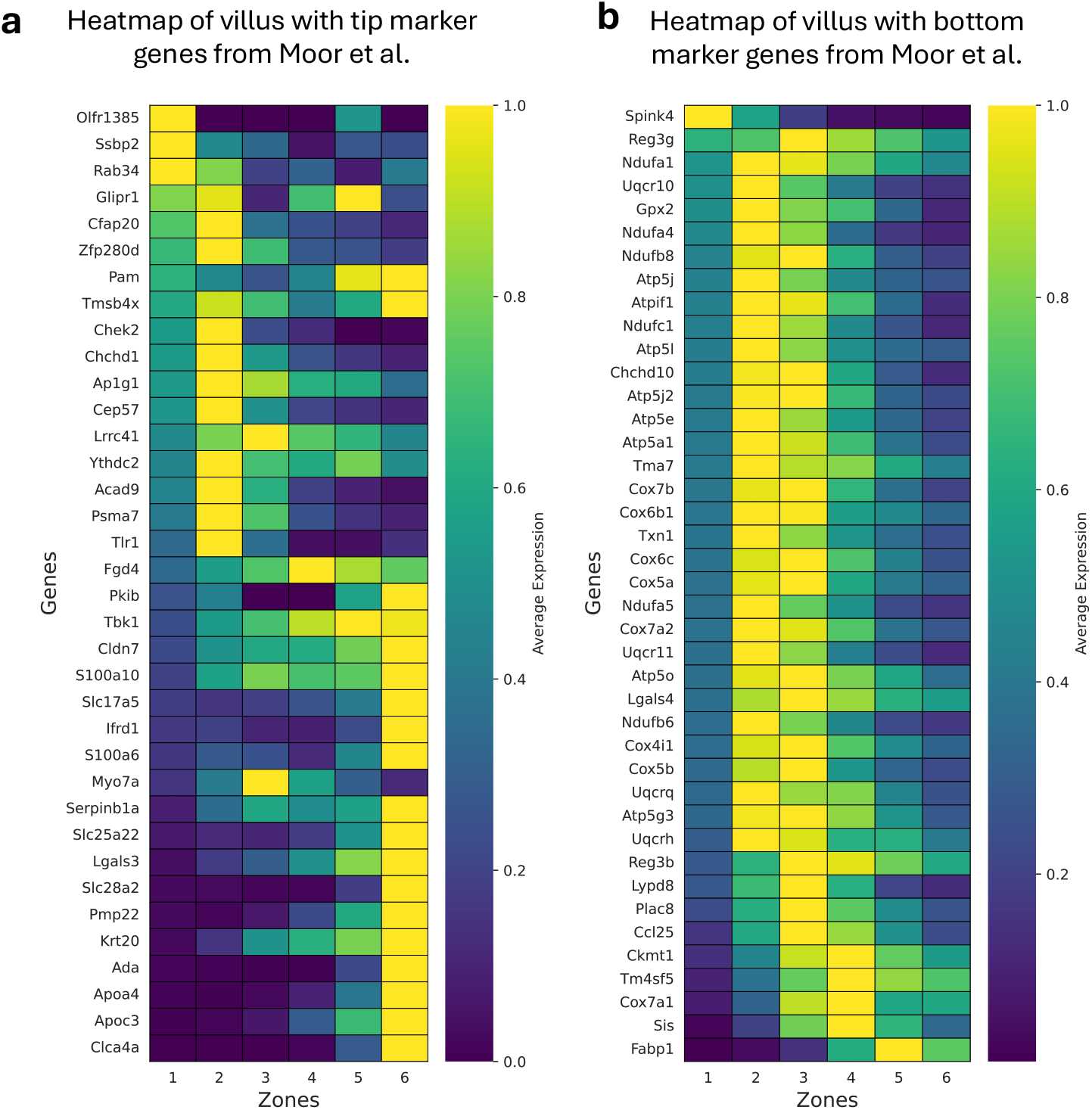
Heatmap of villus tip and bottom marker genes from Moor et al. [49] in the ileal SMURF dataset. (a) Heatmap of villus with tip marker genes from Moor et al. (b) Heatmap of villus with bottom marker genes from Moor et al.

