## Supplementary Table 1 for "SMURF: soft-segmentation for single-cell reconstruction and topological analysis of spatial transcriptomic data"

Zone 1: Significantly Zonated Genes

| Symbol | scores | LFC | pvals | pvals_adj |
|---|---|---|---|---|
| Ang4 | 103.97 | 7.04 | 0 | 0 |
| Defa21 | 103.03 | 7.18 | 0 | 0 |
| AY761184 | 102.77 | 6.94 | 0 | 0 |
| Itln1 | 102.74 | 6.93 | 0 | 0 |
| Lyz1 | 102.21 | 7.01 | 0 | 0 |
| Mptx2 | 100.58 | 6.87 | 0 | 0 |
| Mmp7 | 84.71 | 5.81 | 0 | 0 |
| Spink4 | 82.65 | 5.25 | 0 | 0 |
| Clps | 80.90 | 5.79 | 0 | 0 |
| Olfm4 | 68.85 | 4.61 | 0 | 0 |
| Cd24a | 54.82 | 3.93 | 0 | 0 |
| Pnliprp2 | 54.12 | 4.89 | 0 | 0 |
| Reg4 | 53.11 | 4.71 | 0 | 0 |
| Defa26 | 52.74 | 4.82 | 0 | 0 |
| Nupr1 | 43.33 | 4.26 | 0 | 0 |
| Habp2 | 41.02 | 4.61 | 0 | 0 |
| Car8 | 33.93 | 3.29 | 2.7E-252 | 1E-249 |
| Muc2 | 30.76 | 1.72 | 1E-207 | 2.8E-205 |
| Cracr2a | 29.84 | 3.28 | 1.2E-195 | 3.1E-193 |
| Samd5 | 29.61 | 4.56 | 9.9E-193 | 2.5E-190 |
| Acta2 | 28.70 | 3.07 | 3.6E-181 | 8.3E-179 |
| Selenom | 28.55 | 2.58 | 2.7E-179 | 5.9E-177 |
| Defa35 | 28.06 | 4.36 | 3.4E-173 | 7.2E-171 |
| Qsox1 | 27.90 | 2.66 | 3E-171 | 6.2E-169 |
| Tent5c | 27.77 | 4.20 | 9.7E-170 | 1.9E-167 |
| Lbh | 26.75 | 3.40 | 1.4E-157 | 2.4E-155 |
| Dcn | 26.29 | 3.30 | 2.3E-152 | 4E-150 |
| Slc12a2 | 26.09 | 2.57 | 4.2E-150 | 7.2E-148 |
| Ccl6 | 25.75 | 3.04 | 3.2E-146 | 5.4E-144 |
| Ivns1abp | 25.46 | 1.73 | 5.8E-143 | 9.5E-141 |
| Asph | 24.90 | 2.51 | 8.2E-137 | 1.3E-134 |
| Clca3b | 24.83 | 2.02 | 4.8E-136 | 7.4E-134 |
| Pla2g2f | 24.46 | 3.72 | 3.6E-132 | 5.5E-130 |
| Tmprss2 | 24.23 | 2.32 | 1E-129 | 1.5E-127 |
| Tagln | 24.08 | 2.77 | 4E-128 | 5.8E-126 |
| Agr2 | 23.91 | 1.73 | 2.4E-126 | 3.4E-124 |
| Tmed6 | 23.76 | 4.07 | 9.5E-125 | 1.3E-122 |
| Rgmb | 22.92 | 2.46 | 3E-116 | 3.9E-114 |
| Bambi | 22.01 | 3.98 | 2.5E-107 | 3.1E-105 |
| Tceal9 | 21.95 | 2.43 | 7.9E-107 | 9.5E-105 |
| Tspan1 | 21.41 | 2.05 | 1.1E-101 | 1.3E-99 |

| | | | | |
|---|---|---|---|---|
| Muc3a | 21.39 | 2.26 | 1.7E-101 | 1.9E-99 |
| C3 | 21.29 | 3.99 | 1.3E-100 | 1.49E-98 |
| Hpd | 21.27 | 3.57 | 2E-100 | 2.28E-98 |
| Col1a1 | 21.03 | 2.40 | 3.79E-98 | 4.2E-96 |
| Ramp1 | 20.82 | 2.99 | 2.97E-96 | 3.18E-94 |
| Tpm2 | 20.81 | 2.75 | 3.78E-96 | 4E-94 |
| Col1a2 | 20.59 | 2.57 | 3.33E-94 | 3.49E-92 |
| Reep5 | 20.58 | 3.10 | 3.75E-94 | 3.88E-92 |
| Pglyrp1 | 20.04 | 1.86 | 2.39E-89 | 2.32E-87 |
| Slc1a5 | 19.97 | 2.18 | 9.63E-89 | 9.18E-87 |
| Sparc | 19.81 | 2.34 | 2.22E-87 | 2.1E-85 |
| Ifitm2 | 19.64 | 1.82 | 7.05E-86 | 6.56E-84 |
| Hopx | 19.64 | 2.03 | 7.36E-86 | 6.81E-84 |
| Dpt | 19.24 | 3.01 | 1.84E-82 | 1.67E-80 |
| Sox9 | 19.21 | 2.86 | 3.09E-82 | 2.79E-80 |
| Rab3d | 19.11 | 2.48 | 2.16E-81 | 1.92E-79 |
| Myh11 | 18.99 | 2.53 | 2.21E-80 | 1.96E-78 |
| Smoc2 | 18.81 | 2.58 | 6.02E-79 | 5.24E-77 |
| Mgp | 18.68 | 3.66 | 7.14E-78 | 6.1E-76 |
| Ern2 | 18.36 | 2.63 | 2.83E-75 | 2.35E-73 |
| Stra6l | 18.32 | 2.78 | 6.02E-75 | 4.97E-73 |
| Apoe | 18.01 | 1.72 | 1.59E-72 | 1.28E-70 |
| Ergic1 | 18.00 | 2.38 | 2.09E-72 | 1.68E-70 |
| Bhlha15 | 17.98 | 3.71 | 2.86E-72 | 2.29E-70 |
| Wnt3 | 17.61 | 4.28 | 1.99E-69 | 1.54E-67 |
| Kcnma1 | 17.53 | 3.76 | 8.69E-69 | 6.65E-67 |
| Col3a1 | 17.47 | 1.90 | 2.43E-68 | 1.83E-66 |
| Igfbp7 | 17.33 | 1.87 | 2.81E-67 | 2.06E-65 |
| Actg2 | 17.30 | 2.62 | 4.92E-67 | 3.55E-65 |
| Igfbp4 | 17.09 | 2.55 | 1.64E-65 | 1.16E-63 |
| Myl9 | 16.90 | 2.60 | 4.23E-64 | 2.93E-62 |
| Phlda1 | 16.84 | 3.03 | 1.23E-63 | 8.42E-62 |
| Rgs5 | 16.79 | 2.79 | 3.09E-63 | 2.1E-61 |
| Ptprn2 | 16.71 | 2.83 | 1.19E-62 | 7.94E-61 |
| C4b | 16.43 | 3.21 | 1.15E-60 | 7.3E-59 |
| Copz2 | 16.42 | 3.98 | 1.44E-60 | 9.05E-59 |
| Nucb2 | 16.28 | 3.44 | 1.34E-59 | 8.26E-58 |
| Fgfrl1 | 16.16 | 2.68 | 9.88E-59 | 6.03E-57 |
| Rab15 | 16.01 | 2.33 | 1.07E-57 | 6.4E-56 |
| Flna | 15.90 | 2.37 | 6.04E-57 | 3.54E-55 |
| Vgll4 | 15.90 | 2.31 | 6.57E-57 | 3.82E-55 |
| Rnase4 | 15.86 | 1.54 | 1.27E-56 | 7.32E-55 |
| Stmn1 | 15.53 | 1.56 | 2.08E-54 | 1.17E-52 |
| Fxyd3 | 15.53 | 2.59 | 2.23E-54 | 1.24E-52 |

| | | | | |
|---|---|---|---|---|
| Soat1 | 15.44 | 3.19 | 8.76E-54 | 4.83E-52 |
| C1qtnf1 | 15.25 | 3.81 | 1.77E-52 | 9.59E-51 |
| Sec11c | 15.24 | 1.56 | 1.85E-52 | 1E-50 |
| Ccl9 | 15.09 | 2.69 | 1.77E-51 | 9.33E-50 |
| Sntb1 | 15.00 | 3.13 | 7.69E-51 | 3.99E-49 |
| Syne4 | 14.96 | 3.84 | 1.35E-50 | 6.95E-49 |
| Ica1 | 14.82 | 2.76 | 1.08E-49 | 5.46E-48 |
| Slc1a4 | 14.77 | 3.74 | 2.38E-49 | 1.19E-47 |
| Atp2a3 | 14.72 | 1.61 | 4.69E-49 | 2.33E-47 |
| Lgr5 | 14.64 | 3.19 | 1.53E-48 | 7.49E-47 |
| Atoh1 | 14.56 | 2.17 | 5.04E-48 | 2.43E-46 |
| Kif12 | 14.55 | 3.36 | 5.63E-48 | 2.69E-46 |
| Actn1 | 14.51 | 1.84 | 1.08E-47 | 5.12E-46 |
| Sync | 14.48 | 3.88 | 1.67E-47 | 7.89E-46 |
| Bex1 | 14.41 | 3.47 | 4.62E-47 | 2.15E-45 |
| Igfbp5 | 14.28 | 3.60 | 2.85E-46 | 1.29E-44 |
| Slc27a1 | 14.26 | 2.91 | 3.88E-46 | 1.75E-44 |
| Bmp7 | 14.22 | 2.58 | 7.25E-46 | 3.25E-44 |
| Slc41a1 | 14.19 | 3.12 | 1.13E-45 | 5.02E-44 |
| Csrp1 | 13.99 | 2.15 | 1.91E-44 | 8.34E-43 |
| Pycr1 | 13.88 | 3.21 | 8.84E-44 | 3.82E-42 |
| Asns | 13.76 | 2.49 | 4.35E-43 | 1.86E-41 |
| Grem2 | 13.72 | 5.27 | 7.85E-43 | 3.33E-41 |
| Lrig1 | 13.65 | 1.92 | 2.06E-42 | 8.61E-41 |
| C4bp | 13.58 | 3.80 | 5.52E-42 | 2.29E-40 |
| Mecom | 13.52 | 2.62 | 1.16E-41 | 4.75E-40 |
| Smim38 | 13.50 | 3.81 | 1.53E-41 | 6.21E-40 |
| Ltbp4 | 13.47 | 2.38 | 2.46E-41 | 9.96E-40 |
| Rcn1 | 13.44 | 2.63 | 3.42E-41 | 1.38E-39 |
| Kit | 13.44 | 2.97 | 3.68E-41 | 1.48E-39 |
| Vim | 13.19 | 1.87 | 9.4E-40 | 3.66E-38 |
| Foxa3 | 13.04 | 2.44 | 7.57E-39 | 2.88E-37 |
| Nkd1 | 12.99 | 4.01 | 1.33E-38 | 5.06E-37 |
| Htra3 | 12.94 | 3.17 | 2.53E-38 | 9.54E-37 |
| Msi1 | 12.88 | 3.48 | 5.95E-38 | 2.22E-36 |
| Cald1 | 12.87 | 2.12 | 6.79E-38 | 2.53E-36 |
| Tpd52l1 | 12.71 | 3.84 | 5.01E-37 | 1.84E-35 |
| Rap1gap | 12.70 | 2.16 | 6.07E-37 | 2.22E-35 |
| Vip | 12.68 | 4.18 | 8.11E-37 | 2.94E-35 |
| Gpx3 | 12.65 | 1.67 | 1.16E-36 | 4.18E-35 |
| Smim14 | 12.54 | 1.51 | 4.54E-36 | 1.62E-34 |
| Des | 12.50 | 2.23 | 7.13E-36 | 2.53E-34 |
| Map2k6 | 12.32 | 2.27 | 7.22E-35 | 2.51E-33 |
| Unc5a | 12.32 | 3.54 | 7.48E-35 | 2.6E-33 |

| | | | | |
|---|---|---|---|---|
| Bgn | 12.28 | 1.79 | 1.22E-34 | 4.22E-33 |
| Ifitm3 | 12.26 | 2.33 | 1.52E-34 | 5.25E-33 |
| Ephb3 | 12.21 | 3.13 | 2.66E-34 | 9.09E-33 |
| Grem1 | 12.06 | 3.66 | 1.72E-33 | 5.75E-32 |
| Sfrp1 | 12.03 | 2.88 | 2.61E-33 | 8.66E-32 |
| Cdca7 | 11.96 | 1.82 | 5.86E-33 | 1.92E-31 |
| Sidt1 | 11.95 | 2.80 | 6.62E-33 | 2.17E-31 |
| Aldh1l2 | 11.82 | 4.59 | 2.97E-32 | 9.54E-31 |
| Tnxb | 11.70 | 3.37 | 1.27E-31 | 3.99E-30 |
| Mansc1 | 11.60 | 2.12 | 3.9E-31 | 1.21E-29 |
| Tubb2b | 11.47 | 3.10 | 1.89E-30 | 5.71E-29 |
| Retnlb | 11.42 | 5.37 | 3.17E-30 | 9.56E-29 |
| Tmed9 | 11.33 | 1.52 | 9.3E-30 | 2.77E-28 |
| Slc50a1 | 11.31 | 1.94 | 1.17E-29 | 3.46E-28 |
| Sgsm3 | 11.18 | 1.98 | 4.89E-29 | 1.42E-27 |
| Gtf2i | 11.16 | 1.52 | 6.26E-29 | 1.81E-27 |
| Fzd9 | 10.97 | 4.66 | 5.44E-28 | 1.53E-26 |
| Calcrl | 10.97 | 2.56 | 5.62E-28 | 1.58E-26 |
| Stx17 | 10.94 | 1.76 | 7.17E-28 | 2E-26 |
| Rfc1 | 10.88 | 1.99 | 1.47E-27 | 4.04E-26 |
| Sorcs2 | 10.81 | 2.87 | 2.93E-27 | 8E-26 |
| Gadd45g | 10.79 | 2.23 | 3.9E-27 | 1.06E-25 |
| Slc38a1 | 10.78 | 1.73 | 4.38E-27 | 1.18E-25 |
| Arhgef38 | 10.76 | 1.61 | 5.15E-27 | 1.39E-25 |
| Iqsec1 | 10.71 | 2.21 | 9.41E-27 | 2.51E-25 |
| Ccdc3 | 10.67 | 2.94 | 1.4E-26 | 3.71E-25 |
| Ttc39a | 10.63 | 2.20 | 2.26E-26 | 5.96E-25 |
| Pla2g12a | 10.60 | 2.76 | 2.87E-26 | 7.53E-25 |
| Tmed3 | 10.49 | 1.51 | 9.14E-26 | 2.35E-24 |
| Prelp | 10.40 | 2.88 | 2.4E-25 | 6.08E-24 |
| Fam222a | 10.40 | 3.84 | 2.47E-25 | 6.25E-24 |
| Cgnl1 | 10.40 | 4.06 | 2.6E-25 | 6.56E-24 |
| Stk39 | 10.33 | 1.77 | 5.3E-25 | 1.32E-23 |
| Bex3 | 10.32 | 1.88 | 5.58E-25 | 1.39E-23 |
| Kdelr3 | 10.26 | 2.30 | 1.02E-24 | 2.51E-23 |
| Ces1d | 10.26 | 4.03 | 1.1E-24 | 2.7E-23 |
| Fkbp11 | 10.25 | 2.60 | 1.19E-24 | 2.91E-23 |
| Rassf4 | 10.24 | 1.82 | 1.28E-24 | 3.12E-23 |
| Arfgef3 | 10.22 | 1.78 | 1.56E-24 | 3.78E-23 |
| Ptp4a3 | 10.22 | 1.84 | 1.68E-24 | 4.07E-23 |
| Cxxc5 | 10.15 | 3.46 | 3.47E-24 | 8.26E-23 |
| Tnfrsf19 | 10.14 | 3.17 | 3.52E-24 | 8.35E-23 |
| Cbfa2t3 | 10.12 | 2.34 | 4.39E-24 | 1.03E-22 |
| Maged1 | 10.10 | 1.87 | 5.5E-24 | 1.29E-22 |

| | | | | |
|---|---|---|---|---|
| Myc | 10.09 | 1.57 | 6.14E-24 | 1.43E-22 |
| Pdia2 | 10.06 | 5.13 | 8.68E-24 | 2.02E-22 |
| Scn2b | 10.05 | 2.76 | 8.79E-24 | 2.04E-22 |
| Aqp4 | 10.04 | 2.31 | 1.04E-23 | 2.41E-22 |
| Cln6 | 10.01 | 2.01 | 1.45E-23 | 3.33E-22 |
| Acvr1c | 9.97 | 2.16 | 2.09E-23 | 4.76E-22 |
| Mcf2l | 9.96 | 2.19 | 2.25E-23 | 5.13E-22 |
| Dach1 | 9.95 | 2.01 | 2.48E-23 | 5.64E-22 |
| Fto | 9.95 | 1.99 | 2.55E-23 | 5.8E-22 |
| Spdef | 9.91 | 1.83 | 3.86E-23 | 8.71E-22 |
| Svep1 | 9.90 | 3.09 | 4.05E-23 | 9.13E-22 |
| Fam174b | 9.81 | 2.31 | 9.88E-23 | 2.21E-21 |
| C1qtnf3 | 9.79 | 3.34 | 1.23E-22 | 2.74E-21 |
| Map3k15 | 9.77 | 2.69 | 1.55E-22 | 3.43E-21 |
| Cdc42ep1 | 9.72 | 2.09 | 2.5E-22 | 5.52E-21 |
| Mlph | 9.72 | 1.66 | 2.52E-22 | 5.55E-21 |
| Armcx3 | 9.68 | 2.55 | 3.52E-22 | 7.71E-21 |
| Ints6l | 9.68 | 4.15 | 3.68E-22 | 8.04E-21 |
| Cygb | 9.63 | 2.53 | 6.11E-22 | 1.33E-20 |
| Serping1 | 9.52 | 1.79 | 1.77E-21 | 3.78E-20 |
| Pck2 | 9.52 | 1.79 | 1.79E-21 | 3.8E-20 |
| Cdk6 | 9.50 | 1.83 | 2.16E-21 | 4.57E-20 |
| Cavin1 | 9.44 | 2.45 | 3.73E-21 | 7.81E-20 |
| Ppan | 9.39 | 1.80 | 5.78E-21 | 1.2E-19 |
| Fam129b | 9.38 | 1.68 | 6.67E-21 | 1.38E-19 |
| Cybrd1 | 9.36 | 3.78 | 8.24E-21 | 1.69E-19 |
| Ggh | 9.35 | 2.74 | 9.02E-21 | 1.85E-19 |
| Clec3b | 9.34 | 2.96 | 9.7E-21 | 1.98E-19 |
| Noxa1 | 9.32 | 1.65 | 1.19E-20 | 2.42E-19 |
| Mthfd2 | 9.24 | 2.30 | 2.36E-20 | 4.75E-19 |
| Thbs1 | 9.24 | 2.21 | 2.52E-20 | 5.05E-19 |
| Slit3 | 9.16 | 4.17 | 5.32E-20 | 1.06E-18 |
| Pgm5 | 9.16 | 2.54 | 5.34E-20 | 1.06E-18 |
| Cd44 | 9.15 | 1.89 | 5.6E-20 | 1.11E-18 |
| Zc3h7b | 9.13 | 1.88 | 6.59E-20 | 1.3E-18 |
| Paip2b | 9.10 | 2.03 | 8.75E-20 | 1.71E-18 |
| Cbx3 | 9.09 | 1.78 | 1E-19 | 1.96E-18 |
| Inf2 | 9.07 | 2.57 | 1.19E-19 | 2.31E-18 |
| Timp3 | 9.06 | 2.06 | 1.25E-19 | 2.42E-18 |
| Tgfbr2 | 9.06 | 1.70 | 1.28E-19 | 2.49E-18 |
| Rasef | 9.06 | 1.54 | 1.34E-19 | 2.58E-18 |
| Lzts2 | 9.04 | 1.52 | 1.54E-19 | 2.96E-18 |
| Plch2 | 9.03 | 1.62 | 1.64E-19 | 3.16E-18 |
| Creb3l4 | 9.01 | 2.14 | 2.13E-19 | 4.07E-18 |

| | | | | |
|---|---|---|---|---|
| Naaladl2 | 9.01 | 3.31 | 2.16E-19 | 4.12E-18 |
| Ror2 | 9.00 | 3.47 | 2.29E-19 | 4.38E-18 |
| Slc43a1 | 8.98 | 3.18 | 2.66E-19 | 5.04E-18 |
| Serpina3n | 8.98 | 3.11 | 2.69E-19 | 5.1E-18 |
| Smyd2 | 8.97 | 1.64 | 3.02E-19 | 5.7E-18 |
| Qtrt1 | 8.96 | 2.09 | 3.15E-19 | 5.94E-18 |
| Sytl2 | 8.96 | 1.72 | 3.23E-19 | 6.08E-18 |
| Cnn1 | 8.92 | 2.38 | 4.84E-19 | 9.06E-18 |
| Lama5 | 8.91 | 2.32 | 5E-19 | 9.34E-18 |
| Casc4 | 8.91 | 2.32 | 5.17E-19 | 9.63E-18 |
| Cep78 | 8.90 | 2.70 | 5.47E-19 | 1.02E-17 |
| Lum | 8.85 | 3.45 | 8.83E-19 | 1.62E-17 |
| Dmpk | 8.82 | 2.07 | 1.14E-18 | 2.07E-17 |
| Hcfc2 | 8.82 | 1.76 | 1.19E-18 | 2.16E-17 |
| Sparcl1 | 8.80 | 1.94 | 1.4E-18 | 2.55E-17 |
| Trim27 | 8.79 | 1.54 | 1.49E-18 | 2.7E-17 |
| Slco3a1 | 8.79 | 1.71 | 1.56E-18 | 2.82E-17 |
| Rnase1 | 8.78 | 1.97 | 1.68E-18 | 3.05E-17 |
| Ackr4 | 8.77 | 4.72 | 1.82E-18 | 3.28E-17 |
| Mfap4 | 8.76 | 2.77 | 1.89E-18 | 3.42E-17 |
| Kif21a | 8.76 | 2.06 | 1.94E-18 | 3.48E-17 |
| Insrr | 8.76 | 3.76 | 1.96E-18 | 3.52E-17 |
| Ppic | 8.74 | 2.37 | 2.29E-18 | 4.09E-17 |
| Crispld2 | 8.71 | 2.39 | 2.92E-18 | 5.19E-17 |
| Fmo2 | 8.71 | 2.64 | 3.01E-18 | 5.34E-17 |
| Ptpro | 8.70 | 2.95 | 3.32E-18 | 5.87E-17 |
| Timp2 | 8.66 | 1.94 | 4.71E-18 | 8.25E-17 |
| Myo1b | 8.65 | 2.09 | 5.23E-18 | 9.13E-17 |
| Rhoj | 8.62 | 2.65 | 6.72E-18 | 1.17E-16 |
| Zdhhc14 | 8.61 | 2.44 | 7.07E-18 | 1.23E-16 |
| C1rl | 8.60 | 4.07 | 7.79E-18 | 1.35E-16 |
| Enpp2 | 8.60 | 3.98 | 7.86E-18 | 1.36E-16 |
| Fn1 | 8.58 | 1.81 | 9.89E-18 | 1.7E-16 |
| Nfasc | 8.54 | 3.35 | 1.39E-17 | 2.35E-16 |
| Gpam | 8.53 | 2.23 | 1.44E-17 | 2.42E-16 |
| Pola1 | 8.52 | 1.74 | 1.67E-17 | 2.8E-16 |
| Rarres2 | 8.48 | 1.83 | 2.31E-17 | 3.86E-16 |
| Hid1 | 8.45 | 1.75 | 2.82E-17 | 4.69E-16 |
| Rdh1 | 8.45 | 3.30 | 2.83E-17 | 4.7E-16 |
| Mmrn1 | 8.45 | 3.22 | 2.87E-17 | 4.77E-16 |
| Tgif1 | 8.44 | 1.58 | 3.14E-17 | 5.2E-16 |
| Nrbp2 | 8.44 | 2.07 | 3.3E-17 | 5.46E-16 |
| Ung | 8.41 | 1.85 | 4.01E-17 | 6.6E-16 |
| Hgfac | 8.40 | 1.89 | 4.46E-17 | 7.3E-16 |

| | | | | |
|---|---|---|---|---|
| Aspn | 8.40 | 2.92 | 4.52E-17 | 7.4E-16 |
| Srm | 8.39 | 1.78 | 4.8E-17 | 7.83E-16 |
| Nfix | 8.37 | 1.75 | 5.96E-17 | 9.7E-16 |
| Klk1 | 8.36 | 1.51 | 6.2E-17 | 1.01E-15 |
| Tshz2 | 8.36 | 2.43 | 6.51E-17 | 1.06E-15 |
| Fndc3b | 8.35 | 1.68 | 6.74E-17 | 1.09E-15 |
| Slc16a7 | 8.34 | 4.01 | 7.68E-17 | 1.24E-15 |
| Tmem63a | 8.28 | 1.57 | 1.25E-16 | 2E-15 |
| Gpt2 | 8.25 | 2.77 | 1.62E-16 | 2.58E-15 |
| Igfbp6 | 8.22 | 4.13 | 2.04E-16 | 3.25E-15 |
| St6galnac2 | 8.21 | 1.52 | 2.23E-16 | 3.53E-15 |
| Spock2 | 8.20 | 4.78 | 2.35E-16 | 3.71E-15 |
| Cd244a | 8.20 | 2.30 | 2.36E-16 | 3.73E-15 |
| Cad | 8.20 | 1.75 | 2.42E-16 | 3.82E-15 |
| Clca3a2 | 8.19 | 1.66 | 2.57E-16 | 4.05E-15 |
| Hspb8 | 8.18 | 3.73 | 2.78E-16 | 4.37E-15 |
| Cd34 | 8.14 | 2.46 | 3.88E-16 | 6.07E-15 |
| Aoc3 | 8.14 | 2.80 | 3.9E-16 | 6.1E-15 |
| Ccpg1 | 8.14 | 1.72 | 4.05E-16 | 6.32E-15 |
| Sirpa | 8.13 | 2.69 | 4.21E-16 | 6.57E-15 |
| Agt | 8.13 | 2.68 | 4.24E-16 | 6.61E-15 |
| Lix1l | 8.13 | 2.52 | 4.4E-16 | 6.84E-15 |
| Far1 | 8.10 | 2.20 | 5.39E-16 | 8.36E-15 |
| Mnx1 | 8.09 | 2.98 | 5.91E-16 | 9.15E-15 |
| Mamdc2 | 8.06 | 3.59 | 7.65E-16 | 1.18E-14 |
| Igdcc4 | 8.05 | 2.99 | 8.09E-16 | 1.24E-14 |
| Postn | 8.05 | 2.54 | 8.46E-16 | 1.3E-14 |
| Aebp1 | 8.04 | 3.30 | 8.77E-16 | 1.34E-14 |
| Slc7a5 | 7.99 | 2.43 | 1.33E-15 | 2.01E-14 |
| Sox4 | 7.97 | 2.01 | 1.55E-15 | 2.35E-14 |
| Rbpms | 7.97 | 1.88 | 1.65E-15 | 2.49E-14 |
| Rcan3 | 7.94 | 2.22 | 2.04E-15 | 3.07E-14 |
| Sorbs2 | 7.94 | 2.43 | 2.06E-15 | 3.09E-14 |
| Slain1 | 7.92 | 2.27 | 2.38E-15 | 3.56E-14 |
| Eln | 7.92 | 2.95 | 2.42E-15 | 3.61E-14 |
| Large2 | 7.92 | 2.95 | 2.45E-15 | 3.66E-14 |
| Sgcd | 7.92 | 2.91 | 2.46E-15 | 3.66E-14 |
| Cracr2b | 7.91 | 1.74 | 2.56E-15 | 3.82E-14 |
| Cited4 | 7.91 | 3.14 | 2.63E-15 | 3.91E-14 |
| Cyp20a1 | 7.87 | 2.42 | 3.54E-15 | 5.23E-14 |
| Man2a2 | 7.86 | 1.96 | 3.79E-15 | 5.6E-14 |
| Tgfbr3 | 7.86 | 2.71 | 3.85E-15 | 5.67E-14 |
| Marveld1 | 7.83 | 2.69 | 4.71E-15 | 6.9E-14 |
| Wnt2b | 7.82 | 2.61 | 5.2E-15 | 7.6E-14 |

| | | | | |
|---|---|---|---|---|
| Ceacam10 | 7.80 | 2.16 | 6.38E-15 | 9.3E-14 |
| Slc39a8 | 7.76 | 2.12 | 8.74E-15 | 1.26E-13 |
| Ivd | 7.74 | 1.55 | 9.8E-15 | 1.41E-13 |
| Slc12a8 | 7.74 | 1.73 | 1.02E-14 | 1.47E-13 |
| Atxn10 | 7.73 | 2.07 | 1.07E-14 | 1.53E-13 |
| Epb42 | 7.73 | 4.16 | 1.12E-14 | 1.6E-13 |
| Arl3 | 7.72 | 2.37 | 1.16E-14 | 1.66E-13 |
| Mcc | 7.69 | 2.44 | 1.52E-14 | 2.16E-13 |
| Trim44 | 7.68 | 1.69 | 1.56E-14 | 2.22E-13 |
| Scn1b | 7.66 | 2.85 | 1.86E-14 | 2.63E-13 |
| Dbn1 | 7.66 | 2.89 | 1.87E-14 | 2.65E-13 |
| Slc25a23 | 7.65 | 1.97 | 1.99E-14 | 2.81E-13 |
| Ces1c | 7.64 | 3.63 | 2.2E-14 | 3.11E-13 |
| Scn7a | 7.64 | 3.55 | 2.21E-14 | 3.11E-13 |
| Colca2 | 7.64 | 2.67 | 2.24E-14 | 3.16E-13 |
| Cxcl12 | 7.61 | 1.88 | 2.82E-14 | 3.95E-13 |
| Nav2 | 7.59 | 2.06 | 3.19E-14 | 4.45E-13 |
| Tle2 | 7.58 | 2.84 | 3.46E-14 | 4.82E-13 |
| Zfp428 | 7.55 | 2.80 | 4.26E-14 | 5.91E-13 |
| Tns1 | 7.52 | 2.48 | 5.35E-14 | 7.37E-13 |
| Lepr | 7.52 | 3.10 | 5.61E-14 | 7.71E-13 |
| Eef1akmt3 | 7.51 | 2.97 | 5.72E-14 | 7.85E-13 |
| Lmod1 | 7.51 | 2.65 | 5.79E-14 | 7.95E-13 |
| Glul | 7.51 | 1.84 | 5.84E-14 | 8.01E-13 |
| Klf2 | 7.51 | 2.54 | 5.97E-14 | 8.18E-13 |
| Iffo2 | 7.51 | 2.63 | 5.98E-14 | 8.19E-13 |
| Myrip | 7.51 | 2.61 | 6.08E-14 | 8.32E-13 |
| Ass1 | 7.50 | 2.62 | 6.16E-14 | 8.42E-13 |
| Akap12 | 7.50 | 2.10 | 6.3E-14 | 8.61E-13 |
| Cavin2 | 7.48 | 2.84 | 7.72E-14 | 1.05E-12 |
| Auts2 | 7.45 | 1.71 | 9.18E-14 | 1.24E-12 |
| Ggnbp1 | 7.45 | 2.94 | 9.31E-14 | 1.26E-12 |
| Greb1 | 7.45 | 2.78 | 9.59E-14 | 1.29E-12 |
| Ghr | 7.43 | 1.54 | 1.05E-13 | 1.41E-12 |
| Wfikkn2 | 7.43 | 3.82 | 1.1E-13 | 1.48E-12 |
| Plcb4 | 7.40 | 2.63 | 1.38E-13 | 1.85E-12 |
| Psd | 7.38 | 3.16 | 1.64E-13 | 2.17E-12 |
| Whrn | 7.36 | 1.90 | 1.81E-13 | 2.39E-12 |
| Piezo2 | 7.36 | 3.51 | 1.91E-13 | 2.51E-12 |
| Adamts2 | 7.32 | 3.97 | 2.44E-13 | 3.19E-12 |
| Kcnh2 | 7.30 | 2.67 | 2.86E-13 | 3.71E-12 |
| Enpp5 | 7.29 | 2.76 | 3.03E-13 | 3.93E-12 |
| Bcat2 | 7.29 | 1.67 | 3.06E-13 | 3.96E-12 |
| Akr1b10 | 7.29 | 2.67 | 3.08E-13 | 3.98E-12 |

| | | | | |
|---|---|---|---|---|
| Apod | 7.29 | 4.42 | 3.15E-13 | 4.07E-12 |
| Slc39a13 | 7.29 | 2.60 | 3.19E-13 | 4.12E-12 |
| Esam | 7.29 | 2.54 | 3.21E-13 | 4.15E-12 |
| Homer2 | 7.28 | 1.58 | 3.44E-13 | 4.43E-12 |
| Celsr2 | 7.27 | 3.59 | 3.48E-13 | 4.47E-12 |
| Cpe | 7.23 | 2.28 | 4.76E-13 | 6.08E-12 |
| Scmh1 | 7.22 | 2.09 | 5.26E-13 | 6.7E-12 |
| Slc14a1 | 7.21 | 2.85 | 5.57E-13 | 7.07E-12 |
| Cdk17 | 7.20 | 2.07 | 5.94E-13 | 7.51E-12 |
| Nipal2 | 7.20 | 1.66 | 6.1E-13 | 7.7E-12 |
| Lgals12 | 7.19 | 2.66 | 6.49E-13 | 8.17E-12 |
| Pycr2 | 7.19 | 1.86 | 6.59E-13 | 8.29E-12 |
| Gabbr1 | 7.16 | 2.24 | 8.35E-13 | 1.05E-11 |
| Slc48a1 | 7.16 | 1.67 | 8.36E-13 | 1.05E-11 |
| Anxa5 | 7.14 | 1.56 | 9.03E-13 | 1.13E-11 |
| Haus4 | 7.14 | 1.56 | 9.5E-13 | 1.18E-11 |
| Rora | 7.11 | 2.41 | 1.16E-12 | 1.45E-11 |
| Casp12 | 7.10 | 2.30 | 1.22E-12 | 1.51E-11 |
| Apcdd1 | 7.10 | 3.16 | 1.25E-12 | 1.54E-11 |
| Impdh1 | 7.09 | 1.92 | 1.34E-12 | 1.64E-11 |
| Tex30 | 7.09 | 1.94 | 1.38E-12 | 1.69E-11 |
| Pdlim7 | 7.08 | 2.11 | 1.39E-12 | 1.71E-11 |
| Tmem109 | 7.08 | 1.74 | 1.41E-12 | 1.73E-11 |
| Dagla | 7.08 | 2.37 | 1.47E-12 | 1.79E-11 |
| Pcolce2 | 7.07 | 23.38 | 1.52E-12 | 1.86E-11 |
| Gramd1a | 7.07 | 1.96 | 1.58E-12 | 1.93E-11 |
| Glt1d1 | 7.06 | 3.99 | 1.66E-12 | 2.02E-11 |
| Trit1 | 7.05 | 1.57 | 1.79E-12 | 2.17E-11 |
| Mpp3 | 7.05 | 2.94 | 1.81E-12 | 2.19E-11 |
| Dennd5a | 7.02 | 2.27 | 2.28E-12 | 2.75E-11 |
| Shmt2 | 7.01 | 1.62 | 2.31E-12 | 2.78E-11 |
| Ankrd22 | 7.00 | 4.27 | 2.48E-12 | 2.99E-11 |
| Vav3 | 6.99 | 3.06 | 2.69E-12 | 3.24E-11 |
| Col16a1 | 6.98 | 3.43 | 3.03E-12 | 3.63E-11 |
| Fn3k | 6.97 | 5.94 | 3.1E-12 | 3.72E-11 |
| Adi1 | 6.97 | 1.56 | 3.27E-12 | 3.92E-11 |
| Cdo1 | 6.96 | 1.77 | 3.29E-12 | 3.94E-11 |
| Cbs | 6.96 | 4.37 | 3.4E-12 | 4.05E-11 |
| Bhmt2 | 6.96 | 4.25 | 3.4E-12 | 4.06E-11 |
| Sod3 | 6.96 | 2.13 | 3.47E-12 | 4.13E-11 |
| Plcb1 | 6.95 | 2.49 | 3.71E-12 | 4.41E-11 |
| Adra2a | 6.94 | 2.40 | 3.83E-12 | 4.54E-11 |
| Adgrg1 | 6.93 | 2.19 | 4.19E-12 | 4.97E-11 |
| Tcof1 | 6.92 | 1.54 | 4.42E-12 | 5.22E-11 |

| | | | | |
|---|---|---|---|---|
| Fkbp9 | 6.90 | 2.22 | 5.12E-12 | 6.02E-11 |
| Ascl2 | 6.89 | 2.27 | 5.42E-12 | 6.35E-11 |
| Tmem131l | 6.89 | 2.09 | 5.43E-12 | 6.35E-11 |
| Synj2 | 6.89 | 1.66 | 5.54E-12 | 6.48E-11 |
| Nup210 | 6.88 | 1.61 | 5.81E-12 | 6.79E-11 |
| Vldlr | 6.88 | 2.88 | 5.82E-12 | 6.79E-11 |
| Slc19a1 | 6.88 | 1.57 | 6.18E-12 | 7.19E-11 |
| Pkd2 | 6.87 | 1.93 | 6.53E-12 | 7.59E-11 |
| Znrf3 | 6.87 | 1.74 | 6.55E-12 | 7.6E-11 |
| Bckdhb | 6.86 | 2.03 | 6.92E-12 | 8.01E-11 |
| Acadsb | 6.85 | 1.78 | 7.54E-12 | 8.71E-11 |
| Prim1 | 6.80 | 1.78 | 1.08E-11 | 1.23E-10 |
| C1s1 | 6.79 | 1.63 | 1.09E-11 | 1.24E-10 |
| Zfp367 | 6.77 | 1.68 | 1.32E-11 | 1.49E-10 |
| Synm | 6.76 | 2.31 | 1.39E-11 | 1.57E-10 |
| Hs3st3b1 | 6.75 | 2.76 | 1.45E-11 | 1.64E-10 |
| Pxdn | 6.75 | 2.02 | 1.47E-11 | 1.65E-10 |
| Thsd7a | 6.74 | 3.76 | 1.61E-11 | 1.81E-10 |
| Adpgk | 6.69 | 1.59 | 2.21E-11 | 2.45E-10 |
| Tacc1 | 6.69 | 2.02 | 2.28E-11 | 2.53E-10 |
| Arhgap39 | 6.68 | 1.54 | 2.36E-11 | 2.61E-10 |
| Slc29a1 | 6.68 | 1.93 | 2.43E-11 | 2.68E-10 |
| Itih5 | 6.67 | 1.68 | 2.5E-11 | 2.76E-10 |
| Esrrg | 6.67 | 2.17 | 2.59E-11 | 2.85E-10 |
| Rnf32 | 6.65 | 1.91 | 2.89E-11 | 3.18E-10 |
| Nav1 | 6.65 | 1.78 | 2.98E-11 | 3.26E-10 |
| Tbc1d16 | 6.64 | 2.33 | 3.09E-11 | 3.38E-10 |
| Nudcd1 | 6.63 | 1.65 | 3.44E-11 | 3.75E-10 |
| Soga3 | 6.62 | 4.42 | 3.59E-11 | 3.92E-10 |
| Wnk4 | 6.61 | 1.92 | 3.86E-11 | 4.2E-10 |
| Rapgef3 | 6.60 | 2.41 | 4.21E-11 | 4.57E-10 |
| Erich5 | 6.58 | 4.60 | 4.57E-11 | 4.95E-10 |
| Dkk2 | 6.58 | 4.42 | 4.57E-11 | 4.95E-10 |
| Sulf1 | 6.58 | 3.48 | 4.77E-11 | 5.16E-10 |
| Col5a1 | 6.53 | 2.18 | 6.39E-11 | 6.86E-10 |
| Nt5c3b | 6.50 | 2.10 | 8.15E-11 | 8.69E-10 |
| Ptprs | 6.49 | 1.51 | 8.58E-11 | 9.14E-10 |
| Maged2 | 6.46 | 2.13 | 1.02E-10 | 1.08E-09 |
| Serpinh1 | 6.46 | 1.60 | 1.08E-10 | 1.14E-09 |
| Id4 | 6.45 | 2.04 | 1.13E-10 | 1.19E-09 |
| Mthfd1l | 6.45 | 1.82 | 1.15E-10 | 1.21E-09 |
| Lfng | 6.45 | 2.06 | 1.15E-10 | 1.22E-09 |
| Mcm3 | 6.45 | 1.52 | 1.15E-10 | 1.22E-09 |
| Efna4 | 6.44 | 1.59 | 1.19E-10 | 1.25E-09 |

| | | | | |
|---|---|---|---|---|
| Filip1l | 6.44 | 2.00 | 1.23E-10 | 1.3E-09 |
| Gstm2 | 6.43 | 1.84 | 1.25E-10 | 1.31E-09 |
| Slc7a4 | 6.41 | 1.85 | 1.44E-10 | 1.51E-09 |
| Nbeal2 | 6.41 | 1.59 | 1.49E-10 | 1.56E-09 |
| Lonrf3 | 6.41 | 1.98 | 1.49E-10 | 1.56E-09 |
| Acss3 | 6.40 | 3.69 | 1.52E-10 | 1.59E-09 |
| Hells | 6.40 | 1.53 | 1.53E-10 | 1.59E-09 |
| Tceal8 | 6.40 | 1.55 | 1.56E-10 | 1.63E-09 |
| Sdc3 | 6.39 | 2.79 | 1.63E-10 | 1.7E-09 |
| Inha | 6.35 | 3.26 | 2.09E-10 | 2.16E-09 |
| Itfg2 | 6.35 | 1.51 | 2.18E-10 | 2.24E-09 |
| Sil1 | 6.35 | 1.52 | 2.18E-10 | 2.25E-09 |
| Sema4c | 6.35 | 2.60 | 2.2E-10 | 2.26E-09 |
| Fzd2 | 6.35 | 2.67 | 2.22E-10 | 2.29E-09 |
| Tmem9 | 6.34 | 1.76 | 2.26E-10 | 2.33E-09 |
| Zbtb16 | 6.34 | 2.57 | 2.27E-10 | 2.33E-09 |
| Slc9a3r2 | 6.34 | 1.68 | 2.28E-10 | 2.34E-09 |
| Abcc1 | 6.34 | 2.02 | 2.32E-10 | 2.38E-09 |
| Cib2 | 6.34 | 2.52 | 2.33E-10 | 2.39E-09 |
| Pbx3 | 6.34 | 2.56 | 2.35E-10 | 2.4E-09 |
| Fhl5 | 6.33 | 4.00 | 2.48E-10 | 2.53E-09 |
| Stxbp6 | 6.30 | 1.54 | 2.91E-10 | 2.96E-09 |
| Lysmd2 | 6.30 | 1.74 | 3.07E-10 | 3.11E-09 |
| Pcolce | 6.28 | 2.21 | 3.37E-10 | 3.4E-09 |
| Cyb5r1 | 6.27 | 1.68 | 3.55E-10 | 3.58E-09 |
| Slc7a6 | 6.27 | 2.40 | 3.56E-10 | 3.58E-09 |
| Sema3g | 6.26 | 3.23 | 3.83E-10 | 3.85E-09 |
| Synpo2 | 6.26 | 1.76 | 3.86E-10 | 3.88E-09 |
| Arntl2 | 6.25 | 3.42 | 4.05E-10 | 4.06E-09 |
| Vmac | 6.25 | 2.02 | 4.09E-10 | 4.1E-09 |
| Zdhhc1 | 6.25 | 2.30 | 4.11E-10 | 4.12E-09 |
| Casd1 | 6.23 | 1.69 | 4.72E-10 | 4.71E-09 |
| Syn2 | 6.22 | 5.77 | 4.85E-10 | 4.84E-09 |
| Derl3 | 6.22 | 2.65 | 4.98E-10 | 4.96E-09 |
| Zfhx3 | 6.22 | 2.30 | 5.03E-10 | 5E-09 |
| Pprc1 | 6.21 | 1.60 | 5.35E-10 | 5.31E-09 |
| Nod1 | 6.21 | 1.97 | 5.43E-10 | 5.39E-09 |
| Agtr1a | 6.20 | 5.02 | 5.69E-10 | 5.63E-09 |
| Gfra2 | 6.20 | 4.86 | 5.7E-10 | 5.64E-09 |
| Tmem191c | 6.19 | 2.10 | 6.14E-10 | 6.06E-09 |
| Rasa3 | 6.16 | 1.62 | 7.27E-10 | 7.13E-09 |
| Prps2 | 6.16 | 1.57 | 7.4E-10 | 7.25E-09 |
| A4gnt | 6.15 | 3.58 | 7.63E-10 | 7.47E-09 |
| Bcl7a | 6.15 | 1.77 | 7.77E-10 | 7.61E-09 |

| | | | | |
|---|---|---|---|---|
| Scgn | 6.14 | 3.24 | 8.01E-10 | 7.83E-09 |
| Ifi27l2a | 6.13 | 2.01 | 8.82E-10 | 8.6E-09 |
| Ntn1 | 6.12 | 2.38 | 9.29E-10 | 9.03E-09 |

Zone 2: Significantly Zonated Genes

| Symbol | scores | LFC | pvals | pvals_adj |
|---|---|---|---|---|
| Lgals2 | 56.04 | 3.34 | 0 | 0 |
| Dmbt1 | 52.92 | 3.20 | 0 | 0 |
| Hist1h2ap | 44.63 | 3.16 | 0 | 0 |
| Cps1 | 42.77 | 2.83 | 0 | 0 |
| Pycard | 42.76 | 2.65 | 0 | 0 |
| Aldh1b1 | 42.40 | 2.55 | 0 | 0 |
| Agr2 | 42.23 | 2.63 | 0 | 0 |
| Clca3b | 41.09 | 2.79 | 0 | 0 |
| Gvin1 | 41.04 | 3.04 | 0 | 0 |
| Eef1a1 | 38.73 | 2.35 | 0 | 0 |
| Krt19 | 38.24 | 2.43 | 0 | 0 |
| Stmn1 | 38.23 | 2.92 | 0 | 0 |
| Ivns1abp | 37.60 | 2.30 | 0 | 2E-306 |
| Spink4 | 34.96 | 2.19 | 8.7E-268 | 6.1E-265 |
| Mki67 | 33.91 | 3.03 | 4.1E-252 | 2.7E-249 |
| Ifitm2 | 33.19 | 2.44 | 1.6E-241 | 9.6E-239 |
| Rack1 | 32.69 | 2.21 | 1.9E-234 | 1.1E-231 |
| Hist1h1e | 32.28 | 2.21 | 1.5E-228 | 7.9E-226 |
| Cdca3 | 30.99 | 3.13 | 8.3E-211 | 4.3E-208 |
| Ppp1r1b | 30.58 | 2.03 | 2.2E-205 | 1E-202 |
| Top2a | 30.38 | 2.98 | 9.5E-203 | 4.2E-200 |
| Cluh | 30.19 | 2.66 | 2.9E-200 | 1.2E-197 |
| Olfm4 | 29.60 | 1.92 | 1.4E-192 | 5.6E-190 |
| Tk1 | 28.81 | 3.17 | 1.8E-182 | 6.8E-180 |
| Eef1b2 | 28.56 | 2.01 | 1.9E-179 | 6.8E-177 |
| Myb | 28.37 | 2.64 | 4.9E-177 | 1.7E-174 |
| Oat | 28.27 | 1.86 | 8.7E-176 | 3E-173 |
| Ccnd2 | 28.27 | 2.47 | 9E-176 | 3E-173 |
| Kcnq1 | 28.03 | 2.42 | 8E-173 | 2.6E-170 |
| Tubb5 | 27.61 | 2.63 | 8.2E-168 | 2.6E-165 |
| Uhrf1 | 27.59 | 3.00 | 1.4E-167 | 4.4E-165 |
| Hist1h1b | 27.46 | 2.91 | 5.8E-166 | 1.8E-163 |
| Rbbp7 | 27.44 | 2.37 | 9.1E-166 | 2.8E-163 |
| Gmds | 27.30 | 1.84 | 4.2E-164 | 1.2E-161 |
| Akt1 | 27.19 | 1.97 | 9E-163 | 2.6E-160 |
| Lmnb1 | 26.99 | 2.67 | 1.9E-160 | 5.2E-158 |
| Mlec | 26.95 | 1.77 | 5.4E-160 | 1.4E-157 |
| H2afv | 26.90 | 1.72 | 2E-159 | 5.3E-157 |
| Gpx1 | 26.61 | 1.93 | 5E-156 | 1.3E-153 |
| Stard10 | 26.58 | 1.67 | 1.1E-155 | 2.7E-153 |
| Srsf1 | 26.49 | 2.05 | 1.1E-154 | 2.8E-152 |

| | | | | |
|---|---|---|---|---|
| Lyz1 | 26.48 | 1.69 | 1.6E-154 | 3.9E-152 |
| Csrp2 | 26.40 | 2.69 | 1.4E-153 | 3.3E-151 |
| Tomm5 | 26.25 | 2.59 | 7.1E-152 | 1.7E-149 |
| Uqcr11 | 26.22 | 1.74 | 1.6E-151 | 3.8E-149 |
| Lsm4 | 26.17 | 2.20 | 6.4E-151 | 1.5E-148 |
| Ube2c | 26.03 | 2.93 | 2.3E-149 | 5.3E-147 |
| Shmt1 | 25.39 | 2.79 | 3.2E-142 | 6.9E-140 |
| Uck2 | 25.36 | 2.28 | 6.4E-142 | 1.4E-139 |
| Tuba1b | 25.33 | 2.66 | 1.3E-141 | 2.8E-139 |
| Sf3b3 | 25.31 | 2.25 | 2.5E-141 | 5.3E-139 |
| Elf3 | 25.29 | 1.65 | 3.8E-141 | 7.7E-139 |
| Hnrnpa1 | 25.25 | 2.31 | 1.1E-140 | 2.1E-138 |
| Ybx3 | 25.20 | 2.11 | 3.8E-140 | 7.4E-138 |
| Txn1 | 25.18 | 1.60 | 6.1E-140 | 1.2E-137 |
| Kcne3 | 25.15 | 2.86 | 1.3E-139 | 2.6E-137 |
| Plp2 | 25.11 | 2.44 | 3.5E-139 | 6.8E-137 |
| Lsm2 | 25.09 | 2.64 | 6.3E-139 | 1.2E-136 |
| Irf2bp2 | 25.07 | 1.83 | 1.1E-138 | 2.1E-136 |
| Defa21 | 24.98 | 1.61 | 1.1E-137 | 2E-135 |
| Rpn1 | 24.97 | 1.64 | 1.3E-137 | 2.4E-135 |
| Snrpf | 24.81 | 2.48 | 7.6E-136 | 1.4E-133 |
| Lsm7 | 24.45 | 2.52 | 5.4E-132 | 9.5E-130 |
| Hspd1 | 24.43 | 2.88 | 8.1E-132 | 1.4E-129 |
| Hnrnpm | 24.34 | 2.00 | 8.2E-131 | 1.4E-128 |
| Tfrc | 24.32 | 2.32 | 1.2E-130 | 2.1E-128 |
| Nop56 | 24.22 | 2.65 | 1.4E-129 | 2.4E-127 |
| Hist1h2ae | 24.21 | 3.09 | 1.6E-129 | 2.7E-127 |
| Nop58 | 24.15 | 2.44 | 8.1E-129 | 1.3E-126 |
| Itln1 | 24.00 | 1.52 | 2.7E-127 | 4.4E-125 |
| Cotl1 | 24.00 | 2.11 | 3E-127 | 4.9E-125 |
| Zfp36l2 | 23.95 | 1.69 | 9.8E-127 | 1.6E-124 |
| Slc12a2 | 23.94 | 2.02 | 1.2E-126 | 2E-124 |
| Hist1h1a | 23.90 | 2.57 | 2.9E-126 | 4.5E-124 |
| Ptma | 23.89 | 2.42 | 4.1E-126 | 6.3E-124 |
| Ranbp1 | 23.83 | 2.56 | 1.7E-125 | 2.6E-123 |
| Npm1 | 23.75 | 2.23 | 1E-124 | 1.5E-122 |
| Fgfbp1 | 23.63 | 2.39 | 2E-123 | 3E-121 |
| Gsr | 23.63 | 1.66 | 2E-123 | 3E-121 |
| Birc5 | 23.61 | 3.07 | 3.3E-123 | 4.8E-121 |
| Set | 23.59 | 2.26 | 4.7E-123 | 6.7E-121 |
| Vars | 23.58 | 2.50 | 6.4E-123 | 9.1E-121 |
| Kcnn4 | 23.56 | 2.75 | 9.4E-123 | 1.3E-120 |
| Atic | 23.50 | 2.46 | 4.5E-122 | 6.3E-120 |
| Srsf6 | 23.47 | 1.89 | 8.8E-122 | 1.2E-119 |

| | | | | |
|---|---|---|---|---|
| Snrpd2 | 23.35 | 2.39 | 1.3E-120 | 1.8E-118 |
| Pdcd4 | 23.30 | 1.99 | 4.2E-120 | 5.7E-118 |
| Srsf7 | 23.21 | 2.07 | 3.7E-119 | 4.9E-117 |
| Ssb | 23.19 | 1.79 | 6.4E-119 | 8.6E-117 |
| Mlxipl | 23.14 | 1.70 | 1.9E-118 | 2.6E-116 |
| Cdk1 | 23.08 | 3.13 | 8.1E-118 | 1.1E-115 |
| Ddost | 23.03 | 1.54 | 2.3E-117 | 3E-115 |
| Racgap1 | 22.98 | 3.05 | 6.6E-117 | 8.5E-115 |
| Eif3i | 22.86 | 2.01 | 1.3E-115 | 1.6E-113 |
| Nhp2 | 22.85 | 2.49 | 1.4E-115 | 1.8E-113 |
| Park7 | 22.84 | 2.05 | 1.7E-115 | 2.2E-113 |
| Snu13 | 22.84 | 2.49 | 1.8E-115 | 2.3E-113 |
| Anp32e | 22.76 | 2.32 | 1.1E-114 | 1.3E-112 |
| Tcp1 | 22.76 | 2.29 | 1.3E-114 | 1.5E-112 |
| H1f0 | 22.74 | 1.50 | 1.9E-114 | 2.3E-112 |
| Cdx1 | 22.70 | 1.72 | 4.3E-114 | 5.2E-112 |
| Naa50 | 22.62 | 2.24 | 2.7E-113 | 3.2E-111 |
| Timm8a1 | 22.62 | 2.50 | 2.9E-113 | 3.4E-111 |
| Mad2l1 | 22.61 | 3.08 | 3.4E-113 | 4E-111 |
| Nop10 | 22.60 | 2.21 | 4.7E-113 | 5.6E-111 |
| Hopx | 22.59 | 1.92 | 5.1E-113 | 5.9E-111 |
| Eppk1 | 22.59 | 1.54 | 5.5E-113 | 6.5E-111 |
| Txndc5 | 22.43 | 1.76 | 2.1E-111 | 2.4E-109 |
| Eif5a | 22.36 | 1.86 | 8.9E-111 | 1E-108 |
| Hsp90ab1 | 22.33 | 1.99 | 2E-110 | 2.3E-108 |
| Lsm3 | 22.33 | 2.10 | 2.1E-110 | 2.4E-108 |
| Rrm1 | 22.32 | 2.63 | 2.2E-110 | 2.5E-108 |
| Nusap1 | 22.21 | 3.05 | 2.6E-109 | 2.9E-107 |
| Anp32a | 22.21 | 1.81 | 2.6E-109 | 2.9E-107 |
| Snrpd1 | 22.15 | 2.52 | 1.1E-108 | 1.2E-106 |
| Krt18 | 22.11 | 1.70 | 2.6E-108 | 2.9E-106 |
| Kif11 | 22.11 | 2.98 | 2.6E-108 | 2.9E-106 |
| Mat2a | 22.10 | 2.00 | 3.2E-108 | 3.4E-106 |
| Rab5if | 22.09 | 1.71 | 3.9E-108 | 4.2E-106 |
| Hnrnpab | 22.07 | 1.96 | 6.6E-108 | 7.1E-106 |
| Rrm2 | 21.98 | 3.08 | 4.2E-107 | 4.5E-105 |
| Stard7 | 21.83 | 1.92 | 1.2E-105 | 1.3E-103 |
| Serbp1 | 21.82 | 2.03 | 1.4E-105 | 1.5E-103 |
| Dynll2 | 21.81 | 1.59 | 1.9E-105 | 1.9E-103 |
| Abhd11 | 21.73 | 1.86 | 1.2E-104 | 1.2E-102 |
| Eif3l | 21.72 | 1.90 | 1.4E-104 | 1.5E-102 |
| Ccdc34 | 21.71 | 2.56 | 1.7E-104 | 1.8E-102 |
| Tomm40 | 21.67 | 2.23 | 4.1E-104 | 4.2E-102 |
| Sh3bgrl2 | 21.67 | 1.81 | 4.4E-104 | 4.4E-102 |

| | | | | |
|---|---|---|---|---|
| Snrnp70 | 21.63 | 1.61 | 9.8E-104 | 9.8E-102 |
| Dazap1 | 21.61 | 1.76 | 1.4E-103 | 1.4E-101 |
| Dnmt1 | 21.59 | 2.59 | 2.4E-103 | 2.4E-101 |
| Nono | 21.54 | 1.95 | 6.9E-103 | 6.9E-101 |
| Eef1g | 21.50 | 2.16 | 1.5E-102 | 1.4E-100 |
| H2afx | 21.50 | 2.26 | 1.7E-102 | 1.6E-100 |
| Eif3k | 21.44 | 1.65 | 5.1E-102 | 4.9E-100 |
| Ddx39 | 21.42 | 2.20 | 8.4E-102 | 8E-100 |
| Ccna2 | 21.42 | 3.01 | 9.4E-102 | 9E-100 |
| Trp53i11 | 21.41 | 2.35 | 1E-101 | 9.8E-100 |
| Stip1 | 21.35 | 1.90 | 4.1E-101 | 3.9E-99 |
| Tfdp1 | 21.32 | 1.99 | 7E-101 | 6.6E-99 |
| Slc1a5 | 21.32 | 1.93 | 8.1E-101 | 7.6E-99 |
| Cnn3 | 21.29 | 2.43 | 1.3E-100 | 1.2E-98 |
| Hsd17b10 | 21.29 | 2.17 | 1.3E-100 | 1.21E-98 |
| Cd9 | 21.28 | 1.78 | 1.7E-100 | 1.56E-98 |
| Lrpprc | 21.21 | 2.47 | 8.1E-100 | 7.43E-98 |
| Incenp | 21.18 | 3.03 | 1.3E-99 | 1.21E-97 |
| Esrp1 | 21.18 | 1.85 | 1.4E-99 | 1.27E-97 |
| Ccnd1 | 21.15 | 2.42 | 2.9E-99 | 2.57E-97 |
| Srrt | 21.13 | 2.12 | 4.1E-99 | 3.59E-97 |
| Foxm1 | 21.07 | 3.11 | 1.36E-98 | 1.2E-96 |
| Tardbp | 21.04 | 1.79 | 2.95E-98 | 2.58E-96 |
| Syncrip | 21.03 | 2.03 | 3.23E-98 | 2.81E-96 |
| Nsun2 | 21.00 | 2.23 | 7.07E-98 | 6.12E-96 |
| Noc2l | 20.98 | 2.56 | 1.1E-97 | 9.38E-96 |
| Cct3 | 20.96 | 2.22 | 1.61E-97 | 1.37E-95 |
| Ncl | 20.95 | 2.38 | 2.02E-97 | 1.72E-95 |
| Tceal9 | 20.91 | 1.89 | 4.57E-97 | 3.86E-95 |
| Eif3g | 20.87 | 1.95 | 9.34E-97 | 7.81E-95 |
| H2afy | 20.83 | 1.77 | 2.47E-96 | 2.05E-94 |
| Dtymk | 20.79 | 2.39 | 5.37E-96 | 4.43E-94 |
| Ahsa1 | 20.77 | 1.94 | 7.79E-96 | 6.4E-94 |
| Pcna | 20.75 | 2.06 | 1.18E-95 | 9.64E-94 |
| Psme3 | 20.68 | 1.69 | 4.96E-95 | 4.02E-93 |
| Tmem147 | 20.67 | 2.14 | 6.04E-95 | 4.88E-93 |
| Ephb4 | 20.63 | 2.32 | 1.46E-94 | 1.18E-92 |
| Sae1 | 20.62 | 2.24 | 1.93E-94 | 1.55E-92 |
| Isyna1 | 20.49 | 2.20 | 2.43E-93 | 1.92E-91 |
| Myc | 20.49 | 2.41 | 2.45E-93 | 1.93E-91 |
| Hist1h3c | 20.45 | 3.12 | 6.1E-93 | 4.75E-91 |
| Hspa8 | 20.43 | 1.51 | 8.58E-93 | 6.65E-91 |
| Snrpd3 | 20.39 | 1.94 | 1.87E-92 | 1.44E-90 |
| Ywhae | 20.39 | 1.50 | 2.09E-92 | 1.6E-90 |

| | | | | |
|---|---|---|---|---|
| Impdh2 | 20.38 | 2.31 | 2.75E-92 | 2.08E-90 |
| Snrpa | 20.32 | 1.85 | 8.95E-92 | 6.77E-90 |
| Ndufa10 | 20.29 | 1.62 | 1.53E-91 | 1.15E-89 |
| Rcc1 | 20.27 | 2.51 | 2.6E-91 | 1.95E-89 |
| Golm1 | 20.25 | 1.59 | 3.81E-91 | 2.85E-89 |
| Bzw1 | 20.24 | 1.50 | 4.73E-91 | 3.52E-89 |
| Snd1 | 20.16 | 1.56 | 2.23E-90 | 1.65E-88 |
| Timm13 | 20.08 | 1.61 | 1.06E-89 | 7.83E-88 |
| Hacd1 | 19.99 | 2.04 | 6.25E-89 | 4.6E-87 |
| Rpia | 19.97 | 2.17 | 9.68E-89 | 7.1E-87 |
| Sema5a | 19.95 | 2.30 | 1.47E-88 | 1.07E-86 |
| Knstrn | 19.93 | 2.83 | 2.06E-88 | 1.5E-86 |
| Rpa3 | 19.89 | 2.38 | 4.8E-88 | 3.48E-86 |
| Xpo1 | 19.87 | 2.10 | 8.09E-88 | 5.82E-86 |
| Fasn | 19.78 | 2.27 | 4.23E-87 | 3.03E-85 |
| Idh3b | 19.68 | 1.68 | 2.96E-86 | 2.11E-84 |
| Ppa1 | 19.66 | 1.74 | 4.34E-86 | 3.08E-84 |
| Lbr | 19.58 | 2.21 | 2.35E-85 | 1.65E-83 |
| Sdc4 | 19.58 | 1.51 | 2.36E-85 | 1.66E-83 |
| Ppil1 | 19.49 | 2.24 | 1.35E-84 | 9.39E-83 |
| Cbx5 | 19.44 | 2.37 | 3.22E-84 | 2.24E-82 |
| Timm50 | 19.37 | 2.11 | 1.27E-83 | 8.83E-82 |
| Ehf | 19.29 | 1.71 | 6.19E-83 | 4.26E-81 |
| Dhx9 | 19.26 | 1.72 | 1.18E-82 | 8.09E-81 |
| Tsr1 | 19.25 | 2.40 | 1.35E-82 | 9.25E-81 |
| H2afz | 19.23 | 2.21 | 2.02E-82 | 1.37E-80 |
| Mcm5 | 19.21 | 2.96 | 3.25E-82 | 2.2E-80 |
| Mthfd1 | 19.19 | 2.75 | 4.62E-82 | 3.11E-80 |
| Emg1 | 19.17 | 2.09 | 6.21E-82 | 4.17E-80 |
| Larp1 | 19.11 | 1.65 | 1.96E-81 | 1.31E-79 |
| Tufm | 19.10 | 1.97 | 2.44E-81 | 1.62E-79 |
| Nudc | 19.10 | 1.97 | 2.67E-81 | 1.77E-79 |
| Arhgef38 | 19.09 | 2.09 | 3.22E-81 | 2.12E-79 |
| Cdt1 | 19.07 | 2.36 | 4.19E-81 | 2.74E-79 |
| Cdca7 | 19.01 | 2.26 | 1.37E-80 | 8.87E-79 |
| Dek | 19.00 | 1.97 | 1.69E-80 | 1.09E-78 |
| Hnrnpa2b1 | 18.96 | 1.86 | 3.35E-80 | 2.15E-78 |
| Mybbp1a | 18.94 | 2.41 | 5.51E-80 | 3.52E-78 |
| Ndufc2 | 18.92 | 1.57 | 7.56E-80 | 4.82E-78 |
| Ifrd2 | 18.87 | 2.24 | 1.9E-79 | 1.21E-77 |
| Timm44 | 18.84 | 1.93 | 3.4E-79 | 2.14E-77 |
| Akt1s1 | 18.81 | 1.65 | 6.24E-79 | 3.91E-77 |
| Trp53 | 18.81 | 1.98 | 6.44E-79 | 4.03E-77 |
| Bclaf1 | 18.71 | 1.56 | 4.39E-78 | 2.73E-76 |

| | | | | |
|---|---|---|---|---|
| Acat1 | 18.66 | 1.94 | 1.13E-77 | 6.94E-76 |
| Ruvbl2 | 18.65 | 2.39 | 1.23E-77 | 7.51E-76 |
| Cetn3 | 18.53 | 2.04 | 1.26E-76 | 7.65E-75 |
| Npm3 | 18.51 | 2.65 | 1.63E-76 | 9.88E-75 |
| Snrpg | 18.51 | 2.15 | 1.87E-76 | 1.13E-74 |
| Srek1 | 18.45 | 1.87 | 5.14E-76 | 3.1E-74 |
| Ccnb2 | 18.38 | 2.64 | 2.04E-75 | 1.23E-73 |
| Tacc3 | 18.37 | 2.62 | 2.48E-75 | 1.48E-73 |
| Kif15 | 18.36 | 2.64 | 2.84E-75 | 1.69E-73 |
| Ptcd3 | 18.35 | 2.17 | 3.49E-75 | 2.07E-73 |
| Got2 | 18.33 | 2.05 | 4.37E-75 | 2.59E-73 |
| Naca | 18.33 | 1.68 | 4.94E-75 | 2.92E-73 |
| Sh2d4a | 18.31 | 2.04 | 6.75E-75 | 3.97E-73 |
| Akap8 | 18.30 | 1.70 | 7.64E-75 | 4.48E-73 |
| Ptges3 | 18.30 | 1.92 | 8.03E-75 | 4.69E-73 |
| Ddx39b | 18.29 | 1.82 | 9.12E-75 | 5.32E-73 |
| Tipin | 18.29 | 2.50 | 1.07E-74 | 6.22E-73 |
| Gnl3 | 18.28 | 2.48 | 1.17E-74 | 6.79E-73 |
| Eif3f | 18.26 | 1.62 | 1.59E-74 | 9.2E-73 |
| Uqcc2 | 18.25 | 1.89 | 2.03E-74 | 1.16E-72 |
| Atp13a3 | 18.20 | 1.58 | 5.14E-74 | 2.93E-72 |
| Sfpq | 18.20 | 1.96 | 5.47E-74 | 3.11E-72 |
| Fubp1 | 18.19 | 1.94 | 6.74E-74 | 3.82E-72 |
| Clic6 | 18.17 | 2.51 | 9.66E-74 | 5.46E-72 |
| Elavl1 | 18.16 | 1.67 | 1.15E-73 | 6.47E-72 |
| Dpy30 | 18.15 | 1.98 | 1.36E-73 | 7.65E-72 |
| Fkbp4 | 18.12 | 1.97 | 2.19E-73 | 1.22E-71 |
| Rbm14 | 18.06 | 1.57 | 6.46E-73 | 3.58E-71 |
| Hat1 | 18.04 | 2.34 | 9.91E-73 | 5.48E-71 |
| Ephb2 | 18.03 | 2.43 | 1.06E-72 | 5.85E-71 |
| Hspa9 | 18.01 | 2.22 | 1.75E-72 | 9.59E-71 |
| Gar1 | 18.00 | 2.69 | 2E-72 | 1.09E-70 |
| Maz | 18.00 | 1.78 | 2.02E-72 | 1.1E-70 |
| Hsp90aa1 | 17.97 | 2.10 | 3.56E-72 | 1.94E-70 |
| Taf1d | 17.93 | 2.01 | 6.53E-72 | 3.54E-70 |
| Ipo5 | 17.93 | 2.41 | 7.15E-72 | 3.86E-70 |
| Phb | 17.91 | 2.29 | 1.05E-71 | 5.67E-70 |
| Pbk | 17.88 | 3.73 | 1.61E-71 | 8.63E-70 |
| Pebp1 | 17.87 | 2.01 | 1.88E-71 | 1.01E-69 |
| Micos13 | 17.84 | 1.66 | 3.23E-71 | 1.73E-69 |
| Dars | 17.82 | 2.01 | 4.67E-71 | 2.48E-69 |
| Mbd3 | 17.82 | 1.85 | 4.97E-71 | 2.64E-69 |
| Hcfc1 | 17.76 | 1.95 | 1.37E-70 | 7.24E-69 |
| Trap1 | 17.74 | 2.29 | 2.09E-70 | 1.1E-68 |

| | | | | |
|---|---|---|---|---|
| Idh2 | 17.70 | 1.72 | 3.95E-70 | 2.07E-68 |
| Rbbp4 | 17.65 | 1.63 | 1.09E-69 | 5.65E-68 |
| Mt2 | 17.60 | 1.59 | 2.37E-69 | 1.23E-67 |
| Amot | 17.51 | 2.38 | 1.26E-68 | 6.47E-67 |
| Camk1d | 17.48 | 1.60 | 1.95E-68 | 9.95E-67 |
| Txn2 | 17.45 | 1.59 | 3.26E-68 | 1.66E-66 |
| Ddx21 | 17.44 | 2.13 | 3.98E-68 | 2.02E-66 |
| Tbrg4 | 17.41 | 2.02 | 6.67E-68 | 3.36E-66 |
| Ect2 | 17.37 | 2.65 | 1.31E-67 | 6.57E-66 |
| Umps | 17.37 | 2.39 | 1.43E-67 | 7.16E-66 |
| Lyar | 17.32 | 2.59 | 3.45E-67 | 1.72E-65 |
| Smarcc1 | 17.31 | 2.18 | 4.2E-67 | 2.08E-65 |
| Cftr | 17.28 | 1.94 | 6.3E-67 | 3.12E-65 |
| Rabggtb | 17.28 | 2.00 | 6.79E-67 | 3.35E-65 |
| Nek2 | 17.27 | 2.97 | 8.32E-67 | 4.1E-65 |
| Cfdp1 | 17.25 | 1.95 | 1.15E-66 | 5.64E-65 |
| Lrig1 | 17.23 | 1.89 | 1.6E-66 | 7.85E-65 |
| Apex1 | 17.22 | 2.65 | 1.74E-66 | 8.48E-65 |
| Seh1l | 17.17 | 1.77 | 4.28E-66 | 2.09E-64 |
| Rnps1 | 17.17 | 1.50 | 4.52E-66 | 2.2E-64 |
| Actn1 | 17.16 | 1.70 | 4.96E-66 | 2.41E-64 |
| Aurkb | 17.12 | 2.94 | 1.13E-65 | 5.41E-64 |
| Ptov1 | 17.11 | 1.66 | 1.29E-65 | 6.18E-64 |
| Gtpbp4 | 17.10 | 2.06 | 1.56E-65 | 7.45E-64 |
| Cdk4 | 17.10 | 2.22 | 1.57E-65 | 7.5E-64 |
| Prpf19 | 17.07 | 2.02 | 2.29E-65 | 1.08E-63 |
| Hes1 | 17.04 | 1.99 | 4.13E-65 | 1.95E-63 |
| Slbp | 17.03 | 1.72 | 5.3E-65 | 2.5E-63 |
| Ubap2 | 17.02 | 2.05 | 6.15E-65 | 2.89E-63 |
| Timm10 | 17.01 | 2.31 | 6.56E-65 | 3.08E-63 |
| Polr2f | 17.00 | 1.71 | 8.02E-65 | 3.76E-63 |
| Rpa1 | 16.98 | 2.31 | 1.1E-64 | 5.13E-63 |
| Plk1 | 16.94 | 2.67 | 2.39E-64 | 1.11E-62 |
| Nudt21 | 16.92 | 1.94 | 3.46E-64 | 1.6E-62 |
| Alg8 | 16.89 | 2.69 | 4.95E-64 | 2.29E-62 |
| Actl6a | 16.88 | 2.25 | 6.67E-64 | 3.07E-62 |
| Add3 | 16.87 | 1.63 | 7.5E-64 | 3.44E-62 |
| Acly | 16.85 | 1.58 | 1.13E-63 | 5.18E-62 |
| Ola1 | 16.81 | 1.79 | 2.02E-63 | 9.25E-62 |
| Fh1 | 16.79 | 1.79 | 2.91E-63 | 1.32E-61 |
| Farsb | 16.78 | 2.02 | 3.45E-63 | 1.57E-61 |
| Ptbp1 | 16.76 | 1.53 | 4.42E-63 | 2.01E-61 |
| Rasa4 | 16.70 | 2.15 | 1.29E-62 | 5.85E-61 |
| Pmf1 | 16.70 | 2.67 | 1.31E-62 | 5.91E-61 |

| | | | | |
|---|---|---|---|---|
| Creb3l1 | 16.66 | 1.94 | 2.48E-62 | 1.12E-60 |
| Ndufa4 | 16.66 | 1.52 | 2.62E-62 | 1.18E-60 |
| Smoc2 | 16.65 | 1.95 | 3.05E-62 | 1.37E-60 |
| Ibtk | 16.64 | 1.74 | 3.5E-62 | 1.56E-60 |
| Srsf10 | 16.64 | 2.02 | 3.62E-62 | 1.61E-60 |
| Tmed9 | 16.63 | 1.66 | 4.03E-62 | 1.79E-60 |
| Eif3d | 16.62 | 1.96 | 4.62E-62 | 2.05E-60 |
| U2af1 | 16.62 | 2.04 | 4.65E-62 | 2.06E-60 |
| Snrpa1 | 16.61 | 2.35 | 6.24E-62 | 2.75E-60 |
| Hirip3 | 16.59 | 2.65 | 8.72E-62 | 3.8E-60 |
| Sf3a3 | 16.57 | 2.03 | 1.2E-61 | 5.21E-60 |
| Rtcb | 16.57 | 1.51 | 1.24E-61 | 5.38E-60 |
| Ppm1g | 16.56 | 1.79 | 1.3E-61 | 5.61E-60 |
| Rad54l | 16.56 | 3.09 | 1.31E-61 | 5.65E-60 |
| Tfam | 16.56 | 1.92 | 1.42E-61 | 6.12E-60 |
| Cdca7l | 16.51 | 2.65 | 3.06E-61 | 1.31E-59 |
| Tomm70a | 16.51 | 1.77 | 3.1E-61 | 1.32E-59 |
| Krtcap3 | 16.46 | 1.81 | 6.84E-61 | 2.9E-59 |
| Anapc13 | 16.46 | 1.58 | 6.84E-61 | 2.9E-59 |
| Pdia6 | 16.41 | 1.65 | 1.6E-60 | 6.72E-59 |
| Pus1 | 16.37 | 2.44 | 2.89E-60 | 1.21E-58 |
| Smyd2 | 16.31 | 2.23 | 8.65E-60 | 3.61E-58 |
| Psmb2 | 16.27 | 1.54 | 1.67E-59 | 6.9E-58 |
| Ncapd2 | 16.26 | 2.50 | 1.99E-59 | 8.2E-58 |
| Rnf186 | 16.24 | 1.83 | 2.58E-59 | 1.06E-57 |
| Srsf3 | 16.24 | 1.75 | 2.73E-59 | 1.11E-57 |
| Mis18a | 16.23 | 2.53 | 3.18E-59 | 1.3E-57 |
| Lig1 | 16.21 | 2.79 | 4.58E-59 | 1.86E-57 |
| Cenpx | 16.20 | 2.10 | 4.77E-59 | 1.94E-57 |
| Brix1 | 16.20 | 2.28 | 4.8E-59 | 1.95E-57 |
| Akap1 | 16.19 | 2.04 | 5.48E-59 | 2.22E-57 |
| Cpsf6 | 16.19 | 1.78 | 5.89E-59 | 2.38E-57 |
| Hmgb3 | 16.18 | 2.22 | 7.44E-59 | 2.99E-57 |
| Cdc20 | 16.16 | 2.84 | 1.02E-58 | 4.07E-57 |
| Cdca8 | 16.15 | 2.62 | 1.09E-58 | 4.36E-57 |
| Pold2 | 16.12 | 2.25 | 1.88E-58 | 7.48E-57 |
| Tmem97 | 16.11 | 1.78 | 2.21E-58 | 8.75E-57 |
| Pa2g4 | 16.09 | 2.23 | 2.85E-58 | 1.12E-56 |
| C1qbp | 16.09 | 2.35 | 2.87E-58 | 1.13E-56 |
| Haus1 | 16.09 | 2.68 | 3.13E-58 | 1.23E-56 |
| Eef1d | 16.08 | 2.22 | 3.47E-58 | 1.36E-56 |
| Hnrnpd | 16.07 | 1.77 | 4.16E-58 | 1.62E-56 |
| Ywhag | 16.02 | 1.77 | 8.79E-58 | 3.4E-56 |
| Hdgf | 16.01 | 1.86 | 1.1E-57 | 4.25E-56 |

| | | | | |
|---|---|---|---|---|
| Pck2 | 16.01 | 2.22 | 1.17E-57 | 4.5E-56 |
| Polr2e | 16.00 | 1.90 | 1.22E-57 | 4.7E-56 |
| add45gip1 | 15.99 | 2.15 | 1.39E-57 | 5.34E-56 |
| Nasp | 15.99 | 2.12 | 1.4E-57 | 5.35E-56 |
| Spc24 | 15.98 | 2.78 | 1.77E-57 | 6.72E-56 |
| Pclaf | 15.94 | 2.65 | 3.34E-57 | 1.26E-55 |
| Pdss1 | 15.92 | 1.81 | 4.62E-57 | 1.73E-55 |
| Nap1l4 | 15.90 | 1.81 | 6.1E-57 | 2.28E-55 |
| Fbl | 15.89 | 2.32 | 7.15E-57 | 2.66E-55 |
| Bola2 | 15.89 | 1.78 | 7.75E-57 | 2.88E-55 |
| Psmd7 | 15.88 | 1.56 | 9.39E-57 | 3.48E-55 |
| Jpt2 | 15.87 | 1.65 | 9.87E-57 | 3.65E-55 |
| Slirp | 15.87 | 1.61 | 1.07E-56 | 3.95E-55 |
| Usp10 | 15.83 | 2.09 | 1.83E-56 | 6.74E-55 |
| Ssna1 | 15.80 | 2.07 | 3.33E-56 | 1.22E-54 |
| E2f8 | 15.76 | 2.72 | 5.89E-56 | 2.14E-54 |
| Plekha6 | 15.75 | 1.55 | 6.71E-56 | 2.43E-54 |
| Rab25 | 15.75 | 1.63 | 7.14E-56 | 2.58E-54 |
| Dgcr8 | 15.71 | 2.19 | 1.31E-55 | 4.71E-54 |
| S100a16 | 15.71 | 1.73 | 1.36E-55 | 4.88E-54 |
| Huwe1 | 15.70 | 1.66 | 1.42E-55 | 5.09E-54 |
| Naa38 | 15.67 | 1.59 | 2.31E-55 | 8.22E-54 |
| Hjurp | 15.65 | 1.57 | 3.53E-55 | 1.26E-53 |
| Nsd2 | 15.63 | 1.88 | 4.42E-55 | 1.57E-53 |
| Nans | 15.63 | 1.69 | 4.62E-55 | 1.64E-53 |
| Suv39h1 | 15.60 | 2.43 | 7.19E-55 | 2.52E-53 |
| Trim28 | 15.60 | 2.12 | 7.71E-55 | 2.7E-53 |
| Api5 | 15.59 | 1.68 | 7.91E-55 | 2.77E-53 |
| Rad51 | 15.58 | 2.39 | 9.48E-55 | 3.3E-53 |
| Pgls | 15.57 | 1.76 | 1.2E-54 | 4.14E-53 |
| Magoh | 15.55 | 1.82 | 1.57E-54 | 5.39E-53 |
| Eprs | 15.51 | 2.03 | 2.98E-54 | 1.03E-52 |
| Rnaseh2c | 15.50 | 2.00 | 3.73E-54 | 1.28E-52 |
| Adsl | 15.49 | 2.25 | 3.77E-54 | 1.29E-52 |
| Cnot10 | 15.49 | 1.94 | 4.07E-54 | 1.39E-52 |
| Grpel1 | 15.48 | 1.62 | 4.72E-54 | 1.61E-52 |
| Coq2 | 15.48 | 2.07 | 4.79E-54 | 1.63E-52 |
| Ranbp2 | 15.48 | 1.73 | 4.87E-54 | 1.65E-52 |
| Prc1 | 15.48 | 2.50 | 5.12E-54 | 1.73E-52 |
| Eif2a | 15.47 | 1.74 | 5.62E-54 | 1.9E-52 |
| Tomm6 | 15.47 | 1.83 | 5.84E-54 | 1.97E-52 |
| Gtf2i | 15.46 | 1.59 | 6.32E-54 | 2.13E-52 |
| Dnah8 | 15.43 | 2.93 | 1E-53 | 3.36E-52 |
| Mcm6 | 15.40 | 2.08 | 1.74E-53 | 5.81E-52 |

| | | | | |
|---|---|---|---|---|
| Dnajc15 | 15.39 | 1.97 | 2.05E-53 | 6.83E-52 |
| Wdr83os | 15.38 | 1.68 | 2.23E-53 | 7.42E-52 |
| Mcm7 | 15.38 | 2.38 | 2.39E-53 | 7.91E-52 |
| Tyms | 15.37 | 2.75 | 2.43E-53 | 8.05E-52 |
| Pelp1 | 15.37 | 2.75 | 2.52E-53 | 8.31E-52 |
| Ndufs3 | 15.34 | 1.52 | 4.08E-53 | 1.34E-51 |
| Metap2 | 15.33 | 1.62 | 5.13E-53 | 1.69E-51 |
| Esco2 | 15.30 | 3.17 | 7.72E-53 | 2.53E-51 |
| Rangap1 | 15.29 | 2.06 | 8.69E-53 | 2.84E-51 |
| Fam98b | 15.29 | 2.20 | 8.88E-53 | 2.9E-51 |
| Kars | 15.26 | 1.64 | 1.34E-52 | 4.33E-51 |
| Dnaja3 | 15.22 | 1.96 | 2.76E-52 | 8.9E-51 |
| Denr | 15.21 | 1.88 | 3E-52 | 9.64E-51 |
| Prps1l3 | 15.19 | 2.29 | 4.38E-52 | 1.4E-50 |
| Aimp2 | 15.18 | 1.78 | 4.78E-52 | 1.53E-50 |
| Exosc7 | 15.18 | 2.20 | 5.04E-52 | 1.6E-50 |
| Nucks1 | 15.15 | 2.34 | 8.07E-52 | 2.56E-50 |
| Nme1 | 15.14 | 2.15 | 8.29E-52 | 2.63E-50 |
| Fads2 | 15.12 | 1.87 | 1.16E-51 | 3.67E-50 |
| Tcf3 | 15.07 | 1.84 | 2.65E-51 | 8.35E-50 |
| Ssbp1 | 15.06 | 1.84 | 2.83E-51 | 8.89E-50 |
| Rab3ip | 15.04 | 1.56 | 3.94E-51 | 1.24E-49 |
| Prmt5 | 15.04 | 2.82 | 4.24E-51 | 1.33E-49 |
| Tnfsf13 | 15.04 | 1.83 | 4.31E-51 | 1.34E-49 |
| Ndufaf4 | 15.02 | 2.57 | 5.14E-51 | 1.6E-49 |
| Prdx3 | 15.01 | 1.67 | 6.03E-51 | 1.87E-49 |
| Dut | 15.00 | 2.38 | 7.48E-51 | 2.32E-49 |
| Pfdn6 | 15.00 | 1.75 | 7.55E-51 | 2.33E-49 |
| Timm23 | 15.00 | 1.90 | 7.85E-51 | 2.42E-49 |
| Pold1 | 14.99 | 2.55 | 8.1E-51 | 2.49E-49 |
| Marcks | 14.96 | 1.53 | 1.39E-50 | 4.27E-49 |
| Psmg4 | 14.95 | 2.39 | 1.64E-50 | 5.03E-49 |
| Bdh1 | 14.92 | 1.55 | 2.29E-50 | 6.96E-49 |
| Snrnp40 | 14.92 | 1.93 | 2.56E-50 | 7.77E-49 |
| Aldh18a1 | 14.88 | 1.53 | 4.2E-50 | 1.27E-48 |
| Uchl3 | 14.87 | 1.81 | 5.06E-50 | 1.53E-48 |
| Cbfb | 14.87 | 1.85 | 5.23E-50 | 1.58E-48 |
| Pafah1b3 | 14.86 | 1.75 | 6.44E-50 | 1.94E-48 |
| Dhx15 | 14.85 | 1.72 | 6.96E-50 | 2.09E-48 |
| Sigmar1 | 14.85 | 2.06 | 7.02E-50 | 2.1E-48 |
| Rbm17 | 14.85 | 1.73 | 7.06E-50 | 2.11E-48 |
| Gtse1 | 14.84 | 2.59 | 7.8E-50 | 2.32E-48 |
| Luc7l3 | 14.83 | 1.53 | 9.8E-50 | 2.92E-48 |
| Topbp1 | 14.81 | 2.11 | 1.19E-49 | 3.54E-48 |

| | | | | |
|---|---|---|---|---|
| Bzw2 | 14.80 | 2.09 | 1.51E-49 | 4.49E-48 |
| Smc4 | 14.79 | 2.20 | 1.58E-49 | 4.68E-48 |
| Tspo | 14.79 | 1.79 | 1.71E-49 | 5.06E-48 |
| Nrarp | 14.78 | 2.16 | 1.86E-49 | 5.5E-48 |
| Tfap4 | 14.78 | 2.28 | 2.11E-49 | 6.22E-48 |
| Plch2 | 14.74 | 2.02 | 3.48E-49 | 1.02E-47 |
| Pdk1 | 14.73 | 2.11 | 3.94E-49 | 1.15E-47 |
| Wdr43 | 14.73 | 2.71 | 4.32E-49 | 1.26E-47 |
| Iqgap3 | 14.72 | 2.93 | 4.95E-49 | 1.44E-47 |
| G6pc3 | 14.71 | 2.29 | 5.9E-49 | 1.71E-47 |
| Nsmce4a | 14.70 | 1.51 | 6.39E-49 | 1.84E-47 |
| Oxa1l | 14.67 | 1.71 | 1.07E-48 | 3.08E-47 |
| Tars | 14.66 | 1.62 | 1.1E-48 | 3.14E-47 |
| Nudcd2 | 14.66 | 2.53 | 1.21E-48 | 3.47E-47 |
| Polr1c | 14.64 | 1.77 | 1.67E-48 | 4.77E-47 |
| Gtf2h5 | 14.62 | 1.50 | 2E-48 | 5.71E-47 |
| Hacd3 | 14.62 | 1.52 | 2.18E-48 | 6.19E-47 |
| Anp32b | 14.61 | 2.18 | 2.27E-48 | 6.44E-47 |
| Hnrnpa0 | 14.60 | 1.63 | 2.74E-48 | 7.74E-47 |
| Eml4 | 14.58 | 1.67 | 3.59E-48 | 1.01E-46 |
| Tcerg1 | 14.58 | 1.81 | 3.61E-48 | 1.02E-46 |
| Ppp1r9a | 14.56 | 2.13 | 5.29E-48 | 1.49E-46 |
| Uhrf2 | 14.55 | 1.62 | 5.86E-48 | 1.64E-46 |
| Ilf2 | 14.54 | 2.20 | 6.99E-48 | 1.95E-46 |
| Anln | 14.53 | 2.75 | 7.39E-48 | 2.06E-46 |
| Eif2b1 | 14.45 | 1.87 | 2.59E-47 | 7.18E-46 |
| Rbp7 | 14.44 | 2.76 | 2.7E-47 | 7.46E-46 |
| Nup85 | 14.42 | 2.45 | 3.71E-47 | 1.02E-45 |
| Dele1 | 14.42 | 2.14 | 3.9E-47 | 1.07E-45 |
| Rfc2 | 14.42 | 1.81 | 3.99E-47 | 1.1E-45 |
| Myo5c | 14.41 | 1.84 | 4.78E-47 | 1.31E-45 |
| Iars | 14.40 | 2.13 | 4.82E-47 | 1.32E-45 |
| Nolc1 | 14.39 | 2.48 | 5.8E-47 | 1.58E-45 |
| Kif23 | 14.39 | 2.41 | 5.87E-47 | 1.6E-45 |
| Ppif | 14.35 | 2.04 | 9.94E-47 | 2.69E-45 |
| Cenpe | 14.35 | 2.81 | 1.01E-46 | 2.74E-45 |
| Pola1 | 14.31 | 2.30 | 1.77E-46 | 4.76E-45 |
| Setd6 | 14.31 | 2.17 | 1.88E-46 | 5.04E-45 |
| Mecr | 14.31 | 2.07 | 1.94E-46 | 5.21E-45 |
| Cops7a | 14.31 | 1.58 | 2E-46 | 5.36E-45 |
| Polr2h | 14.30 | 1.83 | 2.13E-46 | 5.71E-45 |
| Wdr36 | 14.29 | 2.15 | 2.42E-46 | 6.47E-45 |
| Dkc1 | 14.28 | 2.43 | 2.75E-46 | 7.32E-45 |
| Mybl2 | 14.28 | 2.67 | 2.77E-46 | 7.36E-45 |

| | | | | |
|---|---|---|---|---|
| Samm50 | 14.27 | 1.54 | 3.27E-46 | 8.67E-45 |
| Pum3 | 14.26 | 2.05 | 3.82E-46 | 1.01E-44 |
| Ung | 14.26 | 2.31 | 4.06E-46 | 1.07E-44 |
| Ilf3 | 14.25 | 2.12 | 4.5E-46 | 1.18E-44 |
| Bccip | 14.24 | 1.74 | 5E-46 | 1.31E-44 |
| Hdac2 | 14.24 | 2.03 | 5.39E-46 | 1.42E-44 |
| Atad3a | 14.24 | 2.00 | 5.46E-46 | 1.43E-44 |
| Sf3a2 | 14.22 | 1.90 | 6.6E-46 | 1.72E-44 |
| Abracl | 14.22 | 1.56 | 6.88E-46 | 1.79E-44 |
| Sf3b5 | 14.22 | 1.99 | 6.96E-46 | 1.81E-44 |
| Limk2 | 14.21 | 1.58 | 7.53E-46 | 1.95E-44 |
| Tra2b | 14.21 | 1.75 | 7.57E-46 | 1.96E-44 |
| Kifc1 | 14.21 | 2.88 | 8.09E-46 | 2.09E-44 |
| Kif18b | 14.20 | 2.89 | 8.57E-46 | 2.21E-44 |
| Ndc1 | 14.20 | 2.86 | 9.05E-46 | 2.33E-44 |
| Melk | 14.20 | 2.97 | 9.09E-46 | 2.34E-44 |
| Cse1l | 14.20 | 2.25 | 9.42E-46 | 2.42E-44 |
| Cdca5 | 14.19 | 3.10 | 1.11E-45 | 2.85E-44 |
| Cdc25b | 14.17 | 2.60 | 1.37E-45 | 3.5E-44 |
| Slc35c1 | 14.17 | 1.54 | 1.42E-45 | 3.63E-44 |
| Etfbkmt | 14.14 | 1.77 | 2.06E-45 | 5.23E-44 |
| Hnrnpu | 14.13 | 1.68 | 2.47E-45 | 6.28E-44 |
| Cdk6 | 14.12 | 2.07 | 2.84E-45 | 7.19E-44 |
| Mlxip | 14.11 | 1.62 | 3.41E-45 | 8.64E-44 |
| Bora | 14.10 | 2.52 | 3.94E-45 | 9.97E-44 |
| Cth | 14.09 | 1.83 | 4.23E-45 | 1.07E-43 |
| Pabpc4 | 14.09 | 1.57 | 4.25E-45 | 1.07E-43 |
| Gart | 14.09 | 2.29 | 4.51E-45 | 1.13E-43 |
| Parp1 | 14.08 | 2.24 | 4.83E-45 | 1.21E-43 |
| Hist1h1d | 14.08 | 2.48 | 4.88E-45 | 1.22E-43 |
| Gjb1 | 14.08 | 1.55 | 4.97E-45 | 1.24E-43 |
| Spc25 | 14.08 | 2.64 | 5.22E-45 | 1.3E-43 |
| Lonp1 | 14.05 | 1.98 | 7.41E-45 | 1.84E-43 |
| Wls | 14.05 | 1.77 | 7.97E-45 | 1.97E-43 |
| Eif4a1 | 14.05 | 1.66 | 7.99E-45 | 1.97E-43 |
| Ankle1 | 14.04 | 2.83 | 9.38E-45 | 2.31E-43 |
| Ccdc43 | 14.01 | 1.98 | 1.29E-44 | 3.18E-43 |
| Snrpe | 13.98 | 1.84 | 2.09E-44 | 5.11E-43 |
| Anapc11 | 13.98 | 1.64 | 2.15E-44 | 5.24E-43 |
| Ppan | 13.98 | 2.19 | 2.15E-44 | 5.24E-43 |
| Eif3e | 13.97 | 1.89 | 2.37E-44 | 5.77E-43 |
| Anapc5 | 13.96 | 1.88 | 2.57E-44 | 6.25E-43 |
| Far2 | 13.93 | 2.24 | 4.26E-44 | 1.04E-42 |
| Ebna1bp2 | 13.93 | 1.82 | 4.28E-44 | 1.04E-42 |

| | | | | |
|---|---|---|---|---|
| Rasgrf2 | 13.93 | 2.15 | 4.32E-44 | 1.05E-42 |
| Zdhhc13 | 13.92 | 1.88 | 4.55E-44 | 1.1E-42 |
| Coq8a | 13.91 | 2.50 | 5.37E-44 | 1.29E-42 |
| Stil | 13.91 | 3.12 | 5.77E-44 | 1.39E-42 |
| Dnajc8 | 13.90 | 1.78 | 6.4E-44 | 1.54E-42 |
| Smpd4 | 13.89 | 2.61 | 7.15E-44 | 1.72E-42 |
| Xrcc6 | 13.89 | 2.44 | 7.2E-44 | 1.73E-42 |
| Dap3 | 13.89 | 1.60 | 7.35E-44 | 1.76E-42 |
| Mak16 | 13.87 | 2.52 | 9.95E-44 | 2.38E-42 |
| Tmem209 | 13.86 | 2.16 | 1.05E-43 | 2.5E-42 |
| Psmg1 | 13.86 | 2.67 | 1.13E-43 | 2.7E-42 |
| Arhgap11a | 13.85 | 2.09 | 1.2E-43 | 2.85E-42 |
| Grsf1 | 13.85 | 1.55 | 1.22E-43 | 2.9E-42 |
| Hsd11b2 | 13.84 | 2.46 | 1.54E-43 | 3.64E-42 |
| Psmg2 | 13.82 | 2.42 | 2.06E-43 | 4.85E-42 |
| Ergic1 | 13.81 | 1.53 | 2.16E-43 | 5.08E-42 |
| Rfc3 | 13.80 | 2.35 | 2.45E-43 | 5.75E-42 |
| Xrn2 | 13.79 | 1.57 | 3.06E-43 | 7.16E-42 |
| Eftud2 | 13.79 | 1.94 | 3.12E-43 | 7.29E-42 |
| Sf3a1 | 13.78 | 1.61 | 3.19E-43 | 7.45E-42 |
| Tmpo | 13.78 | 2.24 | 3.35E-43 | 7.81E-42 |
| Ticrr | 13.77 | 3.29 | 4.02E-43 | 9.34E-42 |
| Rfc4 | 13.76 | 2.67 | 4.14E-43 | 9.6E-42 |
| Kif22 | 13.75 | 3.09 | 4.85E-43 | 1.12E-41 |
| Psma7 | 13.75 | 1.71 | 4.91E-43 | 1.14E-41 |
| Bop1 | 13.73 | 1.98 | 6.81E-43 | 1.57E-41 |
| Map3k20 | 13.73 | 1.77 | 7.05E-43 | 1.62E-41 |
| Agpat4 | 13.72 | 2.25 | 7.72E-43 | 1.77E-41 |
| Mcm4 | 13.71 | 2.82 | 9.25E-43 | 2.12E-41 |
| Trim37 | 13.70 | 2.18 | 1.03E-42 | 2.35E-41 |
| Ddb2 | 13.70 | 2.19 | 1.03E-42 | 2.35E-41 |
| Rangrf | 13.70 | 2.40 | 1.06E-42 | 2.43E-41 |
| Fhod1 | 13.69 | 2.53 | 1.22E-42 | 2.79E-41 |
| Cdk2 | 13.68 | 2.30 | 1.41E-42 | 3.2E-41 |
| Srm | 13.67 | 2.27 | 1.54E-42 | 3.5E-41 |
| Bmp3 | 13.67 | 1.66 | 1.59E-42 | 3.59E-41 |
| Chchd1 | 13.66 | 1.72 | 1.76E-42 | 3.98E-41 |
| Nedd4 | 13.66 | 1.76 | 1.78E-42 | 4.02E-41 |
| Naa10 | 13.66 | 2.42 | 1.87E-42 | 4.22E-41 |
| Hepacam2 | 13.64 | 1.54 | 2.47E-42 | 5.56E-41 |
| Msi2 | 13.63 | 1.62 | 2.8E-42 | 6.3E-41 |
| Glrx5 | 13.61 | 1.76 | 3.34E-42 | 7.48E-41 |
| Phldb1 | 13.59 | 1.81 | 4.54E-42 | 1.01E-40 |
| Ncaph | 13.58 | 2.74 | 5.05E-42 | 1.13E-40 |

| | | | | |
|---|---|---|---|---|
| Krt7 | 13.58 | 1.57 | 5.22E-42 | 1.16E-40 |
| Prom1 | 13.58 | 1.90 | 5.62E-42 | 1.25E-40 |
| Pkdcc | 13.57 | 1.90 | 5.84E-42 | 1.3E-40 |
| Mogs | 13.57 | 2.10 | 6.35E-42 | 1.41E-40 |
| Lzts2 | 13.54 | 1.67 | 9.15E-42 | 2.02E-40 |
| Supt16 | 13.54 | 1.84 | 9.4E-42 | 2.07E-40 |
| Mpp6 | 13.53 | 2.04 | 1E-41 | 2.2E-40 |
| Mtx2 | 13.53 | 1.65 | 1.02E-41 | 2.24E-40 |
| Epb41l2 | 13.52 | 1.72 | 1.12E-41 | 2.46E-40 |
| Fastkd2 | 13.52 | 2.39 | 1.27E-41 | 2.78E-40 |
| Prpf31 | 13.51 | 2.37 | 1.4E-41 | 3.07E-40 |
| Kri1 | 13.51 | 1.94 | 1.44E-41 | 3.15E-40 |
| Pwwp3a | 13.50 | 2.33 | 1.56E-41 | 3.41E-40 |
| Hdac11 | 13.49 | 1.63 | 1.74E-41 | 3.8E-40 |
| Mcm2 | 13.48 | 2.57 | 2.03E-41 | 4.42E-40 |
| Nup155 | 13.48 | 2.45 | 2.13E-41 | 4.63E-40 |
| Arhgef39 | 13.47 | 2.59 | 2.2E-41 | 4.78E-40 |
| Car9 | 13.46 | 1.76 | 2.84E-41 | 6.13E-40 |
| Abhd14a | 13.43 | 2.27 | 3.83E-41 | 8.22E-40 |
| Tpx2 | 13.43 | 2.66 | 3.83E-41 | 8.23E-40 |
| Rnaseh2a | 13.43 | 2.15 | 4.29E-41 | 9.19E-40 |
| Shcbp1 | 13.42 | 3.17 | 4.54E-41 | 9.72E-40 |
| Gjb3 | 13.42 | 2.07 | 4.62E-41 | 9.87E-40 |
| Nifk | 13.42 | 2.24 | 4.75E-41 | 1.01E-39 |
| Slc27a2 | 13.41 | 1.64 | 5.48E-41 | 1.17E-39 |
| Gnpat | 13.41 | 1.98 | 5.62E-41 | 1.19E-39 |
| Cdca2 | 13.40 | 2.50 | 6.09E-41 | 1.29E-39 |
| Syde2 | 13.40 | 2.18 | 6.24E-41 | 1.32E-39 |
| Psmg3 | 13.40 | 2.16 | 6.34E-41 | 1.34E-39 |
| Prdx4 | 13.33 | 2.11 | 1.56E-40 | 3.3E-39 |
| Hibadh | 13.32 | 1.51 | 1.72E-40 | 3.64E-39 |
| Pdia5 | 13.32 | 1.88 | 1.76E-40 | 3.7E-39 |
| Phb2 | 13.32 | 1.60 | 1.77E-40 | 3.72E-39 |
| Bri3bp | 13.31 | 1.62 | 1.92E-40 | 4.04E-39 |
| Nob1 | 13.31 | 2.79 | 2.04E-40 | 4.29E-39 |
| Hsf1 | 13.29 | 1.89 | 2.56E-40 | 5.36E-39 |
| Fam129b | 13.29 | 1.76 | 2.62E-40 | 5.49E-39 |
| Prmt7 | 13.26 | 2.32 | 3.88E-40 | 8.08E-39 |
| Rad54b | 13.26 | 2.85 | 4.07E-40 | 8.48E-39 |
| Ecd | 13.24 | 2.18 | 5.16E-40 | 1.07E-38 |
| Shmt2 | 13.24 | 2.16 | 5.41E-40 | 1.12E-38 |
| Cenpf | 13.23 | 2.87 | 5.68E-40 | 1.18E-38 |
| Ptrh2 | 13.21 | 1.82 | 7.35E-40 | 1.52E-38 |
| Hmgb1 | 13.21 | 1.84 | 7.79E-40 | 1.61E-38 |

| | | | | |
|---|---|---|---|---|
| Trmt112 | 13.20 | 1.88 | 8.84E-40 | 1.82E-38 |
| Adpgk | 13.19 | 2.30 | 9.99E-40 | 2.06E-38 |
| Itga2 | 13.18 | 1.91 | 1.08E-39 | 2.22E-38 |
| Wdr77 | 13.17 | 2.09 | 1.27E-39 | 2.61E-38 |
| Cct6a | 13.16 | 1.89 | 1.51E-39 | 3.09E-38 |
| Alg10b | 13.16 | 1.69 | 1.54E-39 | 3.15E-38 |
| Gtpbp10 | 13.15 | 1.83 | 1.65E-39 | 3.36E-38 |
| Gstz1 | 13.15 | 1.69 | 1.73E-39 | 3.52E-38 |
| Zfp704 | 13.14 | 2.01 | 1.9E-39 | 3.86E-38 |
| Erh | 13.14 | 2.74 | 1.94E-39 | 3.94E-38 |
| Pcbp1 | 13.12 | 1.53 | 2.52E-39 | 5.12E-38 |
| Parn | 13.10 | 2.58 | 3.12E-39 | 6.32E-38 |
| Nup37 | 13.10 | 2.38 | 3.21E-39 | 6.49E-38 |
| Eif1ax | 13.09 | 1.82 | 3.88E-39 | 7.83E-38 |
| Celsr1 | 13.09 | 2.40 | 3.95E-39 | 7.94E-38 |
| Rbm3 | 13.08 | 1.50 | 4.23E-39 | 8.51E-38 |
| Ssrp1 | 13.08 | 1.84 | 4.28E-39 | 8.59E-38 |
| Lrrc31 | 13.07 | 1.74 | 4.95E-39 | 9.93E-38 |
| Rif1 | 13.07 | 1.94 | 5.14E-39 | 1.03E-37 |
| Trmt2a | 13.06 | 2.04 | 5.23E-39 | 1.05E-37 |
| Zfp422 | 13.06 | 1.85 | 5.61E-39 | 1.12E-37 |
| Usp14 | 13.06 | 2.02 | 5.88E-39 | 1.17E-37 |
| Dlgap5 | 13.05 | 3.04 | 6.51E-39 | 1.29E-37 |
| Hmmr | 13.04 | 3.03 | 7.26E-39 | 1.44E-37 |
| Haus4 | 13.04 | 2.17 | 7.72E-39 | 1.52E-37 |
| Snrpb | 13.03 | 1.94 | 8.02E-39 | 1.58E-37 |
| Stk39 | 13.03 | 1.72 | 8.46E-39 | 1.67E-37 |
| Atrx | 13.03 | 1.65 | 8.5E-39 | 1.67E-37 |
| Nol9 | 13.02 | 2.34 | 9.45E-39 | 1.86E-37 |
| Rbmx | 13.02 | 1.94 | 9.58E-39 | 1.88E-37 |
| Fam210a | 13.00 | 1.69 | 1.16E-38 | 2.28E-37 |
| Slfn9 | 12.99 | 3.47 | 1.32E-38 | 2.59E-37 |
| Rassf4 | 12.99 | 1.90 | 1.49E-38 | 2.9E-37 |
| Fam107b | 12.99 | 1.78 | 1.49E-38 | 2.9E-37 |
| Imp3 | 12.98 | 1.88 | 1.61E-38 | 3.13E-37 |
| Ctnnal1 | 12.95 | 2.55 | 2.26E-38 | 4.38E-37 |
| Ccdc58 | 12.95 | 1.69 | 2.27E-38 | 4.39E-37 |
| Gstcd | 12.95 | 2.32 | 2.5E-38 | 4.84E-37 |
| Poc1b | 12.93 | 1.96 | 2.86E-38 | 5.52E-37 |
| Tomm40l | 12.93 | 1.92 | 3.07E-38 | 5.92E-37 |
| Ppp1r8 | 12.90 | 2.11 | 4.31E-38 | 8.32E-37 |
| Prr11 | 12.90 | 2.52 | 4.5E-38 | 8.68E-37 |
| Ddx11 | 12.90 | 2.39 | 4.59E-38 | 8.84E-37 |
| Hmgn5 | 12.89 | 1.82 | 5.2E-38 | 1E-36 |

| | | | | |
|---|---|---|---|---|
| Nat10 | 12.88 | 2.48 | 5.86E-38 | 1.12E-36 |
| Mid1ip1 | 12.87 | 1.72 | 6.97E-38 | 1.33E-36 |
| Ppie | 12.86 | 2.05 | 7.13E-38 | 1.36E-36 |
| Itpa | 12.86 | 1.82 | 7.55E-38 | 1.44E-36 |
| Rad51b | 12.86 | 2.53 | 7.89E-38 | 1.51E-36 |
| Gcsh | 12.85 | 1.89 | 8.39E-38 | 1.6E-36 |
| Bex3 | 12.84 | 1.80 | 9.24E-38 | 1.75E-36 |
| Cops5 | 12.84 | 1.52 | 1.02E-37 | 1.93E-36 |
| Irak1 | 12.83 | 1.59 | 1.12E-37 | 2.11E-36 |
| Naa15 | 12.83 | 1.61 | 1.12E-37 | 2.12E-36 |
| Plagl2 | 12.82 | 1.69 | 1.26E-37 | 2.37E-36 |
| Memo1 | 12.79 | 1.67 | 1.95E-37 | 3.65E-36 |
| Nup62 | 12.77 | 2.31 | 2.48E-37 | 4.63E-36 |
| E2f6 | 12.77 | 2.83 | 2.55E-37 | 4.75E-36 |
| Ncapd3 | 12.76 | 2.37 | 2.6E-37 | 4.83E-36 |
| Nop2 | 12.76 | 2.25 | 2.69E-37 | 5E-36 |
| Khsrp | 12.76 | 1.61 | 2.78E-37 | 5.16E-36 |
| Itpka | 12.76 | 1.98 | 2.78E-37 | 5.16E-36 |
| Nde1 | 12.75 | 2.57 | 3.02E-37 | 5.59E-36 |
| Cct8 | 12.74 | 1.67 | 3.44E-37 | 6.36E-36 |
| Spata5 | 12.73 | 2.51 | 3.88E-37 | 7.17E-36 |
| Nop16 | 12.73 | 2.56 | 4.24E-37 | 7.82E-36 |
| Ammecr1 | 12.72 | 2.11 | 4.37E-37 | 8.05E-36 |
| Cnbp | 12.72 | 1.71 | 4.43E-37 | 8.13E-36 |
| Shtn1 | 12.72 | 1.80 | 4.53E-37 | 8.33E-36 |
| Gfer | 12.71 | 2.14 | 4.94E-37 | 9.06E-36 |
| Sltm | 12.71 | 1.53 | 5.45E-37 | 9.99E-36 |
| Nup210 | 12.70 | 2.04 | 6.22E-37 | 1.14E-35 |
| Knl1 | 12.69 | 2.91 | 6.69E-37 | 1.22E-35 |
| Baz1b | 12.68 | 1.54 | 7.21E-37 | 1.31E-35 |
| Cntrl | 12.68 | 2.23 | 7.28E-37 | 1.32E-35 |
| Ncbp2 | 12.67 | 1.86 | 9.08E-37 | 1.65E-35 |
| Rad18 | 12.64 | 2.80 | 1.22E-36 | 2.2E-35 |
| Tsn | 12.63 | 1.67 | 1.44E-36 | 2.6E-35 |
| Thyn1 | 12.62 | 2.07 | 1.58E-36 | 2.85E-35 |
| Pold3 | 12.62 | 1.92 | 1.61E-36 | 2.9E-35 |
| Cdkn2aipnl | 12.62 | 1.90 | 1.67E-36 | 3E-35 |
| Nfya | 12.61 | 1.69 | 1.84E-36 | 3.31E-35 |
| Dnajc9 | 12.60 | 1.90 | 2.13E-36 | 3.82E-35 |
| Zwilch | 12.59 | 2.75 | 2.52E-36 | 4.49E-35 |
| Noxa1 | 12.57 | 1.72 | 3.25E-36 | 5.79E-35 |
| Fen1 | 12.56 | 2.27 | 3.44E-36 | 6.12E-35 |
| Usp43 | 12.55 | 1.59 | 3.9E-36 | 6.94E-35 |
| Haspin | 12.55 | 2.50 | 4.23E-36 | 7.52E-35 |

| | | | | |
|---|---|---|---|---|
| Ddx56 | 12.54 | 2.06 | 4.74E-36 | 8.4E-35 |
| Upf3b | 12.53 | 1.64 | 4.85E-36 | 8.58E-35 |
| Plekha5 | 12.52 | 1.50 | 5.76E-36 | 1.02E-34 |
| Dnajc2 | 12.51 | 1.91 | 6.61E-36 | 1.17E-34 |
| Trim27 | 12.50 | 1.68 | 7.31E-36 | 1.29E-34 |
| Gsdmc4 | 12.50 | 2.08 | 7.42E-36 | 1.31E-34 |
| Psmd12 | 12.50 | 1.56 | 7.59E-36 | 1.33E-34 |
| Itgam | 12.50 | 2.23 | 7.74E-36 | 1.36E-34 |
| Eif3b | 12.49 | 1.63 | 8.56E-36 | 1.5E-34 |
| Timm21 | 12.48 | 1.98 | 9.78E-36 | 1.71E-34 |
| Pprc1 | 12.47 | 2.43 | 1.14E-35 | 1.98E-34 |
| Aen | 12.47 | 2.88 | 1.14E-35 | 1.99E-34 |
| Mis18bp1 | 12.46 | 2.96 | 1.17E-35 | 2.03E-34 |
| Slc38a1 | 12.46 | 1.57 | 1.24E-35 | 2.15E-34 |
| Parva | 12.46 | 1.96 | 1.3E-35 | 2.25E-34 |
| Tcof1 | 12.45 | 2.06 | 1.38E-35 | 2.38E-34 |
| Tars2 | 12.45 | 2.10 | 1.42E-35 | 2.44E-34 |
| Mars | 12.44 | 1.93 | 1.5E-35 | 2.58E-34 |
| Pds5b | 12.44 | 2.22 | 1.59E-35 | 2.73E-34 |
| Rexo2 | 12.44 | 1.64 | 1.67E-35 | 2.87E-34 |
| Hspg2 | 12.44 | 1.96 | 1.68E-35 | 2.89E-34 |
| Thumpd1 | 12.43 | 2.29 | 1.74E-35 | 2.99E-34 |
| Ncapg | 12.43 | 2.22 | 1.81E-35 | 3.11E-34 |
| Slc44a3 | 12.43 | 1.88 | 1.86E-35 | 3.19E-34 |
| Hnrnph1 | 12.42 | 1.76 | 1.95E-35 | 3.34E-34 |
| Fkbp5 | 12.42 | 1.79 | 2.09E-35 | 3.56E-34 |
| Rab15 | 12.42 | 1.55 | 2.09E-35 | 3.57E-34 |
| Zgrf1 | 12.42 | 3.21 | 2.12E-35 | 3.62E-34 |
| Pdhb | 12.40 | 1.60 | 2.55E-35 | 4.34E-34 |
| Sass6 | 12.40 | 2.42 | 2.71E-35 | 4.6E-34 |
| Elac2 | 12.39 | 2.37 | 2.97E-35 | 5.03E-34 |
| Krr1 | 12.39 | 2.16 | 3.15E-35 | 5.32E-34 |
| Ppia | 12.38 | 1.97 | 3.18E-35 | 5.37E-34 |
| Gsdmc2 | 12.38 | 2.20 | 3.19E-35 | 5.39E-34 |
| L3mbtl2 | 12.38 | 2.59 | 3.44E-35 | 5.8E-34 |
| Smarcad1 | 12.37 | 2.06 | 3.64E-35 | 6.12E-34 |
| Tamm41 | 12.37 | 2.23 | 3.71E-35 | 6.25E-34 |
| Atm | 12.37 | 2.26 | 3.75E-35 | 6.3E-34 |
| Swi5 | 12.37 | 1.64 | 3.9E-35 | 6.55E-34 |
| Rfwd3 | 12.36 | 1.70 | 4.16E-35 | 6.97E-34 |
| Agpat5 | 12.35 | 2.46 | 4.71E-35 | 7.87E-34 |
| Osgep | 12.34 | 2.05 | 5.2E-35 | 8.68E-34 |
| Pgam5 | 12.31 | 1.83 | 7.73E-35 | 1.28E-33 |
| Eif4e | 12.31 | 1.58 | 7.8E-35 | 1.29E-33 |

| | | | | |
|---|---|---|---|---|
| Rpp14 | 12.30 | 1.89 | 8.77E-35 | 1.45E-33 |
| Slc7a1 | 12.30 | 2.62 | 8.82E-35 | 1.46E-33 |
| Blnk | 12.30 | 2.30 | 9.44E-35 | 1.56E-33 |
| Klk1 | 12.28 | 1.62 | 1.09E-34 | 1.81E-33 |
| Med14 | 12.28 | 1.66 | 1.22E-34 | 2E-33 |
| Card10 | 12.27 | 1.67 | 1.27E-34 | 2.1E-33 |
| Hist1h2af | 12.27 | 3.06 | 1.3E-34 | 2.13E-33 |
| Mcph1 | 12.27 | 2.36 | 1.35E-34 | 2.22E-33 |
| Tiam1 | 12.27 | 2.54 | 1.38E-34 | 2.27E-33 |
| Txnl4a | 12.27 | 1.90 | 1.39E-34 | 2.28E-33 |
| Ckap2 | 12.26 | 2.62 | 1.55E-34 | 2.54E-33 |
| Thop1 | 12.24 | 2.13 | 1.84E-34 | 3E-33 |
| Pkp1 | 12.24 | 2.27 | 1.9E-34 | 3.1E-33 |
| Rcc1l | 12.24 | 2.26 | 1.9E-34 | 3.1E-33 |
| Pole | 12.24 | 2.59 | 1.93E-34 | 3.14E-33 |
| Hells | 12.23 | 2.12 | 2.13E-34 | 3.47E-33 |
| Mdn1 | 12.22 | 2.68 | 2.41E-34 | 3.91E-33 |
| Mboat1 | 12.19 | 2.36 | 3.36E-34 | 5.45E-33 |
| Ints1 | 12.19 | 1.53 | 3.65E-34 | 5.9E-33 |
| Rrn3 | 12.19 | 1.72 | 3.72E-34 | 6E-33 |
| Wdr74 | 12.18 | 2.39 | 3.74E-34 | 6.03E-33 |
| Ybx2 | 12.18 | 2.56 | 4.14E-34 | 6.66E-33 |
| Ttk | 12.18 | 2.99 | 4.16E-34 | 6.68E-33 |
| Usp48 | 12.18 | 1.61 | 4.22E-34 | 6.76E-33 |
| Hsph1 | 12.17 | 2.30 | 4.45E-34 | 7.12E-33 |
| Dcun1d5 | 12.17 | 1.53 | 4.57E-34 | 7.3E-33 |
| Eed | 12.16 | 1.54 | 4.82E-34 | 7.7E-33 |
| Trub1 | 12.16 | 2.22 | 4.95E-34 | 7.88E-33 |
| Banf1 | 12.16 | 2.04 | 4.96E-34 | 7.89E-33 |
| Zfp451 | 12.16 | 1.81 | 5.15E-34 | 8.18E-33 |
| Lrfn4 | 12.16 | 2.16 | 5.17E-34 | 8.21E-33 |
| Usp1 | 12.15 | 1.63 | 5.52E-34 | 8.76E-33 |
| Slc16a1 | 12.15 | 1.54 | 5.77E-34 | 9.15E-33 |
| Ptpn18 | 12.15 | 1.80 | 5.86E-34 | 9.29E-33 |
| Cmc2 | 12.13 | 2.11 | 7.37E-34 | 1.16E-32 |
| Bckdk | 12.13 | 1.58 | 7.42E-34 | 1.17E-32 |
| Cyp39a1 | 12.12 | 3.08 | 7.93E-34 | 1.25E-32 |
| Hspe1 | 12.12 | 2.39 | 7.95E-34 | 1.25E-32 |
| Cables2 | 12.12 | 1.54 | 8.05E-34 | 1.26E-32 |
| Akip1 | 12.12 | 2.02 | 8.3E-34 | 1.3E-32 |
| Cpsf3 | 12.11 | 1.74 | 9.13E-34 | 1.43E-32 |
| Cit | 12.11 | 2.60 | 9.77E-34 | 1.53E-32 |
| Lsm8 | 12.11 | 1.73 | 9.82E-34 | 1.53E-32 |
| L1cam | 12.10 | 2.05 | 1.01E-33 | 1.57E-32 |

| | | | | |
|---|---|---|---|---|
| Lsm5 | 12.10 | 2.21 | 1.04E-33 | 1.62E-32 |
| Snrpb2 | 12.08 | 1.75 | 1.35E-33 | 2.1E-32 |
| Gcn1 | 12.07 | 1.65 | 1.45E-33 | 2.25E-32 |
| Lig3 | 12.07 | 2.01 | 1.48E-33 | 2.3E-32 |
| Rnmt | 12.06 | 1.75 | 1.79E-33 | 2.77E-32 |
| Mcm3 | 12.06 | 2.29 | 1.8E-33 | 2.78E-32 |
| Trmt10c | 12.04 | 2.24 | 2.11E-33 | 3.24E-32 |
| Aurka | 12.03 | 2.51 | 2.45E-33 | 3.76E-32 |
| Pfdn2 | 12.03 | 1.90 | 2.51E-33 | 3.85E-32 |
| Sgo1 | 12.03 | 3.01 | 2.52E-33 | 3.87E-32 |
| Srp19 | 12.03 | 1.51 | 2.59E-33 | 3.96E-32 |
| Ccl28 | 12.02 | 1.65 | 2.88E-33 | 4.4E-32 |
| Sorl1 | 12.01 | 1.71 | 3.03E-33 | 4.63E-32 |
| Prdx6 | 11.99 | 1.51 | 4.15E-33 | 6.33E-32 |
| Sgsm3 | 11.98 | 1.62 | 4.4E-33 | 6.71E-32 |
| Coq7 | 11.98 | 1.78 | 4.74E-33 | 7.21E-32 |
| Drg2 | 11.97 | 1.62 | 4.81E-33 | 7.31E-32 |
| Ptpn2 | 11.97 | 1.55 | 5.02E-33 | 7.63E-32 |
| Kat2a | 11.97 | 2.24 | 5.1E-33 | 7.73E-32 |
| Trim59 | 11.97 | 2.84 | 5.16E-33 | 7.82E-32 |
| Slco2b1 | 11.96 | 1.94 | 6.03E-33 | 9.13E-32 |
| Chek2 | 11.94 | 2.48 | 7E-33 | 1.05E-31 |
| Adi1 | 11.94 | 2.08 | 7.11E-33 | 1.07E-31 |
| Engase | 11.94 | 2.45 | 7.36E-33 | 1.11E-31 |
| Lyrm4 | 11.93 | 2.19 | 8.33E-33 | 1.25E-31 |
| Rpf1 | 11.93 | 1.56 | 8.4E-33 | 1.26E-31 |
| Clca3a2 | 11.92 | 1.88 | 8.96E-33 | 1.34E-31 |
| Pwp1 | 11.92 | 2.35 | 9.46E-33 | 1.41E-31 |
| Gcg | 11.92 | 2.39 | 9.7E-33 | 1.45E-31 |
| Dbf4 | 11.91 | 2.34 | 1.04E-32 | 1.54E-31 |
| Ano7 | 11.91 | 1.80 | 1.04E-32 | 1.54E-31 |
| Smc3 | 11.91 | 1.81 | 1.04E-32 | 1.55E-31 |
| Vrk1 | 11.90 | 2.29 | 1.2E-32 | 1.78E-31 |
| Nudt19 | 11.90 | 1.55 | 1.23E-32 | 1.83E-31 |
| Man2c1 | 11.86 | 1.77 | 1.95E-32 | 2.87E-31 |
| Orc6 | 11.86 | 2.46 | 1.95E-32 | 2.88E-31 |
| Ethe1 | 11.84 | 1.55 | 2.42E-32 | 3.56E-31 |
| Pitrm1 | 11.83 | 2.24 | 2.67E-32 | 3.92E-31 |
| Ankrd10 | 11.83 | 1.65 | 2.75E-32 | 4.03E-31 |
| Ckap5 | 11.82 | 1.74 | 2.92E-32 | 4.27E-31 |
| Rap1gap | 11.82 | 1.69 | 2.95E-32 | 4.31E-31 |
| Cchcr1 | 11.81 | 2.74 | 3.37E-32 | 4.93E-31 |
| Ivd | 11.81 | 1.69 | 3.51E-32 | 5.13E-31 |
| Trmt1 | 11.80 | 1.84 | 3.71E-32 | 5.41E-31 |

| | | | | |
|---|---|---|---|---|
| Nadk2 | 11.80 | 1.72 | 4.06E-32 | 5.92E-31 |
| Auts2 | 11.79 | 2.11 | 4.15E-32 | 6.05E-31 |
| Polr3k | 11.79 | 1.75 | 4.37E-32 | 6.36E-31 |
| Acad9 | 11.79 | 1.88 | 4.53E-32 | 6.58E-31 |
| Ahctf1 | 11.79 | 1.57 | 4.62E-32 | 6.71E-31 |
| Plk4 | 11.79 | 2.40 | 4.64E-32 | 6.73E-31 |
| Cenpw | 11.78 | 2.51 | 4.81E-32 | 6.97E-31 |
| Msh6 | 11.78 | 2.37 | 4.98E-32 | 7.21E-31 |
| Grk6 | 11.78 | 1.80 | 5.03E-32 | 7.28E-31 |
| Nup205 | 11.78 | 1.88 | 5.03E-32 | 7.28E-31 |
| Dctpp1 | 11.77 | 2.30 | 5.65E-32 | 8.16E-31 |
| Tsfm | 11.77 | 2.29 | 5.9E-32 | 8.52E-31 |
| Rae1 | 11.75 | 1.58 | 6.73E-32 | 9.68E-31 |
| Fanca | 11.75 | 2.44 | 6.89E-32 | 9.88E-31 |
| Polr2b | 11.72 | 1.72 | 1.03E-31 | 1.47E-30 |
| Neil3 | 11.71 | 2.54 | 1.18E-31 | 1.69E-30 |
| Ints7 | 11.70 | 2.15 | 1.28E-31 | 1.83E-30 |
| Prmt3 | 11.70 | 2.66 | 1.32E-31 | 1.87E-30 |
| Tpd52l2 | 11.69 | 1.62 | 1.39E-31 | 1.98E-30 |
| Lipg | 11.69 | 2.56 | 1.49E-31 | 2.11E-30 |
| Erfe | 11.69 | 3.15 | 1.52E-31 | 2.15E-30 |
| Gpn3 | 11.68 | 2.07 | 1.58E-31 | 2.23E-30 |
| Hnrnpl | 11.66 | 1.51 | 1.94E-31 | 2.74E-30 |
| Scd2 | 11.65 | 1.59 | 2.21E-31 | 3.11E-30 |
| Dph6 | 11.64 | 2.37 | 2.49E-31 | 3.5E-30 |
| Tpsg1 | 11.63 | 1.87 | 2.85E-31 | 3.99E-30 |
| Plce1 | 11.62 | 1.74 | 3.1E-31 | 4.34E-30 |
| Tgif1 | 11.62 | 1.68 | 3.18E-31 | 4.45E-30 |
| Thap12 | 11.62 | 1.65 | 3.39E-31 | 4.73E-30 |
| Rasd2 | 11.62 | 1.82 | 3.45E-31 | 4.81E-30 |
| Ptprs | 11.61 | 2.11 | 3.84E-31 | 5.36E-30 |
| Cpt1a | 11.60 | 1.76 | 4E-31 | 5.58E-30 |
| Nle1 | 11.59 | 2.91 | 4.6E-31 | 6.4E-30 |
| Pitpnb | 11.59 | 1.54 | 4.71E-31 | 6.55E-30 |
| Dnajc11 | 11.58 | 1.81 | 5.17E-31 | 7.19E-30 |
| Bcl7c | 11.58 | 1.65 | 5.38E-31 | 7.46E-30 |
| Trp53bp1 | 11.57 | 2.21 | 5.61E-31 | 7.78E-30 |
| Ccdc88c | 11.57 | 1.63 | 5.62E-31 | 7.79E-30 |
| Smim6 | 11.57 | 1.61 | 5.83E-31 | 8.06E-30 |
| Mcee | 11.56 | 2.03 | 6.77E-31 | 9.36E-30 |
| Tmem70 | 11.55 | 1.68 | 7.28E-31 | 1.01E-29 |
| Xpo4 | 11.55 | 2.23 | 7.62E-31 | 1.05E-29 |
| Mmd | 11.55 | 1.68 | 7.63E-31 | 1.05E-29 |
| Fam136a | 11.54 | 1.65 | 8.3E-31 | 1.14E-29 |

| | | | | |
|---|---|---|---|---|
| Uqcc1 | 11.54 | 1.76 | 8.46E-31 | 1.16E-29 |
| Nup188 | 11.54 | 2.21 | 8.48E-31 | 1.16E-29 |
| Aspm | 11.54 | 2.60 | 8.48E-31 | 1.16E-29 |
| Gusb | 11.53 | 1.67 | 8.99E-31 | 1.23E-29 |
| Paxip1 | 11.52 | 1.78 | 1.05E-30 | 1.44E-29 |
| Cip2a | 11.52 | 2.25 | 1.08E-30 | 1.47E-29 |
| Hmgn1 | 11.52 | 2.68 | 1.1E-30 | 1.5E-29 |
| Uri1 | 11.51 | 1.82 | 1.15E-30 | 1.57E-29 |
| Ncbp1 | 11.50 | 1.82 | 1.29E-30 | 1.76E-29 |
| Gphn | 11.49 | 2.49 | 1.41E-30 | 1.91E-29 |
| Kcnk1 | 11.49 | 1.69 | 1.46E-30 | 1.97E-29 |
| Chaf1a | 11.49 | 2.28 | 1.55E-30 | 2.1E-29 |
| Cct5 | 11.49 | 2.20 | 1.56E-30 | 2.12E-29 |
| Cdo1 | 11.48 | 2.04 | 1.61E-30 | 2.18E-29 |
| Crls1 | 11.48 | 1.57 | 1.7E-30 | 2.3E-29 |
| Exosc8 | 11.47 | 2.59 | 1.9E-30 | 2.57E-29 |
| Stambpl1 | 11.47 | 1.63 | 1.95E-30 | 2.63E-29 |
| Sms | 11.46 | 1.87 | 2.05E-30 | 2.75E-29 |
| Dars2 | 11.46 | 2.08 | 2.16E-30 | 2.9E-29 |
| Cpsf1 | 11.45 | 1.67 | 2.39E-30 | 3.21E-29 |
| Otulin | 11.45 | 1.73 | 2.46E-30 | 3.3E-29 |
| Ehmt1 | 11.44 | 1.67 | 2.68E-30 | 3.58E-29 |
| Rpa2 | 11.44 | 2.04 | 2.75E-30 | 3.68E-29 |
| Cct4 | 11.44 | 1.73 | 2.79E-30 | 3.72E-29 |
| Cep250 | 11.43 | 2.07 | 3.05E-30 | 4.07E-29 |
| Ccdc86 | 11.43 | 2.32 | 3.08E-30 | 4.11E-29 |
| Gpatch4 | 11.42 | 2.33 | 3.28E-30 | 4.36E-29 |
| Eif2b5 | 11.42 | 1.52 | 3.39E-30 | 4.51E-29 |
| Eif1b | 11.42 | 1.72 | 3.43E-30 | 4.55E-29 |
| Arhgap19 | 11.41 | 2.03 | 3.82E-30 | 5.07E-29 |
| Ppat | 11.40 | 2.45 | 4.24E-30 | 5.62E-29 |
| Fkbp3 | 11.39 | 1.96 | 4.47E-30 | 5.91E-29 |
| Mbnl3 | 11.39 | 1.88 | 4.61E-30 | 6.1E-29 |
| Gle1 | 11.38 | 1.87 | 4.99E-30 | 6.6E-29 |
| Stoml2 | 11.38 | 1.76 | 5.04E-30 | 6.65E-29 |
| Cc2d2a | 11.38 | 2.77 | 5.22E-30 | 6.88E-29 |
| Rnaseh2b | 11.38 | 1.99 | 5.34E-30 | 7.04E-29 |
| Psmd14 | 11.37 | 1.56 | 6.12E-30 | 8.06E-29 |
| Dcaf1 | 11.36 | 1.87 | 6.99E-30 | 9.18E-29 |
| Sgf29 | 11.35 | 1.99 | 7.05E-30 | 9.24E-29 |
| Cep55 | 11.35 | 2.87 | 7.08E-30 | 9.28E-29 |
| Yrdc | 11.35 | 2.00 | 7.75E-30 | 1.01E-28 |
| Rph3al | 11.35 | 1.59 | 7.76E-30 | 1.01E-28 |
| Dus3l | 11.34 | 1.79 | 7.87E-30 | 1.03E-28 |

| | | | | |
|---|---|---|---|---|
| Dctd | 11.34 | 2.72 | 8.05E-30 | 1.05E-28 |
| Nol6 | 11.34 | 2.25 | 8.28E-30 | 1.08E-28 |
| Coq5 | 11.34 | 2.02 | 8.47E-30 | 1.1E-28 |
| Bcl11b | 11.34 | 2.04 | 8.71E-30 | 1.13E-28 |
| Atad2 | 11.34 | 2.18 | 8.79E-30 | 1.14E-28 |
| Cinp | 11.33 | 1.91 | 9.24E-30 | 1.2E-28 |
| Mipep | 11.33 | 2.02 | 9.79E-30 | 1.27E-28 |
| Kif14 | 11.32 | 3.06 | 1.08E-29 | 1.4E-28 |
| Cad | 11.32 | 1.90 | 1.09E-29 | 1.41E-28 |
| Wdr46 | 11.31 | 1.88 | 1.23E-29 | 1.58E-28 |
| Cbx3 | 11.30 | 1.70 | 1.29E-29 | 1.66E-28 |
| Mettl5 | 11.30 | 1.71 | 1.34E-29 | 1.72E-28 |
| Gfm1 | 11.30 | 1.74 | 1.37E-29 | 1.76E-28 |
| N6amt1 | 11.30 | 2.56 | 1.37E-29 | 1.76E-28 |
| Fundc2 | 11.29 | 2.11 | 1.47E-29 | 1.89E-28 |
| Prps2 | 11.29 | 2.04 | 1.48E-29 | 1.89E-28 |
| Xpo5 | 11.29 | 2.01 | 1.49E-29 | 1.91E-28 |
| Dynll1 | 11.28 | 1.62 | 1.6E-29 | 2.05E-28 |
| Eif3c | 11.28 | 1.51 | 1.61E-29 | 2.06E-28 |
| Nmt2 | 11.28 | 2.26 | 1.71E-29 | 2.18E-28 |
| Miip | 11.27 | 1.93 | 1.78E-29 | 2.26E-28 |
| Dach1 | 11.27 | 1.86 | 1.87E-29 | 2.38E-28 |
| Cab39l | 11.24 | 1.97 | 2.53E-29 | 3.21E-28 |
| Ice1 | 11.24 | 1.83 | 2.58E-29 | 3.27E-28 |
| Kdm2b | 11.24 | 2.02 | 2.67E-29 | 3.38E-28 |
| Ndufa9 | 11.22 | 1.53 | 3.18E-29 | 4.03E-28 |
| Nr5a2 | 11.22 | 1.53 | 3.33E-29 | 4.22E-28 |
| Tubg1 | 11.22 | 2.21 | 3.34E-29 | 4.22E-28 |
| Qtrt2 | 11.21 | 2.45 | 3.47E-29 | 4.39E-28 |
| Hmbs | 11.21 | 1.91 | 3.58E-29 | 4.52E-28 |
| Dcps | 11.20 | 2.18 | 3.85E-29 | 4.87E-28 |
| Grhl2 | 11.20 | 2.15 | 3.96E-29 | 5E-28 |
| Bnip1 | 11.20 | 1.75 | 3.96E-29 | 5E-28 |
| Lrch1 | 11.20 | 1.68 | 3.97E-29 | 5E-28 |
| Ezh2 | 11.20 | 1.51 | 4.16E-29 | 5.24E-28 |
| Rfc5 | 11.20 | 2.24 | 4.27E-29 | 5.37E-28 |
| Pla2g5 | 11.19 | 2.61 | 4.47E-29 | 5.62E-28 |
| Ddx10 | 11.19 | 1.95 | 4.67E-29 | 5.87E-28 |
| Foxk1 | 11.19 | 1.57 | 4.74E-29 | 5.95E-28 |
| Znrd1 | 11.18 | 1.60 | 5.06E-29 | 6.34E-28 |
| Nup93 | 11.18 | 2.45 | 5.29E-29 | 6.63E-28 |
| Gmnn | 11.18 | 1.76 | 5.36E-29 | 6.7E-28 |
| Polr3h | 11.15 | 2.53 | 7.15E-29 | 8.89E-28 |
| Gemin6 | 11.15 | 2.49 | 7.48E-29 | 9.28E-28 |

| | | | | |
|---|---|---|---|---|
| Rex1bd | 11.14 | 1.71 | 7.89E-29 | 9.8E-28 |
| Utp20 | 11.14 | 2.30 | 8.34E-29 | 1.03E-27 |
| Rad50 | 11.13 | 1.71 | 8.85E-29 | 1.1E-27 |
| BC003965 | 11.13 | 1.83 | 9.02E-29 | 1.12E-27 |
| Smim11 | 11.11 | 1.63 | 1.16E-28 | 1.43E-27 |
| Tcf19 | 11.11 | 2.50 | 1.18E-28 | 1.45E-27 |
| Mif | 11.10 | 2.00 | 1.23E-28 | 1.51E-27 |
| Cdc25c | 11.10 | 3.50 | 1.28E-28 | 1.57E-27 |
| Fars2 | 11.10 | 1.51 | 1.32E-28 | 1.62E-27 |
| Gcdh | 11.09 | 1.92 | 1.48E-28 | 1.82E-27 |
| Lrwd1 | 11.08 | 2.05 | 1.53E-28 | 1.88E-27 |
| Terf1 | 11.08 | 1.92 | 1.54E-28 | 1.89E-27 |
| Tln2 | 11.07 | 2.43 | 1.7E-28 | 2.08E-27 |
| Ralgapa1 | 11.07 | 1.60 | 1.82E-28 | 2.22E-27 |
| Wnk2 | 11.06 | 1.65 | 1.91E-28 | 2.33E-27 |
| Guf1 | 11.06 | 1.73 | 1.95E-28 | 2.37E-27 |
| Nup214 | 11.06 | 1.84 | 1.97E-28 | 2.41E-27 |
| St6galnac2 | 11.06 | 1.52 | 2.02E-28 | 2.46E-27 |
| Slc25a17 | 11.05 | 1.63 | 2.13E-28 | 2.59E-27 |
| Plch1 | 11.05 | 1.89 | 2.13E-28 | 2.6E-27 |
| Thoc3 | 11.04 | 2.02 | 2.36E-28 | 2.87E-27 |
| Syce2 | 11.02 | 2.81 | 2.95E-28 | 3.58E-27 |
| Ddx27 | 11.01 | 1.70 | 3.29E-28 | 3.99E-27 |
| Imp4 | 11.01 | 1.82 | 3.6E-28 | 4.36E-27 |
| Wdr61 | 11.00 | 1.73 | 3.78E-28 | 4.57E-27 |
| Ppp1r14b | 10.99 | 1.87 | 4.09E-28 | 4.93E-27 |
| Selenoh | 10.99 | 2.27 | 4.22E-28 | 5.08E-27 |
| Aspg | 10.99 | 2.24 | 4.37E-28 | 5.27E-27 |
| Tsen34 | 10.97 | 1.58 | 5.18E-28 | 6.22E-27 |
| Grwd1 | 10.97 | 2.46 | 5.25E-28 | 6.3E-27 |
| Spdl1 | 10.97 | 3.22 | 5.34E-28 | 6.4E-27 |
| Patz1 | 10.97 | 1.91 | 5.62E-28 | 6.73E-27 |
| Hdac7 | 10.96 | 2.10 | 5.83E-28 | 6.98E-27 |
| Zfp692 | 10.96 | 1.96 | 6.14E-28 | 7.34E-27 |
| Zfp263 | 10.95 | 1.83 | 6.63E-28 | 7.92E-27 |
| Supv3l1 | 10.95 | 2.16 | 6.94E-28 | 8.27E-27 |
| Rad51ap1 | 10.94 | 2.67 | 7.47E-28 | 8.89E-27 |
| Pllp | 10.93 | 1.62 | 8.22E-28 | 9.79E-27 |
| Gtf2h4 | 10.93 | 1.85 | 8.24E-28 | 9.8E-27 |
| Efcab11 | 10.92 | 3.60 | 8.82E-28 | 1.05E-26 |
| Slc6a7 | 10.91 | 2.28 | 1.02E-27 | 1.21E-26 |
| Zmynd19 | 10.91 | 2.19 | 1.09E-27 | 1.29E-26 |
| Spout1 | 10.90 | 1.85 | 1.12E-27 | 1.33E-26 |
| Eef1akmt2 | 10.90 | 1.77 | 1.16E-27 | 1.37E-26 |

| | | | | |
|---|---|---|---|---|
| Noc3l | 10.87 | 2.81 | 1.55E-27 | 1.83E-26 |
| Ak6 | 10.87 | 1.79 | 1.56E-27 | 1.83E-26 |
| Acbd6 | 10.87 | 1.84 | 1.68E-27 | 1.97E-26 |
| Ddx20 | 10.86 | 2.43 | 1.72E-27 | 2.01E-26 |
| Nsa2 | 10.86 | 1.82 | 1.81E-27 | 2.12E-26 |
| Usp39 | 10.86 | 1.84 | 1.82E-27 | 2.13E-26 |
| Spag5 | 10.85 | 2.37 | 1.99E-27 | 2.33E-26 |
| Rwdd1 | 10.85 | 1.77 | 2.08E-27 | 2.43E-26 |
| Jade2 | 10.84 | 2.14 | 2.14E-27 | 2.49E-26 |
| Tent4a | 10.84 | 1.82 | 2.21E-27 | 2.57E-26 |
| Lsm6 | 10.84 | 1.99 | 2.25E-27 | 2.62E-26 |
| Eef1e1 | 10.84 | 2.02 | 2.35E-27 | 2.73E-26 |
| Znhit3 | 10.83 | 2.29 | 2.4E-27 | 2.79E-26 |
| Asf1a | 10.83 | 2.18 | 2.42E-27 | 2.81E-26 |
| Ssbp3 | 10.83 | 1.67 | 2.43E-27 | 2.82E-26 |
| Prim2 | 10.83 | 2.70 | 2.45E-27 | 2.85E-26 |
| Znhit6 | 10.83 | 2.19 | 2.47E-27 | 2.86E-26 |
| Smc2 | 10.82 | 2.50 | 2.87E-27 | 3.32E-26 |
| Cfap20 | 10.80 | 1.64 | 3.55E-27 | 4.1E-26 |
| Cnot7 | 10.79 | 1.53 | 3.67E-27 | 4.23E-26 |
| Sac3d1 | 10.79 | 2.19 | 3.74E-27 | 4.31E-26 |
| Pom121 | 10.79 | 1.55 | 3.78E-27 | 4.35E-26 |
| Snrnp25 | 10.77 | 2.36 | 4.89E-27 | 5.62E-26 |
| Nup153 | 10.76 | 1.67 | 5.09E-27 | 5.84E-26 |
| Ska2 | 10.76 | 2.18 | 5.27E-27 | 6.04E-26 |
| Foxn2 | 10.76 | 1.52 | 5.3E-27 | 6.07E-26 |
| Brca1 | 10.76 | 2.80 | 5.52E-27 | 6.32E-26 |
| Sapcd2 | 10.74 | 2.50 | 6.46E-27 | 7.37E-26 |
| Bag5 | 10.74 | 1.86 | 6.54E-27 | 7.46E-26 |
| Phip | 10.69 | 1.50 | 1.08E-26 | 1.23E-25 |
| Pole3 | 10.69 | 1.91 | 1.09E-26 | 1.23E-25 |
| Eif3m | 10.69 | 2.16 | 1.11E-26 | 1.25E-25 |
| Cks2 | 10.69 | 2.27 | 1.14E-26 | 1.29E-25 |
| Atxn2 | 10.68 | 1.59 | 1.31E-26 | 1.47E-25 |
| Tmem201 | 10.67 | 2.07 | 1.42E-26 | 1.6E-25 |
| Rpp21 | 10.67 | 1.56 | 1.46E-26 | 1.64E-25 |
| Abcf2 | 10.66 | 1.94 | 1.57E-26 | 1.76E-25 |
| Khdrbs3 | 10.66 | 1.85 | 1.64E-26 | 1.84E-25 |
| Ubr7 | 10.65 | 1.89 | 1.69E-26 | 1.89E-25 |
| Nmd3 | 10.65 | 1.75 | 1.8E-26 | 2E-25 |
| Eif2s3x | 10.65 | 1.61 | 1.83E-26 | 2.04E-25 |
| Nqo2 | 10.64 | 1.57 | 1.86E-26 | 2.07E-25 |
| Haus7 | 10.64 | 2.02 | 1.89E-26 | 2.11E-25 |
| Nectin1 | 10.63 | 1.92 | 2.19E-26 | 2.44E-25 |

| | | | | |
|---|---|---|---|---|
| Scly | 10.63 | 1.53 | 2.24E-26 | 2.49E-25 |
| Tmem63a | 10.62 | 1.52 | 2.41E-26 | 2.68E-25 |
| Magohb | 10.62 | 1.96 | 2.45E-26 | 2.72E-25 |
| Timm9 | 10.61 | 1.96 | 2.56E-26 | 2.83E-25 |
| Oip5 | 10.60 | 3.14 | 2.93E-26 | 3.24E-25 |
| Ube2v2 | 10.60 | 1.79 | 2.96E-26 | 3.27E-25 |
| Kif20a | 10.60 | 2.62 | 3.12E-26 | 3.44E-25 |
| Odf2 | 10.59 | 1.67 | 3.3E-26 | 3.64E-25 |
| Tmco4 | 10.58 | 1.65 | 3.65E-26 | 4.02E-25 |
| Eif3a | 10.57 | 1.78 | 3.95E-26 | 4.34E-25 |
| Rrp9 | 10.57 | 2.46 | 3.98E-26 | 4.37E-25 |
| Msh2 | 10.56 | 1.64 | 4.7E-26 | 5.16E-25 |
| Zranb2 | 10.55 | 1.83 | 4.98E-26 | 5.46E-25 |
| Myef2 | 10.55 | 1.76 | 5.02E-26 | 5.5E-25 |
| Dis3 | 10.55 | 2.27 | 5.21E-26 | 5.71E-25 |
| Eef1akmt4 | 10.54 | 2.38 | 5.58E-26 | 6.1E-25 |
| Wdr18 | 10.53 | 2.24 | 6.05E-26 | 6.62E-25 |
| Cebpz | 10.53 | 2.06 | 6.28E-26 | 6.86E-25 |
| Spdef | 10.53 | 1.55 | 6.56E-26 | 7.15E-25 |
| Psip1 | 10.52 | 1.72 | 7.12E-26 | 7.76E-25 |
| Med22 | 10.52 | 1.83 | 7.14E-26 | 7.78E-25 |
| Eefsec | 10.52 | 2.06 | 7.31E-26 | 7.96E-25 |
| Ncapg2 | 10.51 | 2.42 | 7.74E-26 | 8.42E-25 |
| Gatd3a | 10.49 | 1.82 | 9.43E-26 | 1.02E-24 |
| Cacybp | 10.48 | 1.80 | 1.01E-25 | 1.1E-24 |
| Cebpzos | 10.48 | 1.55 | 1.08E-25 | 1.17E-24 |
| Kif20b | 10.46 | 2.57 | 1.33E-25 | 1.43E-24 |
| Pomt1 | 10.46 | 2.26 | 1.35E-25 | 1.45E-24 |
| Gins1 | 10.46 | 2.28 | 1.38E-25 | 1.48E-24 |
| Bola3 | 10.45 | 1.58 | 1.41E-25 | 1.51E-24 |
| Prelid2 | 10.45 | 2.03 | 1.46E-25 | 1.56E-24 |
| Tmem109 | 10.44 | 1.80 | 1.62E-25 | 1.73E-24 |
| Parp2 | 10.43 | 1.71 | 1.76E-25 | 1.88E-24 |
| Naa40 | 10.43 | 1.92 | 1.77E-25 | 1.89E-24 |
| Nrm | 10.41 | 2.71 | 2.26E-25 | 2.4E-24 |
| Nipsnap1 | 10.40 | 1.87 | 2.41E-25 | 2.55E-24 |
| Fra10ac1 | 10.40 | 1.59 | 2.53E-25 | 2.67E-24 |
| Exosc5 | 10.40 | 1.73 | 2.61E-25 | 2.76E-24 |
| Zpr1 | 10.39 | 1.83 | 2.62E-25 | 2.77E-24 |
| Gne | 10.38 | 1.51 | 2.99E-25 | 3.15E-24 |
| Pgm2l1 | 10.38 | 2.27 | 3.11E-25 | 3.28E-24 |
| Mzt2 | 10.37 | 1.82 | 3.42E-25 | 3.59E-24 |
| Metap1d | 10.36 | 1.85 | 3.95E-25 | 4.14E-24 |
| Adprhl2 | 10.35 | 1.91 | 3.97E-25 | 4.17E-24 |

| | | | | |
|---|---|---|---|---|
| Ylpm1 | 10.35 | 1.60 | 4.01E-25 | 4.2E-24 |
| Rasa3 | 10.35 | 2.00 | 4.29E-25 | 4.49E-24 |
| Galnt12 | 10.33 | 1.53 | 4.98E-25 | 5.21E-24 |
| Gins2 | 10.33 | 2.41 | 5.11E-25 | 5.34E-24 |
| Cmtm7 | 10.33 | 2.34 | 5.38E-25 | 5.61E-24 |
| B3galnt2 | 10.32 | 1.70 | 5.48E-25 | 5.71E-24 |
| Cdkal1 | 10.32 | 1.68 | 5.82E-25 | 6.06E-24 |
| Prim1 | 10.32 | 2.18 | 5.91E-25 | 6.15E-24 |
| Mtap | 10.31 | 2.27 | 6.41E-25 | 6.66E-24 |
| Rad9a | 10.30 | 2.28 | 6.72E-25 | 6.98E-24 |
| Trmt11 | 10.30 | 2.12 | 6.77E-25 | 7.03E-24 |
| Dhx37 | 10.30 | 2.17 | 6.91E-25 | 7.17E-24 |
| Mtif2 | 10.30 | 1.64 | 7.12E-25 | 7.38E-24 |
| Cep192 | 10.30 | 2.01 | 7.19E-25 | 7.45E-24 |
| Asf1b | 10.29 | 2.64 | 7.45E-25 | 7.71E-24 |
| Bub1b | 10.28 | 2.43 | 8.64E-25 | 8.91E-24 |
| Zfp395 | 10.28 | 1.70 | 8.77E-25 | 9.04E-24 |
| Dna2 | 10.27 | 2.73 | 9.72E-25 | 1E-23 |
| Cntrob | 10.27 | 2.43 | 9.98E-25 | 1.03E-23 |
| Dtx4 | 10.26 | 2.05 | 1.03E-24 | 1.06E-23 |
| Aunip | 10.26 | 2.95 | 1.1E-24 | 1.13E-23 |
| Mre11a | 10.25 | 2.01 | 1.2E-24 | 1.23E-23 |
| Pitx2 | 10.24 | 1.60 | 1.38E-24 | 1.41E-23 |
| Zfp507 | 10.23 | 1.90 | 1.39E-24 | 1.42E-23 |
| Ide | 10.23 | 1.52 | 1.42E-24 | 1.45E-23 |
| Naa25 | 10.22 | 1.77 | 1.62E-24 | 1.65E-23 |
| Ddx18 | 10.22 | 2.17 | 1.65E-24 | 1.68E-23 |
| Alkbh1 | 10.21 | 1.98 | 1.74E-24 | 1.77E-23 |
| Alg14 | 10.20 | 1.72 | 2E-24 | 2.02E-23 |
| Vars2 | 10.20 | 2.06 | 2E-24 | 2.02E-23 |
| Utp6 | 10.19 | 1.67 | 2.2E-24 | 2.22E-23 |
| Akr1b3 | 10.19 | 1.53 | 2.22E-24 | 2.23E-23 |
| 0Wsu102e | 10.19 | 2.01 | 2.25E-24 | 2.27E-23 |
| Nars2 | 10.18 | 1.84 | 2.36E-24 | 2.37E-23 |
| Rars2 | 10.18 | 1.59 | 2.46E-24 | 2.47E-23 |
| Bcap29 | 10.18 | 2.45 | 2.47E-24 | 2.48E-23 |
| G2e3 | 10.18 | 1.88 | 2.55E-24 | 2.56E-23 |
| Ttc27 | 10.15 | 2.59 | 3.19E-24 | 3.19E-23 |
| Fbxo5 | 10.15 | 2.23 | 3.24E-24 | 3.23E-23 |
| Lsg1 | 10.14 | 1.75 | 3.66E-24 | 3.65E-23 |
| Txlna | 10.14 | 1.58 | 3.79E-24 | 3.78E-23 |
| Ecsit | 10.13 | 1.83 | 4.16E-24 | 4.13E-23 |
| Kat14 | 10.13 | 1.66 | 4.17E-24 | 4.14E-23 |
| Nfia | 10.11 | 1.56 | 4.95E-24 | 4.91E-23 |

| | | | | |
|---|---|---|---|---|
| Heatr1 | 10.11 | 2.39 | 5.09E-24 | 5.04E-23 |
| Ftsj3 | 10.10 | 2.05 | 5.3E-24 | 5.25E-23 |
| Cox10 | 10.10 | 2.14 | 5.63E-24 | 5.56E-23 |
| Kif18a | 10.10 | 2.62 | 5.67E-24 | 5.6E-23 |
| Mreg | 10.09 | 1.68 | 6.2E-24 | 6.11E-23 |
| AW209491 | 10.09 | 2.39 | 6.36E-24 | 6.27E-23 |
| Swap70 | 10.08 | 1.70 | 6.53E-24 | 6.43E-23 |
| Lsm11 | 10.08 | 2.71 | 6.87E-24 | 6.76E-23 |
| Cd177 | 10.08 | 2.38 | 6.99E-24 | 6.87E-23 |
| Tsr2 | 10.07 | 2.39 | 7.27E-24 | 7.15E-23 |
| Atg16l1 | 10.07 | 1.70 | 7.52E-24 | 7.38E-23 |
| Prmt1 | 10.06 | 2.28 | 8.05E-24 | 7.89E-23 |
| Cdc7 | 10.05 | 2.46 | 9.25E-24 | 9.06E-23 |
| Shq1 | 10.03 | 2.48 | 1.08E-23 | 1.05E-22 |
| Cdc42ep1 | 10.03 | 1.75 | 1.17E-23 | 1.14E-22 |
| Polq | 10.02 | 3.38 | 1.2E-23 | 1.17E-22 |
| Pdcd2 | 10.02 | 1.67 | 1.27E-23 | 1.24E-22 |
| B3gntl1 | 10.02 | 2.54 | 1.27E-23 | 1.24E-22 |
| Ndufaf2 | 10.02 | 1.80 | 1.3E-23 | 1.26E-22 |
| Nup88 | 10.01 | 1.73 | 1.4E-23 | 1.36E-22 |
| Cenpa | 10.01 | 2.84 | 1.4E-23 | 1.36E-22 |
| Taf2 | 10.01 | 1.74 | 1.44E-23 | 1.39E-22 |
| Clpp | 10.01 | 1.60 | 1.45E-23 | 1.4E-22 |
| Aasdhppt | 10.00 | 1.72 | 1.46E-23 | 1.41E-22 |
| Mdc1 | 10.00 | 2.02 | 1.52E-23 | 1.47E-22 |
| Als2cl | 10.00 | 2.44 | 1.54E-23 | 1.49E-22 |
| Ruvbl1 | 9.99 | 2.29 | 1.71E-23 | 1.65E-22 |
| Pex11b | 9.98 | 2.20 | 1.87E-23 | 1.8E-22 |
| Zfp593 | 9.98 | 2.26 | 1.87E-23 | 1.8E-22 |
| Tedc2 | 9.97 | 2.49 | 2.09E-23 | 2E-22 |
| Mto1 | 9.97 | 1.98 | 2.12E-23 | 2.04E-22 |
| Nmral1 | 9.96 | 2.71 | 2.23E-23 | 2.14E-22 |
| Wdhd1 | 9.95 | 2.07 | 2.43E-23 | 2.33E-22 |
| Tlr1 | 9.94 | 2.46 | 2.75E-23 | 2.63E-22 |
| Pfdn4 | 9.94 | 1.86 | 2.79E-23 | 2.67E-22 |
| Bola1 | 9.93 | 2.11 | 2.96E-23 | 2.83E-22 |
| Nr2c2ap | 9.93 | 1.85 | 2.96E-23 | 2.83E-22 |
| Ube2t | 9.93 | 2.68 | 3.23E-23 | 3.08E-22 |
| Fam57a | 9.92 | 2.15 | 3.5E-23 | 3.33E-22 |
| Yars2 | 9.91 | 2.22 | 3.79E-23 | 3.6E-22 |
| Eif2s1 | 9.91 | 1.62 | 3.9E-23 | 3.7E-22 |
| Actr8 | 9.90 | 1.70 | 4.1E-23 | 3.88E-22 |
| Pank1 | 9.90 | 1.81 | 4.13E-23 | 3.91E-22 |
| Nup133 | 9.90 | 2.21 | 4.34E-23 | 4.1E-22 |

| | | | | |
|---|---|---|---|---|
| Pdss2 | 9.89 | 1.81 | 4.38E-23 | 4.14E-22 |
| Utp15 | 9.89 | 1.92 | 4.41E-23 | 4.16E-22 |
| Ncald | 9.89 | 2.15 | 4.55E-23 | 4.29E-22 |
| Rbm28 | 9.89 | 1.99 | 4.72E-23 | 4.45E-22 |
| Emc1 | 9.89 | 1.70 | 4.82E-23 | 4.54E-22 |
| Fam171a1 | 9.88 | 1.84 | 5.02E-23 | 4.72E-22 |
| Tedc1 | 9.86 | 2.01 | 6.05E-23 | 5.67E-22 |
| Cuedc2 | 9.86 | 1.80 | 6.05E-23 | 5.67E-22 |
| Glod4 | 9.85 | 1.58 | 6.68E-23 | 6.25E-22 |
| Dym | 9.85 | 1.51 | 7E-23 | 6.54E-22 |
| Aatf | 9.84 | 2.12 | 7.28E-23 | 6.8E-22 |
| Rpap1 | 9.84 | 2.06 | 7.4E-23 | 6.9E-22 |
| Mycl | 9.84 | 1.86 | 7.57E-23 | 7.06E-22 |
| Tango6 | 9.83 | 1.94 | 8.34E-23 | 7.76E-22 |
| Prr12 | 9.83 | 1.67 | 8.7E-23 | 8.09E-22 |
| Wdr55 | 9.82 | 2.15 | 9.06E-23 | 8.41E-22 |
| Depdc1b | 9.82 | 2.83 | 9.65E-23 | 8.95E-22 |
| Slc12a8 | 9.81 | 1.66 | 9.81E-23 | 9.08E-22 |
| Hnrnpr | 9.81 | 1.65 | 9.89E-23 | 9.15E-22 |
| Mpzl2 | 9.81 | 1.54 | 9.95E-23 | 9.2E-22 |
| Lancl1 | 9.81 | 2.20 | 1.01E-22 | 9.29E-22 |
| Cep83 | 9.81 | 1.82 | 1.04E-22 | 9.57E-22 |
| Cenpp | 9.79 | 3.15 | 1.31E-22 | 1.2E-21 |
| Zfp367 | 9.78 | 1.84 | 1.4E-22 | 1.29E-21 |
| Zan | 9.77 | 2.72 | 1.49E-22 | 1.37E-21 |
| Tnfaip8l1 | 9.76 | 2.20 | 1.68E-22 | 1.54E-21 |
| Rnf145 | 9.75 | 1.67 | 1.8E-22 | 1.65E-21 |
| Msh3 | 9.75 | 1.81 | 1.81E-22 | 1.66E-21 |
| Steap2 | 9.75 | 2.29 | 1.81E-22 | 1.66E-21 |
| Aarsd1 | 9.74 | 1.68 | 1.94E-22 | 1.78E-21 |
| Wdr3 | 9.74 | 2.34 | 2.02E-22 | 1.84E-21 |
| Haus8 | 9.74 | 2.09 | 2.02E-22 | 1.84E-21 |
| Wdr75 | 9.74 | 2.85 | 2.03E-22 | 1.85E-21 |
| Thada | 9.73 | 1.93 | 2.15E-22 | 1.96E-21 |
| Mgme1 | 9.73 | 2.17 | 2.26E-22 | 2.06E-21 |
| Gjb2 | 9.73 | 1.75 | 2.28E-22 | 2.07E-21 |
| Rcc2 | 9.72 | 2.15 | 2.4E-22 | 2.18E-21 |
| Rbm38 | 9.72 | 1.59 | 2.46E-22 | 2.24E-21 |
| Emd | 9.72 | 1.57 | 2.47E-22 | 2.24E-21 |
| Nfix | 9.72 | 1.51 | 2.47E-22 | 2.24E-21 |
| Rhno1 | 9.72 | 1.82 | 2.58E-22 | 2.34E-21 |
| Zbtb41 | 9.71 | 1.75 | 2.71E-22 | 2.46E-21 |
| Glrx3 | 9.70 | 1.80 | 3.04E-22 | 2.75E-21 |
| Zbed3 | 9.68 | 2.34 | 3.69E-22 | 3.33E-21 |

| | | | | |
|---|---|---|---|---|
| Exosc3 | 9.68 | 1.70 | 3.81E-22 | 3.43E-21 |
| Chek1 | 9.67 | 2.56 | 3.91E-22 | 3.53E-21 |
| Impa2 | 9.66 | 2.33 | 4.5E-22 | 4.04E-21 |
| Ccnf | 9.66 | 2.73 | 4.5E-22 | 4.04E-21 |
| Shprh | 9.66 | 1.84 | 4.54E-22 | 4.08E-21 |
| Thg1l | 9.65 | 2.30 | 4.75E-22 | 4.26E-21 |
| Exo1 | 9.64 | 2.87 | 5.2E-22 | 4.66E-21 |
| St7 | 9.63 | 2.39 | 6.25E-22 | 5.59E-21 |
| Arid5b | 9.61 | 1.60 | 7.54E-22 | 6.7E-21 |
| Aaas | 9.61 | 2.11 | 7.6E-22 | 6.75E-21 |
| Atp23 | 9.60 | 2.42 | 8.04E-22 | 7.14E-21 |
| Kpnb1 | 9.60 | 1.70 | 8.25E-22 | 7.32E-21 |
| Scarb2 | 9.59 | 1.53 | 8.64E-22 | 7.66E-21 |
| Ctdspl2 | 9.59 | 1.56 | 8.76E-22 | 7.75E-21 |
| Ugt2b35 | 9.59 | 1.91 | 8.78E-22 | 7.77E-21 |
| Cbx1 | 9.59 | 1.52 | 8.82E-22 | 7.8E-21 |
| Kif2c | 9.59 | 2.80 | 9.08E-22 | 8.02E-21 |
| Aven | 9.59 | 1.88 | 9.2E-22 | 8.13E-21 |
| Mlh1 | 9.58 | 2.04 | 9.44E-22 | 8.33E-21 |
| Ube2s | 9.58 | 1.50 | 9.76E-22 | 8.6E-21 |
| Pms1 | 9.58 | 2.14 | 9.89E-22 | 8.71E-21 |
| Rcl1 | 9.58 | 1.84 | 1.01E-21 | 8.93E-21 |
| Nol10 | 9.57 | 2.57 | 1.07E-21 | 9.44E-21 |
| Ska1 | 9.56 | 2.61 | 1.23E-21 | 1.08E-20 |
| Nelfa | 9.55 | 1.80 | 1.25E-21 | 1.09E-20 |
| Cox18 | 9.55 | 2.53 | 1.28E-21 | 1.12E-20 |
| Rpusd2 | 9.54 | 2.59 | 1.39E-21 | 1.22E-20 |
| Lmnb2 | 9.54 | 1.60 | 1.43E-21 | 1.25E-20 |
| Gars | 9.53 | 1.90 | 1.64E-21 | 1.43E-20 |
| Polr1a | 9.51 | 2.11 | 1.89E-21 | 1.64E-20 |
| Znrd2 | 9.51 | 1.75 | 1.98E-21 | 1.72E-20 |
| Atp2c2 | 9.51 | 1.88 | 1.99E-21 | 1.73E-20 |
| Fam185a | 9.49 | 2.32 | 2.33E-21 | 2.01E-20 |
| Rabepk | 9.49 | 2.34 | 2.34E-21 | 2.02E-20 |
| Nuf2 | 9.49 | 2.30 | 2.35E-21 | 2.03E-20 |
| Rfx5 | 9.49 | 2.05 | 2.38E-21 | 2.05E-20 |
| Acss2 | 9.47 | 1.53 | 2.79E-21 | 2.4E-20 |
| Nup107 | 9.46 | 2.49 | 2.96E-21 | 2.54E-20 |
| Tsnax | 9.46 | 1.70 | 2.97E-21 | 2.55E-20 |
| Kti12 | 9.46 | 1.78 | 3.01E-21 | 2.58E-20 |
| D2hgdh | 9.46 | 1.58 | 3.12E-21 | 2.68E-20 |
| Lrrk1 | 9.46 | 1.56 | 3.13E-21 | 2.68E-20 |
| Las1l | 9.45 | 1.94 | 3.28E-21 | 2.81E-20 |
| Fermt1 | 9.45 | 1.68 | 3.41E-21 | 2.92E-20 |

| | | | | |
|---|---|---|---|---|
| Dhx33 | 9.44 | 1.97 | 3.67E-21 | 3.14E-20 |
| Brip1 | 9.44 | 1.96 | 3.8E-21 | 3.25E-20 |
| Cd44 | 9.44 | 1.61 | 3.84E-21 | 3.28E-20 |
| Tube1 | 9.43 | 3.06 | 4.16E-21 | 3.55E-20 |
| Ap3s2 | 9.42 | 1.56 | 4.3E-21 | 3.67E-20 |
| Frrs1 | 9.41 | 1.71 | 4.91E-21 | 4.17E-20 |
| Thoc6 | 9.41 | 2.32 | 5.01E-21 | 4.26E-20 |
| Nme6 | 9.41 | 2.30 | 5.19E-21 | 4.4E-20 |
| Ache | 9.40 | 2.35 | 5.25E-21 | 4.46E-20 |
| Aste1 | 9.39 | 1.62 | 5.8E-21 | 4.91E-20 |
| Cdc45 | 9.39 | 2.63 | 6.16E-21 | 5.22E-20 |
| Emc8 | 9.39 | 1.90 | 6.2E-21 | 5.25E-20 |
| Mcrip2 | 9.37 | 1.91 | 7.34E-21 | 6.2E-20 |
| Paip2b | 9.36 | 1.55 | 7.77E-21 | 6.55E-20 |
| Lrp8 | 9.36 | 3.89 | 7.79E-21 | 6.57E-20 |
| Mettl23 | 9.36 | 1.90 | 7.94E-21 | 6.69E-20 |
| Bub1 | 9.35 | 2.52 | 8.52E-21 | 7.18E-20 |
| Fpgs | 9.35 | 1.81 | 8.73E-21 | 7.35E-20 |
| Elp1 | 9.35 | 2.05 | 8.98E-21 | 7.55E-20 |
| Kif24 | 9.35 | 2.56 | 9.15E-21 | 7.69E-20 |
| Ckap4 | 9.34 | 1.84 | 9.29E-21 | 7.8E-20 |
| Zwint | 9.34 | 1.67 | 9.64E-21 | 8.08E-20 |
| Cks1b | 9.33 | 2.60 | 1.02E-20 | 8.55E-20 |
| Pfkl | 9.33 | 1.62 | 1.04E-20 | 8.68E-20 |
| Acsm3 | 9.33 | 1.96 | 1.08E-20 | 9.02E-20 |
| Tmtc4 | 9.32 | 1.69 | 1.16E-20 | 9.67E-20 |
| Tmem9 | 9.31 | 1.83 | 1.25E-20 | 1.04E-19 |
| Sclt1 | 9.31 | 2.16 | 1.32E-20 | 1.1E-19 |
| Dmd | 9.31 | 1.64 | 1.34E-20 | 1.11E-19 |
| Nom1 | 9.30 | 1.76 | 1.43E-20 | 1.19E-19 |
| Mccc2 | 9.30 | 1.78 | 1.45E-20 | 1.2E-19 |
| Hgh1 | 9.30 | 2.13 | 1.47E-20 | 1.21E-19 |
| Armc10 | 9.29 | 1.85 | 1.49E-20 | 1.24E-19 |
| Ankrd49 | 9.28 | 1.93 | 1.64E-20 | 1.36E-19 |
| Hpse | 9.28 | 1.58 | 1.68E-20 | 1.39E-19 |
| Maip1 | 9.28 | 1.70 | 1.7E-20 | 1.4E-19 |
| Pole2 | 9.28 | 2.45 | 1.74E-20 | 1.44E-19 |
| Igf1r | 9.27 | 2.04 | 1.8E-20 | 1.48E-19 |
| Wdr76 | 9.27 | 2.76 | 1.85E-20 | 1.52E-19 |
| Ddx51 | 9.27 | 2.51 | 1.91E-20 | 1.57E-19 |
| Ska3 | 9.26 | 2.78 | 2.07E-20 | 1.7E-19 |
| Nanp | 9.25 | 1.85 | 2.22E-20 | 1.81E-19 |
| Clspn | 9.24 | 2.19 | 2.41E-20 | 1.97E-19 |
| Rpf2 | 9.23 | 2.55 | 2.6E-20 | 2.12E-19 |

| | | | | |
|---|---|---|---|---|
| Chtf18 | 9.23 | 2.92 | 2.68E-20 | 2.19E-19 |
| Cdk5rap2 | 9.23 | 2.33 | 2.69E-20 | 2.2E-19 |
| Nln | 9.23 | 1.73 | 2.7E-20 | 2.2E-19 |
| Elp3 | 9.23 | 1.66 | 2.73E-20 | 2.23E-19 |
| Ldah | 9.23 | 1.60 | 2.75E-20 | 2.24E-19 |
| Eny2 | 9.22 | 2.25 | 2.88E-20 | 2.34E-19 |
| Notch1 | 9.22 | 1.85 | 3.07E-20 | 2.5E-19 |
| Fancd2 | 9.21 | 3.03 | 3.14E-20 | 2.56E-19 |
| Mettl1 | 9.21 | 1.99 | 3.34E-20 | 2.71E-19 |
| Zfp369 | 9.21 | 1.98 | 3.37E-20 | 2.74E-19 |
| Ttc37 | 9.20 | 1.73 | 3.42E-20 | 2.78E-19 |
| Rmnd1 | 9.20 | 1.80 | 3.55E-20 | 2.88E-19 |
| Atxn3 | 9.19 | 1.67 | 3.75E-20 | 3.04E-19 |
| Dapk2 | 9.19 | 1.56 | 3.92E-20 | 3.18E-19 |
| Hspbp1 | 9.19 | 2.08 | 3.92E-20 | 3.18E-19 |
| Mms22l | 9.19 | 1.86 | 3.98E-20 | 3.23E-19 |
| Sdr39u1 | 9.18 | 1.67 | 4.21E-20 | 3.4E-19 |
| Ccdc18 | 9.18 | 3.29 | 4.49E-20 | 3.63E-19 |
| Ccdc59 | 9.17 | 1.53 | 4.74E-20 | 3.82E-19 |
| Esrp2 | 9.16 | 1.77 | 5.09E-20 | 4.1E-19 |
| Toe1 | 9.16 | 2.03 | 5.09E-20 | 4.1E-19 |
| Hpf1 | 9.16 | 2.16 | 5.1E-20 | 4.11E-19 |
| Ing5 | 9.16 | 1.76 | 5.34E-20 | 4.3E-19 |
| Polg | 9.16 | 1.66 | 5.44E-20 | 4.37E-19 |
| Rprd1a | 9.15 | 1.78 | 5.8E-20 | 4.66E-19 |
| Cep128 | 9.15 | 2.35 | 5.82E-20 | 4.68E-19 |
| Slc19a1 | 9.15 | 1.58 | 5.87E-20 | 4.71E-19 |
| Mtg1 | 9.14 | 2.33 | 6.2E-20 | 4.97E-19 |
| Sumf2 | 9.13 | 2.70 | 6.63E-20 | 5.3E-19 |
| Mphosph10 | 9.12 | 2.13 | 7.41E-20 | 5.92E-19 |
| Ints11 | 9.12 | 2.04 | 7.82E-20 | 6.24E-19 |
| Nudcd1 | 9.11 | 1.88 | 8.05E-20 | 6.42E-19 |
| Fam53a | 9.11 | 1.91 | 8.19E-20 | 6.53E-19 |
| Ints4 | 9.11 | 1.57 | 8.24E-20 | 6.56E-19 |
| Shank2 | 9.11 | 2.14 | 8.31E-20 | 6.62E-19 |
| Nqo1 | 9.10 | 1.74 | 9.09E-20 | 7.22E-19 |
| Gtf2e2 | 9.10 | 1.51 | 9.28E-20 | 7.37E-19 |
| Nup160 | 9.10 | 2.00 | 9.28E-20 | 7.37E-19 |
| Ipo4 | 9.09 | 1.94 | 9.83E-20 | 7.8E-19 |
| Eef1akmt1 | 9.08 | 1.77 | 1.11E-19 | 8.79E-19 |
| Cdc16 | 9.08 | 1.87 | 1.12E-19 | 8.83E-19 |
| Xrcc1 | 9.07 | 1.85 | 1.15E-19 | 9.07E-19 |
| Ten1 | 9.07 | 1.51 | 1.15E-19 | 9.09E-19 |
| Gemin5 | 9.07 | 2.47 | 1.17E-19 | 9.25E-19 |

| | | | | |
|---|---|---|---|---|
| Larp7 | 9.06 | 1.69 | 1.25E-19 | 9.83E-19 |
| Elp2 | 9.06 | 1.88 | 1.26E-19 | 9.91E-19 |
| Poglut1 | 9.06 | 2.21 | 1.27E-19 | 1E-18 |
| Bcs1l | 9.06 | 2.31 | 1.28E-19 | 1.01E-18 |
| Epn3 | 9.06 | 2.98 | 1.3E-19 | 1.02E-18 |
| Noa1 | 9.06 | 2.31 | 1.35E-19 | 1.06E-18 |
| Traip | 9.06 | 3.39 | 1.35E-19 | 1.06E-18 |
| Utp18 | 9.04 | 1.84 | 1.54E-19 | 1.2E-18 |
| Ovol2 | 9.04 | 1.60 | 1.58E-19 | 1.23E-18 |
| Sumo2 | 9.04 | 1.67 | 1.59E-19 | 1.24E-18 |
| Hr | 9.03 | 1.60 | 1.65E-19 | 1.29E-18 |
| Setdb1 | 9.03 | 1.67 | 1.66E-19 | 1.29E-18 |
| Syk | 9.01 | 1.68 | 1.97E-19 | 1.53E-18 |
| Tmem107 | 9.01 | 2.74 | 1.98E-19 | 1.54E-18 |
| Pcca | 9.00 | 1.65 | 2.25E-19 | 1.74E-18 |
| Haus3 | 9.00 | 1.64 | 2.26E-19 | 1.76E-18 |
| Synj2 | 8.97 | 1.61 | 2.97E-19 | 2.3E-18 |
| Ppt2 | 8.97 | 1.75 | 3.02E-19 | 2.33E-18 |
| Amz2 | 8.97 | 1.56 | 3.1E-19 | 2.39E-18 |
| Pkmyt1 | 8.96 | 1.96 | 3.2E-19 | 2.47E-18 |
| Eif2b3 | 8.96 | 2.23 | 3.27E-19 | 2.53E-18 |
| Alkbh8 | 8.96 | 1.93 | 3.29E-19 | 2.54E-18 |
| Cenpk | 8.95 | 2.56 | 3.44E-19 | 2.65E-18 |
| Creb3l4 | 8.95 | 1.77 | 3.48E-19 | 2.68E-18 |
| BC030867 | 8.93 | 2.85 | 4.09E-19 | 3.14E-18 |
| Exosc1 | 8.92 | 1.99 | 4.72E-19 | 3.61E-18 |
| Apmap | 8.91 | 1.54 | 4.9E-19 | 3.74E-18 |
| Clns1a | 8.91 | 1.58 | 4.92E-19 | 3.76E-18 |
| Zdhhc2 | 8.90 | 2.09 | 5.51E-19 | 4.2E-18 |
| Pradc1 | 8.90 | 2.48 | 5.53E-19 | 4.21E-18 |
| Cep89 | 8.89 | 1.86 | 5.98E-19 | 4.54E-18 |
| Zc3h7b | 8.89 | 1.56 | 5.99E-19 | 4.55E-18 |
| Mettl17 | 8.88 | 1.75 | 6.97E-19 | 5.28E-18 |
| Arhgap39 | 8.87 | 1.59 | 7.01E-19 | 5.31E-18 |
| Sept10 | 8.87 | 1.86 | 7.4E-19 | 5.6E-18 |
| Pdrg1 | 8.86 | 1.61 | 8.07E-19 | 6.1E-18 |
| Gfm2 | 8.86 | 1.54 | 8.09E-19 | 6.1E-18 |
| Macrod1 | 8.86 | 1.67 | 8.17E-19 | 6.16E-18 |
| Mastl | 8.86 | 2.94 | 8.31E-19 | 6.26E-18 |
| Tmem69 | 8.85 | 2.07 | 8.65E-19 | 6.52E-18 |
| Mthfsd | 8.85 | 1.70 | 9.04E-19 | 6.8E-18 |
| Taf1a | 8.85 | 1.81 | 9.13E-19 | 6.86E-18 |
| Adat1 | 8.84 | 2.62 | 9.6E-19 | 7.2E-18 |
| Ercc6l | 8.84 | 3.02 | 9.63E-19 | 7.23E-18 |

| | | | | |
|---|---|---|---|---|
| Cdan1 | 8.83 | 1.99 | 1.05E-18 | 7.88E-18 |
| Mmachc | 8.82 | 2.44 | 1.12E-18 | 8.39E-18 |
| Ndufaf7 | 8.82 | 1.97 | 1.15E-18 | 8.58E-18 |
| Impdh1 | 8.82 | 1.83 | 1.18E-18 | 8.81E-18 |
| Cdca4 | 8.82 | 1.60 | 1.19E-18 | 8.86E-18 |
| Ankrd13b | 8.82 | 1.85 | 1.19E-18 | 8.91E-18 |
| Slc35b4 | 8.81 | 1.67 | 1.23E-18 | 9.16E-18 |
| Qrsl1 | 8.80 | 2.34 | 1.34E-18 | 9.95E-18 |
| Ercc2 | 8.79 | 2.32 | 1.48E-18 | 1.1E-17 |
| Cep295 | 8.79 | 2.17 | 1.54E-18 | 1.14E-17 |
| Parpbp | 8.78 | 2.61 | 1.67E-18 | 1.24E-17 |
| Exosc2 | 8.78 | 2.19 | 1.69E-18 | 1.25E-17 |
| Mtfr2 | 8.77 | 2.85 | 1.75E-18 | 1.3E-17 |
| Sult1c2 | 8.77 | 3.10 | 1.79E-18 | 1.32E-17 |
| Nbeal2 | 8.77 | 1.74 | 1.85E-18 | 1.37E-17 |
| Snx27 | 8.76 | 1.53 | 1.89E-18 | 1.39E-17 |
| Daglb | 8.76 | 1.55 | 1.97E-18 | 1.46E-17 |
| Tmem186 | 8.76 | 1.53 | 2E-18 | 1.47E-17 |
| Hpdl | 8.76 | 3.60 | 2.01E-18 | 1.49E-17 |
| Faim | 8.75 | 1.67 | 2.05E-18 | 1.51E-17 |
| Rwdd4a | 8.75 | 1.56 | 2.16E-18 | 1.59E-17 |
| Pdcd11 | 8.74 | 1.71 | 2.27E-18 | 1.67E-17 |
| Lyrm9 | 8.74 | 1.60 | 2.37E-18 | 1.74E-17 |
| Ckap2l | 8.72 | 2.67 | 2.77E-18 | 2.03E-17 |
| Timm22 | 8.72 | 1.57 | 2.79E-18 | 2.04E-17 |
| Trmt61a | 8.72 | 2.06 | 2.82E-18 | 2.07E-17 |
| Sgo2a | 8.72 | 2.62 | 2.84E-18 | 2.08E-17 |
| Tbc1d31 | 8.71 | 2.36 | 2.96E-18 | 2.17E-17 |
| Ptcd2 | 8.71 | 1.79 | 3.06E-18 | 2.23E-17 |
| Timeless | 8.70 | 2.40 | 3.2E-18 | 2.33E-17 |
| Orc1 | 8.70 | 3.24 | 3.28E-18 | 2.39E-17 |
| Gnl2 | 8.70 | 1.54 | 3.29E-18 | 2.4E-17 |
| Gsdmc3 | 8.70 | 2.23 | 3.32E-18 | 2.42E-17 |
| Fgfr1op | 8.70 | 2.08 | 3.34E-18 | 2.43E-17 |
| Ctu2 | 8.70 | 2.08 | 3.36E-18 | 2.44E-17 |
| Espl1 | 8.69 | 2.40 | 3.51E-18 | 2.55E-17 |
| Bud13 | 8.69 | 1.53 | 3.51E-18 | 2.55E-17 |
| Yars | 8.69 | 1.53 | 3.58E-18 | 2.6E-17 |
| Mxd3 | 8.69 | 2.59 | 3.72E-18 | 2.7E-17 |
| Sfmbt1 | 8.69 | 1.79 | 3.75E-18 | 2.72E-17 |
| Rttn | 8.69 | 2.27 | 3.77E-18 | 2.73E-17 |
| Slc25a35 | 8.68 | 1.77 | 3.85E-18 | 2.79E-17 |
| Trmt6 | 8.68 | 1.71 | 3.98E-18 | 2.88E-17 |
| Fbxo21 | 8.68 | 2.16 | 3.99E-18 | 2.88E-17 |

| | | | | |
|---|---|---|---|---|
| Scap | 8.68 | 1.53 | 4.04E-18 | 2.92E-17 |
| Cwc22 | 8.68 | 1.98 | 4.05E-18 | 2.93E-17 |
| Thoc1 | 8.67 | 1.69 | 4.2E-18 | 3.03E-17 |
| Riok2 | 8.67 | 1.66 | 4.23E-18 | 3.05E-17 |
| Zbtb8a | 8.67 | 1.69 | 4.25E-18 | 3.06E-17 |
| Usp24 | 8.65 | 1.61 | 4.97E-18 | 3.57E-17 |
| Grhpr | 8.64 | 1.66 | 5.62E-18 | 4.03E-17 |
| Eif4a3 | 8.63 | 1.54 | 6.18E-18 | 4.42E-17 |
| Zzz3 | 8.63 | 1.52 | 6.25E-18 | 4.47E-17 |
| Taco1 | 8.63 | 1.65 | 6.37E-18 | 4.55E-17 |
| Clcc1 | 8.62 | 1.67 | 6.64E-18 | 4.74E-17 |
| Fntb | 8.61 | 1.67 | 7.03E-18 | 5.01E-17 |
| Usp46 | 8.61 | 1.87 | 7.17E-18 | 5.11E-17 |
| Stau2 | 8.61 | 1.54 | 7.23E-18 | 5.15E-17 |
| Hist1h3g | 8.61 | 2.88 | 7.38E-18 | 5.25E-17 |
| Card11 | 8.60 | 2.07 | 7.7E-18 | 5.47E-17 |
| Ddx31 | 8.60 | 2.56 | 8.24E-18 | 5.85E-17 |
| Rpp38 | 8.59 | 2.43 | 8.62E-18 | 6.11E-17 |
| Nop14 | 8.59 | 2.10 | 8.67E-18 | 6.14E-17 |
| Tdp1 | 8.59 | 2.22 | 8.7E-18 | 6.16E-17 |
| Ndc80 | 8.59 | 2.85 | 9.06E-18 | 6.41E-17 |
| Sinhcaf | 8.58 | 1.87 | 9.41E-18 | 6.65E-17 |
| Thumpd3 | 8.58 | 1.62 | 9.42E-18 | 6.66E-17 |
| Troap | 8.58 | 2.65 | 9.47E-18 | 6.69E-17 |
| Eef2kmt | 8.57 | 2.02 | 1E-17 | 7.07E-17 |
| Ears2 | 8.57 | 1.98 | 1.01E-17 | 7.15E-17 |
| Emc6 | 8.57 | 1.56 | 1.04E-17 | 7.3E-17 |
| Cyb5r1 | 8.57 | 1.73 | 1.04E-17 | 7.35E-17 |
| Pdcd2l | 8.57 | 1.61 | 1.05E-17 | 7.36E-17 |
| Capn13 | 8.56 | 1.68 | 1.09E-17 | 7.69E-17 |
| Tex10 | 8.56 | 1.67 | 1.11E-17 | 7.79E-17 |
| Elp4 | 8.55 | 1.53 | 1.27E-17 | 8.93E-17 |
| Snapc4 | 8.54 | 1.84 | 1.32E-17 | 9.26E-17 |
| Tbl2 | 8.54 | 1.72 | 1.35E-17 | 9.45E-17 |
| Poglut2 | 8.54 | 2.06 | 1.4E-17 | 9.78E-17 |
| Itgb3bp | 8.54 | 2.07 | 1.4E-17 | 9.8E-17 |
| Bard1 | 8.53 | 2.49 | 1.42E-17 | 9.91E-17 |
| Cenpm | 8.53 | 2.47 | 1.43E-17 | 9.97E-17 |
| Hist1h3a | 8.53 | 3.06 | 1.46E-17 | 1.02E-16 |
| Med16 | 8.53 | 1.57 | 1.47E-17 | 1.02E-16 |
| Mettl16 | 8.53 | 2.16 | 1.49E-17 | 1.04E-16 |
| Cep57 | 8.52 | 1.76 | 1.59E-17 | 1.11E-16 |
| Poldip2 | 8.52 | 1.57 | 1.6E-17 | 1.11E-16 |
| Psrc1 | 8.52 | 2.41 | 1.62E-17 | 1.13E-16 |

| | | | | |
|---|---|---|---|---|
| G3bp1 | 8.51 | 1.82 | 1.69E-17 | 1.17E-16 |
| Zfp397 | 8.51 | 1.61 | 1.81E-17 | 1.26E-16 |
| Hars2 | 8.50 | 1.65 | 1.84E-17 | 1.28E-16 |
| Cstf2 | 8.48 | 1.64 | 2.32E-17 | 1.61E-16 |
| Nip7 | 8.47 | 1.89 | 2.37E-17 | 1.64E-16 |
| Leo1 | 8.47 | 1.57 | 2.47E-17 | 1.71E-16 |
| Ube2g2 | 8.47 | 1.61 | 2.52E-17 | 1.74E-16 |
| Lsm10 | 8.46 | 1.96 | 2.64E-17 | 1.82E-16 |
| Farsa | 8.46 | 1.64 | 2.64E-17 | 1.82E-16 |
| Fktn | 8.46 | 1.95 | 2.69E-17 | 1.86E-16 |
| Tada2a | 8.45 | 1.88 | 3.02E-17 | 2.08E-16 |
| Abitram | 8.44 | 2.50 | 3.16E-17 | 2.17E-16 |
| Cenph | 8.42 | 2.70 | 3.85E-17 | 2.64E-16 |
| Gtf2h3 | 8.42 | 1.52 | 3.88E-17 | 2.66E-16 |
| Fsbp | 8.41 | 2.63 | 3.94E-17 | 2.71E-16 |
| Zfp462 | 8.41 | 1.57 | 3.95E-17 | 2.71E-16 |
| Hibch | 8.41 | 1.90 | 4.01E-17 | 2.75E-16 |
| B4galnt4 | 8.41 | 1.59 | 4.05E-17 | 2.78E-16 |
| Hspa4 | 8.40 | 1.53 | 4.38E-17 | 3E-16 |
| Siva1 | 8.40 | 2.27 | 4.46E-17 | 3.05E-16 |
| Recql | 8.40 | 1.85 | 4.52E-17 | 3.1E-16 |
| Smim4 | 8.38 | 1.58 | 5.48E-17 | 3.73E-16 |
| Gcat | 8.37 | 1.71 | 5.61E-17 | 3.82E-16 |
| Dph5 | 8.36 | 2.09 | 6.21E-17 | 4.21E-16 |
| Exd2 | 8.35 | 1.67 | 6.62E-17 | 4.48E-16 |
| Cdkn2c | 8.34 | 1.63 | 7.61E-17 | 5.14E-16 |
| Amigo1 | 8.34 | 1.73 | 7.65E-17 | 5.16E-16 |
| Utp23 | 8.34 | 2.13 | 7.7E-17 | 5.19E-16 |
| Pwp2 | 8.33 | 2.11 | 7.77E-17 | 5.24E-16 |
| Pfas | 8.33 | 1.74 | 7.79E-17 | 5.25E-16 |
| Hgfac | 8.33 | 1.52 | 8.12E-17 | 5.46E-16 |
| Xrcc5 | 8.33 | 2.15 | 8.3E-17 | 5.58E-16 |
| Rxylt1 | 8.33 | 1.68 | 8.42E-17 | 5.66E-16 |
| Fanci | 8.32 | 2.86 | 8.86E-17 | 5.94E-16 |
| Pank4 | 8.31 | 1.59 | 9.27E-17 | 6.21E-16 |
| Klhl5 | 8.31 | 1.78 | 9.73E-17 | 6.52E-16 |
| Lyplal1 | 8.30 | 1.79 | 1.03E-16 | 6.9E-16 |
| Shisa2 | 8.30 | 1.57 | 1.07E-16 | 7.12E-16 |
| Pde2a | 8.29 | 1.59 | 1.14E-16 | 7.63E-16 |
| Tox | 8.29 | 1.63 | 1.17E-16 | 7.79E-16 |
| Nap1l1 | 8.28 | 1.81 | 1.22E-16 | 8.11E-16 |
| Coasy | 8.28 | 1.62 | 1.24E-16 | 8.23E-16 |
| Prpf4 | 8.27 | 1.71 | 1.3E-16 | 8.59E-16 |
| Dcakd | 8.27 | 1.73 | 1.34E-16 | 8.88E-16 |

| | | | | |
|---|---|---|---|---|
| Vma21 | 8.27 | 1.72 | 1.34E-16 | 8.89E-16 |
| Fignl1 | 8.27 | 2.71 | 1.35E-16 | 8.97E-16 |
| Zfp472 | 8.26 | 1.91 | 1.41E-16 | 9.34E-16 |
| B3galt6 | 8.25 | 2.50 | 1.62E-16 | 1.07E-15 |
| Spef1 | 8.25 | 1.77 | 1.62E-16 | 1.07E-15 |
| Nol11 | 8.25 | 2.04 | 1.63E-16 | 1.07E-15 |
| Orai2 | 8.24 | 1.82 | 1.73E-16 | 1.14E-15 |
| Zfp414 | 8.24 | 1.68 | 1.76E-16 | 1.16E-15 |
| Zkscan8 | 8.24 | 1.87 | 1.79E-16 | 1.17E-15 |
| Haus6 | 8.23 | 1.78 | 1.81E-16 | 1.19E-15 |
| Cenpi | 8.23 | 2.60 | 1.9E-16 | 1.25E-15 |
| Cep72 | 8.22 | 2.03 | 2E-16 | 1.31E-15 |
| E2f2 | 8.22 | 1.89 | 2.11E-16 | 1.38E-15 |
| Uhrf1bp1 | 8.21 | 2.33 | 2.14E-16 | 1.4E-15 |
| Paics | 8.21 | 2.30 | 2.17E-16 | 1.41E-15 |
| Phf6 | 8.21 | 1.70 | 2.19E-16 | 1.43E-15 |
| Ctps | 8.21 | 2.21 | 2.2E-16 | 1.43E-15 |
| Xylb | 8.21 | 2.21 | 2.21E-16 | 1.44E-15 |
| Jmjd7 | 8.21 | 2.23 | 2.23E-16 | 1.45E-15 |
| Sarnp | 8.20 | 1.56 | 2.41E-16 | 1.57E-15 |
| Sned1 | 8.20 | 2.01 | 2.45E-16 | 1.59E-15 |
| Smim8 | 8.19 | 1.87 | 2.59E-16 | 1.68E-15 |
| Tmem39b | 8.19 | 1.77 | 2.65E-16 | 1.71E-15 |
| Arhgef10l | 8.19 | 1.85 | 2.68E-16 | 1.74E-15 |
| Rapgef5 | 8.19 | 1.76 | 2.72E-16 | 1.76E-15 |
| Nedd1 | 8.18 | 1.82 | 2.74E-16 | 1.77E-15 |
| Sipa1l2 | 8.18 | 1.53 | 2.93E-16 | 1.89E-15 |
| Polr3e | 8.17 | 1.53 | 3.21E-16 | 2.07E-15 |
| Slc39a8 | 8.16 | 1.72 | 3.28E-16 | 2.11E-15 |
| Depdc5 | 8.16 | 1.74 | 3.43E-16 | 2.2E-15 |
| Nsun5 | 8.14 | 1.99 | 4.09E-16 | 2.62E-15 |
| Atr | 8.13 | 1.98 | 4.14E-16 | 2.65E-15 |
| Zkscan17 | 8.13 | 2.06 | 4.34E-16 | 2.78E-15 |
| Polr1b | 8.12 | 2.02 | 4.53E-16 | 2.89E-15 |
| Cep63 | 8.12 | 1.58 | 4.58E-16 | 2.93E-15 |
| Cep70 | 8.11 | 1.75 | 4.93E-16 | 3.15E-15 |
| Dtd1 | 8.11 | 2.16 | 5.07E-16 | 3.23E-15 |
| Wtip | 8.10 | 1.91 | 5.43E-16 | 3.46E-15 |
| Slc20a2 | 8.09 | 1.73 | 5.87E-16 | 3.73E-15 |
| Hmg20a | 8.09 | 1.65 | 5.93E-16 | 3.77E-15 |
| Fdx2 | 8.09 | 1.73 | 5.97E-16 | 3.79E-15 |
| Gins3 | 8.09 | 2.89 | 6.06E-16 | 3.84E-15 |
| Tspan12 | 8.09 | 1.62 | 6.13E-16 | 3.89E-15 |
| Cep135 | 8.09 | 1.89 | 6.15E-16 | 3.9E-15 |

| | | | | |
|---|---|---|---|---|
| Ccnd3 | 8.08 | 1.53 | 6.58E-16 | 4.16E-15 |
| Nol8 | 8.07 | 1.88 | 6.84E-16 | 4.32E-15 |
| Noc4l | 8.07 | 1.88 | 6.85E-16 | 4.32E-15 |
| Ndufaf1 | 8.07 | 1.74 | 7.25E-16 | 4.57E-15 |
| Wnk4 | 8.07 | 1.81 | 7.25E-16 | 4.57E-15 |
| Mettl4 | 8.07 | 2.03 | 7.25E-16 | 4.57E-15 |
| Eri3 | 8.06 | 1.52 | 7.72E-16 | 4.85E-15 |
| Myg1 | 8.06 | 1.64 | 7.92E-16 | 4.98E-15 |
| Zfp64 | 8.06 | 1.61 | 7.93E-16 | 4.98E-15 |
| Fam174b | 8.05 | 1.63 | 8.31E-16 | 5.22E-15 |
| Sdhaf3 | 8.05 | 2.76 | 8.36E-16 | 5.25E-15 |
| Smim26 | 8.05 | 1.56 | 8.47E-16 | 5.32E-15 |
| Pbdc1 | 8.04 | 1.80 | 8.84E-16 | 5.54E-15 |
| Eef1aknmt | 8.04 | 2.66 | 8.92E-16 | 5.59E-15 |
| Otud6b | 8.04 | 1.78 | 9.31E-16 | 5.83E-15 |
| Tbl3 | 8.03 | 1.59 | 9.38E-16 | 5.87E-15 |
| Pgap2 | 8.03 | 1.63 | 9.55E-16 | 5.97E-15 |
| Znrf3 | 8.03 | 1.52 | 9.59E-16 | 5.99E-15 |
| Ccdc167 | 8.02 | 2.35 | 1.03E-15 | 6.41E-15 |
| Saal1 | 8.02 | 2.38 | 1.07E-15 | 6.65E-15 |
| Stard5 | 8.02 | 1.97 | 1.08E-15 | 6.75E-15 |
| Tarbp1 | 8.02 | 2.34 | 1.09E-15 | 6.76E-15 |
| Aplf | 8.01 | 2.32 | 1.11E-15 | 6.92E-15 |
| Fastkd5 | 8.01 | 1.91 | 1.12E-15 | 6.99E-15 |
| Ccl11 | 8.01 | 2.25 | 1.14E-15 | 7.09E-15 |
| Fancl | 8.01 | 2.22 | 1.16E-15 | 7.2E-15 |
| Med18 | 8.00 | 1.86 | 1.21E-15 | 7.54E-15 |
| Cnnm3 | 8.00 | 1.72 | 1.22E-15 | 7.57E-15 |
| Gipc1 | 8.00 | 1.65 | 1.23E-15 | 7.66E-15 |
| Wrn | 8.00 | 1.53 | 1.26E-15 | 7.81E-15 |
| Dbt | 7.99 | 1.66 | 1.32E-15 | 8.18E-15 |
| Fam53c | 7.99 | 1.54 | 1.34E-15 | 8.32E-15 |
| Nxt1 | 7.98 | 1.89 | 1.51E-15 | 9.3E-15 |
| Tmem199 | 7.97 | 1.77 | 1.54E-15 | 9.51E-15 |
| Vegfb | 7.97 | 1.51 | 1.54E-15 | 9.51E-15 |
| Efna4 | 7.97 | 1.77 | 1.57E-15 | 9.67E-15 |
| Rtn4rl1 | 7.96 | 1.95 | 1.69E-15 | 1.04E-14 |
| Poc1a | 7.96 | 1.92 | 1.79E-15 | 1.1E-14 |
| Pop5 | 7.96 | 1.63 | 1.79E-15 | 1.1E-14 |
| Cenpq | 7.95 | 2.38 | 1.87E-15 | 1.15E-14 |
| Kif4 | 7.95 | 2.21 | 1.87E-15 | 1.15E-14 |
| Pomgnt1 | 7.95 | 1.72 | 1.89E-15 | 1.16E-14 |
| Abce1 | 7.95 | 1.80 | 1.91E-15 | 1.17E-14 |
| Chaf1b | 7.95 | 1.95 | 1.93E-15 | 1.18E-14 |

| | | | | |
|---|---|---|---|---|
| Pole4 | 7.94 | 1.75 | 1.95E-15 | 1.19E-14 |
| Mtrr | 7.94 | 2.68 | 1.95E-15 | 1.19E-14 |
| Unk | 7.94 | 1.56 | 1.97E-15 | 1.21E-14 |
| Brca2 | 7.94 | 2.60 | 1.98E-15 | 1.21E-14 |
| Depdc1a | 7.93 | 2.47 | 2.25E-15 | 1.37E-14 |
| Mars2 | 7.93 | 2.18 | 2.27E-15 | 1.39E-14 |
| Trip13 | 7.91 | 2.30 | 2.53E-15 | 1.54E-14 |
| Rtel1 | 7.91 | 2.22 | 2.6E-15 | 1.58E-14 |
| Phf20 | 7.90 | 2.11 | 2.7E-15 | 1.64E-14 |
| Nufip1 | 7.90 | 1.83 | 2.72E-15 | 1.65E-14 |
| Nek1 | 7.90 | 2.03 | 2.74E-15 | 1.66E-14 |
| Orc2 | 7.90 | 1.90 | 2.75E-15 | 1.67E-14 |
| Trnau1ap | 7.89 | 1.55 | 3.01E-15 | 1.82E-14 |
| Polr2d | 7.88 | 1.76 | 3.31E-15 | 2E-14 |
| R3hcc1l | 7.87 | 1.60 | 3.49E-15 | 2.11E-14 |
| Agmat | 7.86 | 2.13 | 3.91E-15 | 2.35E-14 |
| Nsmce3 | 7.86 | 1.58 | 3.92E-15 | 2.35E-14 |
| Dus4l | 7.85 | 2.78 | 4.11E-15 | 2.46E-14 |
| Rac3 | 7.85 | 2.69 | 4.2E-15 | 2.51E-14 |
| Pnpt1 | 7.84 | 1.72 | 4.36E-15 | 2.61E-14 |
| Odf2l | 7.84 | 1.66 | 4.62E-15 | 2.76E-14 |
| Dtl | 7.84 | 2.47 | 4.63E-15 | 2.77E-14 |
| Taf6l | 7.83 | 2.41 | 4.72E-15 | 2.81E-14 |
| Wdr4 | 7.83 | 1.77 | 4.85E-15 | 2.89E-14 |
| Fance | 7.83 | 1.73 | 5.04E-15 | 3E-14 |
| Nt5dc1 | 7.82 | 1.66 | 5.16E-15 | 3.07E-14 |
| Knop1 | 7.82 | 1.98 | 5.36E-15 | 3.19E-14 |
| Atpsckmt | 7.82 | 2.05 | 5.45E-15 | 3.24E-14 |
| Fam110a | 7.82 | 2.19 | 5.48E-15 | 3.25E-14 |
| Dnph1 | 7.81 | 2.32 | 5.73E-15 | 3.39E-14 |
| E2f7 | 7.81 | 2.15 | 5.93E-15 | 3.51E-14 |
| Ift88 | 7.80 | 2.15 | 5.96E-15 | 3.53E-14 |
| Mlh3 | 7.80 | 2.51 | 5.97E-15 | 3.53E-14 |
| Hmga2 | 7.80 | 2.71 | 5.99E-15 | 3.54E-14 |
| Rnf8 | 7.80 | 1.61 | 6.02E-15 | 3.56E-14 |
| Nt5m | 7.80 | 1.65 | 6.12E-15 | 3.62E-14 |
| Nek4 | 7.80 | 2.07 | 6.2E-15 | 3.66E-14 |
| Fdxacb1 | 7.80 | 1.91 | 6.43E-15 | 3.79E-14 |
| Crtap | 7.80 | 1.80 | 6.44E-15 | 3.79E-14 |
| Rad17 | 7.78 | 1.65 | 6.98E-15 | 4.11E-14 |
| Dhps | 7.78 | 1.64 | 6.99E-15 | 4.11E-14 |
| Mterf4 | 7.78 | 1.93 | 7.15E-15 | 4.2E-14 |
| Tssc4 | 7.78 | 1.52 | 7.4E-15 | 4.34E-14 |
| Arv1 | 7.78 | 2.19 | 7.52E-15 | 4.41E-14 |

| | | | | |
|---|---|---|---|---|
| Galnt3 | 7.77 | 1.64 | 7.72E-15 | 4.52E-14 |
| Msto1 | 7.75 | 1.83 | 9.27E-15 | 5.41E-14 |
| Alyref | 7.74 | 2.11 | 1.01E-14 | 5.89E-14 |
| Zmym3 | 7.74 | 1.80 | 1.03E-14 | 6E-14 |
| Det1 | 7.72 | 1.83 | 1.12E-14 | 6.52E-14 |
| Isy1 | 7.72 | 1.57 | 1.13E-14 | 6.55E-14 |
| Naf1 | 7.72 | 2.46 | 1.15E-14 | 6.66E-14 |
| Sars2 | 7.72 | 1.99 | 1.18E-14 | 6.85E-14 |
| Aopep | 7.72 | 1.74 | 1.18E-14 | 6.87E-14 |
| Mphosph9 | 7.71 | 2.35 | 1.25E-14 | 7.24E-14 |
| Enkd1 | 7.71 | 2.71 | 1.25E-14 | 7.28E-14 |
| Ift27 | 7.71 | 2.27 | 1.26E-14 | 7.3E-14 |
| Dtwd1 | 7.71 | 2.24 | 1.28E-14 | 7.4E-14 |
| Tctn3 | 7.71 | 2.15 | 1.28E-14 | 7.44E-14 |
| Wdr73 | 7.70 | 2.28 | 1.31E-14 | 7.6E-14 |
| Slc16a6 | 7.70 | 2.02 | 1.32E-14 | 7.62E-14 |
| Pitx1 | 7.70 | 2.22 | 1.34E-14 | 7.78E-14 |
| Cep68 | 7.70 | 2.19 | 1.35E-14 | 7.83E-14 |
| Bcl7a | 7.69 | 1.98 | 1.53E-14 | 8.8E-14 |
| Ice2 | 7.66 | 2.06 | 1.8E-14 | 1.03E-13 |
| Fam199x | 7.66 | 1.76 | 1.83E-14 | 1.05E-13 |
| Nagpa | 7.66 | 1.66 | 1.88E-14 | 1.08E-13 |
| Zfp101 | 7.66 | 1.97 | 1.89E-14 | 1.08E-13 |
| Kitl | 7.65 | 1.61 | 1.97E-14 | 1.13E-13 |
| Gtpbp3 | 7.65 | 1.67 | 1.98E-14 | 1.13E-13 |
| Pno1 | 7.65 | 1.55 | 2.03E-14 | 1.17E-13 |
| Qsox2 | 7.65 | 1.69 | 2.03E-14 | 1.17E-13 |
| Pigb | 7.65 | 2.04 | 2.04E-14 | 1.17E-13 |
| Ncf2 | 7.64 | 1.66 | 2.09E-14 | 1.2E-13 |
| Donson | 7.64 | 1.56 | 2.21E-14 | 1.26E-13 |
| Stimate | 7.62 | 1.81 | 2.47E-14 | 1.41E-13 |
| Eme1 | 7.62 | 3.00 | 2.53E-14 | 1.44E-13 |
| Smo | 7.62 | 2.05 | 2.56E-14 | 1.46E-13 |
| Mtbp | 7.62 | 2.76 | 2.6E-14 | 1.48E-13 |
| Dsn1 | 7.61 | 2.07 | 2.74E-14 | 1.56E-13 |
| Nit2 | 7.61 | 1.63 | 2.79E-14 | 1.59E-13 |
| Kntc1 | 7.60 | 2.29 | 2.98E-14 | 1.69E-13 |
| Prss8 | 7.58 | 1.61 | 3.44E-14 | 1.95E-13 |
| Ubiad1 | 7.58 | 1.87 | 3.58E-14 | 2.03E-13 |
| Ccdc115 | 7.57 | 1.98 | 3.62E-14 | 2.04E-13 |
| Yeats2 | 7.57 | 1.64 | 3.84E-14 | 2.16E-13 |
| Aqp4 | 7.56 | 1.70 | 4.12E-14 | 2.32E-13 |
| Lmf1 | 7.55 | 1.84 | 4.35E-14 | 2.45E-13 |
| Ndufaf5 | 7.55 | 1.68 | 4.36E-14 | 2.46E-13 |

| | | | | |
|---|---|---|---|---|
| Rbm15b | 7.55 | 1.57 | 4.43E-14 | 2.49E-13 |
| Gtf3c4 | 7.55 | 1.64 | 4.47E-14 | 2.52E-13 |
| Ndrg2 | 7.54 | 2.04 | 4.64E-14 | 2.61E-13 |
| Top1mt | 7.54 | 2.27 | 4.72E-14 | 2.65E-13 |
| Ppargc1b | 7.53 | 1.68 | 5.02E-14 | 2.82E-13 |
| Zfp800 | 7.53 | 1.53 | 5.19E-14 | 2.91E-13 |
| Fam72a | 7.53 | 3.04 | 5.22E-14 | 2.92E-13 |
| Ptpn4 | 7.52 | 1.72 | 5.42E-14 | 3.03E-13 |
| Ttf2 | 7.52 | 2.11 | 5.67E-14 | 3.17E-13 |
| Dmac1 | 7.51 | 1.59 | 5.74E-14 | 3.21E-13 |
| Jmjd4 | 7.51 | 2.00 | 5.88E-14 | 3.28E-13 |
| Skp2 | 7.51 | 1.89 | 6.07E-14 | 3.39E-13 |
| Plod3 | 7.51 | 1.57 | 6.1E-14 | 3.41E-13 |
| Lpin1 | 7.51 | 1.87 | 6.1E-14 | 3.41E-13 |
| Zmym1 | 7.50 | 2.43 | 6.45E-14 | 3.59E-13 |
| Tac1 | 7.50 | 2.10 | 6.52E-14 | 3.63E-13 |
| Zfp213 | 7.50 | 1.84 | 6.53E-14 | 3.64E-13 |
| Actr6 | 7.49 | 2.26 | 6.76E-14 | 3.76E-13 |
| Sox4 | 7.48 | 1.63 | 7.42E-14 | 4.13E-13 |
| Slc25a30 | 7.47 | 2.57 | 8.1E-14 | 4.49E-13 |
| Pex10 | 7.47 | 1.96 | 8.13E-14 | 4.51E-13 |
| Abraxas1 | 7.43 | 1.96 | 1.08E-13 | 5.92E-13 |
| Bub3 | 7.43 | 1.51 | 1.12E-13 | 6.13E-13 |
| Pqlc1 | 7.42 | 1.73 | 1.15E-13 | 6.31E-13 |
| Pkn3 | 7.42 | 2.06 | 1.17E-13 | 6.41E-13 |
| Cactin | 7.42 | 1.69 | 1.17E-13 | 6.44E-13 |
| Ints2 | 7.42 | 1.79 | 1.19E-13 | 6.53E-13 |
| Gkap1 | 7.42 | 1.75 | 1.19E-13 | 6.54E-13 |
| Ftsj1 | 7.41 | 1.97 | 1.31E-13 | 7.15E-13 |
| Nup43 | 7.40 | 2.63 | 1.31E-13 | 7.18E-13 |
| Taf4b | 7.40 | 2.37 | 1.35E-13 | 7.38E-13 |
| Kbtbd8 | 7.40 | 2.17 | 1.35E-13 | 7.39E-13 |
| Cenpo | 7.40 | 1.82 | 1.39E-13 | 7.57E-13 |
| Nme2 | 7.40 | 1.54 | 1.39E-13 | 7.6E-13 |
| Slc29a1 | 7.40 | 1.71 | 1.4E-13 | 7.64E-13 |
| Tut1 | 7.39 | 1.54 | 1.45E-13 | 7.9E-13 |
| Zc3h8 | 7.39 | 2.35 | 1.45E-13 | 7.92E-13 |
| Ccdc61 | 7.39 | 1.63 | 1.46E-13 | 7.94E-13 |
| Nfrkb | 7.39 | 1.61 | 1.47E-13 | 7.98E-13 |
| Usp31 | 7.39 | 2.27 | 1.52E-13 | 8.29E-13 |
| Nme4 | 7.38 | 2.27 | 1.53E-13 | 8.32E-13 |
| Acpp | 7.38 | 1.63 | 1.55E-13 | 8.44E-13 |
| Gpld1 | 7.38 | 1.98 | 1.61E-13 | 8.75E-13 |
| Nup54 | 7.38 | 1.84 | 1.61E-13 | 8.76E-13 |

| | | | | |
|---|---|---|---|---|
| Ppp2r1b | 7.37 | 1.74 | 1.74E-13 | 9.4E-13 |
| Pycr2 | 7.35 | 1.50 | 2.02E-13 | 1.09E-12 |
| Lnpk | 7.35 | 1.61 | 2.02E-13 | 1.09E-12 |
| Uspl1 | 7.34 | 1.61 | 2.09E-13 | 1.13E-12 |
| Sik2 | 7.33 | 1.53 | 2.24E-13 | 1.21E-12 |
| Haus5 | 7.33 | 2.41 | 2.27E-13 | 1.22E-12 |
| Sgsh | 7.32 | 1.76 | 2.41E-13 | 1.29E-12 |
| Hdac6 | 7.32 | 1.86 | 2.52E-13 | 1.35E-12 |
| Atad5 | 7.31 | 1.79 | 2.57E-13 | 1.38E-12 |
| Nav2 | 7.31 | 1.66 | 2.58E-13 | 1.38E-12 |
| Rtn4ip1 | 7.31 | 1.65 | 2.59E-13 | 1.39E-12 |
| Pctp | 7.31 | 1.72 | 2.64E-13 | 1.41E-12 |
| Nsl1 | 7.31 | 2.52 | 2.65E-13 | 1.42E-12 |
| Eri2 | 7.31 | 2.42 | 2.71E-13 | 1.45E-12 |
| Slc25a32 | 7.30 | 1.99 | 2.89E-13 | 1.54E-12 |
| Jag1 | 7.30 | 1.75 | 2.95E-13 | 1.57E-12 |
| Gtf3c3 | 7.30 | 1.54 | 2.97E-13 | 1.58E-12 |
| Fam83d | 7.29 | 2.36 | 3.03E-13 | 1.61E-12 |
| Nav1 | 7.29 | 1.59 | 3.06E-13 | 1.63E-12 |
| Wee1 | 7.28 | 1.63 | 3.24E-13 | 1.72E-12 |
| Tmem171 | 7.28 | 1.56 | 3.25E-13 | 1.72E-12 |
| Igsf8 | 7.28 | 1.67 | 3.26E-13 | 1.73E-12 |
| Rrs1 | 7.28 | 2.33 | 3.32E-13 | 1.76E-12 |
| Pstk | 7.28 | 2.28 | 3.46E-13 | 1.83E-12 |
| Klf12 | 7.27 | 1.88 | 3.53E-13 | 1.87E-12 |
| Ccdc130 | 7.27 | 2.22 | 3.56E-13 | 1.88E-12 |
| Cep152 | 7.26 | 2.06 | 3.86E-13 | 2.03E-12 |
| Rhbdd2 | 7.25 | 1.67 | 4.08E-13 | 2.15E-12 |
| Map3k14 | 7.25 | 1.65 | 4.13E-13 | 2.17E-12 |
| Slc17a9 | 7.25 | 1.90 | 4.19E-13 | 2.2E-12 |
| Fbxo45 | 7.24 | 1.86 | 4.34E-13 | 2.28E-12 |
| Abcb8 | 7.24 | 1.50 | 4.5E-13 | 2.37E-12 |
| Zswim3 | 7.22 | 1.68 | 5.01E-13 | 2.63E-12 |
| Rai1 | 7.22 | 1.65 | 5.07E-13 | 2.66E-12 |
| Cdkn3 | 7.22 | 2.41 | 5.11E-13 | 2.68E-12 |
| Miga1 | 7.22 | 1.52 | 5.13E-13 | 2.69E-12 |
| Tfb2m | 7.22 | 1.60 | 5.24E-13 | 2.74E-12 |
| Fancc | 7.21 | 1.93 | 5.39E-13 | 2.82E-12 |
| Tti1 | 7.21 | 1.66 | 5.42E-13 | 2.83E-12 |
| Gorab | 7.21 | 1.87 | 5.54E-13 | 2.89E-12 |
| Entpd6 | 7.21 | 1.59 | 5.68E-13 | 2.97E-12 |
| Adamts15 | 7.21 | 2.14 | 5.72E-13 | 2.98E-12 |
| Mrm3 | 7.20 | 2.83 | 5.98E-13 | 3.11E-12 |
| Zscan22 | 7.20 | 2.77 | 6.03E-13 | 3.14E-12 |

| | | | | |
|---|---|---|---|---|
| Katnbl1 | 7.20 | 1.62 | 6.16E-13 | 3.21E-12 |
| Ccnh | 7.19 | 1.70 | 6.56E-13 | 3.41E-12 |
| Rexo5 | 7.19 | 1.69 | 6.61E-13 | 3.43E-12 |
| Sh3rf2 | 7.18 | 1.68 | 6.79E-13 | 3.52E-12 |
| Nol12 | 7.18 | 1.81 | 6.97E-13 | 3.61E-12 |
| Lonrf3 | 7.18 | 1.75 | 7.09E-13 | 3.68E-12 |
| Lins1 | 7.18 | 2.09 | 7.12E-13 | 3.69E-12 |
| Rcan3 | 7.18 | 1.59 | 7.2E-13 | 3.73E-12 |
| Scmh1 | 7.18 | 1.57 | 7.21E-13 | 3.74E-12 |
| Slc28a3 | 7.17 | 2.06 | 7.25E-13 | 3.75E-12 |
| Clybl | 7.17 | 1.62 | 7.37E-13 | 3.82E-12 |
| Focad | 7.17 | 2.16 | 7.44E-13 | 3.85E-12 |
| Dhodh | 7.17 | 2.35 | 7.48E-13 | 3.87E-12 |
| Zdhhc15 | 7.17 | 2.36 | 7.66E-13 | 3.96E-12 |
| Notch2 | 7.17 | 1.87 | 7.74E-13 | 4E-12 |
| Fam122b | 7.16 | 2.22 | 7.93E-13 | 4.1E-12 |
| Helq | 7.16 | 1.51 | 8.01E-13 | 4.14E-12 |
| Smn1 | 7.16 | 2.17 | 8.02E-13 | 4.14E-12 |
| Ssh1 | 7.16 | 1.64 | 8.1E-13 | 4.18E-12 |
| Mthfd1l | 7.14 | 1.64 | 9.08E-13 | 4.67E-12 |
| Thap1 | 7.13 | 1.72 | 1.01E-12 | 5.18E-12 |
| Pgghg | 7.12 | 1.56 | 1.04E-12 | 5.35E-12 |
| Rpain | 7.11 | 1.65 | 1.14E-12 | 5.84E-12 |
| Rida | 7.10 | 1.90 | 1.21E-12 | 6.18E-12 |
| Siah1b | 7.08 | 1.95 | 1.39E-12 | 7.12E-12 |
| Tubgcp2 | 7.08 | 1.69 | 1.4E-12 | 7.15E-12 |
| Ddx55 | 7.08 | 1.55 | 1.48E-12 | 7.57E-12 |
| Yae1d1 | 7.08 | 2.01 | 1.49E-12 | 7.6E-12 |
| Arhgap8 | 7.06 | 2.21 | 1.68E-12 | 8.54E-12 |
| Surf6 | 7.05 | 1.89 | 1.73E-12 | 8.81E-12 |
| Mettl14 | 7.05 | 1.57 | 1.76E-12 | 8.95E-12 |
| Ccne1 | 7.05 | 2.28 | 1.77E-12 | 8.97E-12 |
| Abhd10 | 7.05 | 1.65 | 1.79E-12 | 9.06E-12 |
| Sdhaf1 | 7.05 | 1.63 | 1.85E-12 | 9.4E-12 |
| Cdk5rap1 | 7.04 | 1.56 | 1.89E-12 | 9.57E-12 |
| Ighmbp2 | 7.04 | 1.63 | 1.94E-12 | 9.82E-12 |
| Dock11 | 7.01 | 1.61 | 2.38E-12 | 1.2E-11 |
| Pex26 | 7.01 | 1.54 | 2.39E-12 | 1.21E-11 |
| Kyat3 | 6.99 | 1.95 | 2.72E-12 | 1.37E-11 |
| Cttnbp2nl | 6.98 | 1.59 | 3E-12 | 1.51E-11 |
| Trmt10a | 6.98 | 2.48 | 3E-12 | 1.51E-11 |
| Gatc | 6.98 | 1.69 | 3.03E-12 | 1.52E-11 |
| Dcaf4 | 6.97 | 1.89 | 3.16E-12 | 1.58E-11 |
| Slc10a7 | 6.97 | 1.56 | 3.26E-12 | 1.63E-11 |

| | | | | |
|---|---|---|---|---|
| Cog8 | 6.96 | 1.56 | 3.32E-12 | 1.66E-11 |
| Gnb1l | 6.96 | 3.09 | 3.34E-12 | 1.67E-11 |
| Suv39h2 | 6.95 | 2.39 | 3.53E-12 | 1.76E-11 |
| Fam111a | 6.95 | 2.14 | 3.55E-12 | 1.78E-11 |
| Galk1 | 6.95 | 1.58 | 3.64E-12 | 1.82E-11 |
| Zfp961 | 6.95 | 2.02 | 3.7E-12 | 1.85E-11 |
| Esrrg | 6.95 | 1.95 | 3.74E-12 | 1.87E-11 |
| Tefm | 6.94 | 1.71 | 3.92E-12 | 1.95E-11 |
| Tfb1m | 6.94 | 2.23 | 3.95E-12 | 1.96E-11 |
| Cyp2w1 | 6.94 | 2.22 | 4E-12 | 1.99E-11 |
| Xrcc2 | 6.93 | 2.62 | 4.14E-12 | 2.05E-11 |
| E2f1 | 6.93 | 2.16 | 4.15E-12 | 2.06E-11 |
| Shroom2 | 6.93 | 1.57 | 4.34E-12 | 2.15E-11 |
| Ggct | 6.92 | 1.77 | 4.59E-12 | 2.27E-11 |
| Bace2 | 6.91 | 1.73 | 4.72E-12 | 2.33E-11 |
| Klhdc4 | 6.91 | 1.72 | 4.75E-12 | 2.35E-11 |
| Twistnb | 6.91 | 1.63 | 5E-12 | 2.47E-11 |
| Prepl | 6.90 | 1.56 | 5.05E-12 | 2.5E-11 |
| Ccdc127 | 6.90 | 1.69 | 5.06E-12 | 2.5E-11 |
| Ccdc112 | 6.88 | 2.83 | 5.85E-12 | 2.88E-11 |
| Ift74 | 6.88 | 1.89 | 5.87E-12 | 2.89E-11 |
| Ercc6 | 6.88 | 1.52 | 5.97E-12 | 2.94E-11 |
| Eif2ak4 | 6.88 | 1.82 | 6.09E-12 | 3E-11 |
| Ubxn2b | 6.88 | 1.78 | 6.11E-12 | 3E-11 |
| Coil | 6.87 | 1.76 | 6.25E-12 | 3.07E-11 |
| Paqr5 | 6.85 | 2.09 | 7.4E-12 | 3.62E-11 |
| Gcfc2 | 6.85 | 1.99 | 7.6E-12 | 3.71E-11 |
| Apex2 | 6.84 | 1.77 | 8.05E-12 | 3.93E-11 |
| Hsf2 | 6.84 | 1.71 | 8.13E-12 | 3.97E-11 |
| Lmf2 | 6.83 | 1.59 | 8.22E-12 | 4E-11 |
| Slf1 | 6.83 | 1.73 | 8.27E-12 | 4.03E-11 |
| Pofut1 | 6.83 | 1.54 | 8.37E-12 | 4.08E-11 |
| Mios | 6.83 | 1.58 | 8.69E-12 | 4.23E-11 |
| Cenpu | 6.82 | 2.61 | 8.98E-12 | 4.37E-11 |
| Morn2 | 6.82 | 2.56 | 8.98E-12 | 4.37E-11 |
| Ap1s3 | 6.82 | 2.20 | 9.1E-12 | 4.42E-11 |
| Ovca2 | 6.81 | 1.78 | 9.98E-12 | 4.83E-11 |
| Fam168a | 6.81 | 1.55 | 9.99E-12 | 4.84E-11 |
| Ctc1 | 6.81 | 1.90 | 1E-11 | 4.86E-11 |
| Phtf1 | 6.80 | 1.65 | 1.05E-11 | 5.08E-11 |
| Map1s | 6.76 | 1.57 | 1.42E-11 | 6.81E-11 |
| Kcnk6 | 6.75 | 1.55 | 1.48E-11 | 7.09E-11 |
| Rmi2 | 6.75 | 2.48 | 1.52E-11 | 7.28E-11 |
| Lpar6 | 6.74 | 1.53 | 1.58E-11 | 7.58E-11 |

| | | | | |
|---|---|---|---|---|
| Slc12a9 | 6.74 | 1.70 | 1.63E-11 | 7.81E-11 |
| Ttll4 | 6.73 | 1.63 | 1.67E-11 | 8E-11 |
| Smyd3 | 6.73 | 1.61 | 1.68E-11 | 8.05E-11 |
| Trmt5 | 6.73 | 2.02 | 1.69E-11 | 8.07E-11 |
| Dimt1 | 6.72 | 2.39 | 1.77E-11 | 8.44E-11 |
| Cenpt | 6.72 | 1.63 | 1.79E-11 | 8.53E-11 |
| Lrrcc1 | 6.72 | 1.54 | 1.8E-11 | 8.58E-11 |
| Xndc1 | 6.72 | 2.01 | 1.88E-11 | 8.93E-11 |
| Sergef | 6.71 | 2.70 | 1.93E-11 | 9.18E-11 |
| Dscc1 | 6.71 | 2.59 | 1.93E-11 | 9.19E-11 |
| Pidd1 | 6.71 | 2.15 | 1.97E-11 | 9.37E-11 |
| Tsen15 | 6.71 | 2.10 | 1.98E-11 | 9.43E-11 |
| Ogfod2 | 6.70 | 1.50 | 2.02E-11 | 9.61E-11 |
| Mks1 | 6.70 | 2.34 | 2.05E-11 | 9.71E-11 |
| Gen1 | 6.70 | 2.33 | 2.05E-11 | 9.73E-11 |
| Ccdc137 | 6.70 | 2.23 | 2.1E-11 | 9.98E-11 |
| Cep97 | 6.70 | 2.23 | 2.11E-11 | 1E-10 |
| Bcl2l12 | 6.68 | 1.64 | 2.33E-11 | 1.1E-10 |
| Repin1 | 6.67 | 2.01 | 2.56E-11 | 1.21E-10 |
| Cenpn | 6.66 | 2.61 | 2.68E-11 | 1.26E-10 |
| Orc5 | 6.66 | 1.55 | 2.77E-11 | 1.3E-10 |
| Rdh13 | 6.65 | 1.64 | 2.85E-11 | 1.34E-10 |
| Smyd5 | 6.65 | 2.06 | 2.92E-11 | 1.37E-10 |
| Srpk2 | 6.65 | 1.67 | 2.95E-11 | 1.38E-10 |
| Ipo7 | 6.65 | 1.57 | 2.95E-11 | 1.39E-10 |
| Pla2g4a | 6.63 | 1.55 | 3.26E-11 | 1.53E-10 |
| Tsen2 | 6.62 | 1.94 | 3.47E-11 | 1.62E-10 |
| Ccdc191 | 6.62 | 1.96 | 3.48E-11 | 1.62E-10 |
| Twnk | 6.62 | 1.60 | 3.56E-11 | 1.66E-10 |
| Cby1 | 6.61 | 1.91 | 3.89E-11 | 1.81E-10 |
| Tchp | 6.61 | 1.62 | 3.97E-11 | 1.85E-10 |
| Ybey | 6.60 | 2.41 | 4E-11 | 1.86E-10 |
| Wdr34 | 6.60 | 2.38 | 4E-11 | 1.86E-10 |
| Adra2a | 6.60 | 2.14 | 4.11E-11 | 1.91E-10 |
| Etaa1 | 6.59 | 1.97 | 4.28E-11 | 1.98E-10 |
| Tmem216 | 6.58 | 2.24 | 4.64E-11 | 2.15E-10 |
| Polr1e | 6.58 | 2.20 | 4.68E-11 | 2.16E-10 |
| Mcat | 6.58 | 1.77 | 4.69E-11 | 2.17E-10 |
| Pinx1 | 6.58 | 2.24 | 4.72E-11 | 2.18E-10 |
| Ctu1 | 6.57 | 1.68 | 4.94E-11 | 2.28E-10 |
| Wdr92 | 6.57 | 1.82 | 5.19E-11 | 2.39E-10 |
| Rwdd3 | 6.56 | 2.91 | 5.27E-11 | 2.43E-10 |
| Cdk10 | 6.55 | 1.76 | 5.86E-11 | 2.69E-10 |
| Cd320 | 6.53 | 1.75 | 6.47E-11 | 2.96E-10 |

| | | | | |
|---|---|---|---|---|
| Tnfaip8 | 6.53 | 1.78 | 6.47E-11 | 2.97E-10 |
| Bysl | 6.53 | 1.97 | 6.72E-11 | 3.08E-10 |
| Slc9a5 | 6.51 | 2.47 | 7.56E-11 | 3.46E-10 |
| Morc4 | 6.51 | 2.45 | 7.6E-11 | 3.47E-10 |
| Uevld | 6.50 | 1.84 | 7.82E-11 | 3.57E-10 |
| Zfp329 | 6.50 | 1.75 | 7.86E-11 | 3.59E-10 |
| Gpn1 | 6.50 | 1.66 | 8.17E-11 | 3.72E-10 |
| Hddc2 | 6.49 | 2.06 | 8.55E-11 | 3.89E-10 |
| Adat2 | 6.49 | 2.02 | 8.61E-11 | 3.91E-10 |
| Sbsn | 6.49 | 2.79 | 8.81E-11 | 4E-10 |
| Heatr3 | 6.48 | 1.84 | 8.97E-11 | 4.07E-10 |
| Arhgap33 | 6.48 | 2.33 | 9.07E-11 | 4.11E-10 |
| H1fx | 6.48 | 1.72 | 9.43E-11 | 4.27E-10 |
| Vav1 | 6.47 | 2.02 | 9.57E-11 | 4.33E-10 |
| Mtg2 | 6.47 | 1.61 | 9.58E-11 | 4.34E-10 |
| Wdr12 | 6.47 | 2.31 | 1.01E-10 | 4.56E-10 |
| Matn2 | 6.46 | 2.12 | 1.02E-10 | 4.6E-10 |
| Clpb | 6.46 | 1.61 | 1.04E-10 | 4.68E-10 |
| Morn1 | 6.46 | 3.00 | 1.06E-10 | 4.79E-10 |
| Rad51c | 6.46 | 3.00 | 1.07E-10 | 4.81E-10 |
| Srbd1 | 6.45 | 1.94 | 1.1E-10 | 4.95E-10 |
| Emp2 | 6.45 | 1.85 | 1.12E-10 | 5.06E-10 |
| Nkapd1 | 6.45 | 1.59 | 1.14E-10 | 5.12E-10 |
| BC052040 | 6.43 | 1.79 | 1.25E-10 | 5.63E-10 |
| Csad | 6.39 | 1.63 | 1.62E-10 | 7.25E-10 |
| Ttc32 | 6.39 | 1.70 | 1.65E-10 | 7.36E-10 |
| Ttr | 6.39 | 2.56 | 1.67E-10 | 7.48E-10 |
| Npat | 6.39 | 1.69 | 1.71E-10 | 7.64E-10 |
| Wdr62 | 6.37 | 2.78 | 1.88E-10 | 8.35E-10 |
| Mex3d | 6.36 | 1.81 | 1.97E-10 | 8.75E-10 |
| Cers4 | 6.35 | 1.67 | 2.1E-10 | 9.34E-10 |
| Phf19 | 6.35 | 2.94 | 2.12E-10 | 9.42E-10 |
| Trp53rka | 6.35 | 2.10 | 2.12E-10 | 9.42E-10 |
| Ccnjl | 6.35 | 1.63 | 2.13E-10 | 9.47E-10 |
| Spryd4 | 6.35 | 1.75 | 2.15E-10 | 9.54E-10 |
| Slain1 | 6.31 | 1.62 | 2.82E-10 | 1.24E-09 |
| Nudt6 | 6.31 | 1.96 | 2.86E-10 | 1.26E-09 |
| Rpp30 | 6.27 | 1.67 | 3.57E-10 | 1.56E-09 |
| Cep78 | 6.27 | 1.64 | 3.6E-10 | 1.58E-09 |
| Mbd4 | 6.26 | 2.57 | 3.73E-10 | 1.64E-09 |
| Txndc16 | 6.26 | 1.56 | 3.78E-10 | 1.66E-09 |
| Mnd1 | 6.26 | 3.85 | 3.79E-10 | 1.66E-09 |
| Spice1 | 6.26 | 2.01 | 3.84E-10 | 1.68E-09 |
| Steap1 | 6.26 | 2.03 | 3.86E-10 | 1.69E-09 |

| | | | | |
|---|---|---|---|---|
| Llph | 6.26 | 1.99 | 3.87E-10 | 1.69E-09 |
| Ankrd26 | 6.26 | 1.88 | 3.95E-10 | 1.73E-09 |
| Dennd6b | 6.24 | 1.83 | 4.27E-10 | 1.86E-09 |
| Cep57l1 | 6.23 | 2.59 | 4.52E-10 | 1.97E-09 |
| Rnaseh1 | 6.23 | 1.76 | 4.53E-10 | 1.97E-09 |
| Cwc27 | 6.22 | 1.56 | 4.84E-10 | 2.1E-09 |
| Clasp2 | 6.22 | 1.59 | 4.87E-10 | 2.11E-09 |
| Taf3 | 6.22 | 1.57 | 4.89E-10 | 2.12E-09 |
| Strada | 6.22 | 1.97 | 4.91E-10 | 2.13E-09 |
| Zbtb12 | 6.22 | 1.91 | 4.95E-10 | 2.15E-09 |
| Prorp | 6.21 | 1.50 | 5.23E-10 | 2.27E-09 |
| Galnt5 | 6.21 | 2.05 | 5.28E-10 | 2.29E-09 |
| Glmn | 6.21 | 2.20 | 5.45E-10 | 2.36E-09 |
| Palb2 | 6.21 | 2.35 | 5.45E-10 | 2.36E-09 |
| Cgas | 6.20 | 2.31 | 5.49E-10 | 2.37E-09 |
| Pask | 6.20 | 2.28 | 5.58E-10 | 2.41E-09 |
| \W554918 | 6.20 | 1.56 | 5.62E-10 | 2.43E-09 |
| Tubgcp5 | 6.19 | 1.55 | 6.09E-10 | 2.63E-09 |
| Zmym6 | 6.17 | 1.52 | 6.67E-10 | 2.87E-09 |
| Ipo11 | 6.17 | 1.80 | 6.75E-10 | 2.9E-09 |
| Slc2a1 | 6.15 | 1.72 | 7.59E-10 | 3.26E-09 |
| Ccdc57 | 6.14 | 2.60 | 8.28E-10 | 3.54E-09 |
| Zfp40 | 6.14 | 2.42 | 8.41E-10 | 3.6E-09 |
| Map3k15 | 6.13 | 1.50 | 8.81E-10 | 3.76E-09 |
| Polr3d | 6.12 | 1.57 | 9.54E-10 | 4.07E-09 |
| Tgs1 | 6.12 | 1.57 | 9.61E-10 | 4.1E-09 |
| Fn3krp | 6.11 | 1.97 | 9.72E-10 | 4.14E-09 |
| Sirt4 | 6.09 | 1.56 | 1.14E-09 | 4.83E-09 |
| Eif3j1 | 6.08 | 2.32 | 1.17E-09 | 4.97E-09 |
| Mad1l1 | 6.08 | 1.85 | 1.21E-09 | 5.12E-09 |
| Cdpf1 | 6.07 | 1.59 | 1.25E-09 | 5.29E-09 |
| Sgsm1 | 6.07 | 1.53 | 1.26E-09 | 5.32E-09 |
| Abt1 | 6.07 | 1.74 | 1.27E-09 | 5.34E-09 |
| Dnaaf2 | 6.07 | 2.16 | 1.27E-09 | 5.35E-09 |
| Mettl2 | 6.07 | 1.69 | 1.27E-09 | 5.35E-09 |
| Rpp40 | 6.07 | 2.07 | 1.31E-09 | 5.51E-09 |
| Pif1 | 6.07 | 2.77 | 1.31E-09 | 5.53E-09 |
| Tmeff1 | 6.07 | 2.69 | 1.32E-09 | 5.55E-09 |
| Trim45 | 6.05 | 3.27 | 1.41E-09 | 5.91E-09 |
| ⁻mem185a | 6.05 | 1.62 | 1.48E-09 | 6.21E-09 |
| Dph2 | 6.03 | 2.03 | 1.66E-09 | 6.95E-09 |
| Kifc5b | 6.03 | 2.97 | 1.67E-09 | 6.99E-09 |
| Socs5 | 6.02 | 1.80 | 1.74E-09 | 7.28E-09 |
| Ccdc77 | 6.02 | 1.87 | 1.78E-09 | 7.44E-09 |

| | | | | |
|---|---|---|---|---|
| Qser1 | 6.01 | 1.76 | 1.84E-09 | 7.66E-09 |
| Lipt1 | 6.01 | 2.56 | 1.85E-09 | 7.7E-09 |
| Tulp3 | 6.01 | 2.48 | 1.86E-09 | 7.76E-09 |
| Fancf | 6.00 | 1.64 | 1.93E-09 | 8.04E-09 |
| Yif1b | 6.00 | 1.92 | 1.98E-09 | 8.22E-09 |
| Arl14ep | 6.00 | 1.86 | 1.99E-09 | 8.28E-09 |
| Acer3 | 5.99 | 1.72 | 2.07E-09 | 8.6E-09 |
| Nme7 | 5.99 | 1.69 | 2.08E-09 | 8.66E-09 |
| Bcl11a | 5.99 | 1.66 | 2.16E-09 | 8.96E-09 |
| Prkdc | 5.98 | 1.87 | 2.28E-09 | 9.46E-09 |
| Mtrf1 | 5.97 | 2.49 | 2.3E-09 | 9.53E-09 |

Zone 3: Significantly Zonated Genes

| Symbol | scores | LFC | pvals | pvals_adj |
|---|---|---|---|---|
| Krt19 | 58.52 | 3.52 | 0 | 0 |
| Lypd8 | 52.37 | 3.12 | 0 | 0 |
| Plac8 | 37.41 | 2.25 | 3.1E-306 | 2E-302 |
| Reg3b | 36.00 | 2.24 | 8.8E-284 | 4.2E-280 |
| Fabp6 | 34.71 | 2.27 | 6E-264 | 2.3E-260 |
| Car4 | 33.78 | 2.57 | 4E-250 | 1.3E-246 |
| Ms4a18 | 31.83 | 2.89 | 2.2E-222 | 6E-219 |
| Reg3g | 29.66 | 1.72 | 2.3E-193 | 5E-190 |
| Ccl25 | 29.60 | 1.85 | 1.5E-192 | 2.8E-189 |
| Aldh1b1 | 28.65 | 1.78 | 1.6E-180 | 2.5E-177 |
| Pycard | 27.49 | 1.82 | 2.1E-166 | 2.9E-163 |
| Oat | 27.07 | 1.85 | 2.2E-161 | 2.9E-158 |
| Mt1 | 26.65 | 1.87 | 1.8E-156 | 2.2E-153 |
| Lgals4 | 26.56 | 1.56 | 2E-155 | 2.3E-152 |
| Maoa | 26.05 | 1.67 | 1.3E-149 | 1.1E-146 |
| Myo1d | 25.28 | 1.74 | 5.5E-141 | 4.1E-138 |
| Lgals2 | 24.61 | 1.57 | 9.5E-134 | 6.7E-131 |
| Uqcrc1 | 24.54 | 1.55 | 5.1E-133 | 3.5E-130 |
| Cyp4f40 | 24.43 | 2.11 | 7.9E-132 | 5.2E-129 |
| Apol10a | 24.08 | 1.73 | 4.1E-128 | 2.6E-125 |
| Gna11 | 22.96 | 1.51 | 1.1E-116 | 6.8E-114 |
| Ppp1r1b | 22.49 | 1.60 | 5.3E-112 | 2.9E-109 |
| Gmds | 22.34 | 1.55 | 1.5E-110 | 7.7E-108 |
| Reg3a | 22.27 | 2.16 | 7.2E-110 | 3.7E-107 |
| Bcl2l15 | 22.11 | 1.79 | 2.5E-108 | 1.2E-105 |
| Casp1 | 22.02 | 1.65 | 2E-107 | 9E-105 |
| Sema4g | 21.92 | 1.52 | 1.5E-106 | 6.7E-104 |
| Prpsap1 | 21.69 | 1.88 | 2.5E-104 | 1.1E-101 |
| Mgam2-ps | 20.70 | 1.96 | 3.69E-95 | 1.41E-92 |
| Gsr | 20.36 | 1.51 | 4E-92 | 1.44E-89 |
| Mt2 | 20.02 | 1.82 | 3.43E-89 | 1.17E-86 |
| Gpx1 | 19.86 | 1.56 | 9.19E-88 | 3.07E-85 |
| Adgrg7 | 18.73 | 1.51 | 2.83E-78 | 8.84E-76 |
| Hkdc1 | 18.64 | 1.65 | 1.53E-77 | 4.48E-75 |
| Arg2 | 17.28 | 1.60 | 6.49E-67 | 1.74E-64 |
| Bmp3 | 17.11 | 2.07 | 1.34E-65 | 3.46E-63 |
| Nlrp6 | 16.86 | 1.58 | 9.24E-64 | 2.23E-61 |
| Nr1h4 | 16.29 | 1.72 | 1.17E-59 | 2.54E-57 |
| Tstd1 | 16.11 | 2.14 | 2.3E-58 | 4.77E-56 |
| Cdkn2b | 15.86 | 1.65 | 1.16E-56 | 2.25E-54 |
| Akr1c19 | 15.84 | 2.05 | 1.7E-56 | 3.27E-54 |

| | | | | |
|---|---|---|---|---|
| Slc26a3 | 14.78 | 1.65 | 2.12E-49 | 3.26E-47 |
| Gnpnat1 | 14.63 | 1.67 | 1.93E-48 | 2.89E-46 |
| Abcc3 | 12.63 | 1.56 | 1.41E-36 | 1.45E-34 |
| Traf4 | 12.28 | 1.52 | 1.14E-34 | 1.12E-32 |
| Scd2 | 11.83 | 1.66 | 2.84E-32 | 2.52E-30 |
| Ms4a12 | 11.56 | 1.77 | 6.69E-31 | 5.62E-29 |
| Gsdmc4 | 11.52 | 2.08 | 9.95E-31 | 8.28E-29 |
| Myo7a | 11.46 | 1.74 | 2.19E-30 | 1.78E-28 |
| ccdc198 | 11.41 | 1.81 | 3.55E-30 | 2.84E-28 |
| Osbpl7 | 11.19 | 1.78 | 4.42E-29 | 3.44E-27 |
| Gsdmc2 | 10.96 | 2.01 | 5.9E-28 | 4.44E-26 |
| Gcnt4 | 10.84 | 1.52 | 2.32E-27 | 1.69E-25 |
| Elovl6 | 10.72 | 1.71 | 8.21E-27 | 5.91E-25 |
| Mcpt1 | 10.71 | 2.45 | 9.55E-27 | 6.82E-25 |
| Prkaa2 | 10.55 | 1.53 | 5.18E-26 | 3.58E-24 |
| Tldc2 | 10.53 | 1.98 | 6.12E-26 | 4.17E-24 |
| Myl7 | 10.11 | 1.91 | 4.82E-24 | 2.83E-22 |
| Dao | 10.00 | 3.03 | 1.58E-23 | 9.1E-22 |
| Hs3st1 | 9.92 | 1.55 | 3.32E-23 | 1.86E-21 |
| Itpka | 9.34 | 1.59 | 9.54E-21 | 4.56E-19 |
| Il18 | 9.23 | 1.84 | 2.81E-20 | 1.29E-18 |
| S100g | 8.96 | 1.88 | 3.19E-19 | 1.39E-17 |
| Ces1f | 8.73 | 1.98 | 2.54E-18 | 1.04E-16 |
| Vstm5 | 8.72 | 1.52 | 2.81E-18 | 1.15E-16 |
| Aldoc | 8.57 | 1.93 | 1.06E-17 | 4.18E-16 |
| Bche | 8.08 | 1.76 | 6.62E-16 | 2.36E-14 |
| Tmem25 | 7.69 | 1.91 | 1.49E-14 | 4.72E-13 |
| Mab21l4 | 7.31 | 1.54 | 2.7E-13 | 7.63E-12 |
| Lss | 7.29 | 1.55 | 3.2E-13 | 8.95E-12 |
| Prelid2 | 7.27 | 1.54 | 3.48E-13 | 9.72E-12 |
| Pwwp2b | 7.23 | 1.52 | 4.91E-13 | 1.35E-11 |
| Bfsp1 | 6.98 | 1.68 | 2.91E-12 | 7.34E-11 |
| Me3 | 6.57 | 1.85 | 5.05E-11 | 1.09E-09 |
| Mcpt2 | 6.54 | 2.18 | 6.22E-11 | 1.33E-09 |
| Slc28a3 | 6.32 | 1.79 | 2.66E-10 | 5.38E-09 |

Zone 4: Significantly Zonated Genes

| Symbol | scores | LFC | pvals | pvals_adj |
|---|---|---|---|---|
| Enpep | 40.22 | 2.41 | 0 | 0 |
| Anpep | 39.67 | 2.48 | 0 | 0 |
| Sis | 38.74 | 2.41 | 0 | 0 |
| Apoa1 | 36.95 | 2.43 | 7.6E-299 | 3.6E-295 |
| Fabp2 | 33.81 | 2.24 | 1.5E-250 | 5.7E-247 |
| Clec2h | 33.61 | 2.05 | 1.2E-247 | 3.8E-244 |
| Cubn | 32.43 | 2.15 | 9E-231 | 1.7E-227 |
| Slc5a1 | 32.37 | 1.98 | 8.2E-230 | 1.4E-226 |
| Reg3b | 32.25 | 1.89 | 3.1E-228 | 4.9E-225 |
| Mep1b | 31.07 | 1.91 | 5.5E-212 | 7.5E-209 |
| Pls1 | 29.79 | 1.84 | 4.9E-195 | 6.2E-192 |
| Naaladl1 | 28.90 | 1.73 | 1.2E-183 | 1.3E-180 |
| Spink1 | 28.49 | 1.77 | 1.5E-178 | 1.5E-175 |
| Slc51a | 28.24 | 1.81 | 1.7E-175 | 1.5E-172 |
| Atp1a1 | 28.17 | 1.65 | 1.3E-174 | 1.1E-171 |
| Crip1 | 28.08 | 1.61 | 2E-173 | 1.6E-170 |
| Maf | 25.75 | 1.68 | 3E-146 | 2.4E-143 |
| Vil1 | 25.20 | 1.55 | 3.7E-140 | 2.6E-137 |
| Maoa | 24.62 | 1.62 | 7.8E-134 | 4.8E-131 |
| Cyp4f14 | 24.34 | 1.55 | 7.3E-131 | 4.3E-128 |
| Slc43a2 | 24.27 | 1.73 | 4E-130 | 2.3E-127 |
| Fabp6 | 24.20 | 1.60 | 2.1E-129 | 1.2E-126 |
| Selenop | 24.07 | 1.60 | 4.9E-128 | 2.7E-125 |
| Aldh1a1 | 23.83 | 2.04 | 1.5E-125 | 7.9E-123 |
| Dpep1 | 23.82 | 1.57 | 2E-125 | 1E-122 |
| Mpp1 | 23.51 | 1.61 | 3.6E-122 | 1.8E-119 |
| Acsl5 | 23.08 | 1.52 | 7.3E-118 | 3.4E-115 |
| Slc13a2 | 23.04 | 1.58 | 2.1E-117 | 9.4E-115 |
| Tgfbi | 22.96 | 1.68 | 1.3E-116 | 5.6E-114 |
| Ace | 22.35 | 1.51 | 1.1E-110 | 4.7E-108 |
| Cndp2 | 22.22 | 1.52 | 2.3E-109 | 9.2E-107 |
| Ggt1 | 22.07 | 1.56 | 5.8E-108 | 2.3E-105 |
| Aspa | 20.18 | 1.54 | 1.6E-90 | 5.46E-88 |
| Khk | 17.90 | 1.54 | 1.16E-71 | 2.95E-69 |
| Slc5a4b | 17.32 | 1.85 | 3.5E-67 | 8.13E-65 |
| Slc5a4a | 16.36 | 1.95 | 3.87E-60 | 8.02E-58 |
| Adh6a | 15.86 | 1.55 | 1.28E-56 | 2.56E-54 |
| Slc5a11 | 15.56 | 1.79 | 1.4E-54 | 2.76E-52 |
| Slc26a3 | 13.40 | 1.61 | 6.23E-41 | 8.85E-39 |
| Slc2a5 | 12.83 | 1.74 | 1.13E-37 | 1.43E-35 |
| Nat8 | 12.21 | 1.57 | 2.59E-34 | 3.01E-32 |

| | | | | |
|---|---|---|---|---|
| Slc25a48 | 12.01 | 1.63 | 3.1E-33 | 3.41E-31 |
| Aldh1a7 | 11.86 | 2.06 | 1.94E-32 | 2.06E-30 |
| Angptl4 | 11.85 | 1.85 | 2.1E-32 | 2.21E-30 |
| Maob | 11.03 | 1.52 | 2.73E-28 | 2.41E-26 |
| Rdh7 | 9.89 | 1.66 | 4.46E-23 | 3.05E-21 |
| Bco2 | 8.18 | 1.91 | 2.8E-16 | 1.24E-14 |
| Gstm3 | 8.04 | 1.99 | 8.85E-16 | 3.76E-14 |
| Rhbg | 7.33 | 1.64 | 2.34E-13 | 7.96E-12 |
| Proz | 7.21 | 1.87 | 5.51E-13 | 1.78E-11 |
| Slc2a7 | 6.41 | 1.51 | 1.5E-10 | 3.63E-09 |

Zone 5: Significantly Zonated Genes

| Symbol | scores | LFC | pvals | pvals_adj |
|---|---|---|---|---|
| Fabp2 | 52.34 | 3.38 | 0 | 0 |
| Selenop | 49.20 | 3.09 | 0 | 0 |
| Apoa1 | 48.53 | 3.16 | 0 | 0 |
| Anpep | 41.50 | 2.62 | 0 | 0 |
| Clca4b | 41.37 | 2.60 | 0 | 0 |
| Slc15a1 | 40.44 | 2.52 | 0 | 0 |
| Ace2 | 39.78 | 2.47 | 0 | 0 |
| Crip1 | 36.15 | 2.01 | 3.9E-286 | 8.2E-283 |
| Slc6a19 | 35.33 | 2.30 | 1.9E-273 | 3.6E-270 |
| Tmigd1 | 32.93 | 2.22 | 9.1E-238 | 1.2E-234 |
| Naaladl1 | 32.88 | 1.94 | 4.6E-237 | 5.4E-234 |
| Sectm1b | 32.73 | 2.23 | 5.3E-235 | 6E-232 |
| Rbp2 | 32.02 | 2.13 | 5.3E-225 | 5.6E-222 |
| Dpp4 | 31.98 | 2.06 | 1.8E-224 | 1.8E-221 |
| Sgk1 | 31.98 | 2.18 | 2E-224 | 1.9E-221 |
| Slc5a1 | 31.57 | 1.93 | 9.1E-219 | 8.3E-216 |
| Spink1 | 31.45 | 1.92 | 4.6E-217 | 4E-214 |
| Pls1 | 31.33 | 1.91 | 1.6E-215 | 1.3E-212 |
| Cdhr5 | 30.88 | 1.82 | 2.2E-209 | 1.7E-206 |
| Cubn | 30.87 | 2.07 | 3.4E-209 | 2.5E-206 |
| Clec2h | 30.76 | 1.86 | 9E-208 | 6.4E-205 |
| Slc34a2 | 30.75 | 1.99 | 1.4E-207 | 9.7E-205 |
| Krt20 | 30.57 | 1.86 | 2.7E-205 | 1.8E-202 |
| Lct | 30.06 | 2.24 | 1.8E-198 | 1.1E-195 |
| Enpp7 | 29.49 | 2.02 | 4.4E-191 | 2.5E-188 |
| Pdzk1 | 28.99 | 2.02 | 7.9E-185 | 4.5E-182 |
| Slc6a8 | 28.49 | 1.80 | 1.7E-178 | 9.2E-176 |
| Cdhr2 | 28.14 | 1.81 | 2.8E-174 | 1.4E-171 |
| Gda | 27.97 | 1.86 | 3.5E-172 | 1.7E-169 |
| Pck1 | 27.38 | 2.14 | 4.8E-165 | 2.3E-162 |
| Rfk | 27.37 | 1.71 | 6E-165 | 2.7E-162 |
| Enpep | 27.12 | 1.68 | 6.5E-162 | 2.9E-159 |
| Ace | 27.11 | 1.80 | 6.7E-162 | 2.9E-159 |
| Slc27a4 | 27.11 | 1.82 | 8.4E-162 | 3.6E-159 |
| Pmp22 | 26.68 | 1.82 | 8.7E-157 | 3.6E-154 |
| Vil1 | 26.57 | 1.59 | 1.4E-155 | 5.6E-153 |
| Slc7a15 | 26.55 | 2.20 | 2.8E-155 | 1.1E-152 |
| Alpi | 25.93 | 1.82 | 3.1E-148 | 1.2E-145 |
| Slc13a2 | 25.80 | 1.78 | 9E-147 | 3.3E-144 |
| Cyp4f14 | 25.76 | 1.62 | 2.6E-146 | 9.5E-144 |
| Lap3 | 25.42 | 1.76 | 1.7E-142 | 5.9E-140 |

| | | | | |
|---|---|---|---|---|
| H2-Aa | 24.74 | 1.51 | 4.4E-135 | 1.4E-132 |
| Mep1b | 24.63 | 1.54 | 6.2E-134 | 2E-131 |
| Prap1 | 24.51 | 1.55 | 1.1E-132 | 3.3E-130 |
| Mme | 24.39 | 1.85 | 2.3E-131 | 7.1E-129 |
| H2-Q2 | 24.06 | 1.60 | 6.6E-128 | 2E-125 |
| Apoc3 | 23.79 | 1.75 | 4.3E-125 | 1.3E-122 |
| Ezr | 23.66 | 1.54 | 8.5E-124 | 2.5E-121 |
| Apob | 23.03 | 1.52 | 2.5E-117 | 6.9E-115 |
| Mxd1 | 22.85 | 1.52 | 1.5E-115 | 4.2E-113 |
| Maf | 22.75 | 1.51 | 1.3E-114 | 3.5E-112 |
| Lgals3 | 22.46 | 1.53 | 9.9E-112 | 2.6E-109 |
| Slc5a12 | 22.03 | 2.10 | 1.5E-107 | 3.7E-105 |
| Asah2 | 21.55 | 1.65 | 4.8E-103 | 1.2E-100 |
| Slc52a3 | 21.10 | 1.66 | 7.3E-99 | 1.71E-96 |
| Ggt1 | 21.02 | 1.51 | 4.58E-98 | 1.07E-95 |
| Aspa | 19.97 | 1.56 | 1.03E-88 | 2.26E-86 |
| Fabp1 | 19.53 | 1.91 | 5.7E-85 | 1.23E-82 |
| Cyp4v3 | 19.39 | 1.62 | 1E-83 | 2.1E-81 |
| Sprr2a3 | 18.66 | 1.57 | 1.06E-77 | 2.08E-75 |
| Slc5a4b | 18.59 | 1.96 | 3.62E-77 | 7.05E-75 |
| Apol7a | 18.42 | 1.60 | 8.73E-76 | 1.66E-73 |
| Stom | 17.42 | 1.55 | 5.7E-68 | 9.71E-66 |
| Treh | 16.96 | 1.69 | 1.73E-64 | 2.83E-62 |
| Exoc3l4 | 16.47 | 1.69 | 6.03E-61 | 9.57E-59 |
| Vnn1 | 16.37 | 1.52 | 2.92E-60 | 4.49E-58 |
| Slc2a2 | 15.91 | 2.20 | 5.56E-57 | 8.16E-55 |
| Dnase1 | 15.37 | 1.58 | 2.52E-53 | 3.43E-51 |
| Bst1 | 15.24 | 1.52 | 1.83E-52 | 2.43E-50 |
| Cyp2d26 | 14.75 | 1.50 | 2.95E-49 | 3.7E-47 |
| Aldh4a1 | 14.53 | 1.59 | 8.37E-48 | 9.97E-46 |
| Cyp3a11 | 13.94 | 1.74 | 3.75E-44 | 4.15E-42 |
| Fam118a | 13.34 | 1.51 | 1.3E-40 | 1.28E-38 |
| Leap2 | 13.26 | 1.85 | 4.05E-40 | 3.92E-38 |
| Slc5a4a | 12.91 | 1.63 | 3.81E-38 | 3.52E-36 |
| Slc6a14 | 12.78 | 1.74 | 2.07E-37 | 1.88E-35 |
| Cyp2b10 | 11.98 | 2.00 | 4.49E-33 | 3.52E-31 |
| Vwa1 | 11.41 | 1.60 | 3.92E-30 | 2.72E-28 |
| Aqp7 | 10.22 | 1.57 | 1.56E-24 | 8.3E-23 |
| Slc47a1 | 9.43 | 1.76 | 4.01E-21 | 1.68E-19 |
| Abcc5 | 8.89 | 1.91 | 6.14E-19 | 2.2E-17 |
| Nat8f5 | 7.78 | 1.71 | 7.36E-15 | 1.94E-13 |
| Tsku | 7.73 | 1.62 | 1.09E-14 | 2.8E-13 |
| Hes2 | 7.58 | 1.77 | 3.48E-14 | 8.53E-13 |
| Slc2a7 | 7.57 | 1.82 | 3.75E-14 | 9.1E-13 |

| | | | | |
|---|---|---|---|---|
| Ifit1 | 7.50 | 1.51 | 6.39E-14 | 1.5E-12 |
| Rnf208 | 7.10 | 1.79 | 1.23E-12 | 2.48E-11 |
| Kcnj13 | 7.08 | 1.69 | 1.4E-12 | 2.8E-11 |
| Rdh16 | 6.98 | 1.79 | 2.9E-12 | 5.55E-11 |
| Slc23a1 | 6.90 | 1.65 | 5.26E-12 | 9.73E-11 |
| Gbp8 | 6.78 | 1.58 | 1.16E-11 | 2.06E-10 |
| Slc28a1 | 6.78 | 1.73 | 1.17E-11 | 2.08E-10 |
| Npl | 6.61 | 1.51 | 3.8E-11 | 6.32E-10 |
| Nxpe3 | 6.42 | 1.96 | 1.32E-10 | 2.07E-09 |
| Car2 | 6.34 | 2.09 | 2.26E-10 | 3.44E-09 |

Zone 6: Significantly Zonated Genes

| Symbol | scores | LFC | pvals | pvals_adj |
|---|---|---|---|---|
| Ada | 83.42 | 5.97 | 0 | 0 |
| Apoa4 | 81.85 | 5.33 | 0 | 0 |
| Clca4a | 66.49 | 5.10 | 0 | 0 |
| Ly6m | 63.18 | 5.46 | 0 | 0 |
| Pmp22 | 62.77 | 3.96 | 0 | 0 |
| Clca4b | 61.42 | 3.75 | 0 | 0 |
| Slc28a2 | 60.84 | 5.10 | 0 | 0 |
| Selenop | 55.95 | 3.52 | 0 | 0 |
| Krt20 | 54.80 | 3.18 | 0 | 0 |
| Fabp2 | 53.64 | 3.47 | 0 | 0 |
| Apoa1 | 51.08 | 3.35 | 0 | 0 |
| Nt5e | 50.52 | 5.34 | 0 | 0 |
| Apob | 50.17 | 3.12 | 0 | 0 |
| Mxd1 | 48.88 | 3.05 | 0 | 0 |
| Muc3 | 47.92 | 2.74 | 0 | 0 |
| Slc23a4 | 46.96 | 4.51 | 0 | 0 |
| Slc6a8 | 46.70 | 2.83 | 0 | 0 |
| Apoc3 | 45.34 | 3.12 | 0 | 0 |
| Apoc2 | 44.86 | 3.19 | 0 | 0 |
| Creb3l3 | 42.91 | 3.58 | 0 | 0 |
| Rbp2 | 42.71 | 2.78 | 0 | 0 |
| Slc15a1 | 41.46 | 2.58 | 0 | 0 |
| Epcam | 41.07 | 2.11 | 0 | 0 |
| Slc34a2 | 38.02 | 2.41 | 0 | 0 |
| Crip1 | 36.76 | 2.09 | 7.2E-296 | 5.3E-293 |
| Alpi | 36.67 | 2.51 | 2.1E-294 | 1.5E-291 |
| Enpp3 | 35.99 | 4.75 | 1.4E-283 | 8.5E-281 |
| Jun | 35.81 | 3.29 | 8.7E-281 | 5.2E-278 |
| Ace2 | 35.51 | 2.20 | 3.2E-276 | 1.8E-273 |
| Slc6a19 | 35.45 | 2.32 | 2.8E-275 | 1.5E-272 |
| Serpinb1a | 35.11 | 2.19 | 4.1E-270 | 2.2E-267 |
| Rasgef1b | 34.97 | 3.70 | 7.6E-268 | 3.9E-265 |
| Ezr | 34.68 | 2.18 | 1.7E-263 | 8.6E-261 |
| Cdhr5 | 34.63 | 2.01 | 9.1E-263 | 4.4E-260 |
| Ccng2 | 34.61 | 2.55 | 2.1E-262 | 9.9E-260 |
| Lgals3 | 34.29 | 2.25 | 9.7E-258 | 4.5E-255 |
| Klf4 | 34.29 | 2.48 | 1E-257 | 4.7E-255 |
| Cdhr2 | 34.19 | 2.16 | 3.7E-256 | 1.6E-253 |
| Cldn7 | 34.19 | 1.91 | 3.9E-256 | 1.7E-253 |
| Cldn4 | 33.75 | 3.64 | 9.5E-250 | 3.8E-247 |
| Acta1 | 33.62 | 3.20 | 7.8E-248 | 3E-245 |

| | | | | |
|---|---|---|---|---|
| Rfk | 33.36 | 2.03 | 6.1E-244 | 2.3E-241 |
| Myo15b | 31.56 | 1.81 | 1.2E-218 | 4.3E-216 |
| Spink1 | 30.76 | 1.85 | 1E-207 | 3.5E-205 |
| H2-K1 | 30.74 | 1.82 | 1.7E-207 | 5.8E-205 |
| Fos | 30.59 | 2.85 | 1.5E-205 | 5E-203 |
| Anpep | 30.13 | 2.00 | 1.8E-199 | 5.9E-197 |
| S100a10 | 29.91 | 1.65 | 1.4E-196 | 4.5E-194 |
| Prap1 | 29.89 | 1.86 | 2.5E-196 | 7.8E-194 |
| Slc27a4 | 29.46 | 1.98 | 8.3E-191 | 2.5E-188 |
| Aldh4a1 | 29.45 | 2.86 | 1.3E-190 | 3.9E-188 |
| Ahnak | 29.03 | 1.93 | 3E-185 | 9.1E-183 |
| Npc1l1 | 28.97 | 2.06 | 1.7E-184 | 5E-182 |
| Pck1 | 28.84 | 2.27 | 6.7E-183 | 1.9E-180 |
| Bst1 | 28.58 | 2.63 | 1.1E-179 | 3.1E-177 |
| Pgm2 | 28.55 | 2.70 | 2.6E-179 | 7.1E-177 |
| Naaladl1 | 28.54 | 1.71 | 3.8E-179 | 1E-176 |
| Sgk1 | 28.54 | 1.99 | 4.2E-179 | 1.1E-176 |
| Slc25a22 | 28.46 | 2.66 | 3.3E-178 | 8.8E-176 |
| Cyp3a13 | 28.27 | 2.24 | 8.6E-176 | 2.2E-173 |
| Zfp36 | 28.15 | 2.04 | 2.6E-174 | 6.7E-172 |
| Susd2 | 27.11 | 3.16 | 8.5E-162 | 2.1E-159 |
| Bmp8a | 27.10 | 4.00 | 9.5E-162 | 2.4E-159 |
| Slc9a3r1 | 27.08 | 1.74 | 1.5E-161 | 3.7E-159 |
| Atf3 | 26.60 | 3.86 | 6.4E-156 | 1.5E-153 |
| Ifrd1 | 26.60 | 2.94 | 7.2E-156 | 1.7E-153 |
| Cystm1 | 25.83 | 1.72 | 3.7E-147 | 8.4E-145 |
| Ceacam1 | 25.36 | 1.53 | 7.2E-142 | 1.6E-139 |
| Plec | 25.19 | 1.75 | 5.6E-140 | 1.2E-137 |
| Rhob | 25.02 | 2.30 | 3.5E-138 | 7.4E-136 |
| Cidec | 25.00 | 4.12 | 5.5E-138 | 1.2E-135 |
| App | 24.75 | 1.52 | 2.8E-135 | 5.8E-133 |
| Sectm1b | 24.63 | 1.74 | 6.7E-134 | 1.4E-131 |
| Clec2h | 24.58 | 1.51 | 2.3E-133 | 4.6E-131 |
| Dnase1 | 24.55 | 2.34 | 4.3E-133 | 8.6E-131 |
| Aplp1 | 24.01 | 2.24 | 2.1E-127 | 4E-125 |
| Lpin2 | 23.98 | 2.30 | 4.8E-127 | 9.1E-125 |
| Gprc5a | 23.93 | 5.22 | 1.6E-126 | 3E-124 |
| Lmo7 | 23.58 | 2.19 | 5.8E-123 | 1.1E-120 |
| Sprr2a3 | 23.54 | 2.01 | 1.6E-122 | 2.8E-120 |
| Hsd17b11 | 23.32 | 1.68 | 2.5E-120 | 4.5E-118 |
| Serpinb6a | 23.17 | 1.65 | 9E-119 | 1.6E-116 |
| Dstn | 23.05 | 1.59 | 1.5E-117 | 2.6E-115 |
| Apol7a | 22.91 | 1.98 | 3.5E-116 | 6E-114 |
| Npc1 | 22.47 | 2.47 | 8E-112 | 1.3E-109 |

| | | | | |
|---|---|---|---|---|
| Max | 22.38 | 1.88 | 6E-111 | 9.8E-109 |
| Lama3 | 22.37 | 2.48 | 8.6E-111 | 1.4E-108 |
| H2-Q2 | 22.36 | 1.50 | 8.9E-111 | 1.4E-108 |
| Ugcg | 22.17 | 1.76 | 6.9E-109 | 1.1E-106 |
| Cyp3a11 | 22.09 | 2.54 | 3.9E-108 | 6E-106 |
| Dnpep | 21.88 | 2.10 | 4E-106 | 6.1E-104 |
| Itprid2 | 21.78 | 1.84 | 3.9E-105 | 5.9E-103 |
| Asah2 | 21.64 | 1.67 | 7.2E-104 | 1.1E-101 |
| Enpp7 | 21.57 | 1.53 | 3.4E-103 | 5E-101 |
| Tmigd1 | 21.53 | 1.51 | 8E-103 | 1.2E-100 |
| Tnks1bp1 | 21.50 | 2.23 | 1.5E-102 | 2.1E-100 |
| Rhoc | 21.50 | 1.87 | 1.6E-102 | 2.3E-100 |
| S100a6 | 21.39 | 2.21 | 1.7E-101 | 2.3E-99 |
| Tmbim1 | 21.26 | 1.70 | 2.4E-100 | 3.36E-98 |
| Fosl2 | 20.98 | 2.07 | 1.04E-97 | 1.41E-95 |
| Tent5a | 20.96 | 2.15 | 1.56E-97 | 2.11E-95 |
| Irf7 | 20.84 | 2.00 | 1.75E-96 | 2.35E-94 |
| Rhod | 20.66 | 2.39 | 7.33E-95 | 9.7E-93 |
| Ptprr | 20.56 | 2.47 | 6.85E-94 | 8.94E-92 |
| Tmem140 | 20.40 | 2.67 | 1.56E-92 | 1.98E-90 |
| Nuak2 | 20.39 | 1.89 | 1.99E-92 | 2.51E-90 |
| Duoxa2 | 20.25 | 2.36 | 3.66E-91 | 4.56E-89 |
| Mme | 20.18 | 1.60 | 1.57E-90 | 1.95E-88 |
| Eps8l3 | 20.11 | 1.58 | 5.85E-90 | 7.19E-88 |
| Selenoi | 20.07 | 1.93 | 1.39E-89 | 1.7E-87 |
| Nudt4 | 20.05 | 1.53 | 1.87E-89 | 2.27E-87 |
| Gtpbp2 | 20.05 | 1.67 | 1.9E-89 | 2.29E-87 |
| Mtmr11 | 19.84 | 1.89 | 1.23E-87 | 1.47E-85 |
| Plin3 | 19.60 | 1.54 | 1.43E-85 | 1.68E-83 |
| Emp1 | 19.58 | 1.87 | 2.19E-85 | 2.54E-83 |
| Noct | 19.22 | 2.35 | 2.3E-82 | 2.64E-80 |
| Trim25 | 19.12 | 1.81 | 1.75E-81 | 1.98E-79 |
| Hbegf | 19.10 | 2.77 | 2.76E-81 | 3.11E-79 |
| Acot12 | 19.03 | 4.98 | 8.84E-81 | 9.91E-79 |
| S100a11 | 18.92 | 1.68 | 7.24E-80 | 8.07E-78 |
| Kyat1 | 18.81 | 2.75 | 6.01E-79 | 6.58E-77 |
| Abhd2 | 18.62 | 1.70 | 2.37E-77 | 2.54E-75 |
| Skil | 18.51 | 1.92 | 1.64E-76 | 1.75E-74 |
| Ptrh1 | 18.33 | 2.55 | 4.69E-75 | 4.86E-73 |
| Rhof | 18.00 | 2.41 | 1.9E-72 | 1.94E-70 |
| Igf2 | 17.98 | 2.83 | 2.56E-72 | 2.59E-70 |
| Ston2 | 17.92 | 2.71 | 7.68E-72 | 7.71E-70 |
| Grina | 17.74 | 1.60 | 1.99E-70 | 1.97E-68 |
| Ptger4 | 17.55 | 2.67 | 6.39E-69 | 6.21E-67 |

| | | | | |
|---|---|---|---|---|
| Oasl1 | 17.49 | 2.80 | 1.72E-68 | 1.66E-66 |
| Marcksl1 | 17.40 | 2.21 | 7.63E-68 | 7.31E-66 |
| Treh | 17.16 | 1.72 | 5.58E-66 | 5.29E-64 |
| Slc9a2 | 17.13 | 1.72 | 9.27E-66 | 8.74E-64 |
| Hspa12a | 17.07 | 2.27 | 2.43E-65 | 2.28E-63 |
| Lratd1 | 16.82 | 2.21 | 1.6E-63 | 1.47E-61 |
| Stk17b | 16.71 | 1.53 | 1.17E-62 | 1.06E-60 |
| Gzma | 16.68 | 1.98 | 1.82E-62 | 1.64E-60 |
| Duox2 | 16.58 | 1.66 | 1.05E-61 | 9.42E-60 |
| Isg15 | 16.53 | 2.36 | 2.36E-61 | 2.1E-59 |
| Fam83b | 16.31 | 2.17 | 8.88E-60 | 7.76E-58 |
| Pdlim2 | 16.26 | 1.90 | 1.98E-59 | 1.72E-57 |
| Tuft1 | 16.18 | 2.17 | 7.09E-59 | 6.15E-57 |
| Dusp6 | 16.08 | 2.14 | 3.26E-58 | 2.81E-56 |
| Sfn | 16.02 | 1.97 | 9.96E-58 | 8.48E-56 |
| Bmp8b | 15.98 | 4.03 | 1.68E-57 | 1.42E-55 |
| Apol9a | 15.90 | 3.56 | 5.87E-57 | 4.95E-55 |
| Lhfpl2 | 15.78 | 2.40 | 4.58E-56 | 3.83E-54 |
| Slc39a4 | 15.77 | 1.63 | 4.78E-56 | 3.98E-54 |
| Igsf23 | 15.76 | 2.00 | 5.45E-56 | 4.5E-54 |
| Lrrfip1 | 15.73 | 1.83 | 8.84E-56 | 7.26E-54 |
| Itgb5 | 15.65 | 1.56 | 3.59E-55 | 2.92E-53 |
| Gramd1b | 15.49 | 1.98 | 4.21E-54 | 3.34E-52 |
| Insig2 | 15.46 | 1.72 | 6.34E-54 | 5E-52 |
| Cyp4b1 | 15.36 | 1.86 | 2.91E-53 | 2.27E-51 |
| Hsd17b2 | 15.23 | 1.61 | 2.21E-52 | 1.68E-50 |
| Sprr1a | 15.20 | 4.06 | 3.38E-52 | 2.54E-50 |
| Ifit1bl2 | 15.06 | 1.67 | 3.04E-51 | 2.25E-49 |
| Smpdl3b | 15.04 | 1.95 | 3.85E-51 | 2.84E-49 |
| Cox6b2 | 14.94 | 2.71 | 1.83E-50 | 1.32E-48 |
| Tubb2a | 14.93 | 2.78 | 2.24E-50 | 1.61E-48 |
| Ndfip2 | 14.89 | 1.52 | 3.87E-50 | 2.77E-48 |
| Stk10 | 14.73 | 2.05 | 4.03E-49 | 2.82E-47 |
| Plekha1 | 14.62 | 1.79 | 2.19E-48 | 1.51E-46 |
| Ccl5 | 14.60 | 1.95 | 2.95E-48 | 2.02E-46 |
| Itgb6 | 14.55 | 2.35 | 6.08E-48 | 4.12E-46 |
| Oas3 | 14.53 | 2.37 | 8.33E-48 | 5.59E-46 |
| Prdm1 | 14.44 | 3.00 | 2.71E-47 | 1.8E-45 |
| Ceacam18 | 14.44 | 2.96 | 2.94E-47 | 1.95E-45 |
| Slc6a14 | 14.43 | 2.05 | 3.38E-47 | 2.22E-45 |
| Ephx2 | 14.27 | 1.76 | 3.47E-46 | 2.24E-44 |
| Cxcl14 | 14.26 | 2.55 | 4.03E-46 | 2.59E-44 |
| Vegfa | 14.16 | 1.66 | 1.57E-45 | 1E-43 |
| Gal3st2 | 14.12 | 1.52 | 2.82E-45 | 1.79E-43 |

| | | | | |
|---|---|---|---|---|
| Pxdc1 | 14.10 | 2.43 | 4.04E-45 | 2.55E-43 |
| Sgk2 | 14.03 | 2.53 | 1E-44 | 6.24E-43 |
| Aqp7 | 13.97 | 2.10 | 2.29E-44 | 1.41E-42 |
| Ptpre | 13.48 | 1.82 | 2.03E-41 | 1.17E-39 |
| Arl14 | 13.36 | 2.72 | 1.1E-40 | 6.12E-39 |
| Relb | 13.31 | 1.51 | 1.98E-40 | 1.09E-38 |
| Ypel5 | 13.25 | 1.67 | 4.63E-40 | 2.53E-38 |
| Aqp3 | 13.24 | 2.56 | 5.06E-40 | 2.75E-38 |
| Fam102a | 13.23 | 1.73 | 6.02E-40 | 3.27E-38 |
| Unc5b | 12.88 | 1.64 | 5.7E-38 | 2.95E-36 |
| Tmem43 | 12.87 | 2.10 | 6.72E-38 | 3.46E-36 |
| Cd55 | 12.75 | 2.58 | 3.32E-37 | 1.69E-35 |
| Dgat2 | 12.67 | 1.57 | 8.31E-37 | 4.16E-35 |
| Junb | 12.66 | 1.53 | 9.81E-37 | 4.88E-35 |
| Tmcc3 | 12.56 | 1.51 | 3.64E-36 | 1.8E-34 |
| Slc17a5 | 12.32 | 2.14 | 7.43E-35 | 3.46E-33 |
| Synpo | 12.27 | 1.65 | 1.28E-34 | 5.82E-33 |
| Tlcd2 | 12.16 | 2.57 | 4.9E-34 | 2.18E-32 |
| Leng9 | 12.07 | 2.02 | 1.56E-33 | 6.81E-32 |
| Entpd8 | 11.87 | 1.58 | 1.75E-32 | 7.25E-31 |
| Slc16a3 | 11.39 | 2.04 | 4.56E-30 | 1.75E-28 |
| Il1rl1 | 11.35 | 3.45 | 7.07E-30 | 2.7E-28 |
| Usp35 | 11.22 | 2.66 | 3.36E-29 | 1.24E-27 |
| Rap2b | 11.11 | 2.33 | 1.1E-28 | 3.98E-27 |
| Sertad1 | 11.10 | 1.54 | 1.21E-28 | 4.37E-27 |
| Creb3l2 | 11.07 | 1.75 | 1.68E-28 | 5.99E-27 |
| Cd36 | 11.06 | 2.05 | 1.89E-28 | 6.69E-27 |
| Lamb3 | 11.00 | 1.76 | 3.88E-28 | 1.35E-26 |
| Clic1 | 10.98 | 1.51 | 4.76E-28 | 1.64E-26 |
| Dusp10 | 10.72 | 2.60 | 7.97E-27 | 2.57E-25 |
| Acsf2 | 10.69 | 1.67 | 1.1E-26 | 3.52E-25 |
| Unc93a2 | 10.64 | 1.74 | 1.91E-26 | 6.07E-25 |
| Gch1 | 10.57 | 2.10 | 3.94E-26 | 1.23E-24 |
| Ido1 | 10.55 | 2.47 | 5.14E-26 | 1.59E-24 |
| St3gal1 | 10.40 | 1.93 | 2.61E-25 | 7.73E-24 |
| Plau | 10.39 | 2.86 | 2.79E-25 | 8.25E-24 |
| Ifit1 | 10.17 | 2.08 | 2.8E-24 | 7.91E-23 |
| March3 | 10.15 | 3.47 | 3.38E-24 | 9.5E-23 |
| Pmaip1 | 9.99 | 1.64 | 1.75E-23 | 4.74E-22 |
| Ppm1j | 9.91 | 2.71 | 3.92E-23 | 1.05E-21 |
| Trim15 | 9.73 | 1.70 | 2.23E-22 | 5.71E-21 |
| Rab30 | 9.61 | 2.64 | 7.52E-22 | 1.88E-20 |
| Fbxo32 | 9.48 | 2.03 | 2.59E-21 | 6.24E-20 |
| Prkx | 9.42 | 1.89 | 4.71E-21 | 1.12E-19 |

| | | | | |
|---|---|---|---|---|
| Cd274 | 9.29 | 2.71 | 1.51E-20 | 3.46E-19 |
| Egr1 | 9.20 | 2.10 | 3.49E-20 | 7.77E-19 |
| Foxq1 | 8.85 | 2.60 | 9.14E-19 | 1.86E-17 |
| Capn2 | 8.76 | 1.55 | 1.98E-18 | 3.94E-17 |
| Dedd2 | 8.68 | 1.53 | 3.91E-18 | 7.67E-17 |
| Col6a4 | 8.67 | 2.07 | 4.21E-18 | 8.21E-17 |
| Nckap5 | 8.51 | 2.09 | 1.68E-17 | 3.12E-16 |
| Gas2l3 | 8.48 | 2.33 | 2.3E-17 | 4.22E-16 |
| Rapgef2 | 8.44 | 1.75 | 3.06E-17 | 5.59E-16 |
| Ankrd37 | 8.28 | 1.86 | 1.24E-16 | 2.16E-15 |
| Zfp37 | 8.24 | 2.36 | 1.68E-16 | 2.89E-15 |
| Slc6a18 | 8.22 | 2.42 | 2.07E-16 | 3.55E-15 |
| Asap1 | 8.13 | 2.03 | 4.22E-16 | 7.02E-15 |
| Plxna2 | 8.09 | 1.69 | 6.11E-16 | 1.01E-14 |
| Slc46a1 | 8.07 | 1.99 | 7.24E-16 | 1.19E-14 |
| P2ry2 | 8.03 | 1.57 | 1E-15 | 1.61E-14 |
| Gzmb | 7.99 | 1.64 | 1.3E-15 | 2.07E-14 |
| Nts | 7.87 | 2.04 | 3.45E-15 | 5.35E-14 |
| Rtp4 | 7.80 | 1.71 | 6.1E-15 | 9.22E-14 |
| Inhba | 7.78 | 3.76 | 7.19E-15 | 1.08E-13 |
| Lipa | 7.78 | 1.56 | 7.52E-15 | 1.13E-13 |
| Gal3st2b | 7.63 | 1.64 | 2.37E-14 | 3.39E-13 |
| Utp14b | 7.63 | 2.42 | 2.38E-14 | 3.4E-13 |
| Lpar3 | 7.51 | 2.23 | 5.87E-14 | 8.03E-13 |
| Sct | 7.35 | 1.57 | 1.97E-13 | 2.57E-12 |
| Rarres1 | 7.27 | 3.64 | 3.47E-13 | 4.43E-12 |
| Ccdc120 | 7.18 | 2.48 | 7.23E-13 | 8.98E-12 |
| Nyap1 | 7.11 | 1.68 | 1.13E-12 | 1.37E-11 |
| Maff | 6.91 | 1.74 | 4.98E-12 | 5.67E-11 |
| Ttll2 | 6.87 | 2.15 | 6.26E-12 | 7.06E-11 |
| Pdk4 | 6.77 | 3.25 | 1.32E-11 | 1.43E-10 |
| Rsad2 | 6.63 | 1.94 | 3.3E-11 | 3.42E-10 |
| Ubd | 6.63 | 1.59 | 3.42E-11 | 3.53E-10 |
| Wnt5a | 6.61 | 2.07 | 3.75E-11 | 3.85E-10 |
| Acsl3 | 6.60 | 1.51 | 4.2E-11 | 4.29E-10 |
| Foxo6 | 6.51 | 1.88 | 7.33E-11 | 7.31E-10 |
| Plk3 | 6.50 | 1.86 | 8.22E-11 | 8.15E-10 |
| Gml2 | 6.40 | 3.48 | 1.53E-10 | 1.47E-09 |
| Smox | 6.40 | 1.54 | 1.54E-10 | 1.47E-09 |
| Dusp1 | 6.31 | 1.50 | 2.76E-10 | 2.57E-09 |
| Ltb4r2 | 6.20 | 1.85 | 5.66E-10 | 5.08E-09 |
| Glp2r | 6.09 | 2.31 | 1.15E-09 | 9.93E-09 |
