## Supplementary Text for "SMURF: soft-segmentation for single-cell reconstruction and topological analysis of spatial transcriptomic data"

#### Spatial scRNA-seq Mirrors Single-Cell Statistics: Gene-Wise NB Fits SMURF Data

Bulk RNA-seq analysis aggregates gene expression measurements from large pools of cells, providing a population-level snapshot of transcriptional activity within a sample [1]. Many computational tools for analyzing bulk RNA-seq data are built on statistical distributions that model the discrete and often overdispersed count data generated in these experiments. The Poisson model assumes that the variance equals the mean; however, this assumption may prove inadequate when overdispersion is present. The Negative Binomial (NB) model overcomes this limitation by incorporating an additional dispersion parameter, which quantifies the extent to which the variance exceeds the mean [2, 3, 4].

In contrast to bulk RNA-seq, scRNA-seq data typically contain a high degree of observed zero counts (commonly referred to as "dropouts"). Zero counts arise from

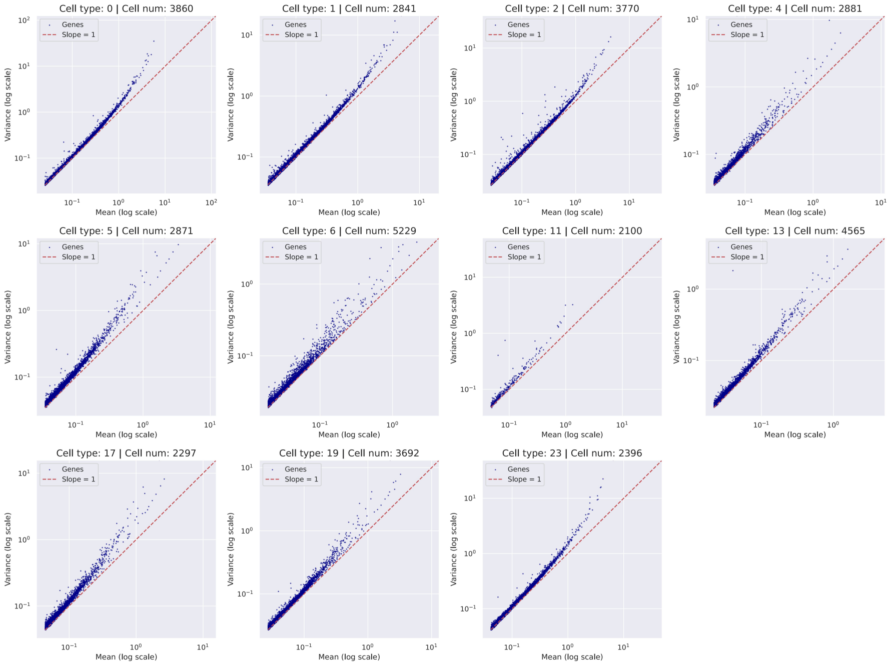


**Fig. S1**: Mean vs. Variance (mouse brain dataset)

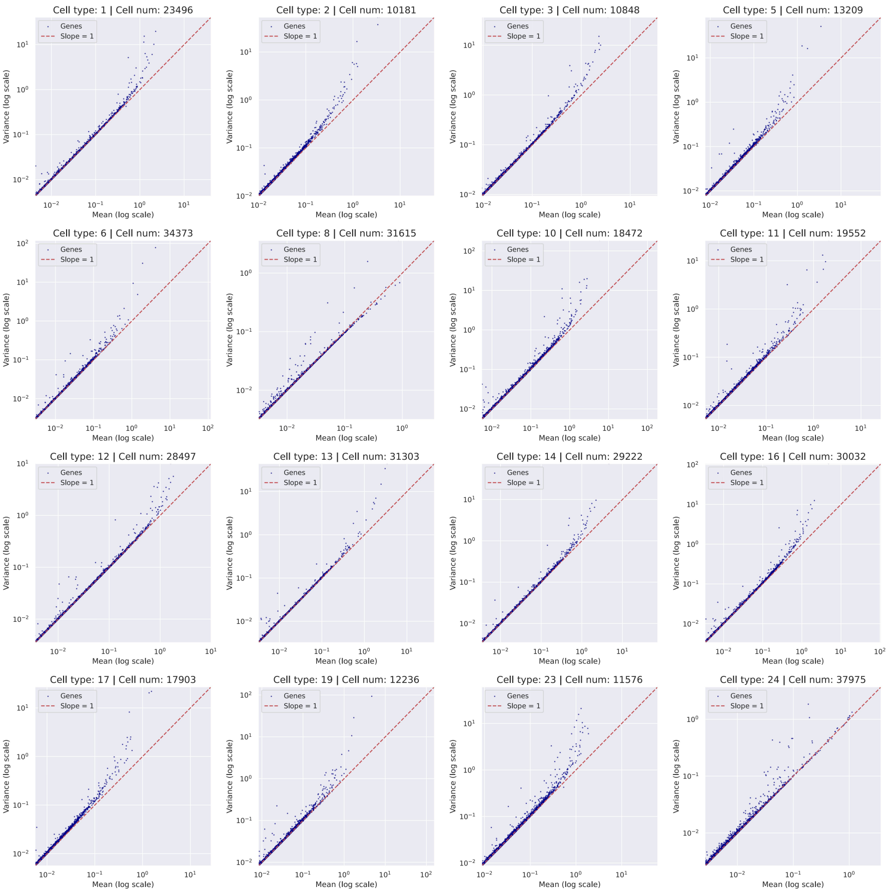


**Fig. S2**: Mean vs. Variance (mouse small intestine dataset)

both biological and technical factors. When excessive zero counts occur, the zero-inflated negative binomial (ZINB) model can be employed. This model effectively separates the NB component from the processes generating zero counts, offering a more flexible fit to the heterogeneous count distributions observed in single-cell transcriptomic data [5, 6]. However, recent studies on UMI-based protocols have demonstrated that many zero counts can be adequately modeled using simpler Poisson or NB distributions after accounting for cell-type heterogeneity, thereby reducing the need for additional zero-inflation components [7, 8, 9].

These insights have spurred the development of a wide range of computational methods for both bulk and single-cell RNA-seq analyses. For instance, tools such as edgeR [3] and DESeq2 [4] employ NB models to account for biological variability. Additionally, scvi is a deep generative framework that allows users to assume that counts follow various distributions — including the NB, ZINB, Poisson, or even the normal distribution — to capture the inherent variability in single-cell transcriptomic data [10].

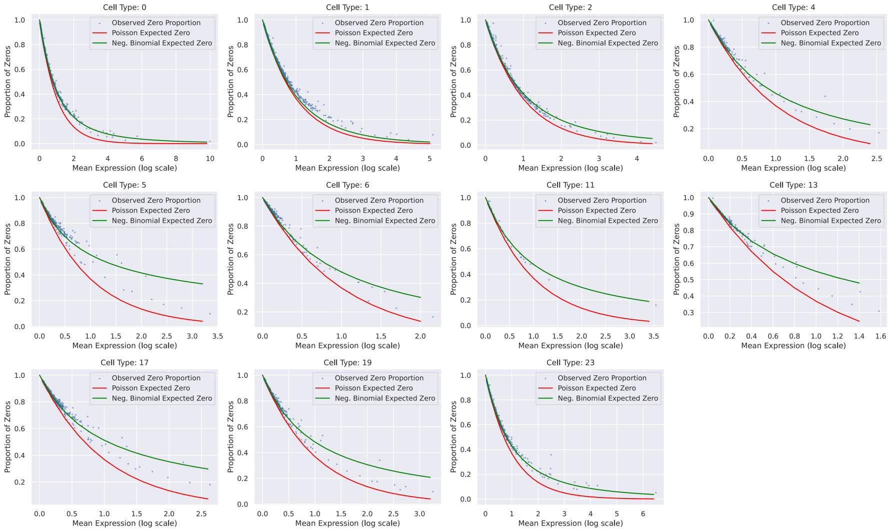


**Fig. S3**: Comparisons of zero proportions in different clusters of the mouse brain dataset

**Table S1**: Mouse Brain Data

| Cell type | Cell number | Gene number | Model | Number and percentage of genes over x percentage points away from expected fraction | | | | | | | |
|---|---|---|---|---|---|---|---|---|---|---|---|
| | | | | 1 pp count | 1 pp percentage | 5 pp count | 5 pp percentage | 10 pp count | 10 pp percentage | 20 pp count | 20 pp percentage |
| 0 | 3993 | 1155 | 1. Poisson | 1088.00 | 94.20 | 344.00 | 29.78 | 180.00 | 15.58 | 94.00 | 8.14 |
| 0 | 3993 | 1155 | 2. NB common | 373.00 | 32.29 | 65.00 | 5.63 | 25.00 | 2.16 | 8.00 | 0.69 |
| 0 | 3993 | 1155 | 3. NB gene-wise | 163.00 | 14.11 | 52.00 | 4.50 | 25.00 | 2.16 | 14.00 | 1.21 |
| 0 | 3993 | 1155 | 4. ZINB | 165.00 | 14.29 | 52.00 | 4.50 | 25.00 | 2.16 | 14.00 | 1.21 |
| 5 | 2497 | 467 | 1. Poisson | 465.00 | 99.57 | 236.00 | 50.54 | 122.00 | 26.12 | 64.00 | 13.70 |
| 5 | 2497 | 467 | 2. NB common | 260.00 | 55.67 | 49.00 | 10.49 | 17.00 | 3.64 | 10.00 | 2.14 |
| 5 | 2497 | 467 | 3. NB gene-wise | 103.00 | 22.06 | 28.00 | 6.00 | 14.00 | 3.00 | 6.00 | 1.28 |
| 5 | 2497 | 467 | 4. ZINB | 103.00 | 22.06 | 28.00 | 6.00 | 14.00 | 3.00 | 6.00 | 1.28 |
| 7 | 4214 | 79 | 1. Poisson | 79.00 | 100.00 | 46.00 | 58.23 | 17.00 | 21.52 | 8.00 | 10.13 |
| 7 | 4214 | 79 | 2. NB common | 53.00 | 67.09 | 11.00 | 13.92 | 6.00 | 7.59 | 2.00 | 2.53 |
| 7 | 4214 | 79 | 3. NB gene-wise | 14.00 | 17.72 | 2.00 | 2.53 | 0.00 | 0.00 | 0.00 | 0.00 |
| 7 | 4214 | 79 | 4. ZINB | 14.00 | 17.72 | 2.00 | 2.53 | 0.00 | 0.00 | 0.00 | 0.00 |
| 8 | 2884 | 44 | 1. Poisson | 43.00 | 97.73 | 33.00 | 75.00 | 22.00 | 50.00 | 12.00 | 27.27 |
| 8 | 2884 | 44 | 2. NB common | 41.00 | 93.18 | 18.00 | 40.91 | 11.00 | 25.00 | 5.00 | 11.36 |
| 8 | 2884 | 44 | 3. NB gene-wise | 17.00 | 38.64 | 4.00 | 9.09 | 2.00 | 4.55 | 0.00 | 0.00 |
| 8 | 2884 | 44 | 4. ZINB | 17.00 | 38.64 | 4.00 | 9.09 | 2.00 | 4.55 | 0.00 | 0.00 |
| 10 | 2621 | 312 | 1. Poisson | 312.00 | 100.00 | 227.00 | 72.76 | 110.00 | 35.26 | 55.00 | 17.63 |
| 10 | 2621 | 312 | 2. NB common | 179.00 | 57.37 | 38.00 | 12.18 | 15.00 | 4.81 | 7.00 | 2.24 |
| 10 | 2621 | 312 | 3. NB gene-wise | 68.00 | 21.79 | 20.00 | 6.41 | 15.00 | 4.81 | 7.00 | 2.24 |
| 10 | 2621 | 312 | 4. ZINB | 67.00 | 21.47 | 20.00 | 6.41 | 15.00 | 4.81 | 7.00 | 2.24 |
| 12 | 2059 | 73 | 1. Poisson | 73.00 | 100.00 | 72.00 | 98.63 | 71.00 | 97.26 | 44.00 | 60.27 |
| 12 | 2059 | 73 | 2. NB common | 71.00 | 97.26 | 54.00 | 73.97 | 27.00 | 36.99 | 13.00 | 17.81 |
| 12 | 2059 | 73 | 3. NB gene-wise | 15.00 | 20.55 | 6.00 | 8.22 | 4.00 | 5.48 | 3.00 | 4.11 |
| 12 | 2059 | 73 | 4. ZINB | 12.00 | 16.44 | 6.00 | 8.22 | 4.00 | 5.48 | 3.00 | 4.11 |
| 14 | 3216 | 97 | 1. Poisson | 75.00 | 77.32 | 47.00 | 48.45 | 27.00 | 27.84 | 13.00 | 13.40 |
| 14 | 3216 | 97 | 2. NB common | 73.00 | 75.26 | 33.00 | 34.02 | 15.00 | 15.46 | 5.00 | 5.15 |
| 14 | 3216 | 97 | 3. NB gene-wise | 43.00 | 44.33 | 13.00 | 13.40 | 7.00 | 7.22 | 1.00 | 1.03 |
| 14 | 3216 | 97 | 4. ZINB | 41.00 | 42.27 | 13.00 | 13.40 | 7.00 | 7.22 | 1.00 | 1.03 |
| 15 | 3697 | 35 | 1. Poisson | 32.00 | 91.43 | 17.00 | 48.57 | 13.00 | 37.14 | 6.00 | 17.14 |
| 15 | 3697 | 35 | 2. NB common | 31.00 | 88.57 | 14.00 | 40.00 | 7.00 | 20.00 | 0.00 | 0.00 |
| 15 | 3697 | 35 | 3. NB gene-wise | 14.00 | 40.00 | 2.00 | 5.71 | 0.00 | 0.00 | 0.00 | 0.00 |
| 15 | 3697 | 35 | 4. ZINB | 14.00 | 40.00 | 2.00 | 5.71 | 0.00 | 0.00 | 0.00 | 0.00 |
| 18 | 2142 | 53 | 1. Poisson | 52.00 | 98.11 | 52.00 | 98.11 | 45.00 | 84.91 | 26.00 | 49.06 |
| 18 | 2142 | 53 | 2. NB common | 52.00 | 98.11 | 34.00 | 64.15 | 18.00 | 33.96 | 9.00 | 16.98 |
| 18 | 2142 | 53 | 3. NB gene-wise | 27.00 | 50.94 | 9.00 | 16.98 | 4.00 | 7.55 | 1.00 | 1.89 |
| 18 | 2142 | 53 | 4. ZINB | 27.00 | 50.94 | 9.00 | 16.98 | 4.00 | 7.55 | 1.00 | 1.89 |
| 19 | 3939 | 852 | 1. Poisson | 630.00 | 73.94 | 140.00 | 16.43 | 72.00 | 8.45 | 28.00 | 3.29 |
| 19 | 3939 | 852 | 2. NB common | 276.00 | 32.39 | 64.00 | 7.51 | 27.00 | 3.17 | 10.00 | 1.17 |
| 19 | 3939 | 852 | 3. NB gene-wise | 178.00 | 20.89 | 58.00 | 6.81 | 30.00 | 3.52 | 15.00 | 1.76 |
| 19 | 3939 | 852 | 4. ZINB | 179.00 | 21.01 | 58.00 | 6.81 | 30.00 | 3.52 | 15.00 | 1.76 |
| 23 | 3034 | 111 | 1. Poisson | 111.00 | 100.00 | 77.00 | 69.37 | 44.00 | 39.64 | 23.00 | 20.72 |
| 23 | 3034 | 111 | 2. NB common | 97.00 | 87.39 | 46.00 | 41.44 | 25.00 | 22.52 | 11.00 | 9.91 |
| 23 | 3034 | 111 | 3. NB gene-wise | 36.00 | 32.43 | 10.00 | 9.01 | 8.00 | 7.21 | 5.00 | 4.50 |
| 23 | 3034 | 111 | 4. ZINB | 36.00 | 32.43 | 10.00 | 9.01 | 8.00 | 7.21 | 5.00 | 4.50 |

Note: pp=percentage points.

The observed counts in cells from spatial transcriptomics data are frequently even lower than what is observed in dissociation-based scRNA-seq. For example, SMURF identified a median of 426 UMIs per cell in the mouse brain, and only 165 UMIs

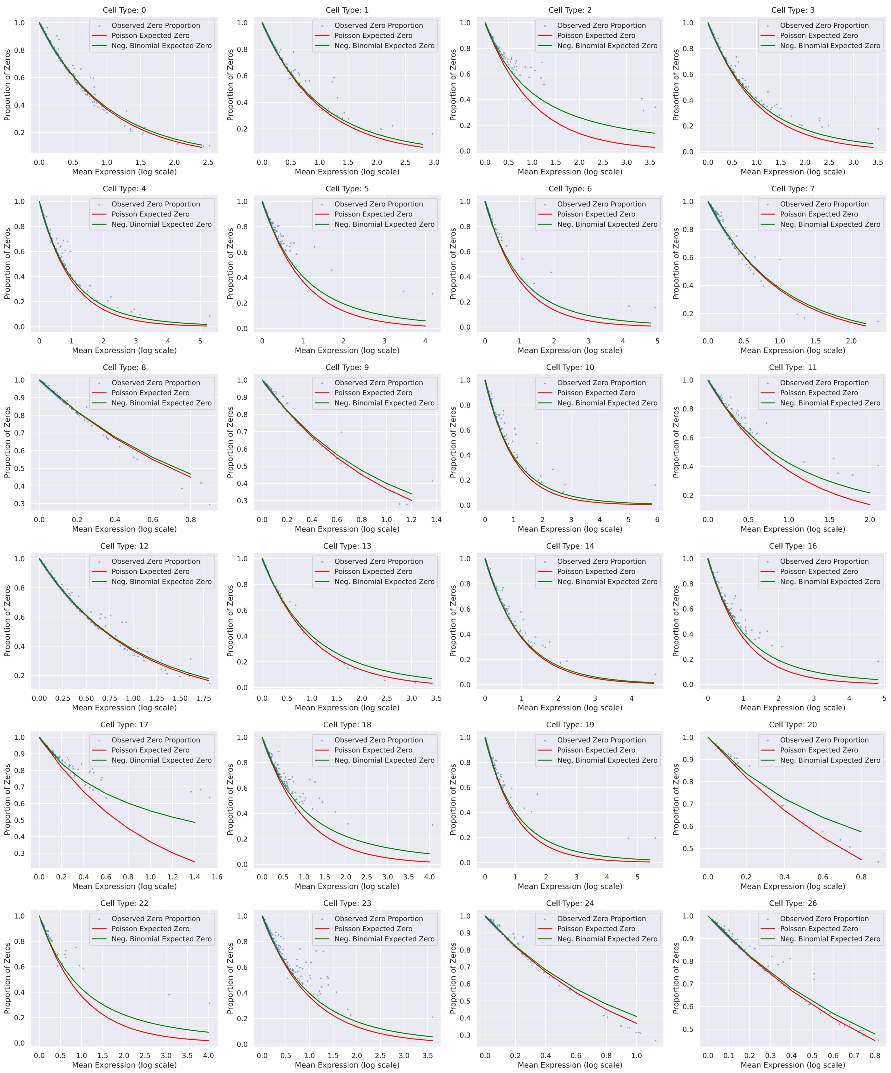


**Fig. S4**: Comparisons of zero proportions in different clusters of the mouse small intestine dataset

per cell in the mouse small intestine. In contrast, current scRNA-seq protocols can capture more than 7,000 UMIs per cell when deeply sequenced [11]. Despite the exploding interest in spatial transcriptomics methods, we do not know the best model to fit these datasets that accounts for the high frequency of zero counts.

To evaluate the distribution of SMURF-based spatial single-cell data, we began by examining dispersion under the assumption that cells within the same cluster are homogeneous. We then analyzed different cell types separately, focusing on those with more than 1,000 cells (mouse brain) and those with more than 10,000 cells (mouse small intestine). Focusing on genes with more than 100 total counts across all cells,

**Table S2**: Mouse Small Intestine Data

| Cell type | Cell number | Gene number | Model | Number and percentage of genes over x percentage points away from expected fraction | | | | | | | |
|---|---|---|---|---|---|---|---|---|---|---|---|
| | | | | 1 pp count | 1 pp percentage | 5 pp count | 5 pp percentage | 10 pp count | 10 pp percentage | 20 pp count | 20 pp percentage |
| 1 | 20105 | 6754 | 1. Poisson | 66.00 | 0.98 | 18.00 | 0.27 | 12.00 | 0.18 | 8.00 | 0.12 |
| 1 | 20105 | 6754 | 2. NB common | 59.00 | 0.87 | 16.00 | 0.24 | 10.00 | 0.15 | 6.00 | 0.09 |
| 1 | 20105 | 6754 | 3. NB gene-wise | 11.00 | 0.16 | 5.00 | 0.07 | 4.00 | 0.06 | 2.00 | 0.03 |
| 1 | 20105 | 6754 | 4. ZINB | 11.00 | 0.16 | 5.00 | 0.07 | 4.00 | 0.06 | 2.00 | 0.03 |
| 2 | 11405 | 3599 | 1. Poisson | 62.00 | 1.72 | 23.00 | 0.64 | 14.00 | 0.39 | 7.00 | 0.19 |
| 2 | 11405 | 3599 | 2. NB common | 61.00 | 1.69 | 21.00 | 0.58 | 13.00 | 0.36 | 7.00 | 0.19 |
| 2 | 11405 | 3599 | 3. NB gene-wise | 9.00 | 0.25 | 2.00 | 0.06 | 2.00 | 0.06 | 2.00 | 0.06 |
| 2 | 11405 | 3599 | 4. ZINB | 9.00 | 0.25 | 2.00 | 0.06 | 2.00 | 0.06 | 2.00 | 0.06 |
| 3 | 19358 | 8725 | 1. Poisson | 71.00 | 0.81 | 22.00 | 0.25 | 14.00 | 0.16 | 10.00 | 0.11 |
| 3 | 19358 | 8725 | 2. NB common | 61.00 | 0.70 | 22.00 | 0.25 | 14.00 | 0.16 | 7.00 | 0.08 |
| 3 | 19358 | 8725 | 3. NB gene-wise | 23.00 | 0.26 | 6.00 | 0.07 | 3.00 | 0.03 | 2.00 | 0.02 |
| 3 | 19358 | 8725 | 4. ZINB | 23.00 | 0.26 | 6.00 | 0.07 | 3.00 | 0.03 | 2.00 | 0.02 |
| 4 | 13598 | 1985 | 1. Poisson | 77.00 | 3.88 | 27.00 | 1.36 | 16.00 | 0.81 | 7.00 | 0.35 |
| 4 | 13598 | 1985 | 2. NB common | 56.00 | 2.82 | 18.00 | 0.91 | 9.00 | 0.45 | 3.00 | 0.15 |
| 4 | 13598 | 1985 | 3. NB gene-wise | 9.00 | 0.45 | 1.00 | 0.05 | 0.00 | 0.00 | 0.00 | 0.00 |
| 4 | 13598 | 1985 | 4. ZINB | 10.00 | 0.50 | 1.00 | 0.05 | 0.00 | 0.00 | 0.00 | 0.00 |
| 5 | 31265 | 7721 | 1. Poisson | 125.00 | 1.62 | 44.00 | 0.57 | 24.00 | 0.31 | 14.00 | 0.18 |
| 5 | 31265 | 7721 | 2. NB common | 81.00 | 1.05 | 30.00 | 0.39 | 16.00 | 0.21 | 11.00 | 0.14 |
| 5 | 31265 | 7721 | 3. NB gene-wise | 34.00 | 0.44 | 9.00 | 0.12 | 3.00 | 0.04 | 1.00 | 0.01 |
| 5 | 31265 | 7721 | 4. ZINB | 34.00 | 0.44 | 9.00 | 0.12 | 4.00 | 0.05 | 1.00 | 0.01 |
| 6 | 38613 | 7412 | 1. Poisson | 44.00 | 0.59 | 14.00 | 0.19 | 9.00 | 0.12 | 0.00 | 0.00 |
| 6 | 38613 | 7412 | 2. NB common | 46.00 | 0.62 | 14.00 | 0.19 | 10.00 | 0.13 | 1.00 | 0.01 |
| 6 | 38613 | 7412 | 3. NB gene-wise | 21.00 | 0.28 | 9.00 | 0.12 | 8.00 | 0.11 | 1.00 | 0.01 |
| 6 | 38613 | 7412 | 4. ZINB | 22.00 | 0.30 | 10.00 | 0.13 | 9.00 | 0.12 | 2.00 | 0.03 |
| 8 | 21261 | 7322 | 1. Poisson | 138.00 | 1.88 | 43.00 | 0.59 | 24.00 | 0.33 | 15.00 | 0.20 |
| 8 | 21261 | 7322 | 2. NB common | 148.00 | 2.02 | 49.00 | 0.67 | 28.00 | 0.38 | 15.00 | 0.20 |
| 8 | 21261 | 7322 | 3. NB gene-wise | 112.00 | 1.53 | 42.00 | 0.57 | 30.00 | 0.41 | 15.00 | 0.20 |
| 8 | 21261 | 7322 | 4. ZINB | 110.00 | 1.50 | 42.00 | 0.57 | 30.00 | 0.41 | 15.00 | 0.20 |
| 10 | 21947 | 5826 | 1. Poisson | 39.00 | 0.67 | 15.00 | 0.26 | 7.00 | 0.12 | 5.00 | 0.09 |
| 10 | 21947 | 5826 | 2. NB common | 35.00 | 0.60 | 13.00 | 0.22 | 6.00 | 0.10 | 5.00 | 0.09 |
| 10 | 21947 | 5826 | 3. NB gene-wise | 10.00 | 0.17 | 4.00 | 0.07 | 2.00 | 0.03 | 0.00 | 0.00 |
| 10 | 21947 | 5826 | 4. ZINB | 10.00 | 0.17 | 4.00 | 0.07 | 2.00 | 0.03 | 0.00 | 0.00 |
| 11 | 23973 | 6470 | 1. Poisson | 77.00 | 1.19 | 28.00 | 0.43 | 19.00 | 0.29 | 11.00 | 0.17 |
| 11 | 23973 | 6470 | 2. NB common | 55.00 | 0.85 | 19.00 | 0.29 | 12.00 | 0.19 | 7.00 | 0.11 |
| 11 | 23973 | 6470 | 3. NB gene-wise | 45.00 | 0.70 | 20.00 | 0.31 | 11.00 | 0.17 | 6.00 | 0.09 |
| 11 | 23973 | 6470 | 4. ZINB | 44.00 | 0.68 | 20.00 | 0.31 | 11.00 | 0.17 | 6.00 | 0.09 |
| 12 | 10841 | 5917 | 1. Poisson | 130.00 | 2.20 | 48.00 | 0.81 | 29.00 | 0.49 | 18.00 | 0.30 |
| 12 | 10841 | 5917 | 2. NB common | 85.00 | 1.44 | 30.00 | 0.51 | 21.00 | 0.35 | 15.00 | 0.25 |
| 12 | 10841 | 5917 | 3. NB gene-wise | 50.00 | 0.85 | 19.00 | 0.32 | 10.00 | 0.17 | 4.00 | 0.07 |
| 12 | 10841 | 5917 | 4. ZINB | 48.00 | 0.81 | 19.00 | 0.32 | 10.00 | 0.17 | 4.00 | 0.07 |
| 13 | 18292 | 7184 | 1. Poisson | 96.00 | 1.34 | 35.00 | 0.49 | 21.00 | 0.29 | 12.00 | 0.17 |
| 13 | 18292 | 7184 | 2. NB common | 81.00 | 1.13 | 29.00 | 0.40 | 20.00 | 0.28 | 9.00 | 0.13 |
| 13 | 18292 | 7184 | 3. NB gene-wise | 20.00 | 0.28 | 3.00 | 0.04 | 3.00 | 0.04 | 2.00 | 0.03 |
| 13 | 18292 | 7184 | 4. ZINB | 21.00 | 0.29 | 3.00 | 0.04 | 3.00 | 0.04 | 2.00 | 0.03 |
| 14 | 17651 | 7922 | 1. Poisson | 82.00 | 1.04 | 38.00 | 0.48 | 18.00 | 0.23 | 10.00 | 0.13 |
| 14 | 17651 | 7922 | 2. NB common | 62.00 | 0.78 | 27.00 | 0.34 | 12.00 | 0.15 | 9.00 | 0.11 |
| 14 | 17651 | 7922 | 3. NB gene-wise | 49.00 | 0.62 | 19.00 | 0.24 | 12.00 | 0.15 | 6.00 | 0.08 |
| 14 | 17651 | 7922 | 4. ZINB | 47.00 | 0.59 | 19.00 | 0.24 | 12.00 | 0.15 | 6.00 | 0.08 |
| 15 | 28810 | 10140 | 1. Poisson | 71.00 | 0.70 | 32.00 | 0.32 | 19.00 | 0.19 | 6.00 | 0.06 |
| 15 | 28810 | 10140 | 2. NB common | 86.00 | 0.85 | 32.00 | 0.32 | 20.00 | 0.20 | 6.00 | 0.06 |
| 15 | 28810 | 10140 | 3. NB gene-wise | 88.00 | 0.87 | 30.00 | 0.30 | 23.00 | 0.23 | 11.00 | 0.11 |
| 15 | 28810 | 10140 | 4. ZINB | 98.00 | 0.97 | 31.00 | 0.31 | 23.00 | 0.23 | 11.00 | 0.11 |
| 17 | 10088 | 4852 | 1. Poisson | 82.00 | 1.69 | 33.00 | 0.68 | 21.00 | 0.43 | 15.00 | 0.31 |
| 17 | 10088 | 4852 | 2. NB common | 94.00 | 1.94 | 36.00 | 0.74 | 24.00 | 0.49 | 16.00 | 0.33 |
| 17 | 10088 | 4852 | 3. NB gene-wise | 81.00 | 1.67 | 31.00 | 0.64 | 20.00 | 0.41 | 16.00 | 0.33 |
| 17 | 10088 | 4852 | 4. ZINB | 88.00 | 1.81 | 31.00 | 0.64 | 20.00 | 0.41 | 16.00 | 0.33 |
| 19 | 25711 | 8981 | 1. Poisson | 33.00 | 0.37 | 12.00 | 0.13 | 10.00 | 0.11 | 7.00 | 0.08 |
| 19 | 25711 | 8981 | 2. NB common | 30.00 | 0.33 | 12.00 | 0.13 | 12.00 | 0.13 | 8.00 | 0.09 |
| 19 | 25711 | 8981 | 3. NB gene-wise | 28.00 | 0.31 | 12.00 | 0.13 | 10.00 | 0.11 | 8.00 | 0.09 |
| 19 | 25711 | 8981 | 4. ZINB | 29.00 | 0.32 | 12.00 | 0.13 | 10.00 | 0.11 | 8.00 | 0.09 |
| 21 | 11165 | 6814 | 1. Poisson | 160.00 | 2.35 | 61.00 | 0.90 | 36.00 | 0.53 | 23.00 | 0.34 |
| 21 | 11165 | 6814 | 2. NB common | 125.00 | 1.83 | 50.00 | 0.73 | 31.00 | 0.45 | 19.00 | 0.28 |
| 21 | 11165 | 6814 | 3. NB gene-wise | 53.00 | 0.78 | 21.00 | 0.31 | 6.00 | 0.09 | 3.00 | 0.04 |
| 21 | 11165 | 6814 | 4. ZINB | 51.00 | 0.75 | 21.00 | 0.31 | 6.00 | 0.09 | 3.00 | 0.04 |
| 26 | 16372 | 6483 | 1. Poisson | 21.00 | 0.32 | 7.00 | 0.11 | 4.00 | 0.06 | 2.00 | 0.03 |
| 26 | 16372 | 6483 | 2. NB common | 29.00 | 0.45 | 8.00 | 0.12 | 3.00 | 0.05 | 2.00 | 0.03 |
| 26 | 16372 | 6483 | 3. NB gene-wise | 15.00 | 0.23 | 7.00 | 0.11 | 2.00 | 0.03 | 1.00 | 0.02 |
| 26 | 16372 | 6483 | 4. ZINB | 16.00 | 0.25 | 7.00 | 0.11 | 2.00 | 0.03 | 1.00 | 0.02 |
| 27 | 29083 | 7515 | 1. Poisson | 28.00 | 0.37 | 7.00 | 0.09 | 4.00 | 0.05 | 1.00 | 0.01 |
| 27 | 29083 | 7515 | 2. NB common | 30.00 | 0.40 | 8.00 | 0.11 | 4.00 | 0.05 | 1.00 | 0.01 |
| 27 | 29083 | 7515 | 3. NB gene-wise | 19.00 | 0.25 | 7.00 | 0.09 | 4.00 | 0.05 | 2.00 | 0.03 |
| 27 | 29083 | 7515 | 4. ZINB | 22.00 | 0.29 | 7.00 | 0.09 | 5.00 | 0.07 | 2.00 | 0.03 |

Note: pp=percentage points.

we plotted the mean against the variance (see Figures S1 and S2). The results show that the variance consistently exceeds the mean, indicating marked overdispersion, particularly for genes with higher mean expression.

Subsequently, we examined the underlying sources of low counts in spatial single-cell data to assess which modeling strategies best account for the high prevalence of zeros. We analyzed the data using both Poisson and NB (with gene-wise dispersion) distributions to estimate the expected probability of zeros and compared these estimates with the observed zero frequencies (see Figures S3 and S4). For most genes and cell types, the observed percentage of zeros is higher than that predicted by the Poisson model yet closely aligns with the predictions of the NB model.

For a quantitative assessment, we applied the Poisson model, the NB model with common dispersion, the NB model with gene-wise dispersion, and the ZINB model. For each dataset, we computed the number and percentage of genes that deviate from the expected fraction of zero counts by more than 1, 5, 10, and 20 percentage points. Our findings reveal that both the Poisson model and the NB model with common dispersion deviate considerably from the empirical data, whereas the NB model with gene-wise dispersion aligns closely with the observed values. Moreover, the ZINB model performs nearly identically to the NB model with gene-specific dispersion, indicating that once gene-level dispersion is modeled, additional zero-inflation offers little further improvement.

In summary, our analysis demonstrates that the NB model, particularly when implemented with gene-wise dispersion, more accurately captures the variability and zero-count distributions observed in SMURF-based spatial single-cell datasets. As a result, algorithms and pipelines originally developed for single-cell RNA-seq can be effectively applied to these spatial single-cell data.

### Normalization artifact in the 8 $\mu$m method

We noticed an artifact specific to the 8 $\mu$m method. Because of the way this method normalizes count data, if the data contains spatial regions where few transcripts are captured, there is a severe data imbalance that skews the expression values for some genes. For example, Slc1a2 is an important gene primarily expressed in the CA3 subregion of the mouse hippocampus [12, 13]. However, 8 $\mu$m normalization of count data for this gene is discordant with ISH data from the Allen Brain Atlas. In contrast, the single-cell data generated by SMURF is much more concordant with the ISH data (main text Extended Data Figure 1a). This improvement arises because SMURF reallocates transcripts to individual cells based on spatial continuity. This corrects the undersampling artifacts that plague sparsely populated regions.

### Identification of Highly Zonated Genes

For each zone (1–6), we first restricted candidates to genes detected in more than 100 cells within the corresponding dataset. We then used Scanpy differential-expression results to call zone-enriched genes under zone-specific thresholds on $\log_2$ fold-change (LFC) and adjusted $p$-value. To ensure high specificity, we excluded genes significantly enriched in other epithelial zones and in non-epithelial compartments (smooth muscle, immune, fibroblasts, endothelial), as well as Goblet, Tuft, Peyer's patch, and M cell-related set, using the exact criteria listed below.

***Zone 1:***

Selection: LFC $> 1.8$ and adjusted $p < 10^{-40}$. Exclusions: genes enriched in Zone 2 (LFC $> 3$, $p < 10^{-50}$), Zone 3 (LFC $> 0$, $p < 10^{-50}$), or Zone 4 (LFC $> 0$, $p < 10^{-50}$); in smooth muscle (LFC $> 0$, $p < 10^{-30}$), immune (LFC $> 0$, $p < 10^{-30}$), Goblet (LFC $> 0$, $p < 10^{-30}$), Peyer's patch (LFC $> 0$, $p < 10^{-30}$), M cells-related (LFC $> 0$, $p < 10^{-30}$), Tuft (LFC $> 0$, $p < 10^{-30}$), Endothelial (LFC $> 0$, $p < 10^{-30}$), or Fibroblasts (LFC $> 2.8$, $p < 10^{-40}$).

***Zone 2:***

Selection: LFC $> 2.0$ and adjusted $p < 10^{-50}$. Exclusions: genes enriched in Zone 1 (LFC $> 0$, $p < 10^{-100}$), and in smooth muscle, immune, Goblet, Peyer's patch, Enterocyte + M cells, Tuft, Endothelial, or Fibroblasts (all with LFC $> 0$, $p < 10^{-100}$).

***Zone 3:***
Selection: LFC $> 1.4$ and adjusted $p < 10^{-30}$. Exclusions: genes enriched in Zone 2, Zone 6, or Zone 5 (all with LFC $> 0$, $p < 10^{-100}$).
***Zone 4:***
Selection: LFC $> 1.5$ and adjusted $p < 10^{-10}$. Exclusions: genes enriched in Zone 6 or Zone 3 (both with LFC $> 0$, $p < 10^{-100}$).
***Zone 5:***
Selection: LFC $> 1.5$ and adjusted $p < 10^{-10}$. Exclusions: genes enriched in Zone 6, Zone 3, or Zone 4 (all with LFC $> 0$, $p < 10^{-100}$), and in the immune dataset (LFC $> 0$, $p < 10^{-100}$).
***Zone 6:***
Selection: LFC $> 2.0$ and adjusted $p < 10^{-20}$. Exclusions: genes enriched in Zone 4, Zone 3, or Zone 5 (all with LFC $> 0$, $p < 10^{-50}$), and in the immune dataset (LFC $> 0$, $p < 10^{-50}$).

### Transcription Factor related Results

Here we show the results from SCENIC analysis [14]. For each zone, we calculated average TF-regulon scores and kept regulons with average activity scores exceeding 0.05.

| TF | 1 | 2 | 3 | 4 | 5 | 6 | TF | 1 | 2 | 3 | 4 | 5 | 6 |
|---|---|---|---|---|---|---|---|---|---|---|---|---|---|
| Ahctf1(+) | 0.054 | 0.080 | 0.077 | 0.060 | 0.053 | 0.053 | Atf3(+) | 0.040 | 0.052 | 0.073 | 0.089 | 0.117 | 0.163 |
| Atf7(+) | 0.038 | 0.054 | 0.043 | 0.035 | 0.033 | 0.032 | Bach1(+) | 0.034 | 0.048 | 0.067 | 0.084 | 0.094 | 0.102 |
| Bclaf1(+) | 0.053 | 0.065 | 0.051 | 0.041 | 0.037 | 0.036 | Bptf(+) | 0.027 | 0.051 | 0.050 | 0.038 | 0.035 | 0.035 |
| Cdx2(+) | 0.040 | 0.054 | 0.054 | 0.047 | 0.042 | 0.041 | Cebpa(+) | 0.066 | 0.078 | 0.077 | 0.068 | 0.062 | 0.057 |
| Cebpb(+) | 0.026 | 0.047 | 0.052 | 0.053 | 0.053 | 0.052 | Cebpd(+) | 0.072 | 0.079 | 0.088 | 0.089 | 0.091 | 0.091 |
| Cebpg(+) | 0.045 | 0.048 | 0.053 | 0.055 | 0.054 | 0.054 | Chd2(+) | 0.065 | 0.069 | 0.068 | 0.066 | 0.065 | 0.064 |
| Churc1(+) | 0.081 | 0.093 | 0.102 | 0.086 | 0.070 | 0.068 | Cnot3(+) | 0.035 | 0.054 | 0.047 | 0.035 | 0.030 | 0.030 |
| Creb1(+) | 0.060 | 0.071 | 0.059 | 0.050 | 0.048 | 0.048 | Creb3l1(+) | 0.044 | 0.058 | 0.043 | 0.033 | 0.031 | 0.030 |
| Crem(+) | 0.035 | 0.062 | 0.051 | 0.032 | 0.028 | 0.026 | Cux1(+) | 0.095 | 0.075 | 0.085 | 0.092 | 0.095 | 0.098 |
| E2f8(+) | 0.058 | 0.079 | 0.063 | 0.046 | 0.042 | 0.040 | Ehf(+) | 0.035 | 0.065 | 0.056 | 0.036 | 0.024 | 0.019 |
| Elf1(+) | 0.048 | 0.067 | 0.071 | 0.070 | 0.066 | 0.060 | Elf2(+) | 0.044 | 0.057 | 0.050 | 0.044 | 0.044 | 0.049 |
| Elf3(+) | 0.049 | 0.068 | 0.059 | 0.046 | 0.041 | 0.040 | Elk4(+) | 0.050 | 0.068 | 0.075 | 0.072 | 0.072 | 0.075 |
| Ep300(+) | 0.043 | 0.066 | 0.081 | 0.082 | 0.078 | 0.074 | Esrra(+) | 0.048 | 0.071 | 0.095 | 0.099 | 0.093 | 0.085 |
| Ets1(+) | 0.043 | 0.048 | 0.063 | 0.075 | 0.092 | 0.105 | Fos(+) | 0.021 | 0.029 | 0.056 | 0.108 | 0.142 | 0.167 |
| Fosl2(+) | 0.039 | 0.044 | 0.060 | 0.075 | 0.100 | 0.127 | Foxf1(+) | 0.116 | 0.095 | 0.117 | 0.142 | 0.148 | 0.138 |
| Foxn2(+) | 0.129 | 0.118 | 0.111 | 0.116 | 0.116 | 0.116 | Foxo1(+) | 0.050 | 0.044 | 0.045 | 0.046 | 0.047 | 0.051 |
| Foxo4(+) | 0.051 | 0.055 | 0.054 | 0.051 | 0.052 | 0.051 | Gabpa(+) | 0.046 | 0.063 | 0.064 | 0.054 | 0.049 | 0.050 |
| Gata5(+) | 0.056 | 0.076 | 0.072 | 0.055 | 0.046 | 0.040 | Gata6(+) | 0.052 | 0.056 | 0.053 | 0.050 | 0.050 | 0.053 |
| Gatad1(+) | 0.036 | 0.054 | 0.056 | 0.045 | 0.038 | 0.038 | Hes1(+) | 0.043 | 0.059 | 0.045 | 0.032 | 0.026 | 0.024 |
| Hic1(+) | 0.041 | 0.054 | 0.080 | 0.104 | 0.111 | 0.094 | Hmg20b(+) | 0.035 | 0.058 | 0.065 | 0.070 | 0.074 | 0.071 |
| Hnf1a(+) | 0.043 | 0.063 | 0.050 | 0.035 | 0.031 | 0.030 | Hnf4a(+) | 0.044 | 0.064 | 0.069 | 0.062 | 0.062 | 0.066 |
| Hnf4g(+) | 0.044 | 0.066 | 0.094 | 0.105 | 0.102 | 0.095 | Id2(+) | 0.048 | 0.062 | 0.060 | 0.055 | 0.053 | 0.049 |
| Irf6(+) | 0.059 | 0.076 | 0.085 | 0.082 | 0.079 | 0.080 | Irf7(+) | 0.033 | 0.048 | 0.078 | 0.109 | 0.136 | 0.155 |
| Irf8(+) | 0.049 | 0.057 | 0.059 | 0.053 | 0.049 | 0.046 | Jun(+) | 0.042 | 0.052 | 0.069 | 0.087 | 0.118 | 0.160 |
| Junb(+) | 0.058 | 0.075 | 0.122 | 0.163 | 0.180 | 0.184 | Jund(+) | 0.045 | 0.070 | 0.080 | 0.074 | 0.072 | 0.072 |
| Klf3(+) | 0.053 | 0.067 | 0.089 | 0.097 | 0.101 | 0.107 | Klf4(+) | 0.044 | 0.053 | 0.087 | 0.126 | 0.161 | 0.188 |
| Klf5(+) | 0.094 | 0.198 | 0.196 | 0.115 | 0.070 | 0.056 | Maf(+) | 0.019 | 0.058 | 0.159 | 0.268 | 0.273 | 0.222 |
| Mafb(+) | 0.047 | 0.046 | 0.072 | 0.108 | 0.123 | 0.122 | Mafg(+) | 0.038 | 0.053 | 0.049 | 0.042 | 0.041 | 0.042 |
| Max(+) | 0.038 | 0.050 | 0.077 | 0.101 | 0.123 | 0.142 | Mbd2(+) | 0.046 | 0.065 | 0.055 | 0.043 | 0.041 | 0.041 |
| Mlx(+) | 0.035 | 0.045 | 0.059 | 0.061 | 0.056 | 0.049 | Mlxipl(+) | 0.087 | 0.171 | 0.165 | 0.112 | 0.085 | 0.066 |
| Mxd1(+) | 0.043 | 0.068 | 0.105 | 0.138 | 0.158 | 0.168 | Mxd3(+) | 0.073 | 0.081 | 0.068 | 0.060 | 0.060 | 0.064 |
| Myb(+) | 0.120 | 0.122 | 0.090 | 0.087 | 0.087 | 0.089 | Myc(+) | 0.063 | 0.080 | 0.059 | 0.043 | 0.039 | 0.038 |
| Ncoa2(+) | 0.144 | 0.114 | 0.122 | 0.138 | 0.140 | 0.139 | Neurod1(+) | 0.032 | 0.053 | 0.086 | 0.109 | 0.150 | 0.204 |
| Nfya(+) | 0.064 | 0.073 | 0.059 | 0.050 | 0.048 | 0.047 | Nfyc(+) | 0.058 | 0.071 | 0.073 | 0.062 | 0.057 | 0.055 |
| Nkx2-3(+) | 0.033 | 0.053 | 0.040 | 0.031 | 0.034 | 0.040 | Nr1h4(+) | 0.040 | 0.054 | 0.063 | 0.058 | 0.052 | 0.048 |
| Nr1i2(+) | 0.040 | 0.059 | 0.078 | 0.096 | 0.092 | 0.084 | Nr1i3(+) | 0.023 | 0.049 | 0.061 | 0.058 | 0.050 | 0.041 |
| Nr2f2(+) | 0.066 | 0.079 | 0.094 | 0.095 | 0.093 | 0.097 | Ovol1(+) | 0.109 | 0.107 | 0.104 | 0.100 | 0.105 | 0.113 |
| Ovol2(+) | 0.040 | 0.059 | 0.054 | 0.038 | 0.041 | 0.049 | Pbx1(+) | 0.025 | 0.049 | 0.065 | 0.060 | 0.051 | 0.044 |
| Pdlim5(+) | 0.072 | 0.067 | 0.072 | 0.072 | 0.069 | 0.069 | Plagl2(+) | 0.106 | 0.085 | 0.087 | 0.089 | 0.092 | 0.088 |
| Ppard(+) | 0.033 | 0.061 | 0.064 | 0.055 | 0.045 | 0.036 | Pura(+) | 0.049 | 0.065 | 0.062 | 0.055 | 0.054 | 0.052 |
| Rad21(+) | 0.050 | 0.051 | 0.043 | 0.038 | 0.038 | 0.038 | Rfx5(+) | 0.057 | 0.067 | 0.066 | 0.058 | 0.057 | 0.055 |
| Rxra(+) | 0.052 | 0.066 | 0.082 | 0.085 | 0.091 | 0.101 | Sirt6(+) | 0.024 | 0.038 | 0.054 | 0.064 | 0.063 | 0.055 |
| Smad1(+) | 0.044 | 0.064 | 0.053 | 0.042 | 0.037 | 0.034 | Smarcc2(+) | 0.089 | 0.091 | 0.092 | 0.090 | 0.090 | 0.087 |
| Sox13(+) | 0.107 | 0.084 | 0.091 | 0.100 | 0.104 | 0.105 | Sp1(+) | 0.044 | 0.063 | 0.049 | 0.039 | 0.035 | 0.034 |
| Sp3(+) | 0.074 | 0.065 | 0.068 | 0.071 | 0.072 | 0.073 | Spdef(+) | 0.062 | 0.057 | 0.050 | 0.044 | 0.039 | 0.036 |
| Srebf2(+) | 0.054 | 0.076 | 0.077 | 0.065 | 0.058 | 0.055 | Srf(+) | 0.064 | 0.056 | 0.085 | 0.113 | 0.119 | 0.121 |
| Stat1(+) | 0.051 | 0.063 | 0.078 | 0.084 | 0.081 | 0.071 | Stat2(+) | 0.041 | 0.056 | 0.082 | 0.108 | 0.127 | 0.138 |
| Stat6(+) | 0.055 | 0.078 | 0.082 | 0.074 | 0.069 | 0.069 | Tbl1xr1(+) | 0.212 | 0.163 | 0.170 | 0.185 | 0.193 | 0.198 |
| Tcf7l2(+) | 0.044 | 0.053 | 0.061 | 0.063 | 0.061 | 0.062 | Tead1(+) | 0.068 | 0.059 | 0.058 | 0.058 | 0.057 | 0.058 |
| Tfdp1(+) | 0.050 | 0.068 | 0.053 | 0.039 | 0.035 | 0.034 | Tfeb(+) | 0.025 | 0.051 | 0.053 | 0.041 | 0.036 | 0.036 |
| Thrb(+) | 0.056 | 0.067 | 0.062 | 0.054 | 0.051 | 0.051 | Trp53(+) | 0.057 | 0.066 | 0.051 | 0.038 | 0.037 | 0.038 |
| Usf1(+) | 0.059 | 0.069 | 0.065 | 0.060 | 0.056 | 0.053 | Usf2(+) | 0.053 | 0.063 | 0.063 | 0.061 | 0.061 | 0.061 |
| Xbp1(+) | 0.045 | 0.067 | 0.065 | 0.054 | 0.047 | 0.043 | Ybx1(+) | 0.066 | 0.081 | 0.076 | 0.065 | 0.059 | 0.056 |
| Yy1(+) | 0.044 | 0.053 | 0.050 | 0.048 | 0.052 | 0.057 | Zbtb14(+) | 0.059 | 0.064 | 0.054 | 0.047 | 0.046 | 0.045 |
| Zbtb21(+) | 0.085 | 0.074 | 0.089 | 0.091 | 0.091 | 0.089 | Zbtb33(+) | 0.029 | 0.055 | 0.055 | 0.047 | 0.038 | 0.028 |
| Zbtb41(+) | 0.061 | 0.072 | 0.052 | 0.040 | 0.038 | 0.035 | Zbtb7a(+) | 0.063 | 0.066 | 0.069 | 0.068 | 0.066 | 0.063 |
| Zbtb7b(+) | 0.041 | 0.062 | 0.070 | 0.067 | 0.066 | 0.067 | Zfp467(+) | 0.067 | 0.161 | 0.193 | 0.202 | 0.252 | 0.291 |
| Zfp992(+) | 0.087 | 0.099 | 0.119 | 0.131 | 0.128 | 0.121 | Zfx(+) | 0.047 | 0.062 | 0.063 | 0.056 | 0.052 | 0.051 |

### Lite (Non-GPU) version of SMURF and example results

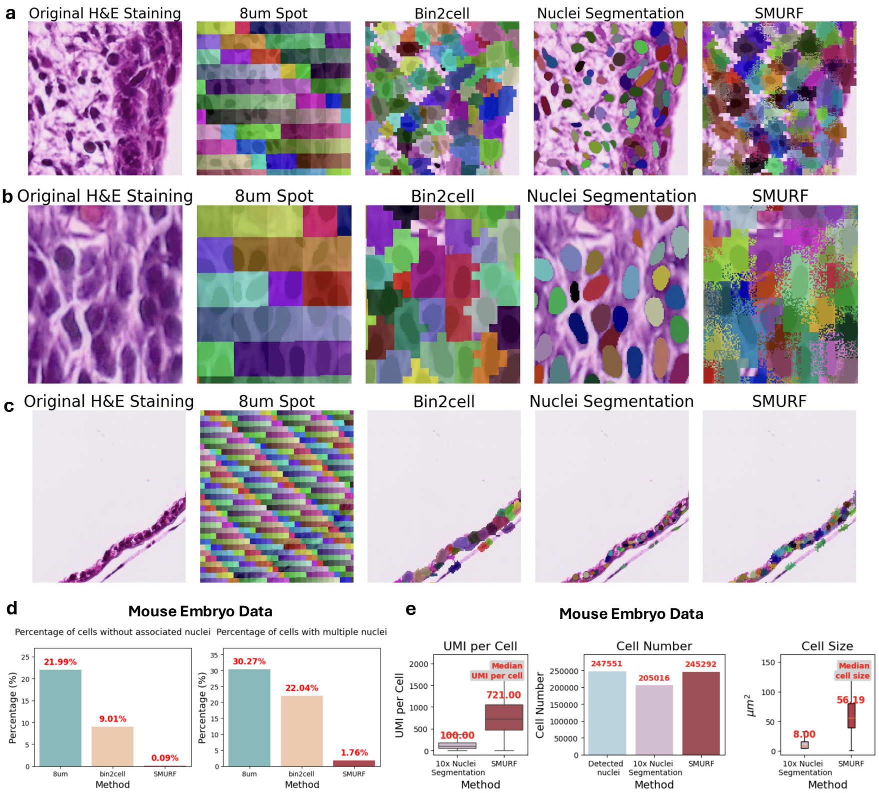


**Fig. S5**: Mouse Embryo Dataset

The deconvolution module of SMURF relies on a PyTorch-based deep learning implementation that demanded substantial GPU resources. To offer a more accessible option for all users, we introduce a Lite version which does not require access to GPUs. The lite version streamlines the algorithm by deconvolving shared spots only using information on the spot's distance from the center of its corresponding nucleus. We applied this lite version to a public mouse embryo dataset (H&E staining) [15] and human tonsil dataset (IF staining) [16] to demonstrate its effectiveness.

**Fig. S6**: Human Tonsil Example

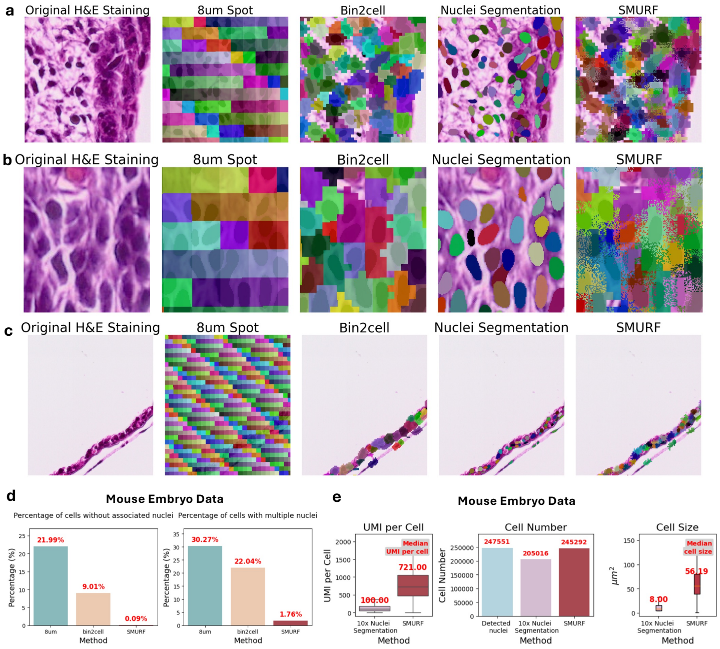


For the mouse embryo example (Supplementary Figure S5) and the human tonsil example (Supplementary Figure S6), we visualized how SMURF-lite assigned capture spots to nuclei in three distinct sections, and compared those results to assigned obtained using 8 $\mu$m method, Bin2cell, and nuclei segmentation methods. Following the format shown in Figure 2a-b of the main text, we presented H&E/IF-stained sections alongside capture spot-to-nuclei assignments from the 8 $\mu$m method, Bin2cell, and SMURF-lite for both the mouse embryo (Supplementary Figure S5a-c) and human tonsil (Supplementary Figure S6a-c), each in a separate randomly selected section. SMURF-lite reliably achieves single-cell resolution by assigning transcripts from overlapping and non-overlapping capture spots to cells, outperforming the 8 $\mu$m method, Bin2cell, and nuclei segmentation, especially in densely populated regions.

For quantitative analysis, we employed the same procedure as in the main text (Figure 2c) to compare the 8 $\mu$m method, Bin2cell, and SMURF-lite based on the percentages of cells lacking an associated nucleus and those containing multiple nuclei (Supplementary Figure S5d and Supplementary Figure S6d). In the mouse embryo data, only 0.15% of SMURF-lite-defined cells lacked a nucleus, whereas this figure was approximately 146.6-fold higher (21.99%) and 94.8-fold higher (14.22%) when using Bin2cell and the 8 $\mu$m method, respectively. Likewise, 2.75% of SMURF-lite-defined cells contained multiple nuclei, contrasting with about 11.0-fold (30.27%) and 20.2-fold (55.37%) higher rates for Bin2cell and the 8 $\mu$m method. In the human tonsil data, only 0.76% of SMURF-lite-defined cells were without a nucleus, roughly

15.2-fold (11.52%) and 21.8-fold (16.53%) higher in Bin2cell and the 8 $\mu$m method, respectively, while 3.80% of SMURF-lite-defined cells included multiple nuclei, about 12.7-fold (48.30%) and 14.4-fold (54.66%) higher in Bin2cell and the 8 $\mu$m method, respectively. These findings demonstrate that the lite version of SMURF more accurately assigns transcripts to cells at single-cell resolution than the 8 $\mu$m or Bin2cell method.

Concluding that Bin2cell and the 8 $\mu$m method do not reliably achieve single-cell resolution with Visium HD data, we next compared SMURF-lite directly to the nuclei segmentation method as Figure 2e of the main text. Since SMURF-lite includes capture spots that do not overlap nuclei, we hypothesized it would yield larger median cell sizes and higher transcript counts per cell. Indeed, when analyzing both mouse embryo and human tonsil Visium HD data with SMURF-lite or nuclei segmentation (Supplementary Figure S5e and Supplementary Figure S6e, left), SMURF-lite produced a 7.21-fold increase in median UMIs per cell (721 vs. 100) and a 7.02-fold increase in median cell area (56.19 $\mu m^2$ vs. 8 $\mu m^2$) for the mouse embryo data. For the human tonsil data, SMURF-lite provided a 1.85-fold increase in median UMIs per cell (205 vs. 111) and a 2-fold increase in median cell area (48 $\mu m^2$ vs. 24 $\mu m^2$). Both methods identified comparable numbers of cells (Supplementary Figure S5e and Supplementary Figure S6e, middle), consistent with their shared reliance on segmented nuclei as the basis for defining cells. Collectively, these data indicate that SMURF-lite assigns substantially more transcripts to each cell than the nuclei segmentation method by incorporating capture spots beneath cell bodies, thus leading to larger overall cell sizes.

In total, SMURF-lite surpasses Bin2cell, the 8 $\mu$m method, and nuclei segmentation by more accurately assigning transcripts to cells at single-cell resolution, thereby offering reliable spatial single-cell results through SMURF for researchers lacking GPU resources.

[12] Ke Cheng, Chong Chen, Qinji Zhou, Xiangyu Chen, and Peng Xie. Deficit of neuronal eaat2 impairs hippocampus ca3 neuron's activity and may induce depressive like behaviors. *Journal of Advanced Research*, 2025.

[13] Allen Institute for Brain Science. Allen mouse brain atlas [dataset]. http://mouse.brain-map.org, 2004. Accessed: 2025-03-24.

[14] Sara Aibar, Carlos B. González-Blas, Thomas Moerman, Vân A. Huynh-Thu, Hana Imrichova, Geert Hulselmans, Florian Rambow, José C. Marine, Pierre Geurts, and Stein Aerts. SCENIC: single-cell regulatory network inference and clustering. *Nature Methods*, 14(11):1083–1086, 2017.

[15] 10x Genomics. Visium hd cytassist gene expression libraries of mouse embryo. https://www.10xgenomics.com/datasets/visium-hd-cytassist-gene-expression-libraries-of-mouse-embryo. Accessed: 28 February 2025.

[16] 10x Genomics. Visium hd human tonsil fresh frozen if. https://cf.10xgenomics.com/samples/spatial-exp/3.1.1/Visium_HD_Human_Tonsil_Fresh_Frozen_IF/Visium_HD_Human_Tonsil_Fresh_Frozen_IF_web_summary.html. Dataset version 3.1.1. Accessed: 28 February 2025.
